# A systematic evaluation of 41 DNA methylation predictors across 101 data preprocessing and normalization strategies highlights considerable variation in algorithm performance

**DOI:** 10.1101/2021.09.29.462387

**Authors:** Anil P.S. Ori, Ake T Lu, Steve Horvath, Roel A Ophoff

## Abstract

**Background:** DNA methylation (DNAm) based predictors hold great promise to serve as clinical tools for health interventions and disease management. While these algorithms often have high prediction accuracy and are associated with many disease-related phenotypes, the reliability of their performance remains to be determined. We therefore conducted a systematic evaluation across 101 different data processing strategies that preprocess and normalize DNAm data and assessed how each analytical strategy affects the reliability and prediction accuracy of 41 DNAm-based predictors.

**Results:** Our analyses were conducted in a large EPIC DNAm sample of the Jackson Heart Study (N=2,053) that included 146 pairs of technical replicate samples. By estimating the average absolute agreement between replicate pairs, we show that 32 out of 41 predictors (78%) demonstrate excellent test-retest reliability when appropriate data processing and normalization steps are implemented. Across all pairs of predictors, we find a moderate correlation in performance across analytical strategies (mean rho=0.40, SD=0.27), highlighting significant heterogeneity in performance across algorithms within a choice of an analytical pipeline. (Un)successful removal of technical variation furthermore significantly impacts downstream phenotypic association analysis, such as all-cause mortality risk associations.

**Conclusions:** We show that DNAm-based algorithms are sensitive to technical variation. The right choice of data processing and normalization pipeline is important to achieve reproducible estimates and improve prediction accuracy in downstream phenotypic association analyses. For each of the 41 DNAm predictors, we report its test-retest reliability and provide the best performing analytical strategy as a guideline for the research community. As DNAm-based predictors become more and more widely used, both for research purposes as well as for clinic applications, our work helps improve their performance and standardize their implementation.

## Introduction

DNA methylation (DNAm) is a form of epigenetic regulation that is essential for human development and implicated in health and disease[1,2]. Through advancements in biological technology, large-scale DNA methylation profiling has become more affordable and widely used. Microarray technologies now enable the simultaneous interrogation of DNAm states of more than 850,000 CpG dinucleotides across the genome, using the latest EPIC array[3]. An application of DNAm data has been in developing DNAm-based algorithms to predict health-related phenotypes, including blood cell type proportions[4,5], ageing[6–13], all-cause mortality risk[14–17], cancer risk[18,19], body-mass-index (BMI), and smoking signatures[20], among others. These molecular predictors have great potential for clinical applications. A thorough and systematic investigation of their performance has however not been conducted so far.

Unlike the genome, the DNA methylome is of dynamic nature and largely explained by non-shared individual environments[21]. Like other high-throughput molecular data, DNAm can furthermore be impacted by variation in laboratory conditions, sample handling, reagents and/or equipment used[22]. Technical variation is often widespread and tackling such effects is of critical importance to study biological variation in any -omic analysis, including DNAm. Over the years, a plethora of methods have been developed to identify and remove unwanted technical variations from DNAm data[23–29]. Previous studies have investigated the impact of specific methods on outcomes of DNAm analysis and demonstrated the importance of correcting for probe design type, batch effects, and hidden confounders while the effect of different normalization strategies gave mixed results[30–33]. A systematic and unbiased evaluation of commonly used data preprocessing and normalization strategies of DNAm data for the application of DNAm-based predictors has however not yet been conducted. DNAm is an important tool to study health and disease and understanding how analytical strategies impact algorithm performance is critical for method standardization and implementation for both research and clinical purposes.

Here, we performed a comprehensive investigation of 41 DNAm predictors and evaluated algorithm performance by measuring their test-retest reliability across 101 data preprocessing and normalization strategies in the Jackson Heart Study (JHS)[34]. The JHS has collected a large sample of 850K EPIC DNAm arrays in blood that includes 146 pairs of technical replicates. These replicates represent identical DNA samples that were assayed twice at independent time points. The agreement in DNAm predictor estimate between technical replicates after data preprocessing and normalization allowed us to quantify the degree to which an analytical strategy can successfully remove unwanted technical variation. We report the best test-retest reliability for each predictor and demonstrate how reducing technical variation is critical for optimal algorithm performance in downstream phenotypic analyses. Our work emphasizes the importance of data processing and normalization of DNAm data and provides best practices to optimize the performance and reliability of DNAm predictors.

## Results

To evaluate how unwanted technical variation in DNAm data impacts the performance of DNAm-based predictors, we implemented 101 data processing and normalization strategies in the JHS dataset. For each analytical strategy, which we will refer to as a “pipeline”, we then extracted beta values and calculated estimates of 41 DNAm-based predictors in (1) JHS data1: a sample of 146 technical replicate pairs and (2) JHS data 2: a general sample of 1,761 non-replicate samples that do not overlap with the individuals in the replicate dataset. Figure 1 shows an overview of our analysis plan. In the sample of technical replicates, we quantified the average absolute agreement between replicate pair values (i.e. reliability) by means of the ICC for each DNAm predictor and each pipeline separately (41 predictors × 101 pipelines = 4,141 ICC analysis). We also generated DNAm estimates in the general sample. This allowed us to correlate the ICC of a pipeline that was estimated in the sample of replicates with predictor estimates in the independent general JHS sample.

**Figure 1.**
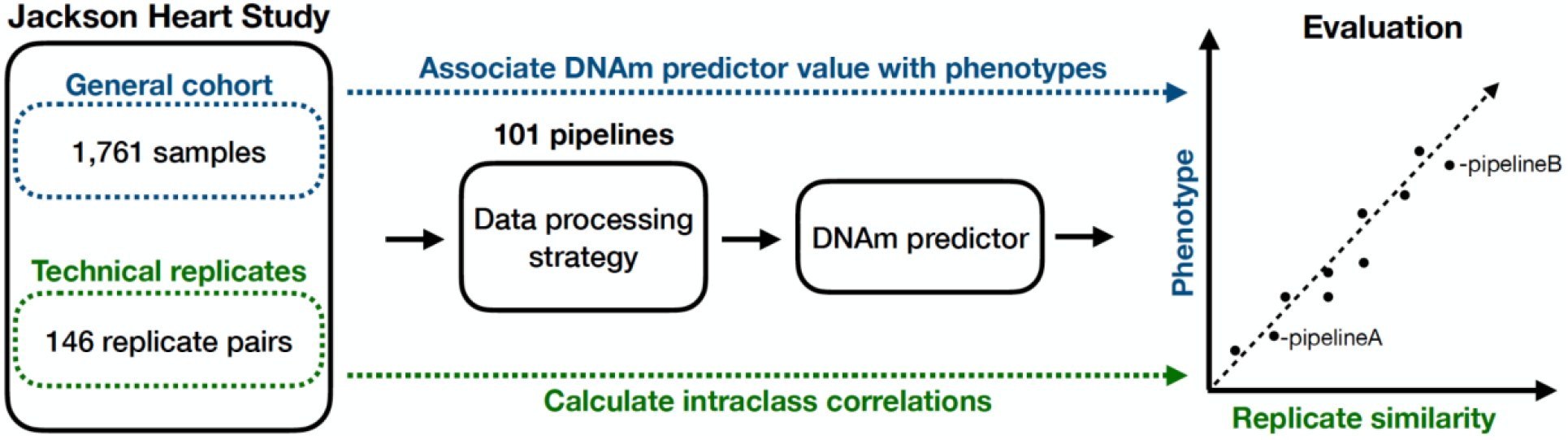
Schematic overview of analysis plan to evaluate DNAm algorithm performance. DNAm analyses are conducted using DNAm EPIC array samples in JHS. JHS includes a significant number of technical replicate pairs thereby allowing for a careful investigation of how the removal of unwanted technical variation impacts DNAm algorithm performance across 101 data processing pipelines. JHS has also collected information on disease-related phenotypes, including mortality status after follow-up. This allowed us to assess how removal of technical variation in DNAm predictor estimates by a data processing pipeline impacts downstream phenotypic association analyses.

We calculated the ICC estimates derived from a two-way random effect model to assess the reliability of each predictor for each data processing pipelines. The ICC is a zero to one estimate that quantifies the average absolute agreement across technical replicate pairs that were processed at a different occasion. We also calculated five other types of ICCs and found high concordance between the different ICC measures (mean rho=0.99, SD=0.01, see Figure S1). Table S5 reports all ICC statistics for each DNAm predictor and pipeline. In the remainder of the paper we will refer to ICC(2,1) as ICC, unless stated otherwise.

### Most DNAm-based predictors yield high reliability when the best analytical pipeline is implemented

Table 1 shows all 41 DNAm predictors alongside general information on each algorithm and corresponding ICC statistics, including the data processing and normalization pipeline that yielded the highest reliability for each predictor. Across all predictors and pipelines (N=4,141), we observed a significant degree of similarity between replicates (all ICC P-values < 0.05/4,141). The median across all ICC estimates is 0.93 with a range of 0.22-0.99.

**Table 1.**
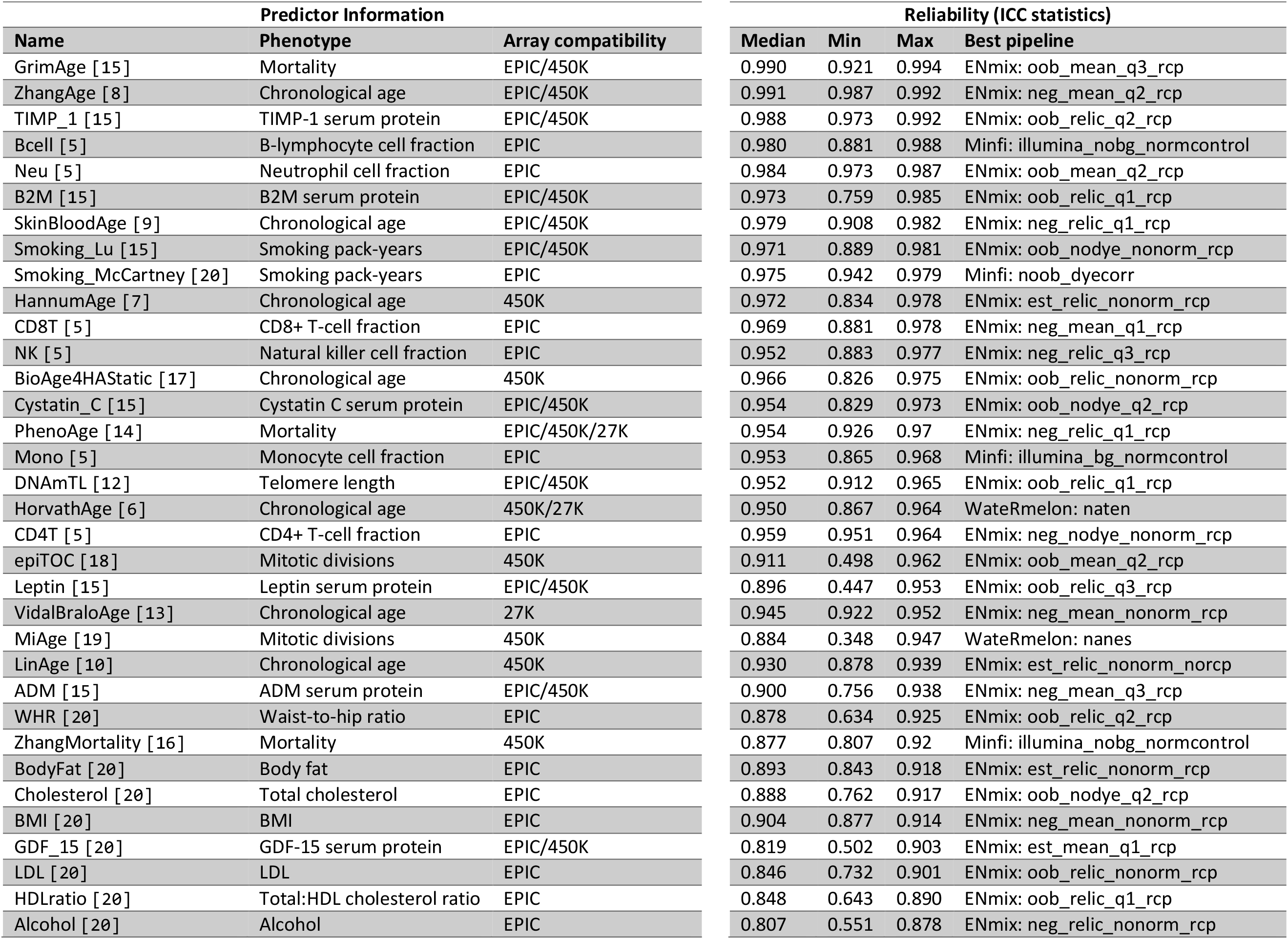
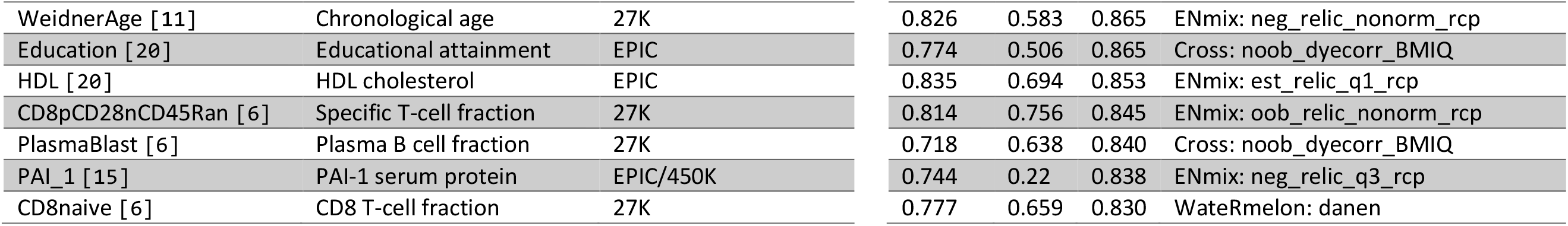
Overview of predictor reliability and best performing data processing pipelines. Shown are general information on each DNAm-based predictor alongside their corresponding ICC statistics from the reliability analysis. The name of the predictor, the phenotype it is trained on, and DNAm array compatibility are listed on the left side of the table. ICC statistics are listed on the right side of the table. For each predictor, across 101 pipelines, the median, minimum, and maximum ICC are listed. Predictors are ranked by the maximum ICC. The final column reports the best performing data processing pipelines (i.e., the pipeline with the highest reliability).

| Predictor Information |  |  |
| --- | --- | --- |
| Name | Phenotype | Array compatibility |
| GrimAge [15] | Mortality | EPIC/450K |
| ZhangAge [8] | Chronological age | EPIC/450K |
| TIMP_1 [15] | TIMP-1 serum protein | EPIC/450K |
| Bcell [5] | B-lymphocyte cell fraction | EPIC |
| Neu [5] | Neutrophil cell fraction | EPIC |
| B2M [15] | B2M serum protein | EPIC/450K |
| SkinBloodAge [9] | Chronological age | EPIC/450K |
| Smoking_Lu [15] | Smoking pack-years | EPIC/450K |
| Smoking_McCartney [20] | Smoking pack-years | EPIC |
| HannumAge [7] | Chronological age | 450K |
| CD8T [5] | CD8+ T-cell fraction | EPIC |
| NK [5] | Natural killer cell fraction | EPIC |
| BioAge4HAStatic [17] | Chronological age | 450K |
| Cystatin_C [15] | Cystatin C serum protein | EPIC/450K |
| PhenoAge [14] | Mortality | EPIC/450K/27K |
| Mono [5] | Monocyte cell fraction | EPIC |
| DNAmTL [12] | Telomere length | EPIC/450K |
| HorvathAge [6] | Chronological age | 450K/27K |
| CD4T [5] | CD4+ T-cell fraction | EPIC |
| epiTOC [18] | Mitotic divisions | 450K |
| Leptin [15] | Leptin serum protein | EPIC/450K |
| VidalBraloAge [13] | Chronological age | 27K |
| MiAge [19] | Mitotic divisions | 450K |
| LinAge [10] | Chronological age | 450K |
| ADM [15] | ADM serum protein | EPIC/450K |
| WHR [20] | Waist-to-hip ratio | EPIC |
| ZhangMortality [16] | Mortality | 450K |
| BodyFat [20] | Body fat | EPIC |
| Cholesterol [20] | Total cholesterol | EPIC |
| BMI [20] | BMI | EPIC |
| GDF_15 [20] | GDF-15 serum protein | EPIC/450K |
| LDL [20] | LDL | EPIC |
| HDLratio [20] | Total:HDL cholesterol ratio | EPIC |
| Alcohol [20] | Alcohol | EPIC |

| Reliability (ICC statistics) |  |  |  |
| --- | --- | --- | --- |
| Median | Min | Max | Best pipeline |
| 0.990 | 0.921 | 0.994 | ENmix: oob_mean_q3_rcp |
| 0.991 | 0.987 | 0.992 | ENmix: neg_mean_q2_rcp |
| 0.988 | 0.973 | 0.992 | ENmix: oob_relic_q2_rcp |
| 0.980 | 0.881 | 0.988 | Minfi: illumina_nobg_normcontrol |
| 0.984 | 0.973 | 0.987 | ENmix: oob_mean_q2_rcp |
| 0.973 | 0.759 | 0.985 | ENmix: oob_relic_q1_rcp |
| 0.979 | 0.908 | 0.982 | ENmix: neg_relic_q1_rcp |
| 0.971 | 0.889 | 0.981 | ENmix: oob_nodye_nonorm_rcp |
| 0.975 | 0.942 | 0.979 | Minfi: noob_dyecorr |
| 0.972 | 0.834 | 0.978 | ENmix: est_relic_nonorm_rcp |
| 0.969 | 0.881 | 0.978 | ENmix: neg_mean_q1_rcp |
| 0.952 | 0.883 | 0.977 | ENmix: neg_relic_q3_rcp |
| 0.966 | 0.826 | 0.975 | ENmix: oob_relic_nonorm_rcp |
| 0.954 | 0.829 | 0.973 | ENmix: oob_nodye_q2_rcp |
| 0.954 | 0.926 | 0.97 | ENmix: neg_relic_q1_rcp |
| 0.953 | 0.865 | 0.968 | Minfi: illumina_bg_normcontrol |
| 0.952 | 0.912 | 0.965 | ENmix: oob_relic_q1_rcp |
| 0.950 | 0.867 | 0.964 | Watermelon: naten |
| 0.959 | 0.951 | 0.964 | ENmix: neg_nodye_nonorm_rcp |
| 0.911 | 0.498 | 0.962 | ENmix: oob_mean_q2_rcp |
| 0.896 | 0.447 | 0.953 | ENmix: oob_relic_q3_rcp |
| 0.945 | 0.922 | 0.952 | ENmix: neg_mean_nonorm_rcp |
| 0.884 | 0.348 | 0.947 | Watermelon: nanes |
| 0.930 | 0.878 | 0.939 | ENmix: est_relic_nonorm_norcp |
| 0.900 | 0.756 | 0.938 | ENmix: neg_mean_q3_rcp |
| 0.878 | 0.634 | 0.925 | ENmix: oob_relic_q2_rcp |
| 0.877 | 0.807 | 0.92 | Minfi: illumina_nobg_normcontrol |
| 0.893 | 0.843 | 0.918 | ENmix: est_relic_nonorm_rcp |
| 0.888 | 0.762 | 0.917 | ENmix: oob_nodye_q2_rcp |
| 0.904 | 0.877 | 0.914 | ENmix: neg_mean_nonorm_rcp |
| 0.819 | 0.502 | 0.903 | ENmix: est_mean_q1_rcp |
| 0.846 | 0.732 | 0.901 | ENmix: oob_relic_nonorm_rcp |
| 0.848 | 0.643 | 0.890 | ENmix: oob_relic_q1_rcp |
| 0.807 | 0.551 | 0.878 | ENmix: neg_relic_nonorm_rcp |

**Table 1. Overview of predictor reliability and best performing data processing pipelines.**
|  |  |  |  |  |  |  |
| --- | --- | --- | --- | --- | --- | --- |
| WeidnerAge [11] | Chronological age | 27K | 0.826 | 0.583 | 0.865 | ENmix: neg_relic_nonorm_rcp |
| Education [20] | Educational attainment | EPIC | 0.774 | 0.506 | 0.865 | Cross: noob_dyecorr_BMIQ |
| HDL [20] | HDL cholesterol | EPIC | 0.835 | 0.694 | 0.853 | ENmix: est_relic_q1_rcp |
| CD8pCD28nCD45Ran [6] | Specific T-cell fraction | 27K | 0.814 | 0.756 | 0.845 | ENmix: oob_relic_nonorm_rcp |
| PlasmaBlast [6] | Plasma B cell fraction | 27K | 0.718 | 0.638 | 0.840 | Cross: noob_dyecorr_BMIQ |
| PAI_1 [15] | PAI-1 serum protein | EPIC/450K | 0.744 | 0.22 | 0.838 | ENmix: neg_relic_q3_rcp |
| CD8naive [6] | CD8 T-cell fraction | 27K | 0.777 | 0.659 | 0.830 | WateRmelon: danen |

The GrimAge predictor reports the highest reliability (ICC=0.994, P=6.6e-144), followed by ZhangAge (ICC=0.992, P=8.4e-132), and TIMP_1 (ICC=0.992, P=8.5e-133). In fact, 32 out of 41 predictors (78%) reach a reliability of an ICC > 0.9 with at least one data processing pipeline. The predictors with higher ICCs have more narrow ICC distributions than predictors with lower ICCs (see Figure 2), suggesting that predictors with higher reliability are more robust to the choise of data processing pipelines. The predictors with the lowest reliability are CD8pCD28nCD45RAn (ICC=0.85, P=1.63e-41), PlasmaBlast (ICC=0.84, P=7.19e-52), PAI-1 (ICC=0.84, P=2.80e-40), and CD8_naive (ICC=0.83, P=1.17e-39).

**Figure 2.**
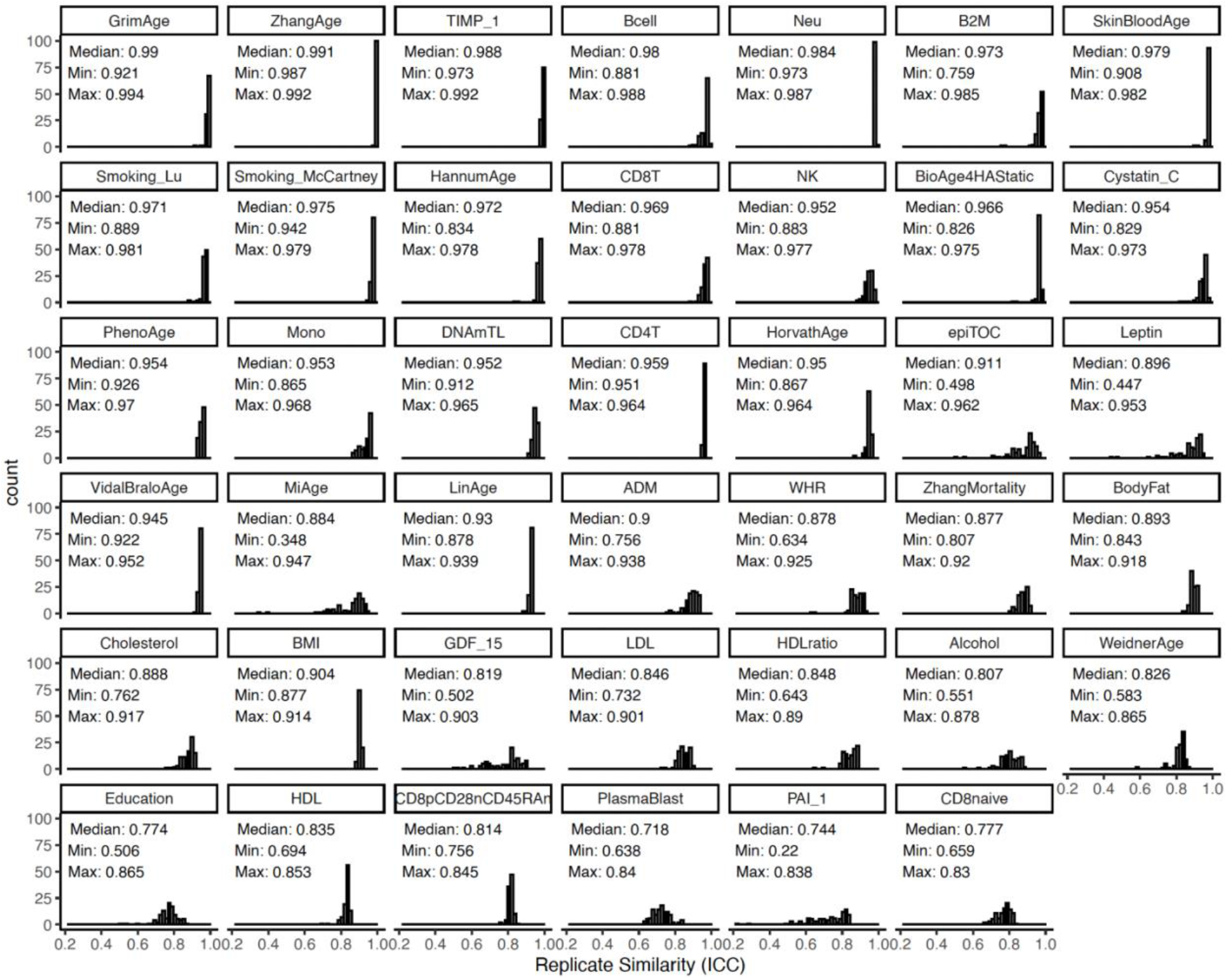
The distribution of intraclass correlations across pipelines for each DNAm algorithm. For each predictor, a histogram of ICC values across 101 pipelines is shown. The ICC quantifies the degree of absolute agreement between estimator values of a pair of technical replicates. The predictors are ranked based on their max ICC value. The name of the predictor is printed on top. In each panel, the median, lowest, and highest ICC value of a corresponding data processing pipeline for that predictor is shown as well.

Across pipelines and predictors (N=4,141), the ENmix package yielded higher reliability (median ICC=0.93, range=0.61-0.99) than the minfi (median ICC=0.91, range=0.22-0.99) and wateRmelon (median ICC=0.91, range=0.49-0.99) packages. Among the best performance of each 41 DNAm predictors, i.e. achieving the highest reliability, 32 (78%), 4 (10%), and 3 (7%) predictors were from the ENmix, minfi, and wateRmelon package, respectively. Among ENmix pipelines; out-of-band (OOB) background estimation (15 out of 32), REgression on Logarithm of Internal Control probes (RELIC) dye-bias correction (19 out of 32), no quantile normalization (12 out of 32), and the Regression on Correlated Probes (RCP) probe-type bias correction (31 out of 32) yielded the highest reliability most often (see Figure S2). Two ENmix pipelines achieved the highest reliability for three predictors. The analytical pipeline that included OOB background estimation, RELIC dye-bias correction, no normalization, and RCP probe-type bias correction (i.e. “ENmix:oob_relic_nonorm_rcp”) performed best for the BioAge4HAStatic, LDL, and CD8pCD28nCD45RAn predictors. The pipeline that included OOB background estimation, RELIC dye-bias correction, quantile normalization, and RCP probe-type bias correction (i.e. “ENmix: oob_relic_q1_rcp”) performed best for the B2M, DNAmTL, and HDLratio predictors.

### There is significant heterogeneity in pipeline performance across predictors

Among the 41 best performing pipelines (i.e. the pipeline with the largest ICC value for each of the 41 predictors), there are 27 different data processing and normalization strategies, which highlights significant heterogeneity in choice of best pipeline between predictors. As ICC differences between pipelines of a predictor can be small and pipelines beyond the highest ICC may also be informative, we calculated the median rank across the 41 predictors for each of the 101 pipelines (see Table S6). The pipeline with the best median rank (at 15) across predictors is the “ENmix: oob_relic_q1_rcp”. While this observation suggests this pipeline yields the best average performance across predictors, it still scored average to low for multiple predictors. For example, for the BMI predictor the “ENmix: oob_relic_q1_rcp” pipeline had one of the lowest ranks (ICC = 0.89, rank = 91). It is also important to note that a data processing pipeline can also introduce more spurious variation instead of removing technical variation. That is, the raw data pipeline that does not apply any data processing and normalization yielded a median rank of 85 (range: 7 to 100). For the CD4T and CD8 naive predictors, the raw data pipeline ranked as the seventh best performing pipeline highlighting that most pipelines perform worse than no data processing at all for these two predictors. The “Minfi: raw_quantile_strat” and “Minfi: illumina_bg_quantile_strat” had the lowest median rank of 100 and yielded the lowest reliability for 17 and 9 predictors, respectively (Table S5).

To assess the concordance in pipeline performance across predictors more formally, we calculated the rank correlation in pipeline reliability between all pairs of predictors. In Figure 3 we visualize the result of this analysis via a clustered correlation heatmap.

**Figure 3.**
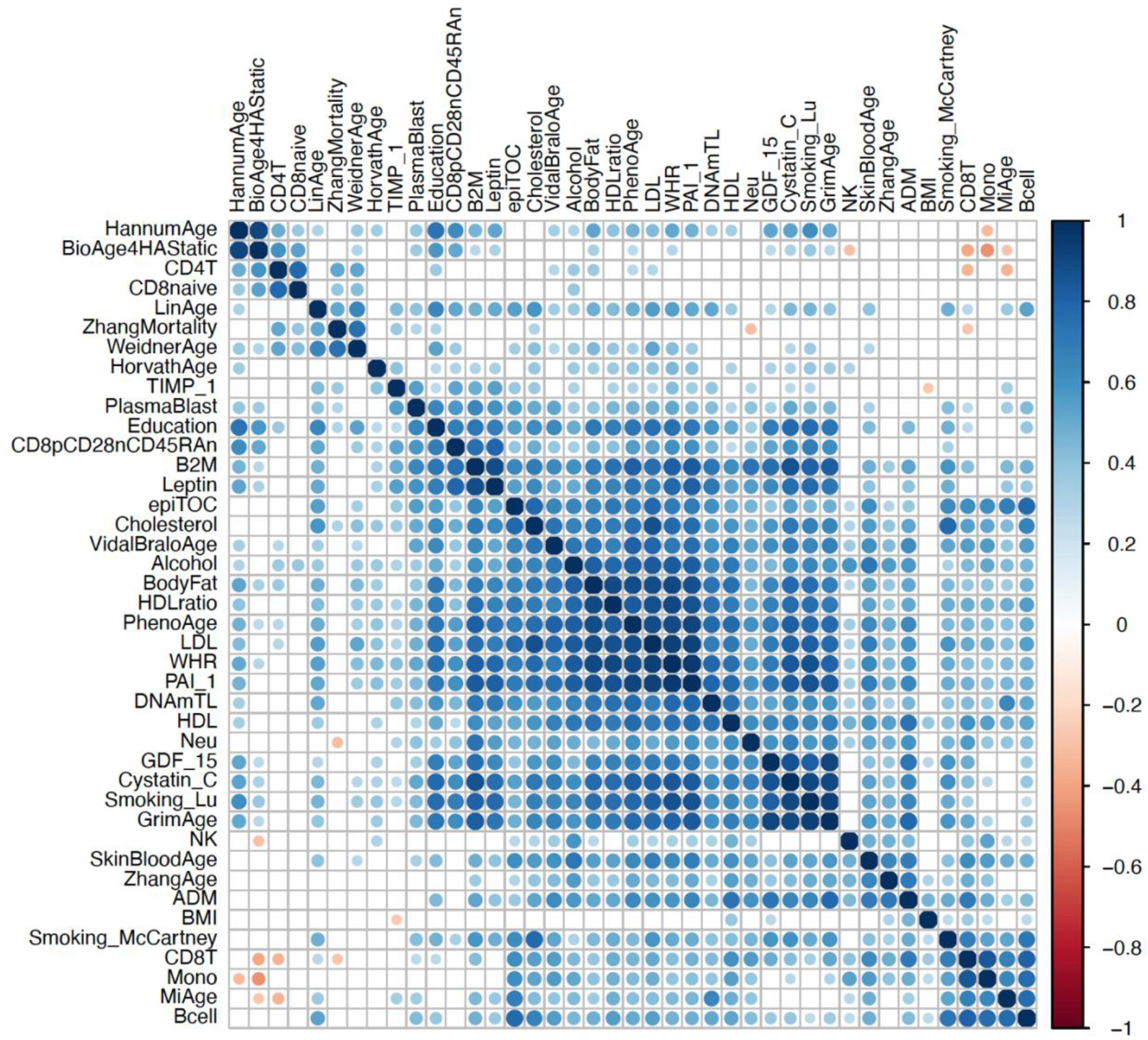
DNAm predictors have a moderate degree of concordance in performance between pipelines. Shown is a clustered correlation heatmap of pipeline reliability concordance between predictors. The color coding depicts Spearman’s rho and clustering is performed using hierarchical clustering. Only correlations with a P-value < 0.01 are colored.

For some predictors the ranking in pipeline performance is very similar. For example, the GrimAge, Smoking_Lu, Cystatin_C, and GDF_15 predictors show strong concordance (mean rho = 0.92). As noted, these four predictors were developed in the same dataset and the Cystatin_C, GDF_15, and Smoking_Lu estimates are included in the GrimAge algorithm. Across all pairs of predictors, we find a moderate correlation in pipeline performance (mean rho=0.40, SD=0.27). Some predictors however show little to no concordance with other predictors. The ranking of pipelines of the BMI and NK predictor, for example, have a mean rank correlation of 0.14 (SD=0.20) and 0.21 (SD=0.24), respectively, with that of other predictors. For a handful of predictor-pairs we even observe a negative correlation, suggesting that pipelines that yield high reliability for one predictor yield low reliability for another. Pipeline performance of the BioAge4HAStatic and Mono predictors for example have a correlation of −0.45 (P=2.1e-06). Our findings thus far show that specific pipelines are more effective in removing unwanted technical variation for a predictor and that significant heterogeneity exists in pipeline performance across predictors.

### The choice of data processing pipeline impacts downstream analysis of predictors

Next, we evaluated if the performance of a pipeline can also affect downstream phenotypic analyses of a predictor. For these analyses, we used the general JHS data 2 sample. For each pipeline, we calculated the mean and standard deviation (SD) of the predictor estimate distribution in the general JHS sample. For each predictor, we then correlated these two statistics (i.e., the mean and SD) with the ICC estimates of the pipelines obtained in the technical replicate sample. We find that the choice of pipeline has a significant impact on the distribution of the predictor estimate. Of the 41 predictors, 33 (80%) are significantly impacted on the distribution of their estimates after Bonferroni correction P<0.0012). For 22 predictors (54%), we find a significant correlation for both the mean and standard deviation. For DNAmTL, we, for example, observe a negative correlation between the performance of a pipeline and the mean of the estimate distribution (rho=−0.71, P<2.2e-16) and a positive correlation with the standard deviation of the estimate distribution (rho=0.79, P<2.2e-16). The best performing pipeline yields a mean estimate of 6.83 kilobases (SD=0.34). The least performing pipeline yields a mean estimate of 7.20 kilobases (SD=0.29). This shows that the more effective a pipeline is in removing technical variation, the lower the DNAm-based predicted estimate of telomere length and the larger the variation between individuals. The direction of effect of the relationship between pipeline performance and the mean and standard deviation of the DNAm variables varies between predictors as well. HorvathAge, for example, is impacted on its standard deviation (rho=0.39, P=5.6e-05) but not on the mean (rho=−0.10, P=0.27). HDLratio is impacted on its mean but unlike DNAmTL shows a positive correlation with pipeline performance (rho=0.38, P=9.8e-05). HDLratio is not impacted on the standard deviation of its distribution (rho=0.00, P=0.96). Correlation plots and correlation statistics of all predictors are shown in Supplementary Note 1. A full overview of test statistics can be found in Table S7.

Several DNAm age predictors are known to predict all-cause mortality risk. We therefore examined if pipeline performance also impacts their association with mortality risk. We focus on four predictors: HorvathAge, PhenoAge, GrimAge, and ZhangAge. Each predictor has different training characteristics and captures a different aspect of biological age and/or mortality risk[41]. ZhangAge is a blood-based DNAm clock and was developed on the largest training dataset and shown not to be associated with mortality risk despite its improved precision[8]. We find that pipeline performance significantly impacts downstream analysis for all four predictors (Figure 4).

**Figure 4.**
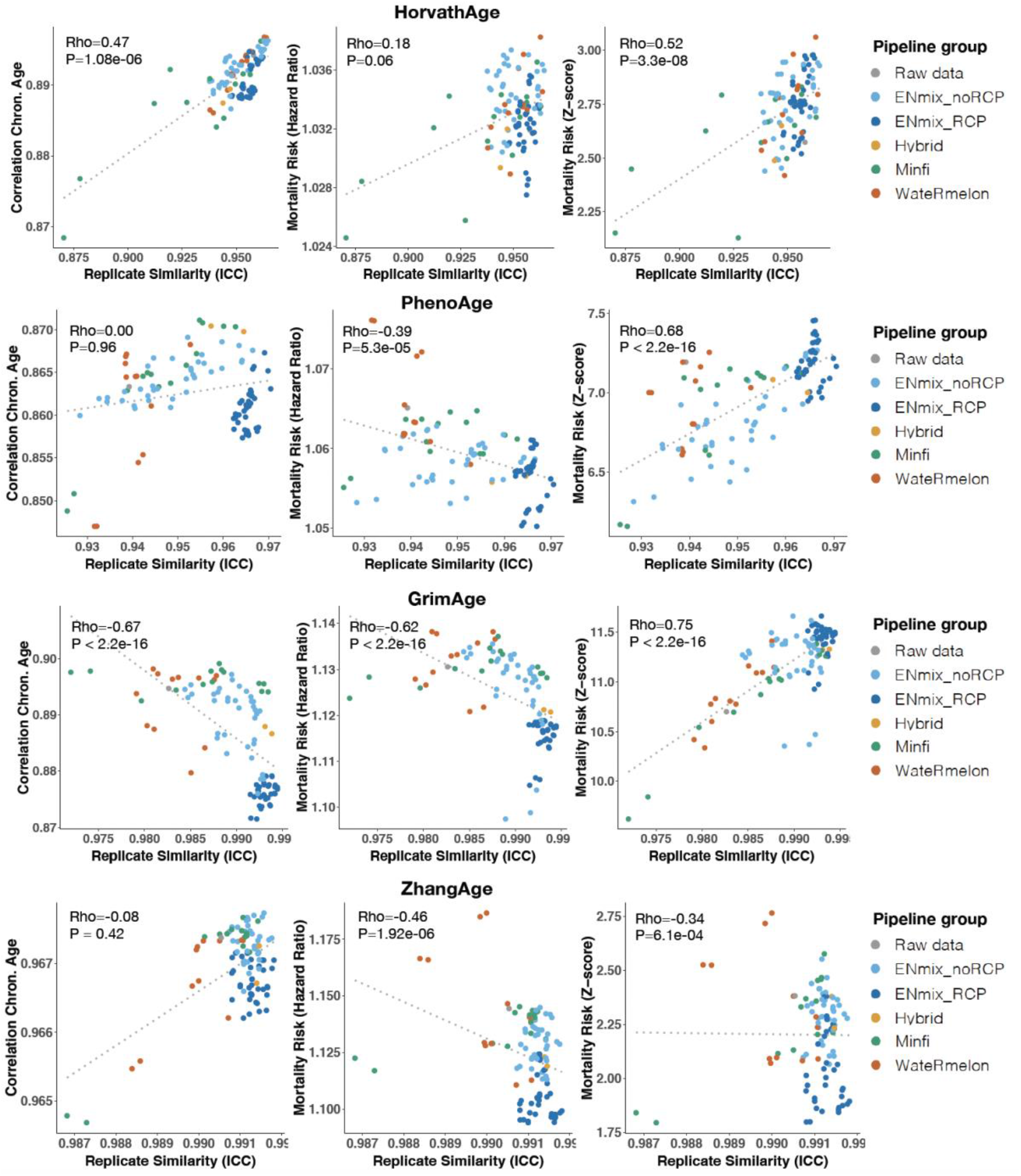
Pipeline performance impacts downstream analyses of DNAm age predictors. Shown are association between pipeline ICC and the correlation with chronological age (left panels), the hazard ratio of mortality risk prediction (middle panel), and the z-score of the mortality risk prediction (right panels) for Horvath Age (top row), PhenoAge (2nd row), GrimAge (3rd row), and ZhangAge (bottom row). Pipelines are color-coded by package/method. Spearman rank correlation statistics are shown in the top left corners.

For HorvathAge, pipelines that achieve greater reliability also achieve a greater correlation between HorvathAge and chronological age (rho=0.47, P=1.8e06). Better performing pipelines furthermore achieve greater power to predict all-cause mortality (rho=0.52, P=3.3e-08). For PhenoAge, we did not find an effect on the correlation with chronological age but did find the survival analysis to be significantly impacted. Better performing pipelines achieve greater power for PhenoAge (rho=0.68, P<2.2e-16) but also a smaller hazard ratio (rho=−0.39, P=5.3e-05), suggesting that unsuccessful removal of technical variation in DNAm data can inflate the magnitude of mortality risk. In contrast to our findings for HorvathAge, we found that better performing pipelines produced a lower correlation with chronological age for GrimAge (rho=−0.67, P < 2.2e-16). Similar to PhenoAge, we found that pipelines that achieve greater reliability yield more significant associations with mortality for GrimAge (rho=0.75, P<2.2e-16) but also a smaller hazard ratio (rho=−0.62, P<2.2e-16). The most reliable pipeline reports a significant hazard ratio of 1.12 (SE=0.01, P=1.60e-30), which verifies GrimAge as a strong predictor of all-cause mortality, especially when spurious technical variation is appropriately accounted for. For ZhangAge, we found no impact on the correlation with chronological age. Better performing pipelines produced smaller and less significant effects in associations with all-cause mortality. The most reliable pipeline produced a non-significant hazard ratio of 1.10 (SE=0.05, P=0.06), confirming that ZhangAge does not predict mortality risk. Taken together, using the general JHS sample, we demonstrate how pipeline performance has a significant impact on downstream phenotypic analysis of DNAm predictors.

### Predictor reliability is inversely associated with sample size of the training dataset

To assess if specific features of the predictors are associated with higher reliability, we investigated the number of CpG probes and the sample size of the training dataset in relation to the ICC of the best performing pipeline (see Figure S3). Using predictors for which such information was available, we find that the sample size of the dataset in which a predictor was developed is inversely associated with the observed predictor reliability (N=37, rho=−0.39, P=0.02). We did not find a significant association between the number of predictor CpG probes and reliability of a predictor (N=37, rho=−0.21, P=0.20).

### A smaller number of replicate pairs can be used to measure reliability

In our analyses, we made use of a large number of replicate pairs. We therefore assessed how sample size affected our measure of reliability and if a smaller number of replicate pairs yield similar findings. Across reliabilities from all pipelines and predictors, we observe good concordance (rho > 0.94) with as low as ten replicate pairs compared with measures obtained from larger sample sizes (Figure S4). Differences however exist between predictors with some predictors still requiring a larger number of replicate pairs (Supplementary Note 2).

## Discussion

DNAm-based predictors are emerging as powerful new methods to study health and disease, but little is known about the reliability of the estimates they produce. To investigate their performance, we carried out a systematic evaluation of 41 predictors across 101 data processing and normalization strategies and assessed to what degree algorithm performance is impacted by (un)successful removal of technical variation. Leveraging a large technical replicate sample in the JHS, we demonstrate that the choice of analytical pipeline has a significant impact on the reliability of predictors as well as on the outcomes of downstream phenotypic analyses. We highlight that specific pipelines are more effective in removing unwanted technical variation for a predictor but that significant heterogeneity exists in pipeline performance across predictors. Pipelines of the ENmix package achieved the highest reliability and were most frequently represented among the best performing pipelines. As research on DNAm-based predictors will continue to grow, our work provides best practices for the research community to help standardize their implementation and improve their performance.

To quantify method performance, we used a type of intraclass correlation that measures test-retest reliability by assessing the degree of absolute similarity between technical replicate pairs. Guidelines from reliability research suggest that ICC values less than 0.5 are indicative of poor reliability, values between 0.5 and 0.75 indicate moderate reliability, values between 0.75 and 0.9 indicate good reliability, and values greater than 0.90 indicate excellent reliability[40]. The ICC range of best performing pipelines across predictors was 0.83-0.99, indicating good to excellent reliability for these predictors. For 32 out of 41 predictors (78%), we found excellent reliability (ICC > 0.9) for at least one data processing pipeline. Several predictors show a reliability close to 1, which demonstrates that repeated collections of DNAm data yield almost the same predictor estimate and highlights their potential as a biomarker for health-related outcomes. Among predictors with high reliability are predictors of mortality risk, smoking behavior, blood cell types, and cancer risk. Demonstrating internal validity for these DNAm tools is important for research purposes but even more so for their potential utilization for health management and disease prediction in the clinic. GrimAge, a strong predictor of all-cause mortality, for example, has the highest test-retest reliability of 0.994. This finding demonstrates excellent test-retest reliability based on technical replicates from the same biological sample. It remains an open question if the measured reliability translates to repeated measures of DNA samples extracted from different blood draws at the same time point or across time points. The analytical framework we applied can however be easily extended to study design of other types of (biological) replicates. Establishing method reliability in other contexts of technical and biological variation is an important next step for future research.

We found that the choice of analytical pipeline is essential as multiple data processing strategies produced poor reliability (ICC<0.5) for several predictors. For some predictors, like for CD4T and CD8 naive T cells, using the raw data achieves higher reliability than most data processing pipelines. This highlights that analytical decisions on how to best prepare DNAm data require careful consideration as certain data processing and normalization steps can even reduce algorithm performance. Among the best performing pipelines of each predictor, we found significant heterogeneity across predictors. That is, there are 27 unique pipelines across the 41 predictors. On average, pipelines of the Enmix package achieved the highest reliability most frequently. While there is no one optimal pipeline to use for all predictors, several data processing steps stand out as producing high reliability for multiple predictors. For example, almost half of the best performing pipelines make use of the RELIC dye-bias correction method. RELIC uses the information between pairs of internal normalization control probes to correct for differences between color channels that measure intensity levels of the array[28]. The EPIC array contains 85 pairs of controls that target the same DNA region in housekeeping genes and contain no underlying CpG sites. RELIC uses the relationship between the pairs of controls to correct for dye-bias on intensity values for the whole array. Another data processing step that produced high reliability is the RCP probe type-bias correction method. 31 out of 41 of the best performing pipelines make use of this data processing step. RCP uses the existing correlation between pairs of nearby type I and II probes to adjust the beta values of all type II probes[27]. Both RELIC and RCP have been shown to reduce technical variation in DNAm data and are implemented in the ENmix package. While both approaches are effective in removing unwanted technical variation, we still recommend using the best performing pipeline for a specific predictor as reported in Table 1 as RELIC and RCP both show heterogeneity in performance across predictors.

The choice of analytical pipeline does not only impact the test-retest reliability of a predictor but also significantly affects downstream phenotypic analyses. We show that 80% of predictors are impacted on the mean and/or standard deviation of their distribution in the general JHS cohort. We furthermore analyzed DNAm clocks and showed that the strength of correlation between DNAm age and chronological age is affected in opposite directions for HorvathAge and GrimAge. While the correlation with chronological age becomes stronger with better performing pipelines for HorvathAge, the correlation becomes weaker for GrimAge. For DNAm clocks that are shown to be associated with mortality risk, successful removal of technical variation produced smaller hazard ratios but more significant associations. This highlights that not appropriately accounting for technical variation can decrease statistical power and inflate risk estimates for these predictors. It also shows that despite the narrow distribution of reliability estimates for these predictors, for example GrimAge has an ICC range of 0.921-0.994 indicating excellent reliability across all pipelines, the choice of pipeline still impacts downstream association analyses. We note that in our association analysis with mortality risk, we adjusted for chronological age, and still found that the choice of pipeline influences the outcome of the analysis. This is different from findings of a previous study that reported that the choice of pipeline influences the mean of DNAm age but not the DNAm age acceleration residual[42]. This study however only compared three data processing and normalization strategies and could have missed this effect as it did not perform a systematic evaluation across many pipelines. Finally, we confirm that ZhangAge, a DNAm clock developed in the largest blood based DNAm dataset, does not associate with mortality risk.

We also investigated if specific characteristics of a predictor impacted the measured reliability. We found that the sample size of the training dataset has a moderate inverse relationship with the reliability of a predictor. This suggests that predictors developed in larger training datasets are more sensitive to technical variation than predictors developed in a smaller dataset. This relationship could for example arise if larger training datasets on average have more technical factors that are not properly accounted for. The ZhangAge predictor, however, was developed in the largest training dataset and shows the second to highest reliability of all predictors we investigated. This indicates that other factors in addition to sample size of the training dataset are likely to play a role as well. ZhangAge was developed using 65 training sets across 14 cohorts, where each training set had a certain number (ranging between 1 and 13) of cohorts randomly sampled from the 14 cohorts[8]. This strategy is, as far as we know, unique to this predictor and may have helped select for CpG probes that are less impacted by technical variation due to its many training sets of different randomly assigned cohort compositions. As training datasets with large sample sizes are essential to developing more accurate DNA-based predictors, a strategy to randomize the potential effect of technical factors, like was implemented for the development of ZhangAge, could be worthwhile to consider for new predictors as well. We did not find a significant relationship between the number of CpG probes and the observed reliability of a predictor.

Our study comes with limitations. First, we measured reliability using technical replicate in one study. A different cohort or different types of repeated measures may yield different outcomes. Ideally, one would use study-specific replicate samples and assess if similar best practices are achieved or if alternative strategies are more appropriate to remove technical variation most optimally for that specific study. If future studies have the means to include replicate samples, they should aim to include at least ten replicate pairs. We determined that for most predictors a sample size of ten replicate pairs can already provide meaningful insights into their reliability. Second, several predictors were not fully compatible with the EPIC array platform. Predictors that were developed on older DNAm array platforms showed lower reliability. Missing probes could have affected the outcome of our analysis. Having said that, as the older 27K and 450K DNAm array platforms are discontinued, any future application of predictors that are not fully compatible with the EPIC array will face a similar challenge.

## Conclusion

In summary, this study demonstrates that considerable variation exists in the performance of DNAm-based predictors depending on the data processing and normalization strategy implemented. Analytical pipelines that best remove unwanted technical variation in DNAm data achieve excellent test-retest reliability for most predictors thereby demonstrating their potential as biomarkers for health-related outcomes. DNAm is an important tool to study health and disease. As the number of DNAm predictors continues to rise, understanding how best to improve and implement these algorithms will be essential for downstream clinical applications.

## Methods

### Cohort descriptions

The Jackson Heart Study is a large observational study of African American individuals from the Jackson, Mississippi (USA), metropolitan area[34]. JHS seeks to study the causes and disparities in cardiovascular health and related phenotypes in African Americans. Data and biological materials have been collected from 5,306 participants. For a subset of the cohort, peripheral blood samples were collected at baseline and subsequently used to quantify DNA methylation using the Illumina Infinium MethylationEPIC BeadChip that covers over 850,000 CpG sties. These samples have been included in previous DNAm studies[15,35]. See Table S1 for cohort characteristics. In our analysis, we included individuals for which DNAm data, phenotypic variables, and mortality data were available (N=1,909, 62.2% women, mean (SD) of age = 56.1 (12.4) years). For 146 individuals, technical replicates were collected. We therefore divided this dataset into two samples; 1) a general cohort sample that does not include technical replicate pairs (N=1,761, 62.6% women, mean(SD) of age=56.0 (12.3)) and 2) a technical replicate sample (N=146, 57.5% women, mean (SD) of age=57.4 (14.0)). Replicate pairs represent DNAm samples that were assayed twice using the EPIC array at separate occasions but originate from the same DNA extraction sample.

### Data preprocessing and normalization strategies

To perform a systematic evaluation of available data preprocessing and normalization strategies, we incorporated all methods that are available through the commonly used R packages minfi[36], wateRmelon[23], and ENmix[25]. Within the same package, we implemented all possible combinations of background correction, dye-bias correction, probe correction, and data normalizations as was feasible within the structure of the package. In total, this yielded 101 strategies to prepare DNAm data (Table S2). For each sample, raw intensity values were read from IDAT files into an RGChannelSetExtended object in the R programming environment using the read.metharray() function in minfi. Sample quality control was performed by excluding samples with more than 5% of CpG sites with a detection P-value greater than 0.05 (using the pfilter() function in the wateRmelon package) and by removing outlying samples based on a low median of chipwide (un)methylation across CpG sites (using the getQC() function in minfi). In total, 44 samples were removed. No probes were filtered out to minimize missing probes in downstream DNAm prediction analysis. Data processing and normalization were then executed in batches of 96 samples for computational efficiency. The output of each analytical pipeline was a matrix with beta values for each sample. Table S3 shows an overview of our sample quality control analysis.

### DNAm-based predictors

DNAm predictor estimates were calculated using regression coefficients as reported by the corresponding study unless stated otherwise. Custom R scripts were implemented that take as input a matrix of EPIC array beta values and output predicted estimates as a linear combination of weighted CpG methylation levels. For DNAm clocks, inverse transformation was applied to calibrate the DNAm age estimates in units of years, as required by the algorithm. For instance, Horvath’s epigenetic clock regressed log-linear age (that leveraged age at 20) on DNA methylation levels and required this calibration step.

Next, we briefly describe the different predictors included in our study. Table S4 presents an overview of predictor characteristics. For full details on each predictor, we refer to their corresponding studies.

#### DNAm clocks

The following predictors all output a form of DNAm age and capture a different aspect of biological age depending on characteristics of their training dataset. The Hannum clock uses 71 CpG probes and was developed in a whole blood 450K DNAm dataset of 656 individuals[7]. The Horvath clock was developed using 3,931 multi-tissue and -cell type samples using both 27K and 450K array samples[6]. The Horvath clock uses 353 CpG probes that are present on both arrays. The BioAge4HAStatic clock is an extended measure of the Hannum clock and defined by forming a weighted average of Hannum’s estimate with 3 cell types that are known to change with age: naïve (CD45RA+CCR7+) cytotoxic T cells, exhausted (CD28−CD45RA−) cytotoxic T cells, and plasmablasts[17]. The Weidner clock uses 3 CpG and was developed in a 27K DNAm dataset of whole blood samples from 575 individuals[11]. The Lin clock uses 99 CpG and was developed in a dataset of 450K array whole blood samples of 656 individuals[10]. The VidalBralo clock uses 8 CpG probes and was developed in a dataset of 450K array whole blood tissue of 390 individuals[13]. The Skin & Blood clock uses 391 CpG probes and was developed in a dataset of 450K and EPIC arrays of a mixture of human fibroblasts, skin tissue, buccal cells, endothelial cells, whole blood, and cord blood samples (N=896)[9]. The Zhang clock uses 514 CpG probes and was developed in a dataset of EPIC and 450K arrays of 13,566 samples. The majority of the samples were derived from whole blood with a small subsample from saliva tissue[8].

#### Mitotic clocks

The MiAge calculator uses 268 CpG probes and was developed on 4,020 samples of 8 cancer types using 450K DNAm arrays[19]. MiAge outputs an estimate of mitotic age (total number of lifetime cell divisions) for a given human tissue. The epiTOC calculator was developed in a 450K DNAm dataset of 650 whole blood samples. EpiTOC uses a subset of 385 Polycomb group targets promoter CpGs to predict an estimate of age acceleration in cancer. EpiTOC yields a score, denoted “pcgtAge”, as the average DNAm over CpG sites, representing the age-cumulative increase in DNAm at these sites due to putative cell-replication errors[18].

#### Mortality risk estimators

The Zhang mortality score is defined by a weighted average of 10 CpGs that are associated with mortality status[16]. The Zhang mortality score predictor was trained on a discovery cohort of whole blood 450K DNAm samples from 954 individuals (N=402 deceased at follow-up) and validated in a cohort of 1,000 individuals (N=231 deceased at follow-up). The second mortality estimator, Levine clock, is a predictor of “phenotypic age”, which is a DNAm surrogate of the composite score based on ten mortality markers (9 clinical markers + chronological age)[37]. A training cohort of 456 whole blood samples were then used to identify 513 CpGs predictive of phenotypic age. Only probes available on the 27K, 450K, and the EPIC array platform were used in their analysis. The linear combination of the weighted 513 CpGs is called “DNAm PhenoAge”. The third mortality risk estimator isGrimAge from Lu et al., which is defined by a composite score based on seven DNAm-based plasma protein markers, DNAm-based pack years of smoking, chronological age and gender[15]. GrimAge used a training dataset of whole blood samples of 1,731 individuals. The DNA methylation profiling was based on the 450K beadchip but the biomarker was trained on the CpGs present on both the 450K and the EPIC array in order to ensure compatibility for both platforms. GrimAge was calculated using a python executable that was developed by the authors of the original study, which also outputs several DNAm-based plasma protein markers, three blood cell types, and pack years of smoking (see below).

#### Plasma protein markers

DNAm-based estimators were developed for the following seven plasma proteins; adrenomedullin (ADM), beta-2-microglobulin (B2M), Cystatin-C, growth differentiation factor 15 (GDF-15), leptin, plasmin activator inhibitor 1 (PAI-1), tissue inhibitor metalloproteinases 1 (TIMP-1). These plasma proteins were measured using an immunoassay and the predictor trained using a whole blood 450k DNAm dataset of 1,731 individuals in Framingham Heart Study (FHS) cohort[15]. ADM, B2M, cystatin-C, GDF-15, leptin, PAI-1, and TIMP-1 are defined by 186, 91, 87, 137, 187, 211, and 42 CpGs, respectively. Each of these individual estimates were calculated using the GrimAge python executable.

#### Smoking predictors

Two DNAm-based smoking predictors were included in our analysis. The Lu estimator was trained using a whole blood 450K DNAm dataset of 1,731 individuals in FHS and uses 172 CpGs for prediction, which is a component of GrimAge[15]. We estimated Lu pack years of smoking using the GrimAge python executable. The McCartney estimator was developed using EPIC DNAm data (only probes that are also present on the 450K platform) of 3,444 individuals[20]. The McCartney estimator uses 233 CpGs and outputs, similar to the Lu predictor, the number of pack years of smoking.

#### Blood cell type estimator

We included DNAm-based blood cell type estimators for nine cell types in our analysis. For neutrophils (Neu), B cells, monocytes (Mono), natural killer cells (NK), CD4+ T cells (CD4T), and CD8+ T cells (CD8T), estimators were developed using 850K EPIC DNAm data from magnetic sorted cells[5]. These six cell types were estimated using the estimateCellProp(refdata=“FlowSorted.Blood.EPIC”, nprobes=50) function of the ENmix R package. Plasma B cells (PlasmaBlasts), naive CD8+ T cells, and CD8+, CD28−,CD45RA− T cells (CD8pCD28nCD45RAn), were estimated based on the Horvath method [38] and computed using the same python executable as was used for the GrimAge estimator. These estimates are the same estimates that can be obtained through the online DNAm Age Calculator; https://dnamage.genetics.ucla.edu/.

#### Other estimators

We also included DNAm-based estimators that are developed for body-mass-index (BMI, in kg/m^2^), alcohol (units: per week), educational attainment (Edu, in years), total cholesterol (in mmol/L), HDL cholesterol (in mmol/L), LDL with remnant cholesterol (in mmol/L), total:HDL cholesterol ratio (HDL_ratio), waist-to-hip ratio (WHR), body fat (in %). These estimators were developed in a whole blood EPIC DNAm dataset (only probes that are also present on the 450K platform) of between 2,819 to 5,036 individuals and used between 205 to 1,109 CpG sites to predict DNAm-based estimates[20]. Finally, we also included an estimator of leukocyte telomere length (TL). This DNAm-based TL predictor was developed in a whole blood 450K/EPIC DNAm dataset of 2,256 individuals and uses 140 CpGs[12].

### Statistical analyses

In the sample of technical replicates, the intraclass correlation (ICC) was calculated using the ICC() function of the R *psych* package (v2.1.3). More specifically, we use ICC(2,1), which is a type of ICC that calculates reliability from a single-measurement using a two-way random effects model[39,40]. ICC(2,1) assumes absolute agreement, which means the estimates of the replicates are expected to have exactly the same value. We also calculated ICC(1,1), ICC(3,1), ICC(1,k), ICC(2,k), ICC(3,k) for comparison with other ICC types.

In the general JHS sample (i.e., without technical replicates), we calculated multiple statistical measures on the distribution of the output estimates of each predictor. The coefficient of variation was calculated by dividing the standard deviation by the mean of the distribution of the estimates. DNAm age acceleration residual (ΔAge) was calculated by regressing DNAm age on chronological age using the lm() function in R. To relate DNAm predictor estimates with mortality risk, a Cox proportional hazards regression model was fit using the coxph() function of the *survival* package (v3.2). Finally, to assess if the above statistical properties change depending on the type of data processing pipeline used, we calculated Spearman correlations between the ICC calculated in the replicate JHS sample and the various statistics generated in the general JHS sample across the 101 pipelines For this we use the cor.test(method=“spearman”) function of the *stats* package. The statistical analyses were performed in R (v4.0.3).

## Declarations

### Ethics approval and consent to participate

All participants included in this study provided consent per procedures of the JacksonHeart Study.

### Availability of data and materials

All DNA methylation and phenotypic information are available through the Jackson Heart Study. To submit a request for data, complete a data request form: https://www.jacksonheartstudy.org/Research/Study-Data/Data-Access

R code for the 101 DNA methylation data processing and normalization pipelines can be downloaded here: https://github.com/anilpsori/DNAm_pipelines_and_biomarkers. This GitHub repository also contains R scripts to calculate estimates for DNAm predictors that are described in the manuscript.

### Competing interests

UC Regents (the employer of SH and ATL) has filed patents surrounding several epigenetic biomarkers of aging (including GrimAge) which list SH and ATL as inventors. The other authors declare that they have no competing interests.

### Funding

ATL and SH were supported by NIH Grant 1U01AG060908 – 01.

## Acknowledgements

The Jackson Heart Study (JHS) is supported and conducted in collaboration with Jackson State University (HHSN268201800013I), Tougaloo College (HHSN268201800014I), the Mississippi State Department of Health (HHSN268201800015I) and the University of Mississippi Medical Center (HHSN268201800010I, HHSN268201800011I and HHSN268201800012I) contracts from the National Heart, Lung, and Blood Institute (NHLBI) and the National Institute on Minority Health and Health Disparities (NIMHD). The authors also wish to thank the staffs and participants of the JHS. We furthermore thank James G. Wilson for his contributions to the study.

## Author contributions

APSO, SH, and RAO conceived of the study. APSO performed data analyses and primary interpretation of results and writing of the manuscript. ATL developed the GrimAge executable and provided code to estimate the GrimAge predictors and run the survival analyses. SH and RAO oversaw the work. JGW and SH provided access to data of the JacksonHeart Study. All authors read, gave input on, and approved the final manuscript.

## Disclaimers

The views expressed in this manuscript are those of the authors and do not necessarily represent the views of the National Heart, Lung, and Blood Institute; the National Institutes of Health; or the U.S. Department of Health and Human Services.

