## Supplementary Figures for "A systematic evaluation of 41 DNA methylation predictors across 101 data preprocessing and normalization strategies highlights considerable variation in algorithm performance"

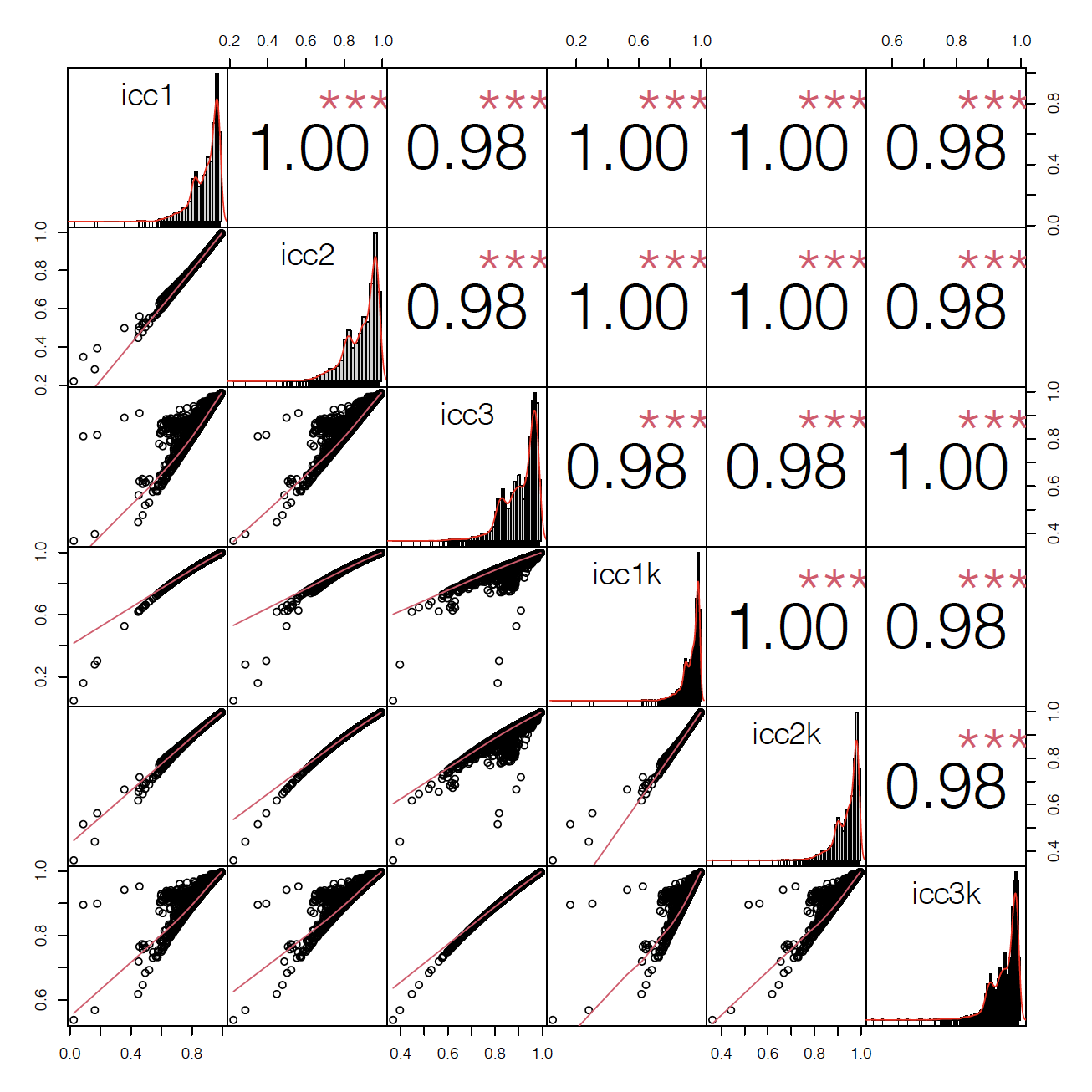

**Figure S1. Comparative analysis of ICC types across DNAm-based predictors and data processing pipelines.** Shown are the bivariate scatterplots (left bottom) and the Spearman correlation (right top) between ICC types across all pipelines and predictors (N= 101x41 = 4141). The distribution of each ICC type is shown on the diagonal. ***P-values < 2.2e-16. This figure was made using the chart.Correlation() function of the PerformanceAnalytics R package (v2.0.4).

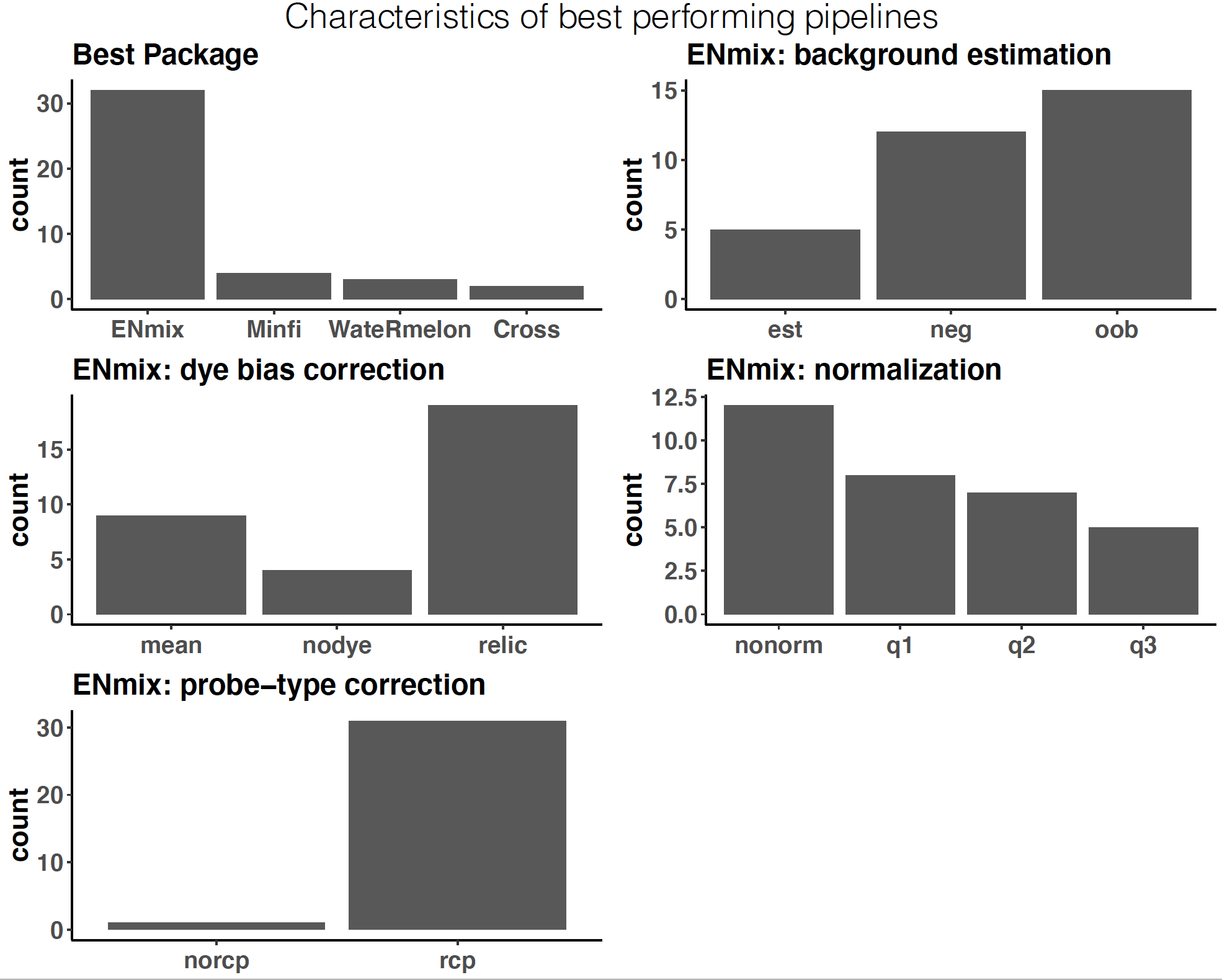

**Figure S2. Characteristics of best performing pipelines of predictors.** These graphs are based on the 42 best performing data processing pipelines (i.e., pipeline with the highest reliability of each predictors). Top left shows the corresponding package. 32 out of 41 pipelines are part of the Enmix package. The top right shows which background estimations ranked among the 32 Enmix pipelines. Middle left shows the ENmix dye bias correction method. Middle right shows the Enmix normalization method. The bottom graph shows if a pipeline used probe-type bias correction (i.e. “RCP method”).

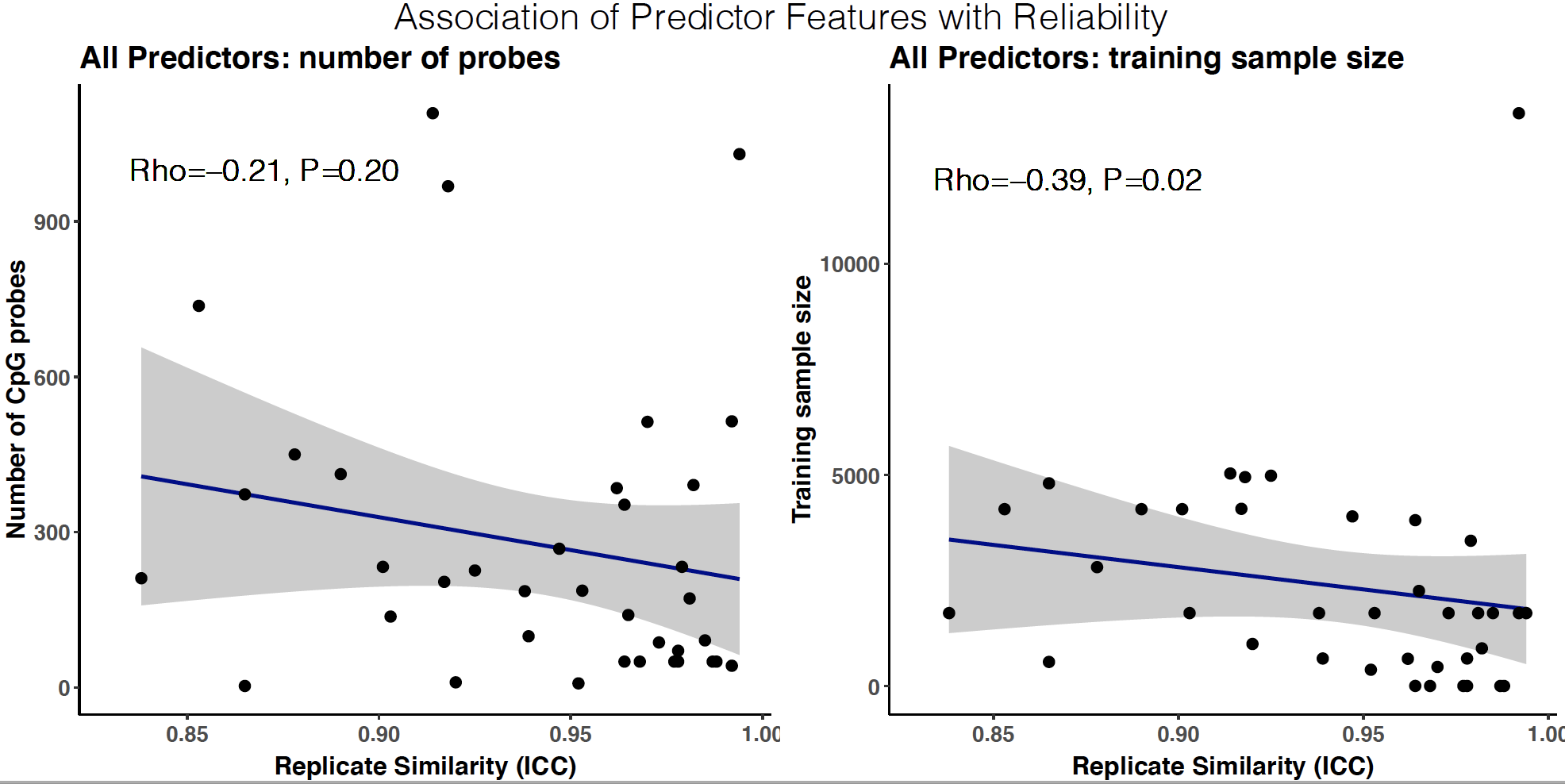

**Figure S3. Association of predictor features with reliability.** Shown are scatter plots of the relationships between predictor features (i.e., training sample size and number of CpG probes) and the reliability (i.e., ICC) of the best performing pipeline for each predictor. Shown are the statistics of the correlation test (method=”spearman”) and a corresponding regression line.

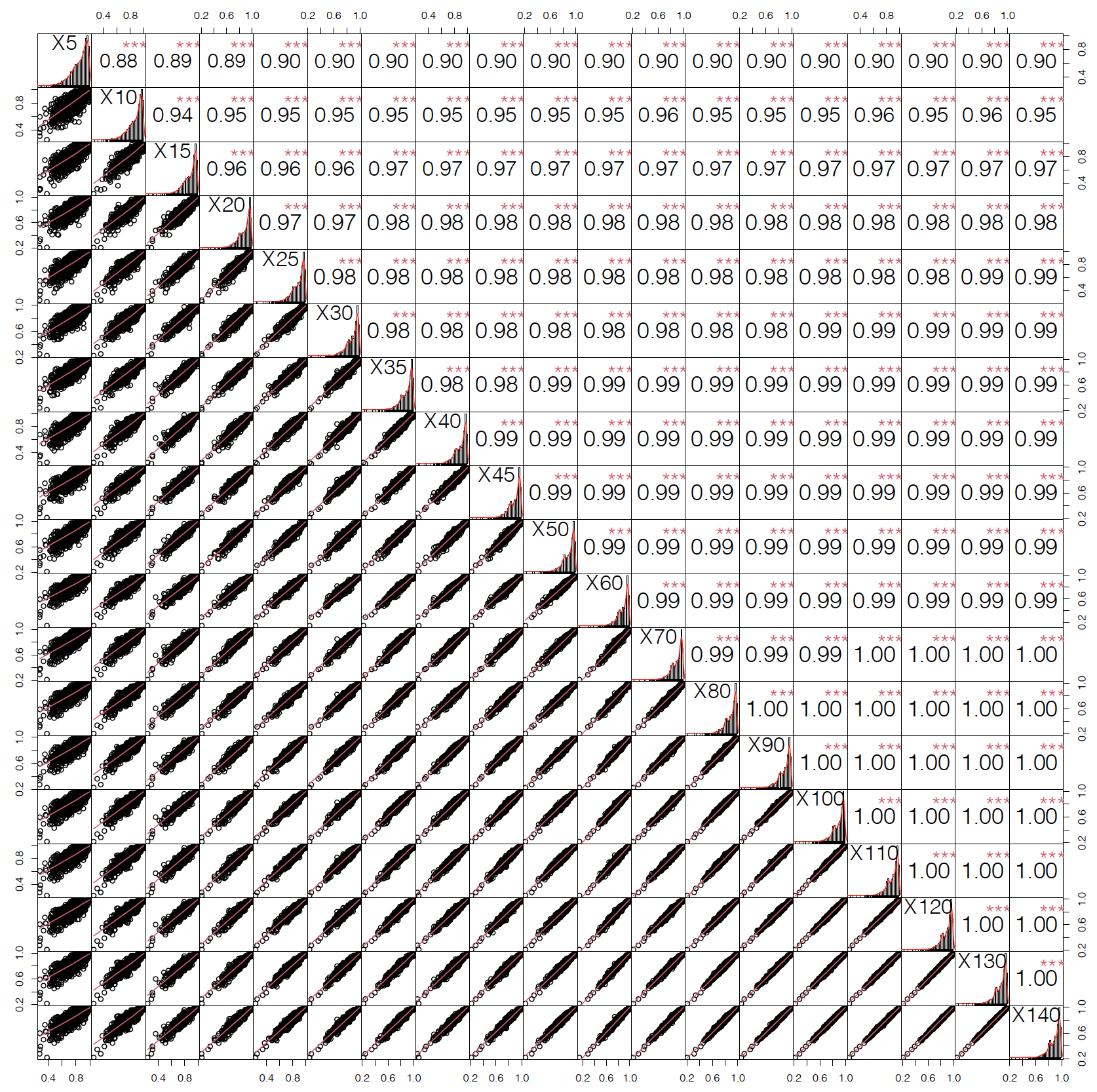

**Figure S4. Reliability measures across different sample sizes of replicate pairs.** Shown are the bivariate scatterplots (left bottom) and the Spearman correlation (right top) between the interclass correlations (all pipelines and predictors (N= 101x41 = 4141)) obtained across different sample sizes of replicate pairs. The sample size of the set of replicate pairs is shown on the diagonal across. For each sample size, we performed a bootstrap analysis in which we randomly selected the specified number of pairs from the total of 146 replicate pairs and computed the intraclass correlation across ten independent samplings. We then computed the mean intraclass correlation across these ten samplings and correlated this obtained mean ICC across different sets of replicate pairs. ***P-values < 2.2e-16. This figure was made using the chart.Correlation() function of the PerformanceAnalytics R package (v2.0.4).
