## Supplemental Note 1 for "A systematic evaluation of 41 DNA methylation predictors across 101 data preprocessing and normalization strategies highlights considerable variation in algorithm performance"

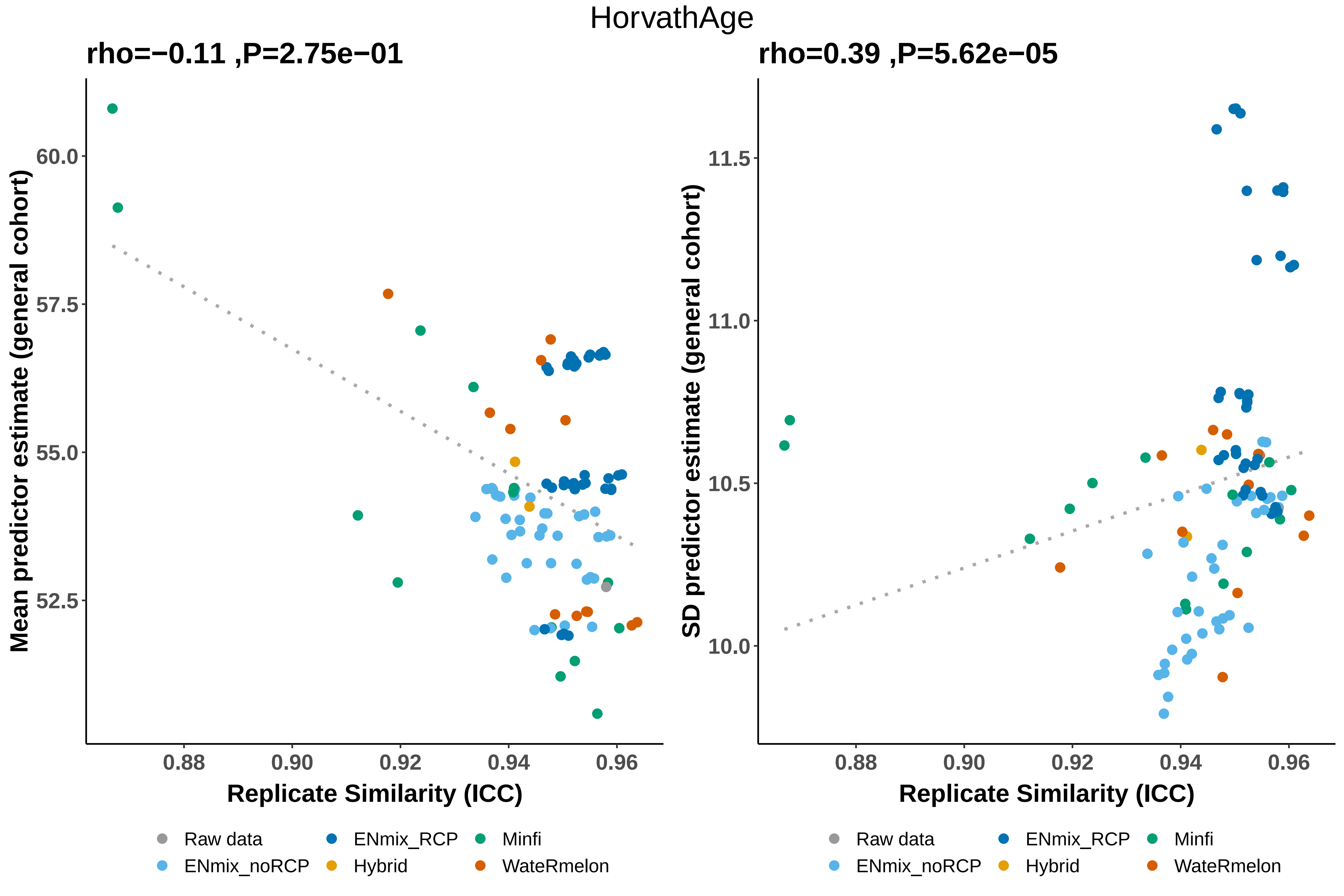
HorvathAge
rho=−0.11 ,P=2.75e−01
rho=0.39 ,P=5.62e−05
Mean predictor estimate (general cohort)
SD predictor estimate (general cohort)
60.0
57.5
55.0
52.5
11.5
11.0
10.5
10.0
0.88
0.90
0.92
0.94
0.96
Replicate Similarity (ICC)
Raw data
ENmix_noRCP
ENmix_RCP
Hybrid
Minfi
WateRmelon

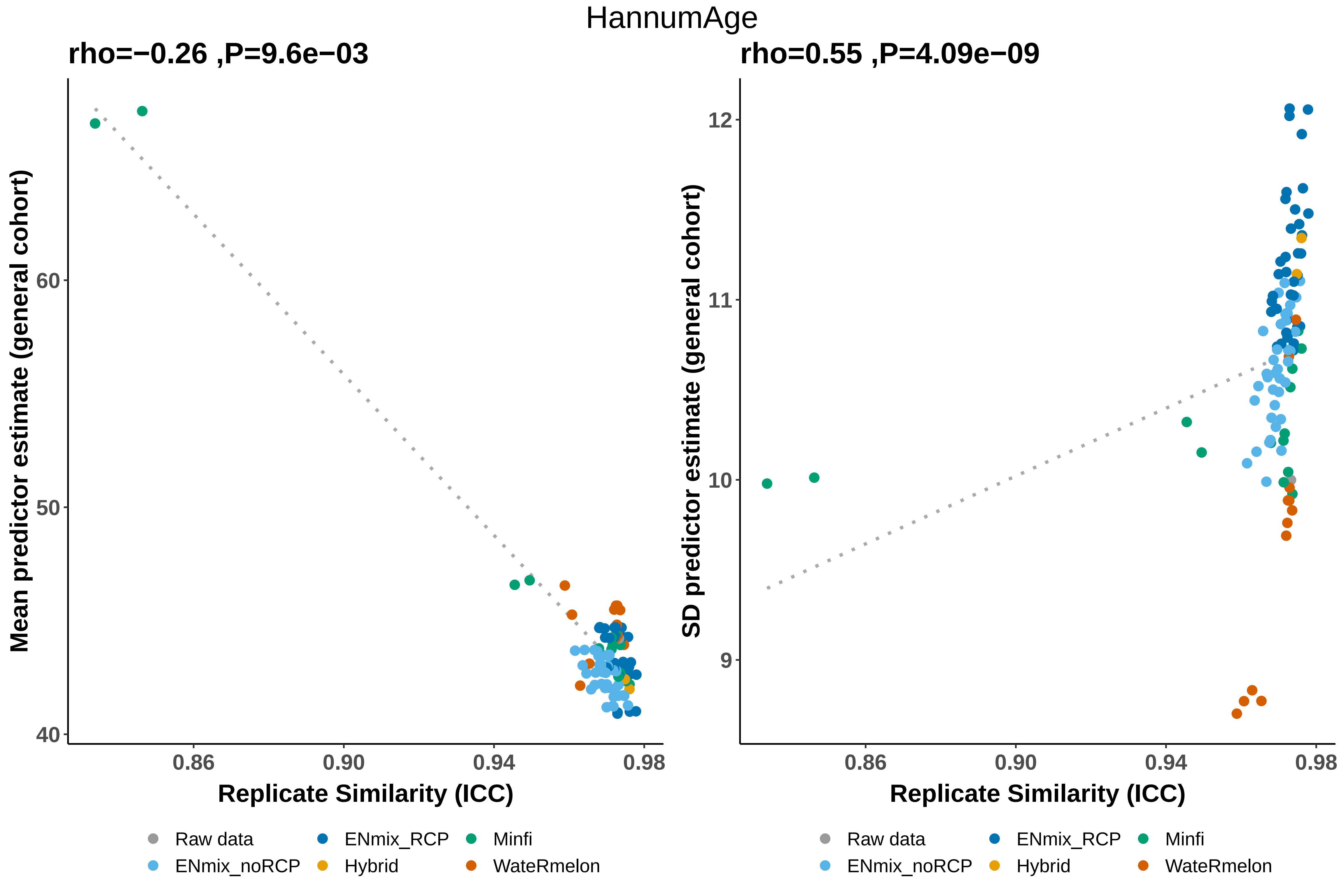
HannumAge
rho=−0.26 ,P=9.6e−03
rho=0.55 ,P=4.09e−09
Mean predictor estimate (general cohort)
SD predictor estimate (general cohort)
60
50
40
12
11
10
9
0.86
0.90
0.94
0.98
Replicate Similarity (ICC)
Raw data
ENmix_RCP
Minfi
ENmix_noRCP
Hybrid
WateRmelon

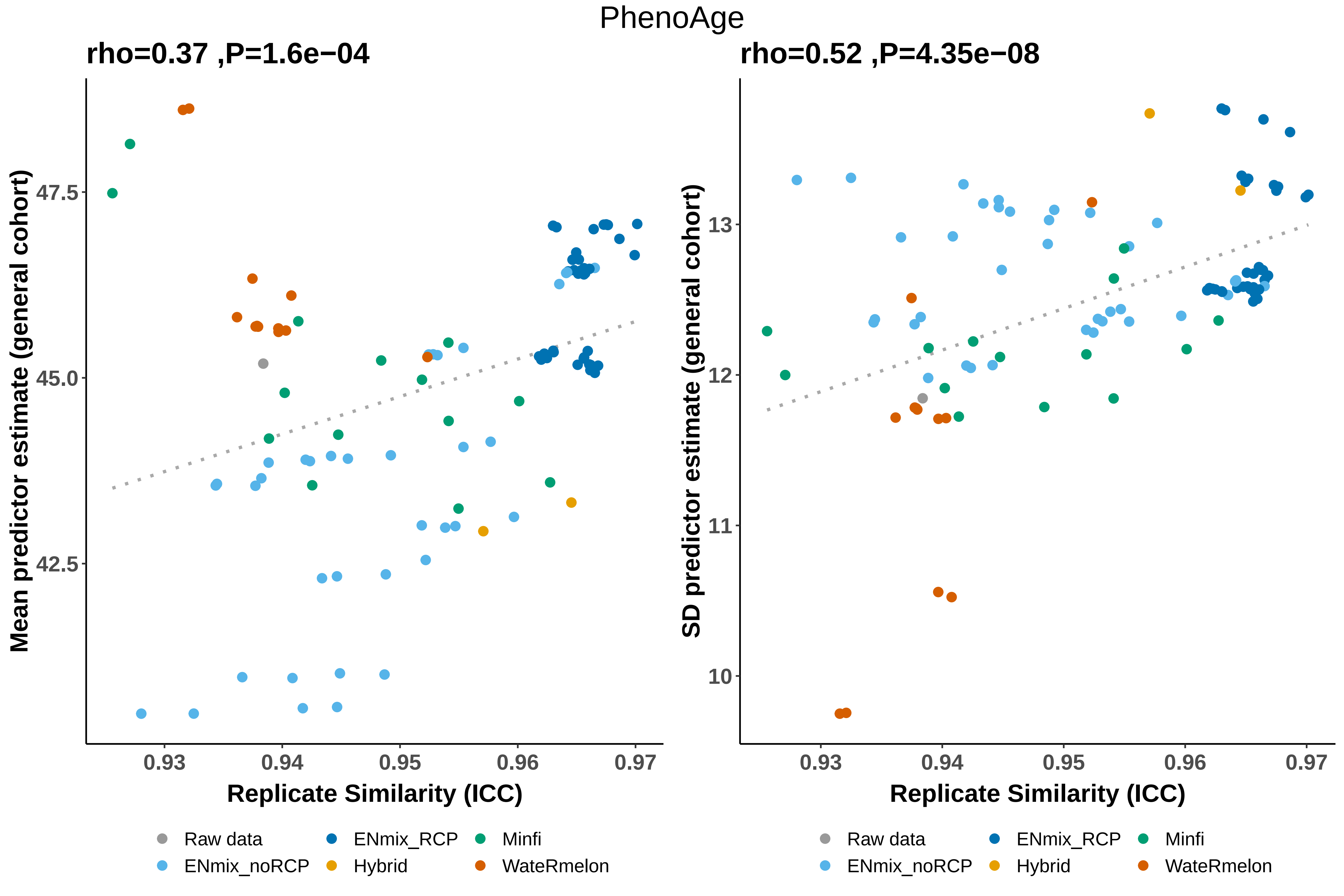
PhenoAge
rho=0.37 ,P=1.6e−04
rho=0.52 ,P=4.35e−08
Mean predictor estimate (general cohort)
SD predictor estimate (general cohort)
47.5
45.0
42.5
13
12
11
10
0.93
0.94
0.95
0.96
0.97
Replicate Similarity (ICC)
Replicate Similarity (ICC)
Raw data
ENmix_noRCP
ENmix_RCP
Hybrid
Minfi
WateRmelon
Raw data
ENmix_noRCP
ENmix_RCP
Hybrid
Minfi
WateRmelon

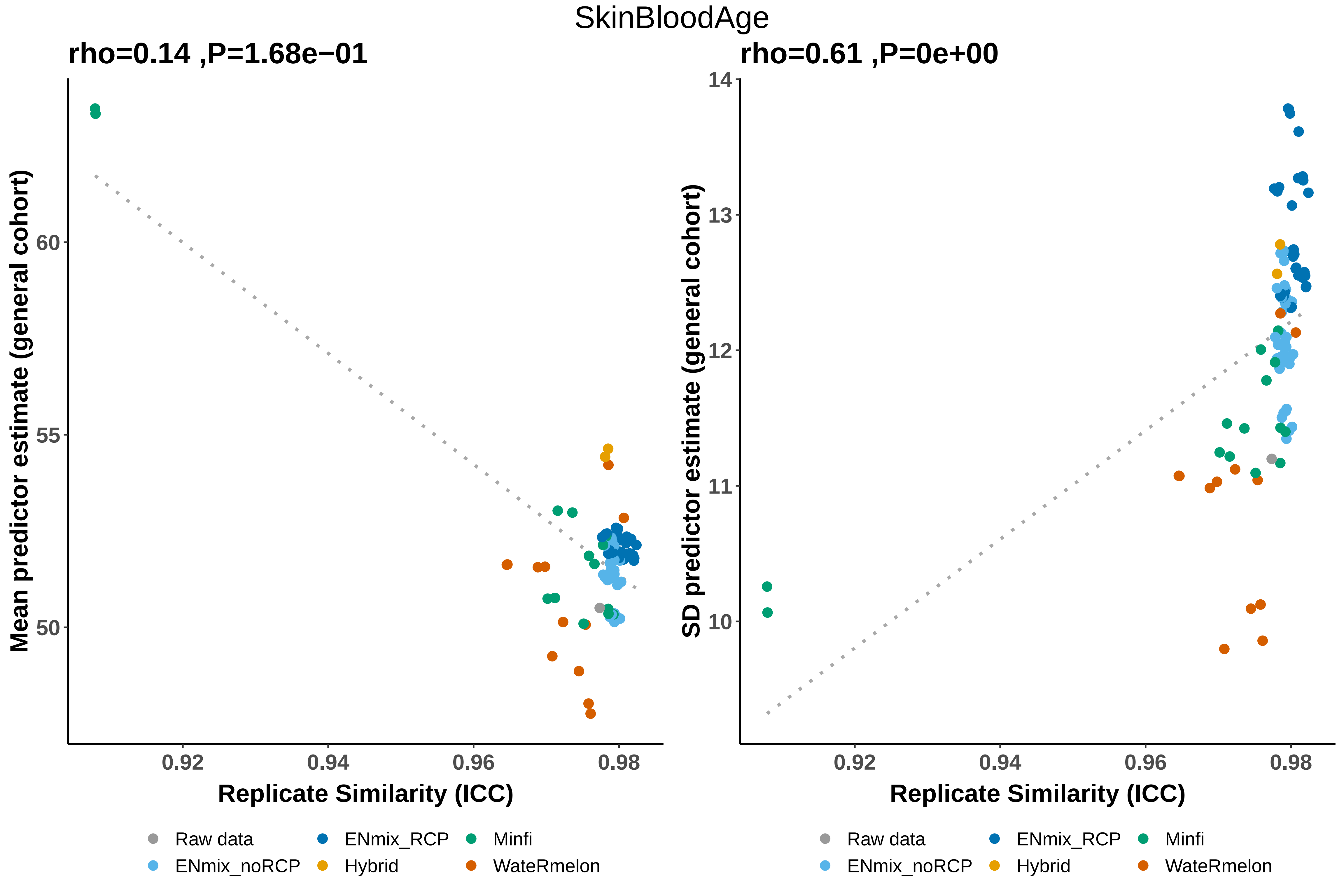

SkinBloodAge
rho=0.14 ,P=1.68e−01
rho=0.61 ,P=0e+00
Mean predictor estimate (general cohort)
SD predictor estimate (general cohort)
60
55
50
14
13
12
11
10
0.92
0.94
0.96
0.98
Replicate Similarity (ICC)
Replicate Similarity (ICC)
Raw data
ENmix_RCP
Minfi
ENmix_noRCP
Hybrid
WateRmelon

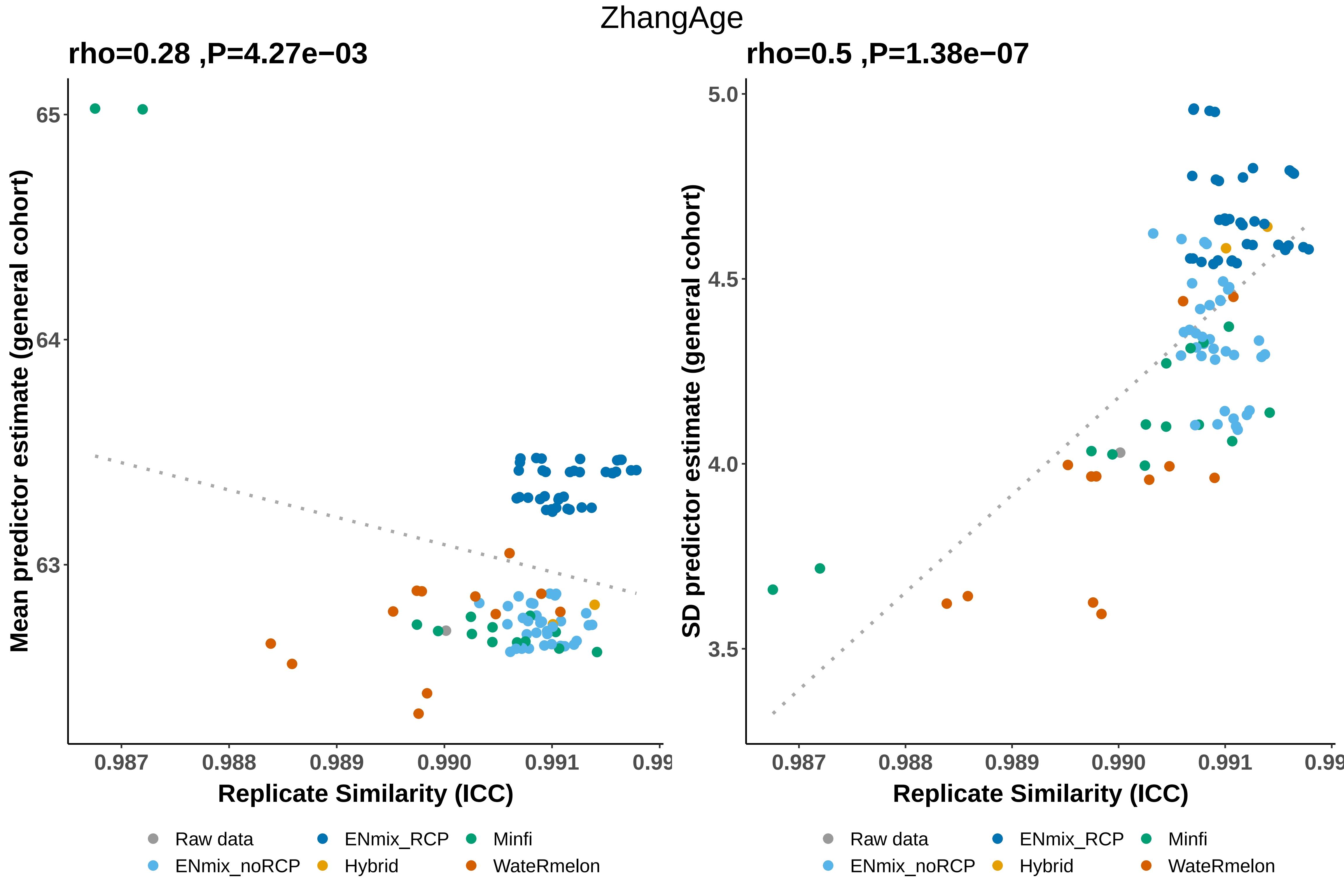
ZhangAge
rho=0.28 ,P=4.27e−03
rho=0.5 ,P=1.38e−07
Mean predictor estimate (general cohort)
SD predictor estimate (general cohort)
65
64
63
5.0
4.5
4.0
3.5
0.987
0.988
0.989
0.990
0.991
Replicate Similarity (ICC)
Raw data
ENmix_RCP
Minfi
ENmix_noRCP
Hybrid
WateRmelon

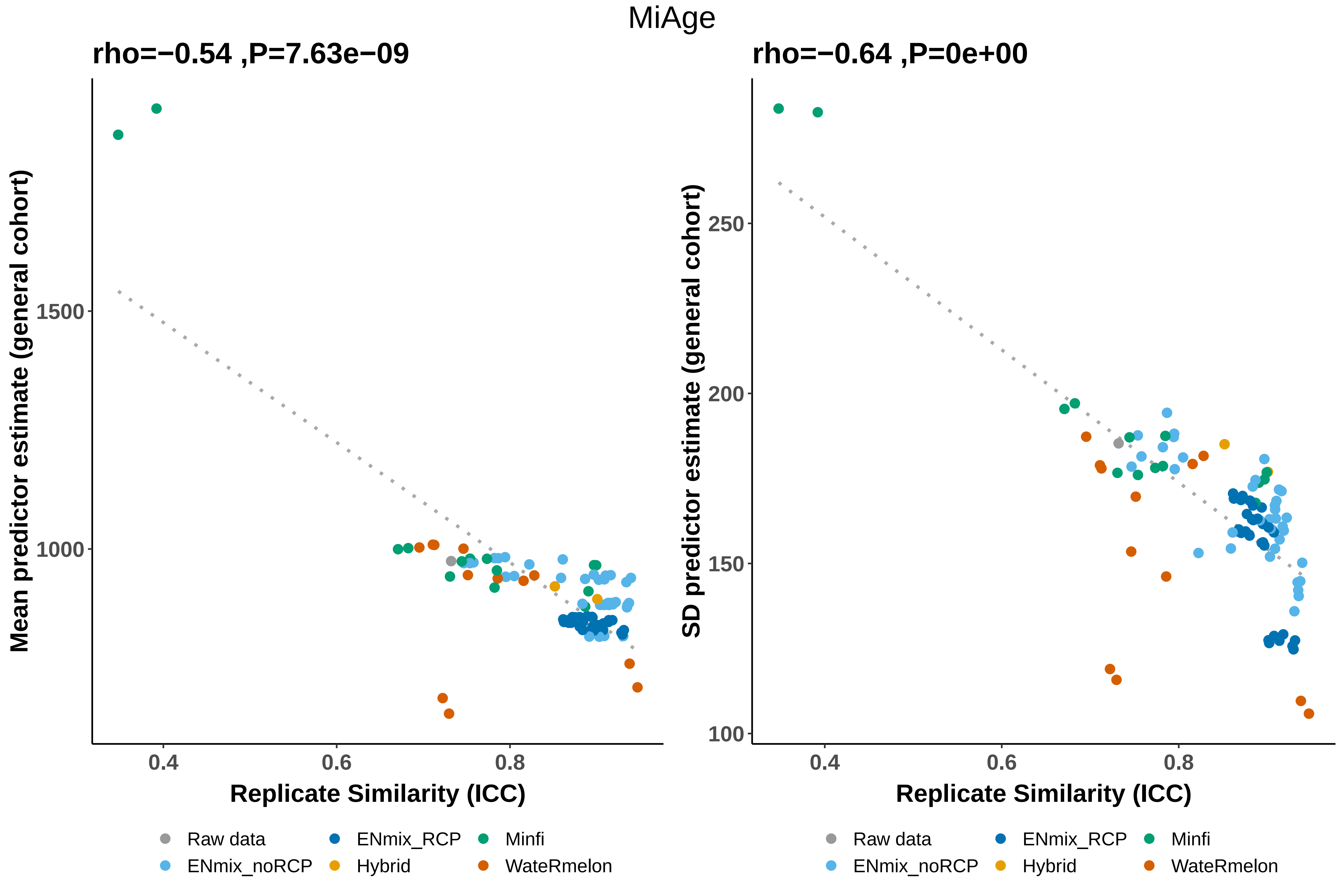
MiAge
rho=−0.54 ,P=7.63e−09
Mean predictor estimate (general cohort)
1500
1000
0.4
0.6
0.8
Replicate Similarity (ICC)
rho=−0.64 ,P=0e+00
SD predictor estimate (general cohort)
250
200
150
100
0.4
0.6
0.8
Replicate Similarity (ICC)
Raw data
ENmix_noRCP
ENmix_RCP
Hybrid
Minfi
WateRmelon

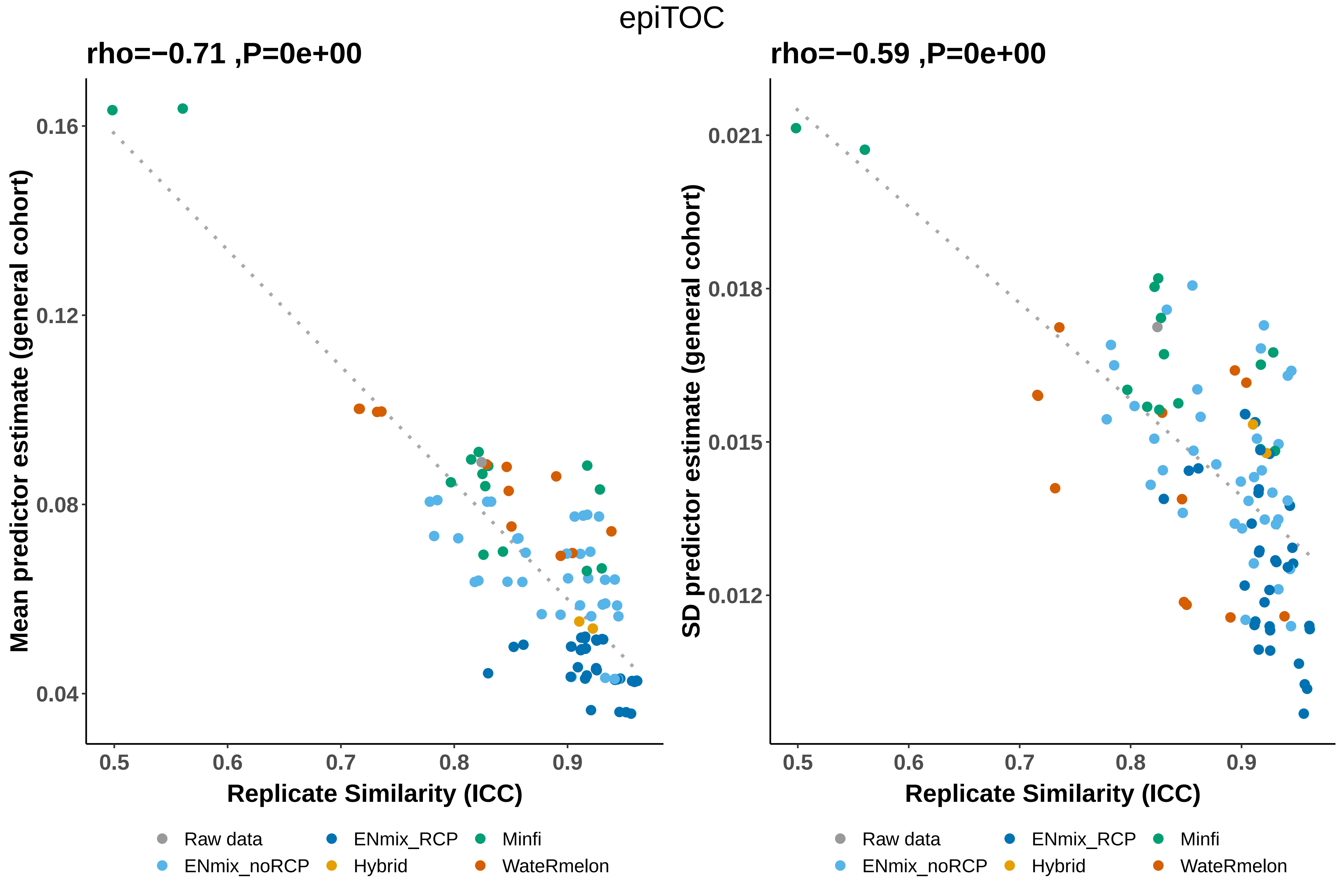

epiTOC
rho=−0.71 ,P=0e+00
rho=−0.59 ,P=0e+00
Mean predictor estimate (general cohort)
SD predictor estimate (general cohort)
0.16
0.12
0.08
0.04
0.021
0.018
0.015
0.012
0.5
0.6
0.7
0.8
0.9
Replicate Similarity (ICC)
Replicate Similarity (ICC)
Raw data
ENmix_noRCP
ENmix_RCP
Hybrid
Minfi
WateRmelon

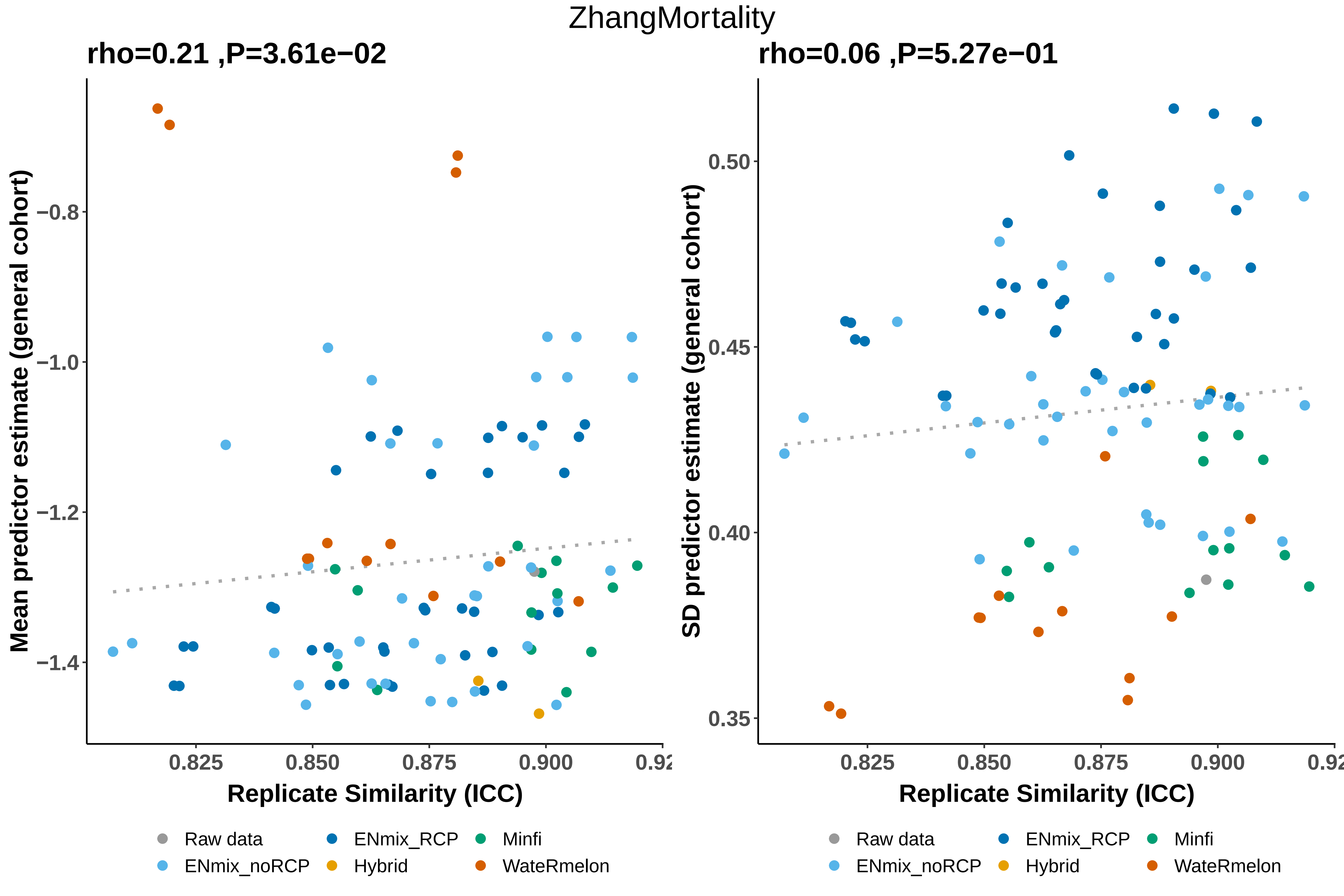

ZhangMortality
rho=0.21 ,P=3.61e−02
rho=0.06 ,P=5.27e−01
Mean predictor estimate (general cohort)
SD predictor estimate (general cohort)
−0.8
−1.0
−1.2
−1.4
0.50
0.45
0.40
0.35
0.825
0.850
0.875
0.900
Replicate Similarity (ICC)
Replicate Similarity (ICC)
Raw data
ENmix_noRCP
ENmix_RCP
Hybrid
Minfi
WateRmelon

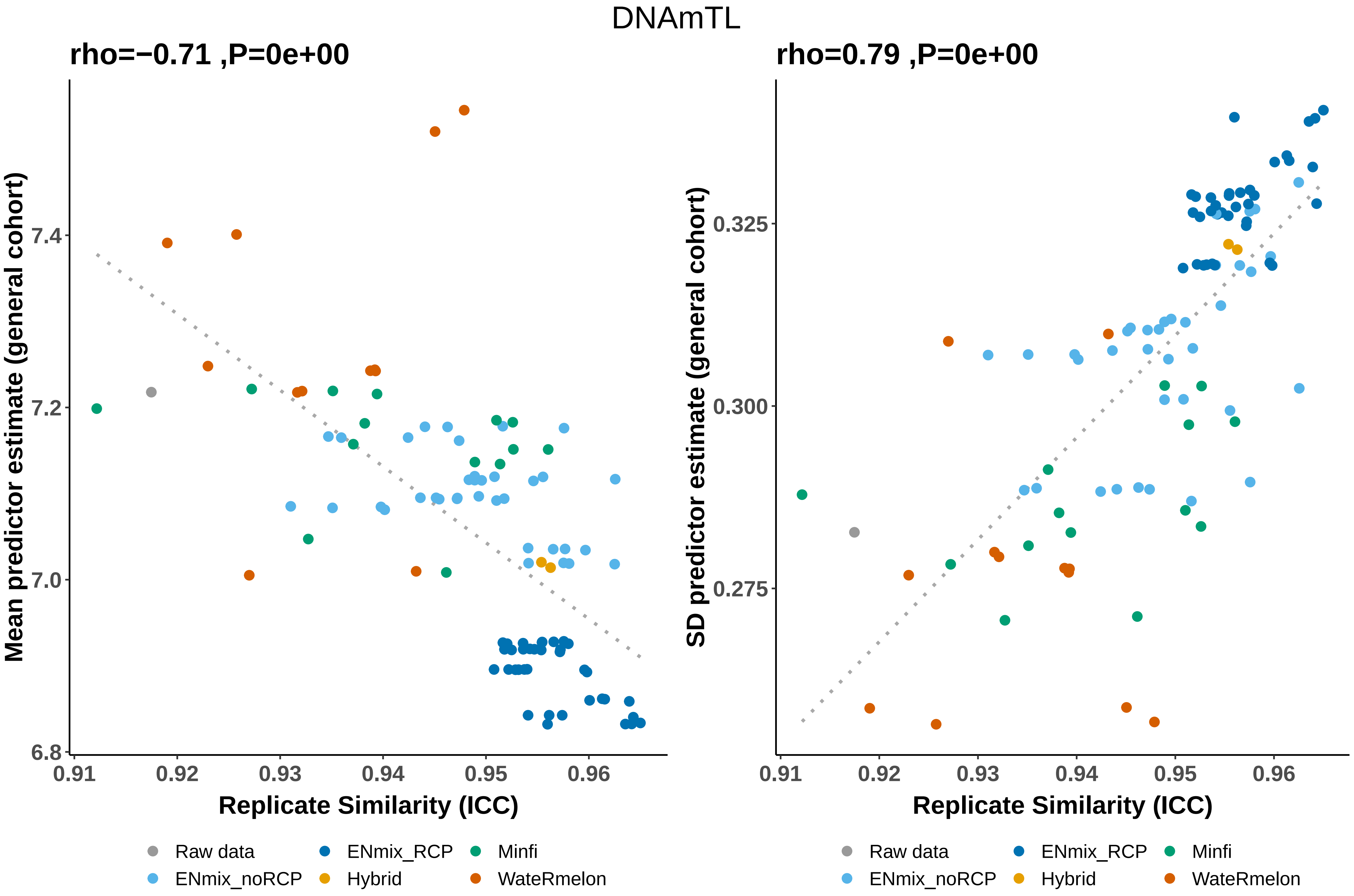

DNAmTL
rho=−0.71 ,P=0e+00
Mean predictor estimate (general cohort)
7.4
7.2
7.0
6.8
0.91
0.92
0.93
0.94
0.95
0.96
Replicate Similarity (ICC)
rho=0.79 ,P=0e+00
SD predictor estimate (general cohort)
0.325
0.300
0.275
0.91
0.92
0.93
0.94
0.95
0.96
Replicate Similarity (ICC)
Raw data
ENmix_noRCP
ENmix_RCP
Hybrid
Minfi
WateRmelon
Raw data
ENmix_noRCP
ENmix_RCP
Hybrid
Minfi
WateRmelon

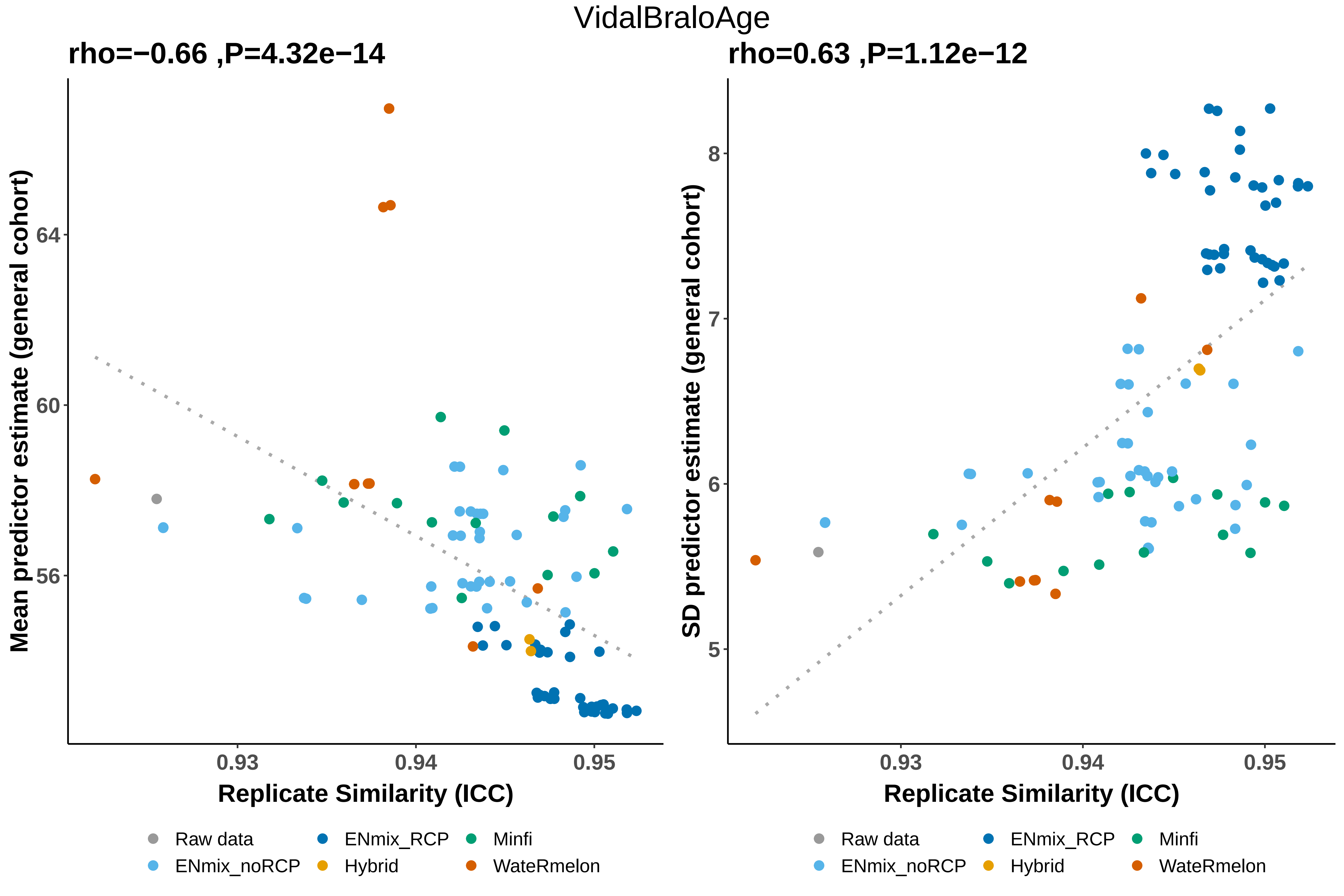

VidalBraloAge
rho=−0.66 ,P=4.32e−14
rho=0.63 ,P=1.12e−12
Mean predictor estimate (general cohort)
SD predictor estimate (general cohort)
Replicate Similarity (ICC)
Replicate Similarity (ICC)
64
60
56
8
7
6
5
0.93
0.94
0.95
0.93
0.94
0.95
Raw data
ENmix_RCP
Minfi
ENmix_noRCP
Hybrid
WateRmelon
Raw data
ENmix_RCP
Minfi
ENmix_noRCP
Hybrid
WateRmelon

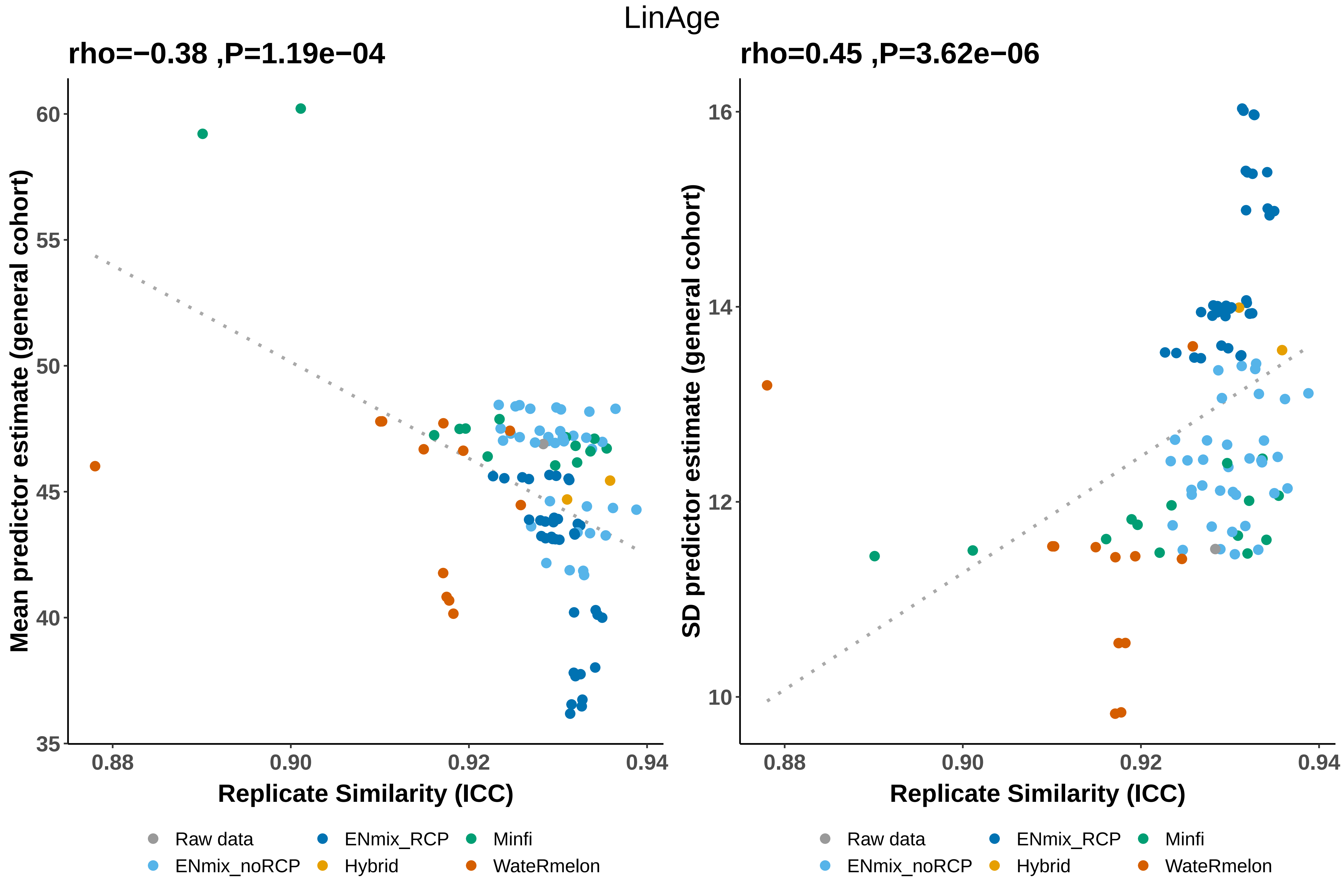
LinAge
rho=−0.38 ,P=1.19e−04
rho=0.45 ,P=3.62e−06
Mean predictor estimate (general cohort)
SD predictor estimate (general cohort)
60
55
50
45
40
35
16
14
12
10
0.88
0.90
0.92
0.94
Replicate Similarity (ICC)
Raw data
ENmix_RCP
Minfi
ENmix_noRCP
Hybrid
WateRmelon

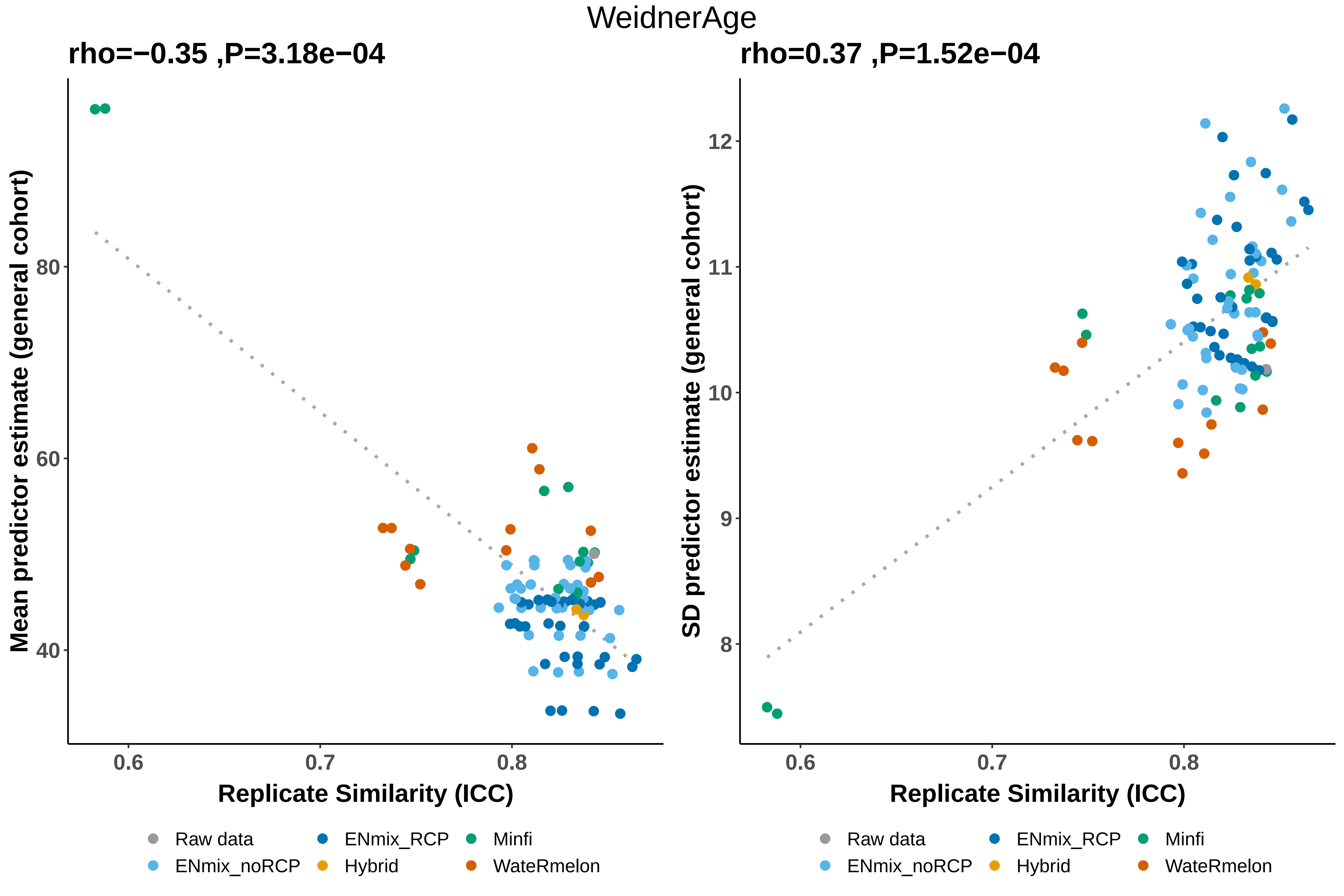

WeidnerAge
rho=−0.35 ,P=3.18e−04
rho=0.37 ,P=1.52e−04
Mean predictor estimate (general cohort)
SD predictor estimate (general cohort)
Replicate Similarity (ICC)
Replicate Similarity (ICC)
40
60
80
8
9
10
11
12
0.6
0.7
0.8
Raw data
ENmix_noRCP
ENmix_RCP
Hybrid
Minfi
WateRmelon

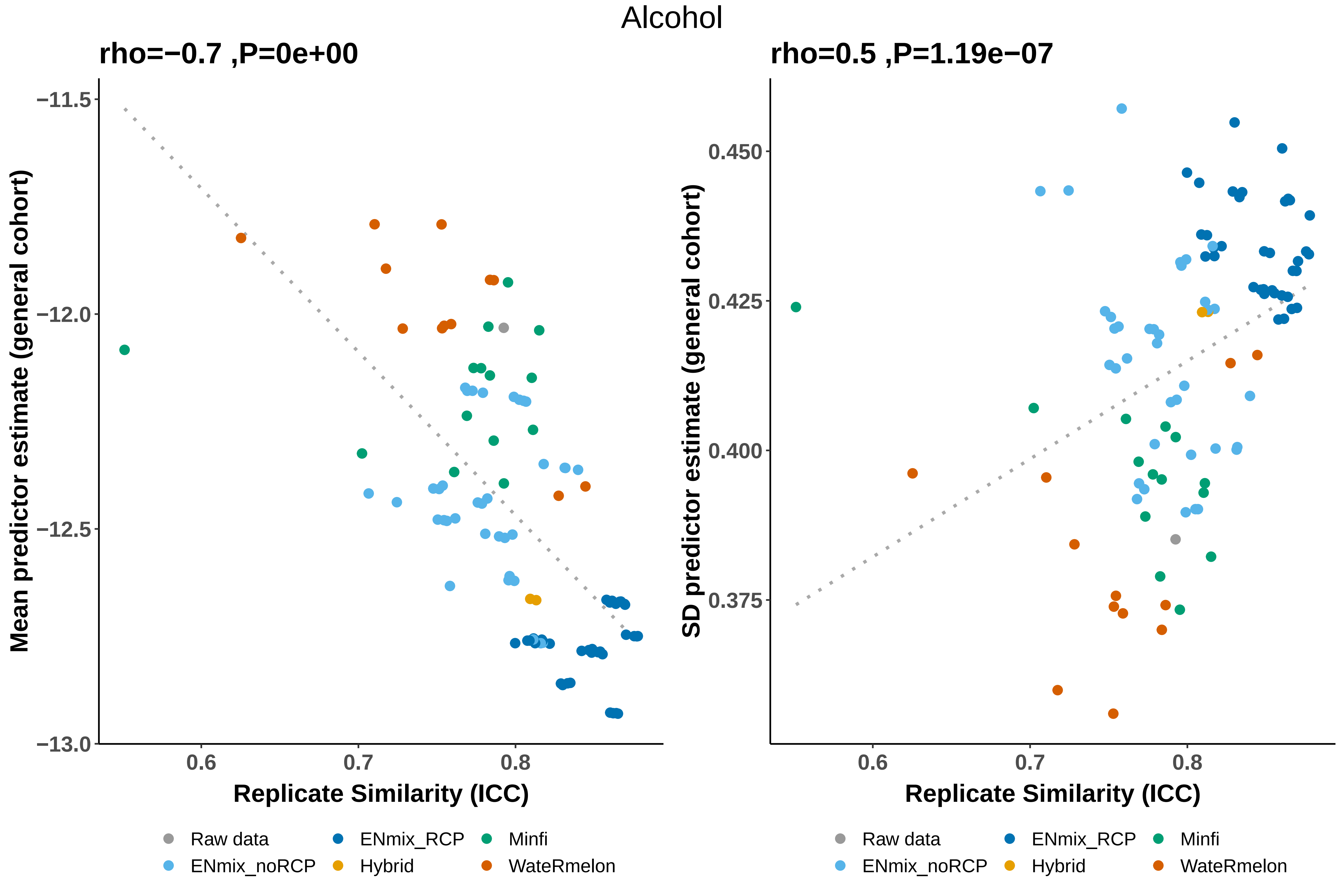

Alcohol
rho=−0.7 ,P=0e+00
rho=0.5 ,P=1.19e−07
Mean predictor estimate (general cohort)
SD predictor estimate (general cohort)
−11.5
−12.0
−12.5
−13.0
0.450
0.425
0.400
0.375
0.6
0.7
0.8
Replicate Similarity (ICC)
Raw data
ENmix_RCP
Minfi
ENmix_noRCP
Hybrid
WateRmelon

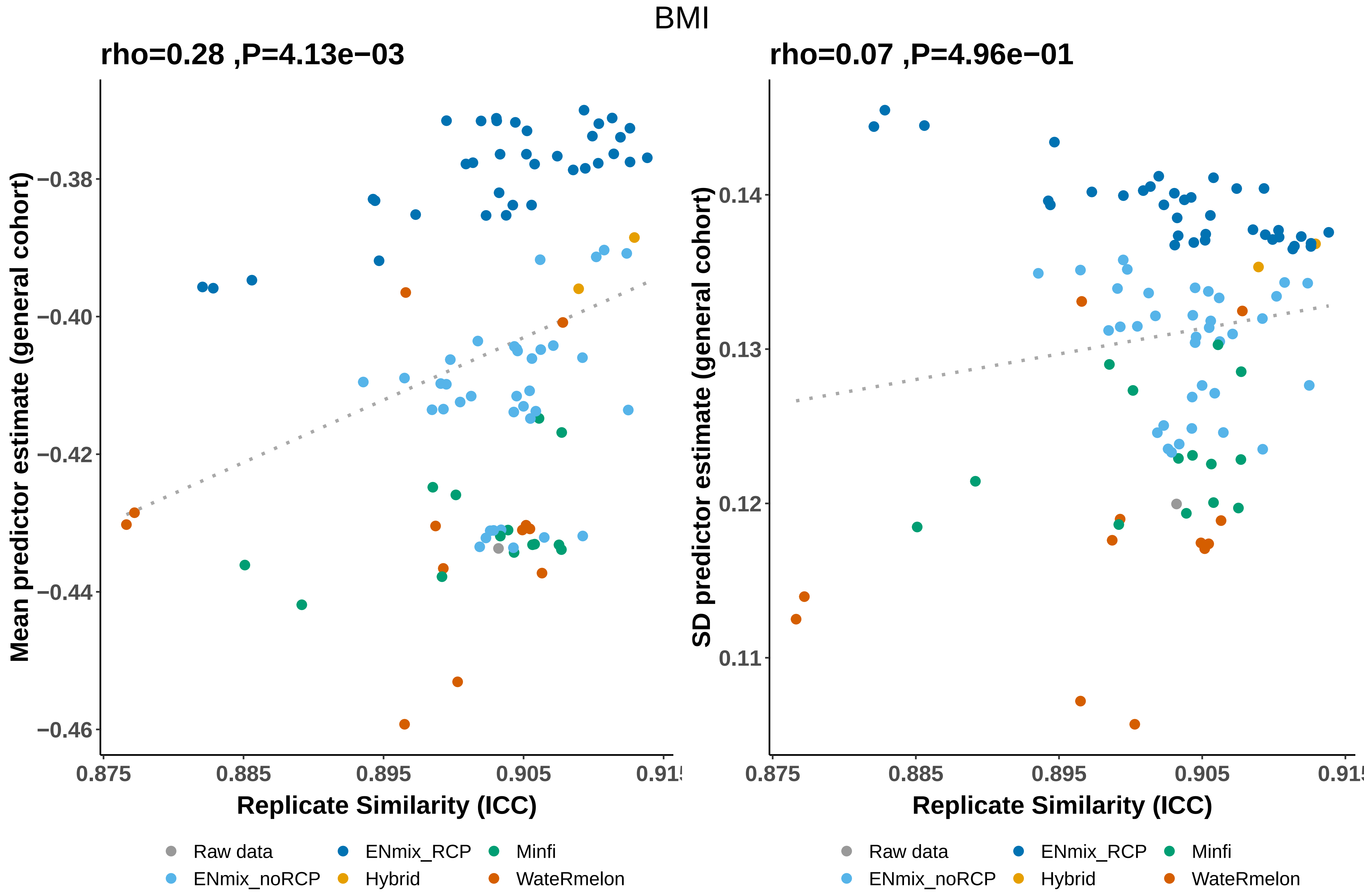

BMI
rho=0.28 ,P=4.13e−03
rho=0.07 ,P=4.96e−01
Mean predictor estimate (general cohort)
SD predictor estimate (general cohort)
−0.38
−0.40
−0.42
−0.44
−0.46
0.14
0.13
0.12
0.11
0.875
0.885
0.895
0.905
Replicate Similarity (ICC)
Raw data
ENmix_noRCP
ENmix_RCP
Hybrid
Minfi
WateRmelon

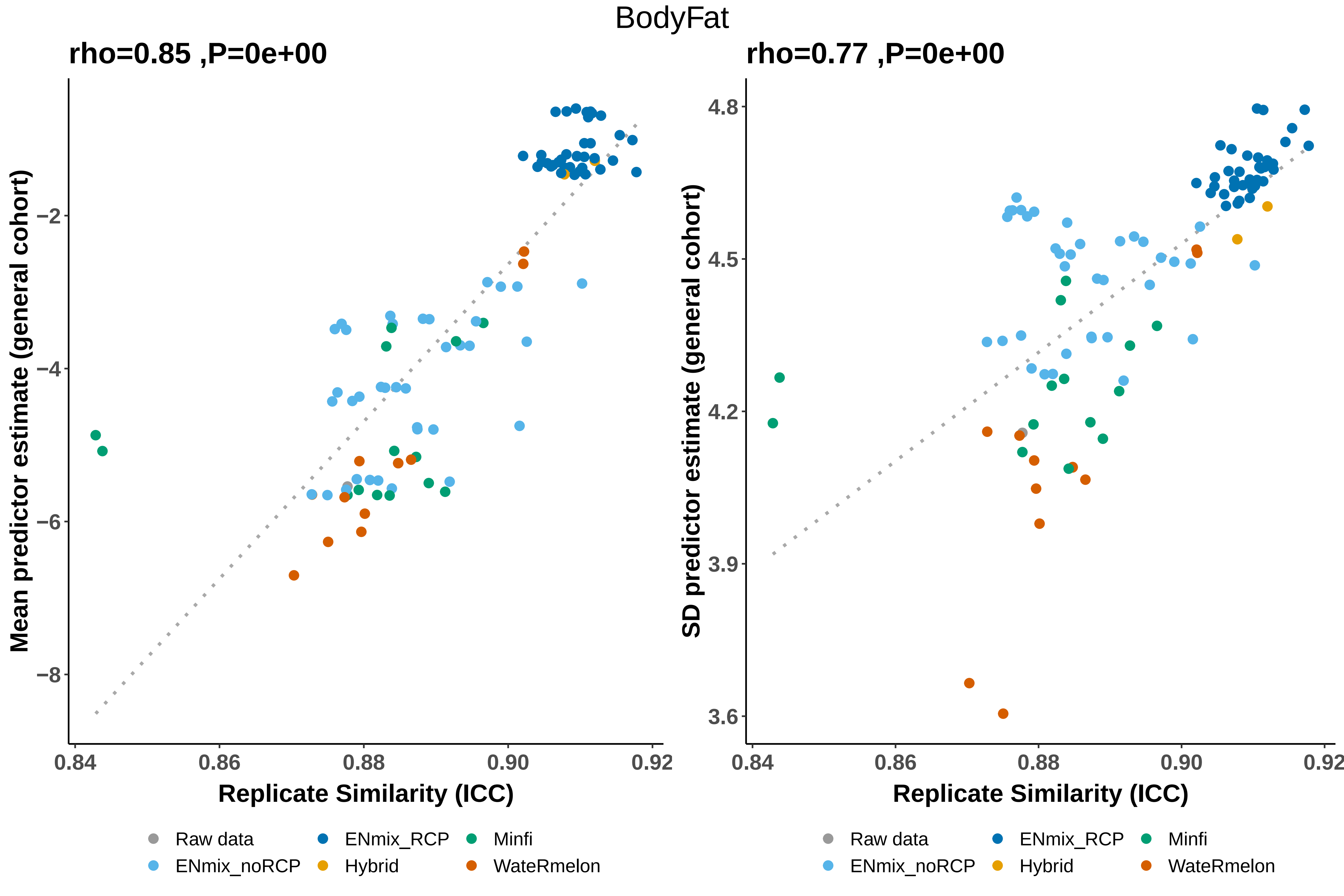
BodyFat
rho=0.85 ,P=0e+00
rho=0.77 ,P=0e+00
Mean predictor estimate (general cohort)
−2
−4
−6
−8
0.84
0.86
0.88
0.90
0.92
Replicate Similarity (ICC)
SD predictor estimate (general cohort)
4.8
4.5
4.2
3.9
3.6
0.84
0.86
0.88
0.90
0.92
Replicate Similarity (ICC)
Raw data
ENmix_RCP
Minfi
ENmix_noRCP
Hybrid
WateRmelon

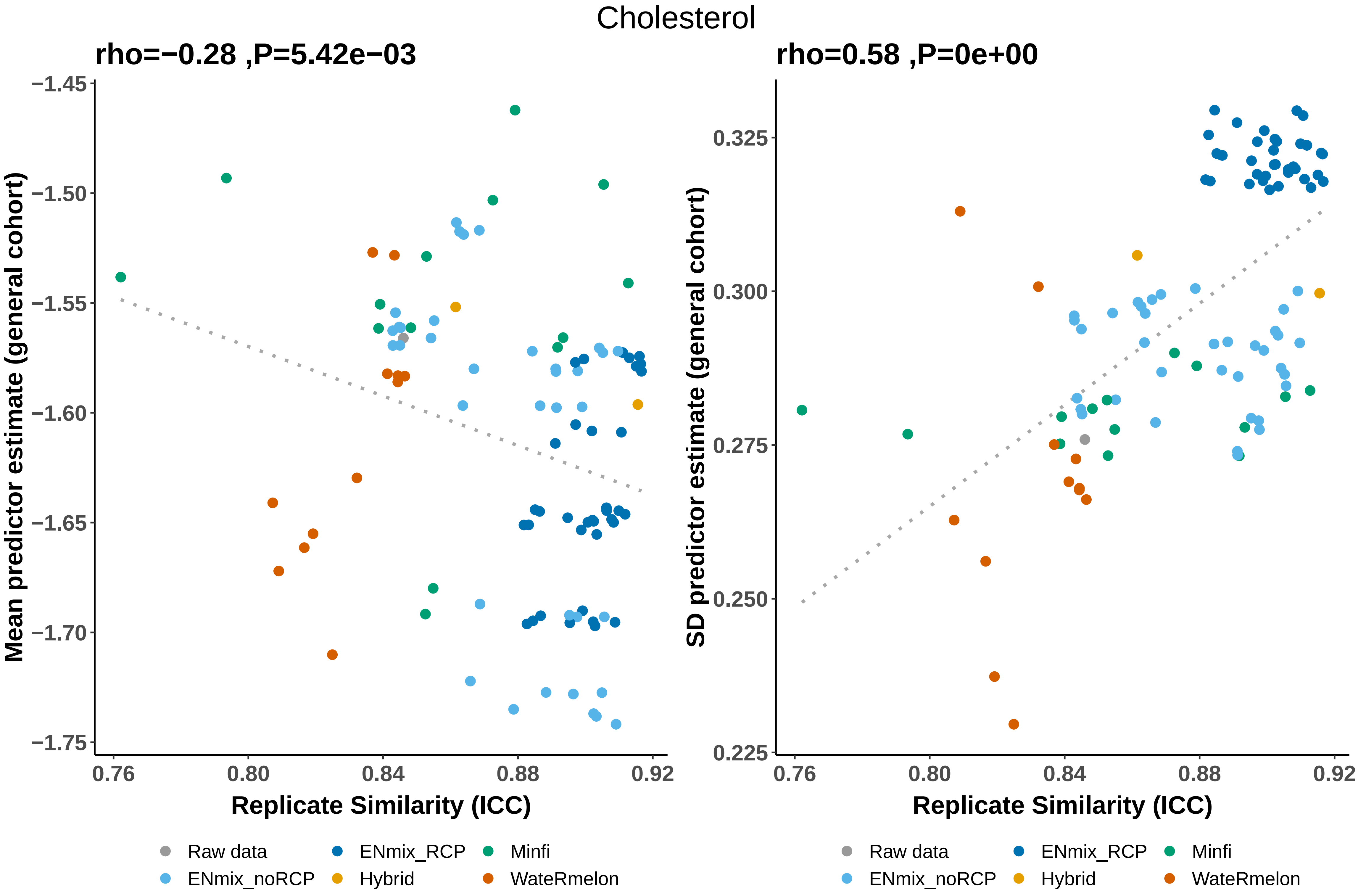

Cholesterol
rho=−0.28 ,P=5.42e−03
rho=0.58 ,P=0e+00
Mean predictor estimate (general cohort)
SD predictor estimate (general cohort)
Replicate Similarity (ICC)
Replicate Similarity (ICC)
−1.45
−1.50
−1.55
−1.60
−1.65
−1.70
−1.75
0.325
0.300
0.275
0.250
0.225
0.76
0.80
0.84
0.88
0.92
Raw data
ENmix_noRCP
ENmix_RCP
Hybrid
Minfi
WateRmelon

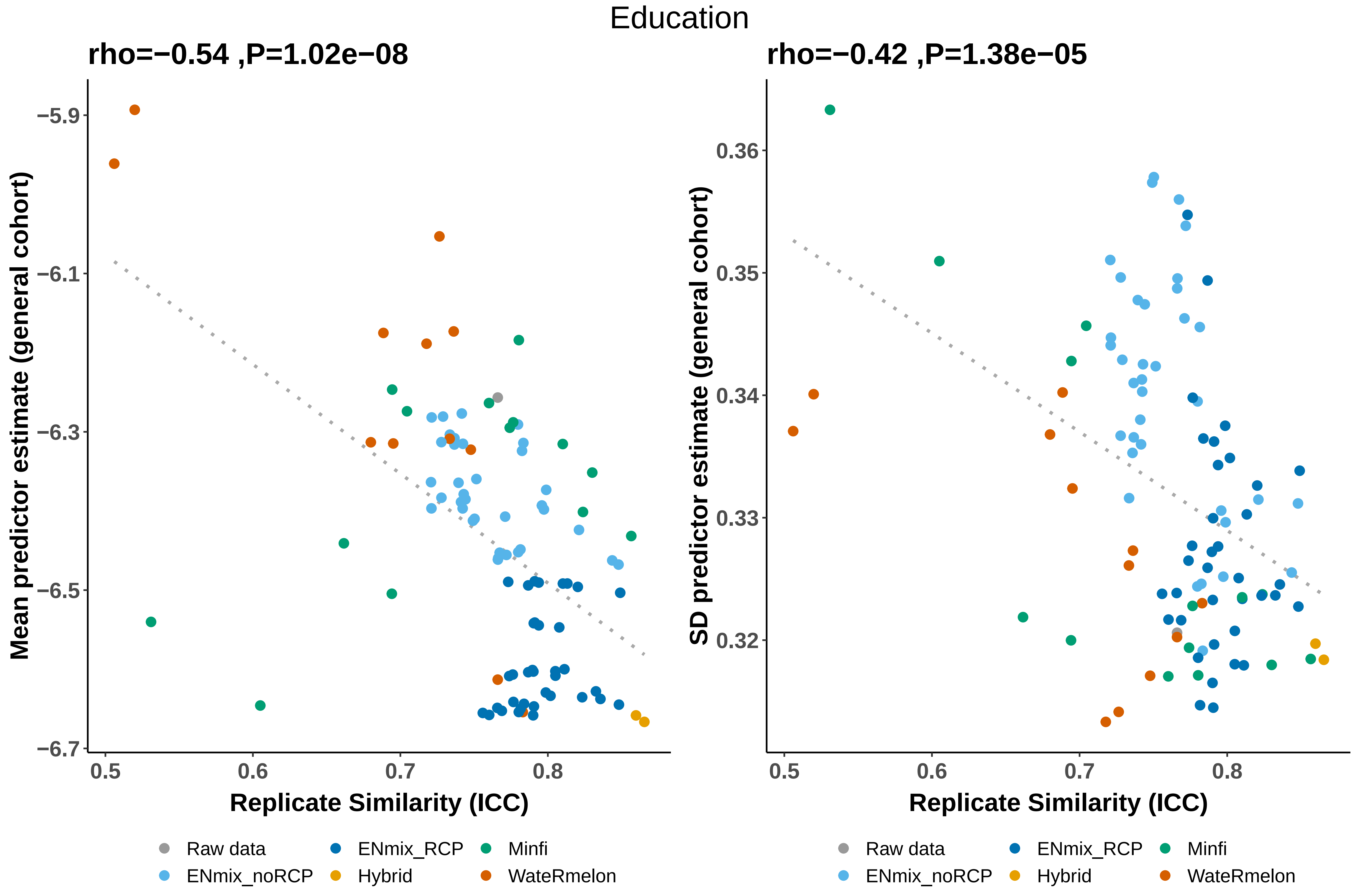

Education
rho=−0.54 ,P=1.02e−08
rho=−0.42 ,P=1.38e−05
Mean predictor estimate (general cohort)
SD predictor estimate (general cohort)
−5.9
−6.1
−6.3
−6.5
−6.7
0.36
0.35
0.34
0.33
0.32
0.5
0.6
0.7
0.8
Replicate Similarity (ICC)
Raw data
ENmix_RCP
Minfi
ENmix_noRCP
Hybrid
WateRmelon

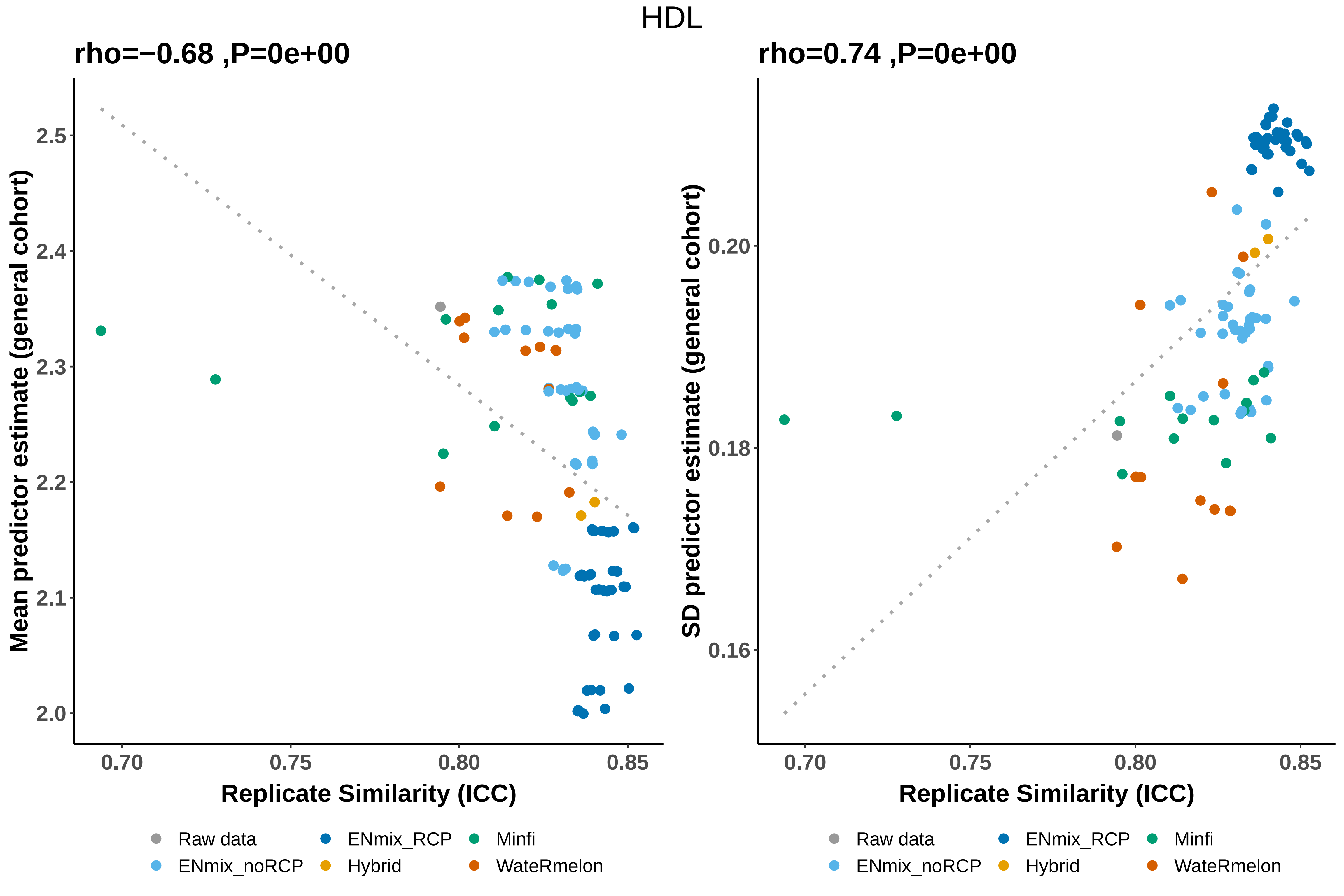
HDL
rho=−0.68 ,P=0e+00
rho=0.74 ,P=0e+00
Mean predictor estimate (general cohort)
SD predictor estimate (general cohort)
Replicate Similarity (ICC)
Replicate Similarity (ICC)
2.5
2.4
2.3
2.2
2.1
2.0
0.20
0.18
0.16
0.70
0.75
0.80
0.85
Raw data
ENmix_RCP
Minfi
ENmix_noRCP
Hybrid
WateRmelon

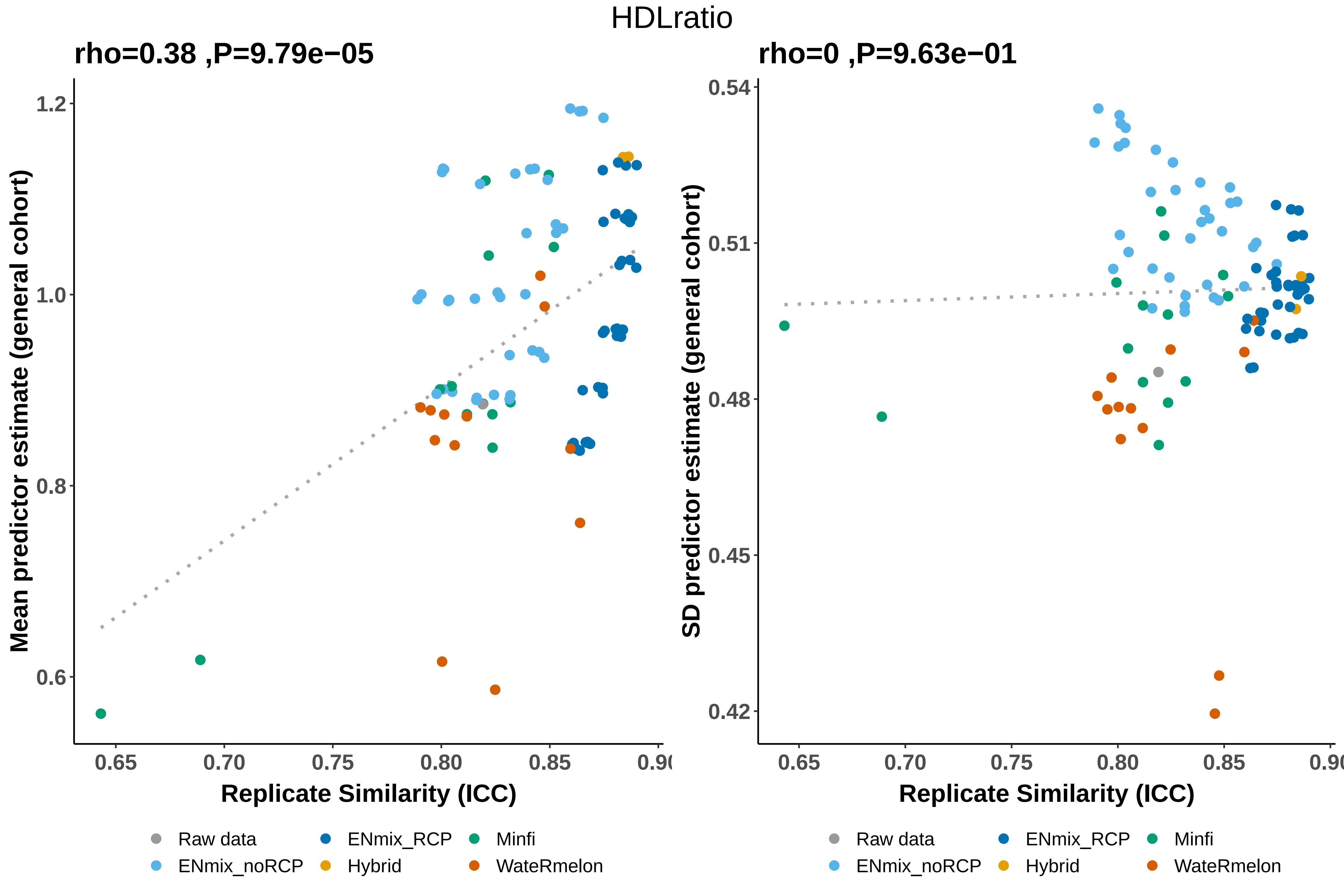

HDLratio
rho=0.38 ,P=9.79e−05
rho=0 ,P=9.63e−01
Mean predictor estimate (general cohort)
SD predictor estimate (general cohort)
1.2
1.0
0.8
0.6
0.54
0.51
0.48
0.45
0.42
0.65
0.70
0.75
0.80
0.85
0.90
Replicate Similarity (ICC)
Replicate Similarity (ICC)
Raw data
ENmix_noRCP
ENmix_RCP
Hybrid
Minfi
WateRmelon
Raw data
ENmix_noRCP
ENmix_RCP
Hybrid
Minfi
WateRmelon

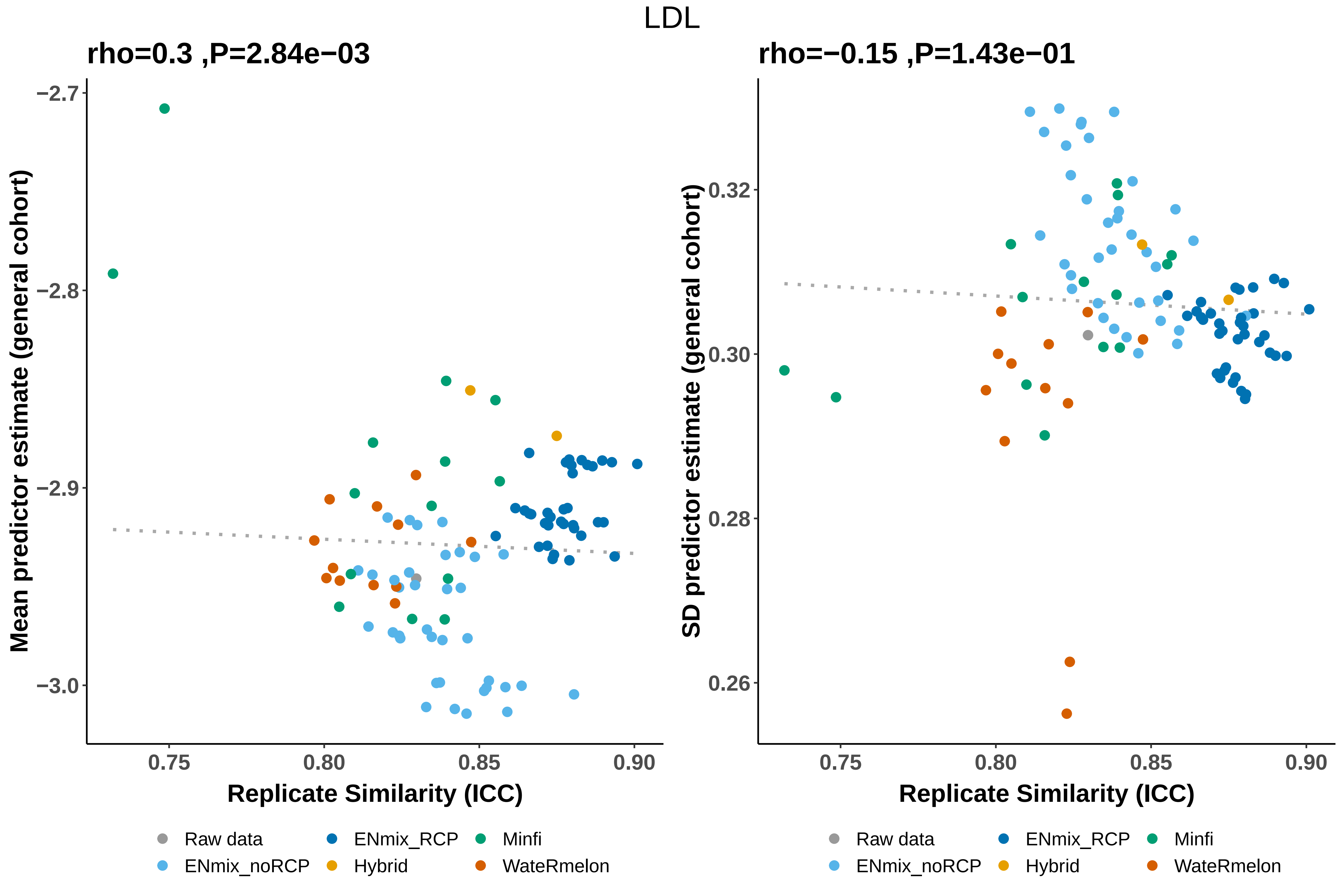

LDL
rho=0.3 ,P=2.84e−03
rho=−0.15 ,P=1.43e−01
Mean predictor estimate (general cohort)
SD predictor estimate (general cohort)
−2.7
−2.8
−2.9
−3.0
0.32
0.30
0.28
0.26
0.75
0.80
0.85
0.90
Replicate Similarity (ICC)
Raw data
ENmix_RCP
Minfi
ENmix_noRCP
Hybrid
WateRmelon

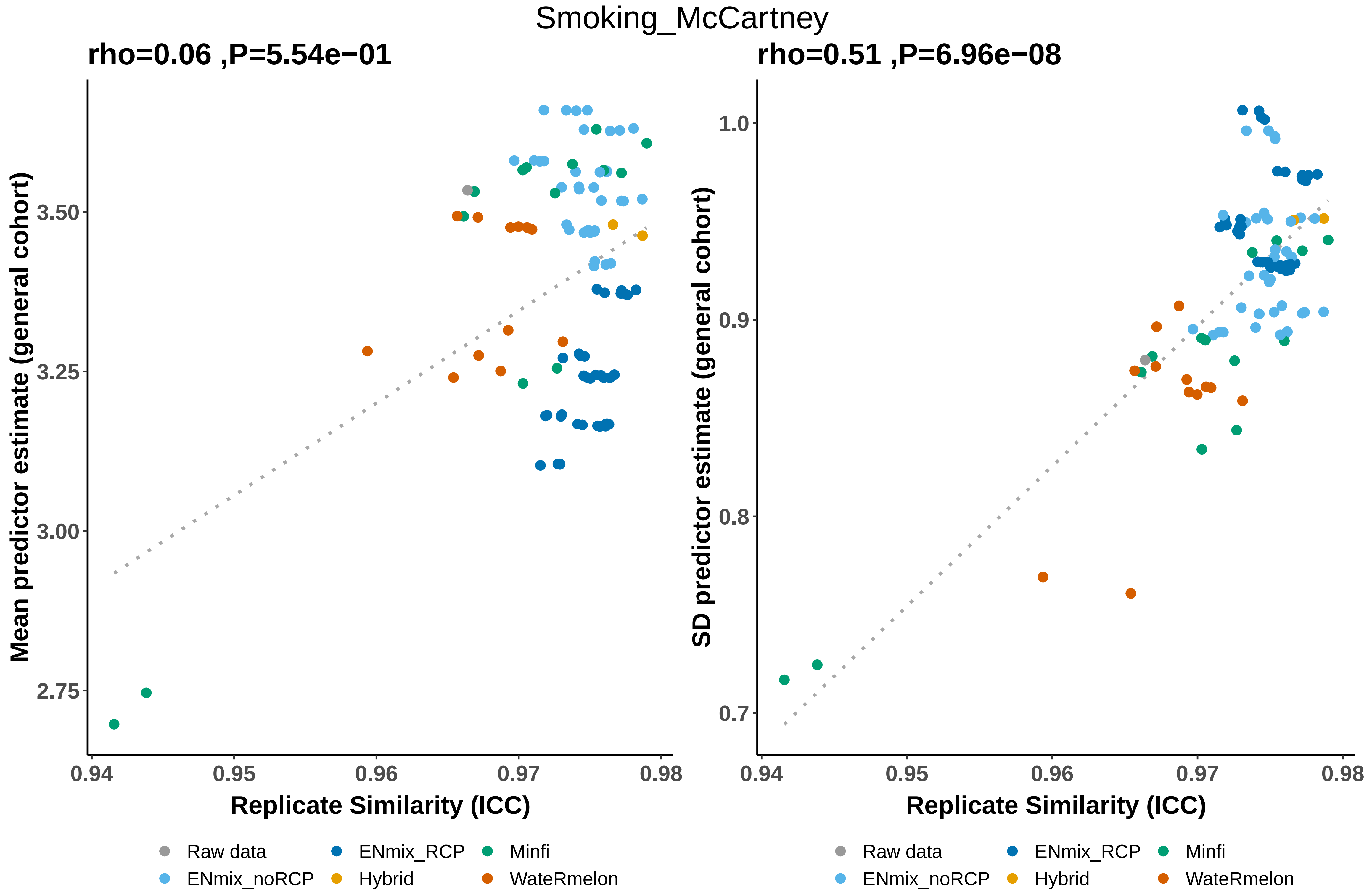
Smoking_McCartney
rho=0.06 ,P=5.54e−01
rho=0.51 ,P=6.96e−08
Mean predictor estimate (general cohort)
SD predictor estimate (general cohort)
3.50
3.25
3.00
2.75
1.0
0.9
0.8
0.7
0.94
0.95
0.96
0.97
0.98
Replicate Similarity (ICC)
Replicate Similarity (ICC)
Raw data
ENmix_noRCP
ENmix_RCP
Hybrid
Minfi
WateRmelon
Raw data
ENmix_noRCP
ENmix_RCP
Hybrid
Minfi
WateRmelon

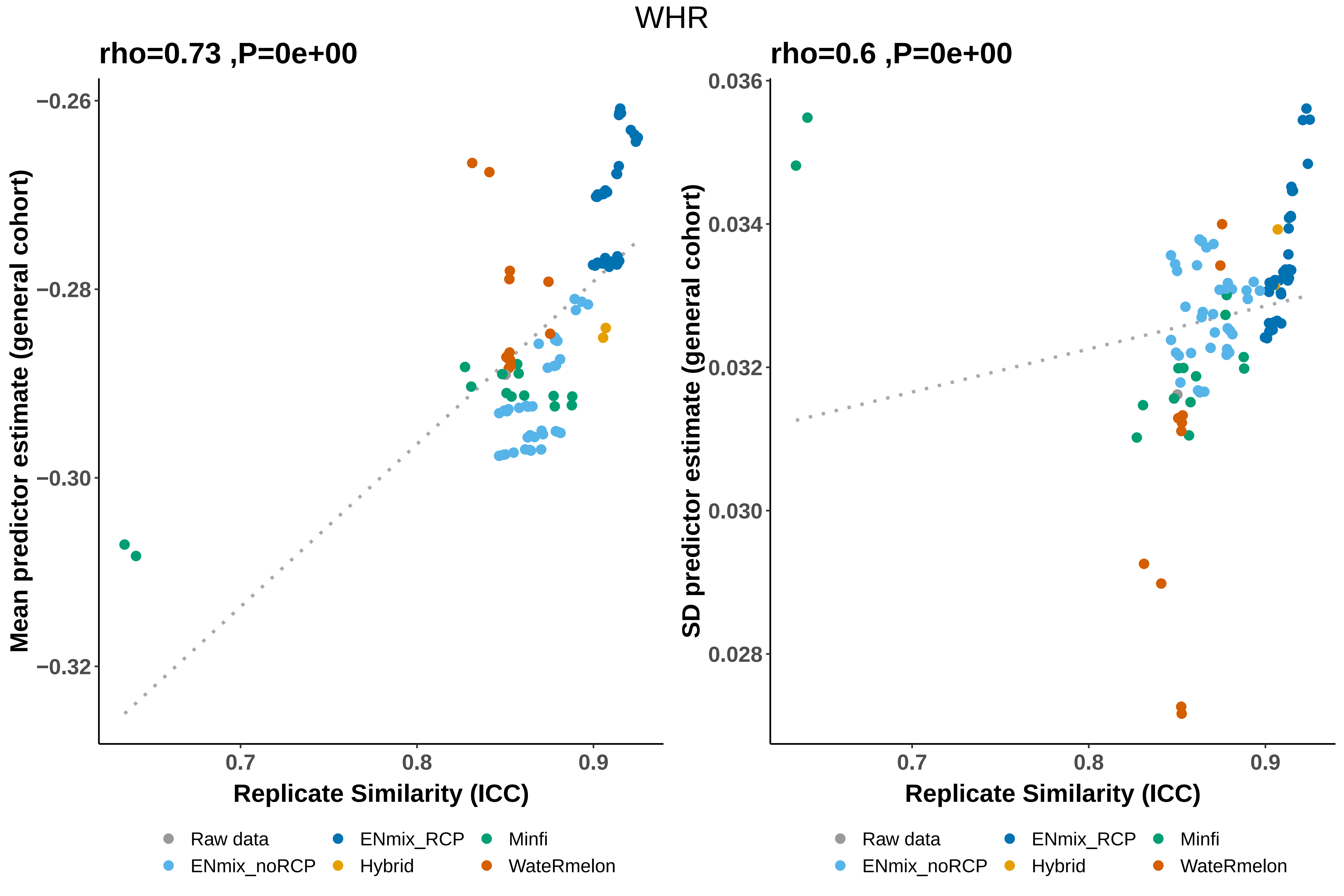
WHR
rho=0.73 ,P=0e+00
rho=0.6 ,P=0e+00
Mean predictor estimate (general cohort)
SD predictor estimate (general cohort)
−0.26
−0.28
−0.30
−0.32
0.036
0.034
0.032
0.030
0.028
0.7
0.8
0.9
Replicate Similarity (ICC)
Raw data
ENmix_noRCP
ENmix_RCP
Hybrid
Minfi
WateRmelon

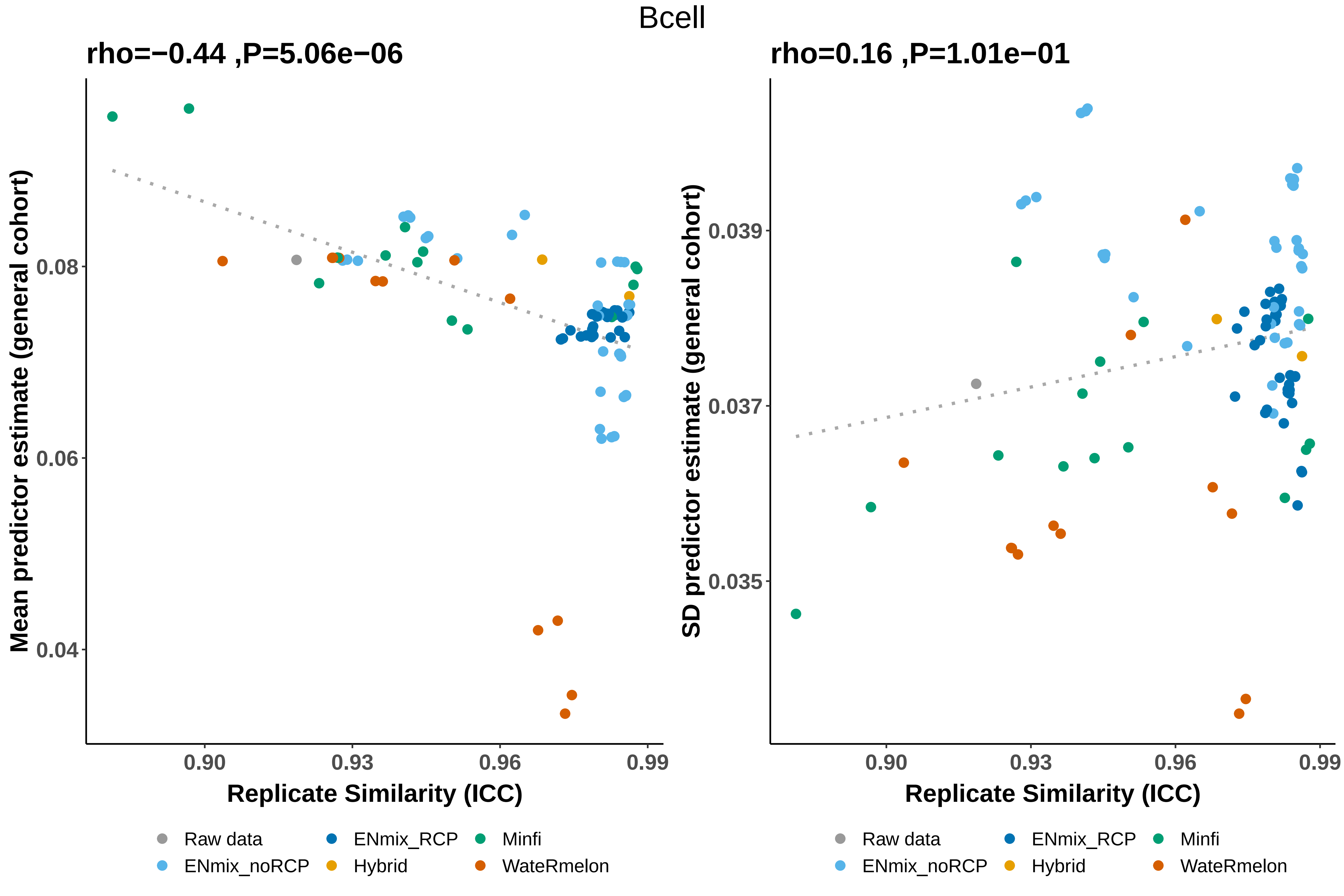

Bcell
rho=−0.44 ,P=5.06e−06
Mean predictor estimate (general cohort)
0.08
0.06
0.04
0.90
0.93
0.96
0.99
Replicate Similarity (ICC)
Raw data
ENmix_noRCP
ENmix_RCP
Hybrid
Minfi
WateRmelon
rho=0.16 ,P=1.01e−01
SD predictor estimate (general cohort)
0.039
0.037
0.035
0.90
0.93
0.96
0.99
Replicate Similarity (ICC)
Raw data
ENmix_noRCP
ENmix_RCP
Hybrid
Minfi
WateRmelon

CD4T

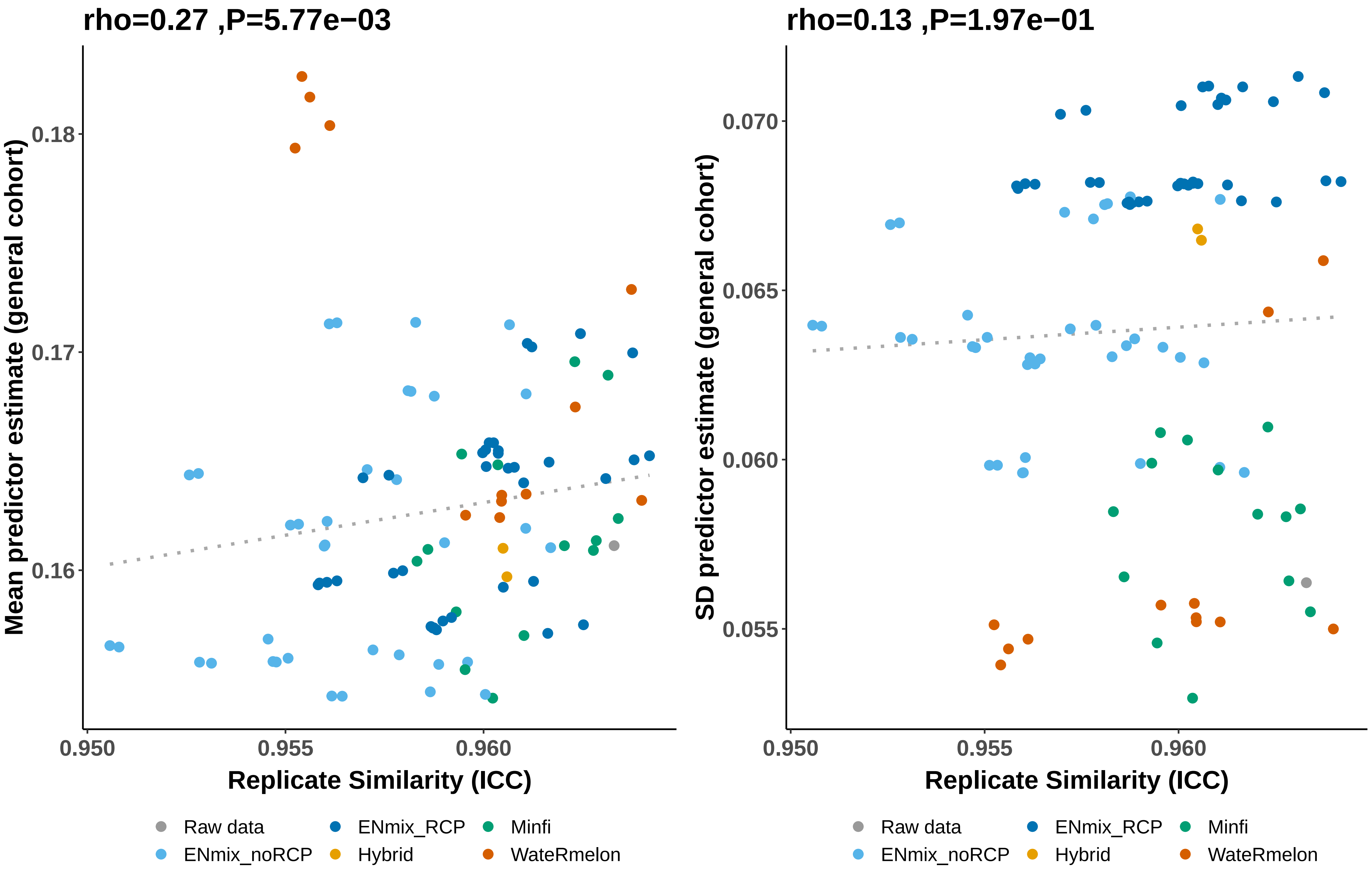
rho=0.27 ,P=5.77e−03
Mean predictor estimate (general cohort)
0.18
0.17
0.16
0.950
0.955
0.960
Replicate Similarity (ICC)
rho=0.13 ,P=1.97e−01
SD predictor estimate (general cohort)
0.070
0.065
0.060
0.055
0.950
0.955
0.960
Replicate Similarity (ICC)
Raw data
ENmix_noRCP
ENmix_RCP
Hybrid
Minfi
WateRmelon

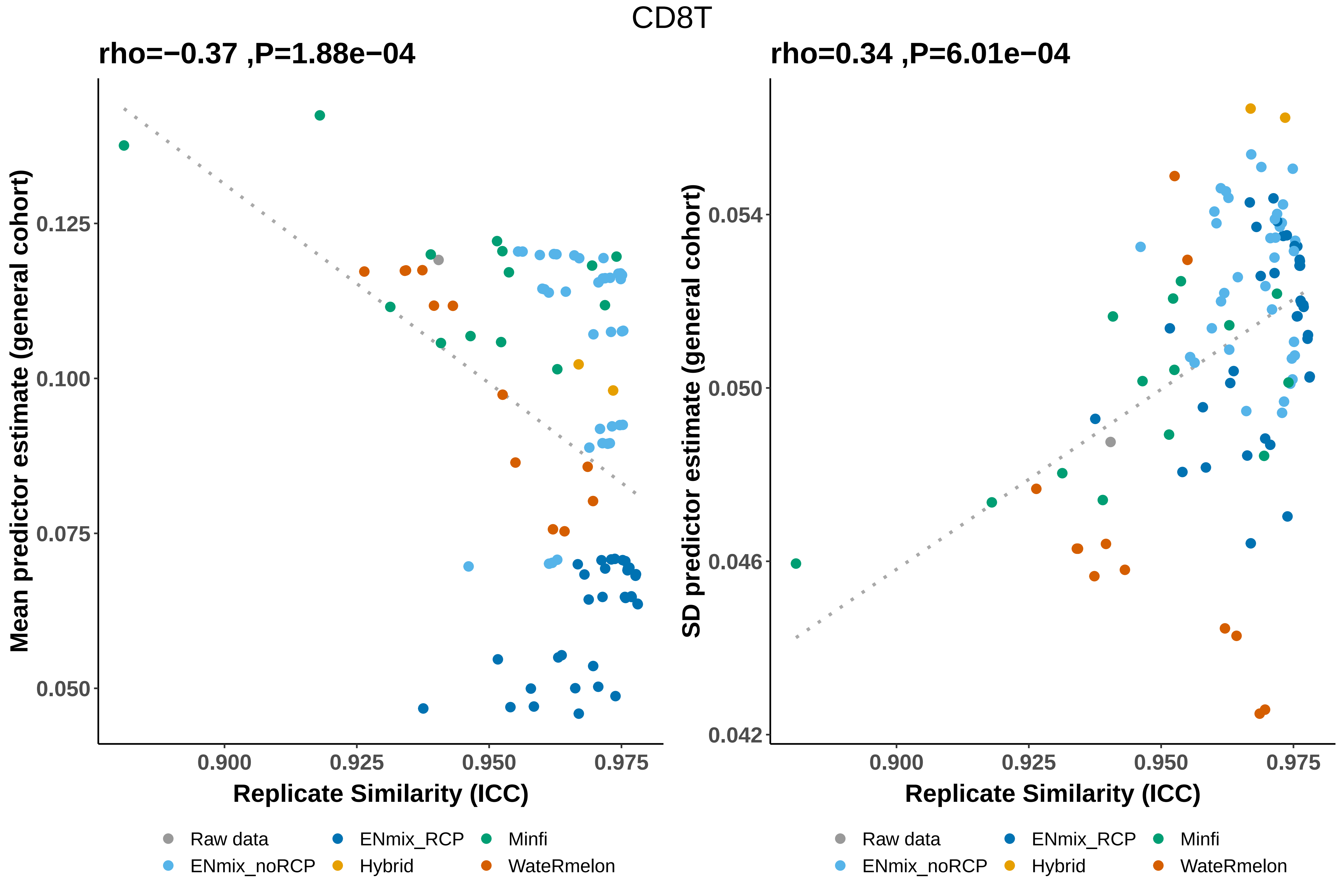

CD8T
rho=−0.37 ,P=1.88e−04
Mean predictor estimate (general cohort)
0.125
0.100
0.075
0.050
0.900
0.925
0.950
0.975
Replicate Similarity (ICC)
rho=0.34 ,P=6.01e−04
SD predictor estimate (general cohort)
0.054
0.050
0.046
0.042
0.900
0.925
0.950
0.975
Replicate Similarity (ICC)
Raw data
ENmix_RCP
Minfi
ENmix_noRCP
Hybrid
WateRmelon

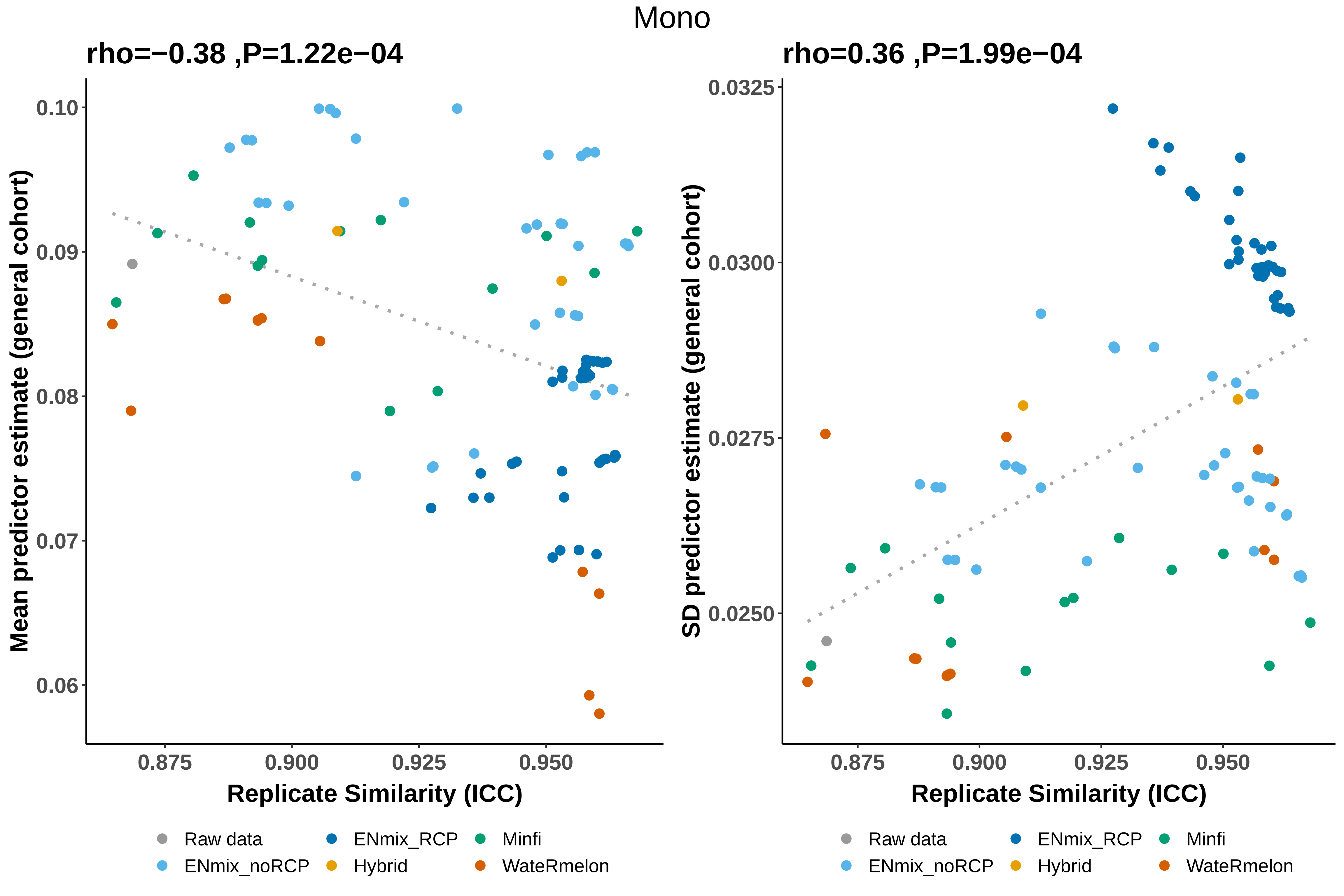

Mono
rho=−0.38 ,P=1.22e−04
Mean predictor estimate (general cohort)
0.10
0.09
0.08
0.07
0.06
0.875
0.900
0.925
0.950
Replicate Similarity (ICC)
rho=0.36 ,P=1.99e−04
SD predictor estimate (general cohort)
0.0325
0.0300
0.0275
0.0250
0.875
0.900
0.925
0.950
Replicate Similarity (ICC)
Raw data
ENmix_noRCP
ENmix_RCP
Hybrid
Minfi
WateRmelon

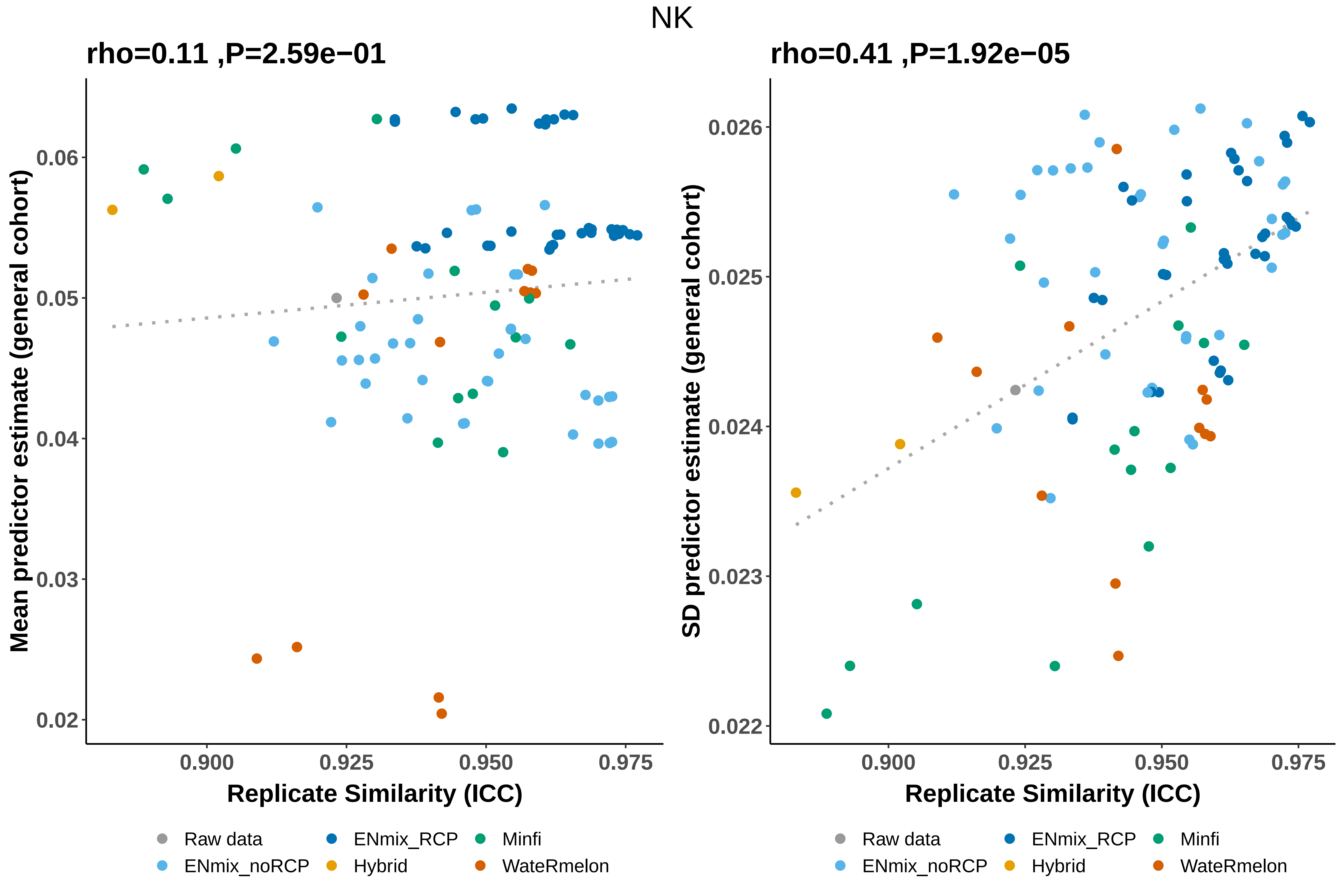

NK
rho=0.11 ,P=2.59e−01
rho=0.41 ,P=1.92e−05
Mean predictor estimate (general cohort)
SD predictor estimate (general cohort)
0.02
0.03
0.04
0.05
0.06
0.022
0.023
0.024
0.025
0.026
0.900
0.925
0.950
0.975
Replicate Similarity (ICC)
Raw data
ENmix_noRCP
ENmix_RCP
Hybrid
Minfi
WateRmelon

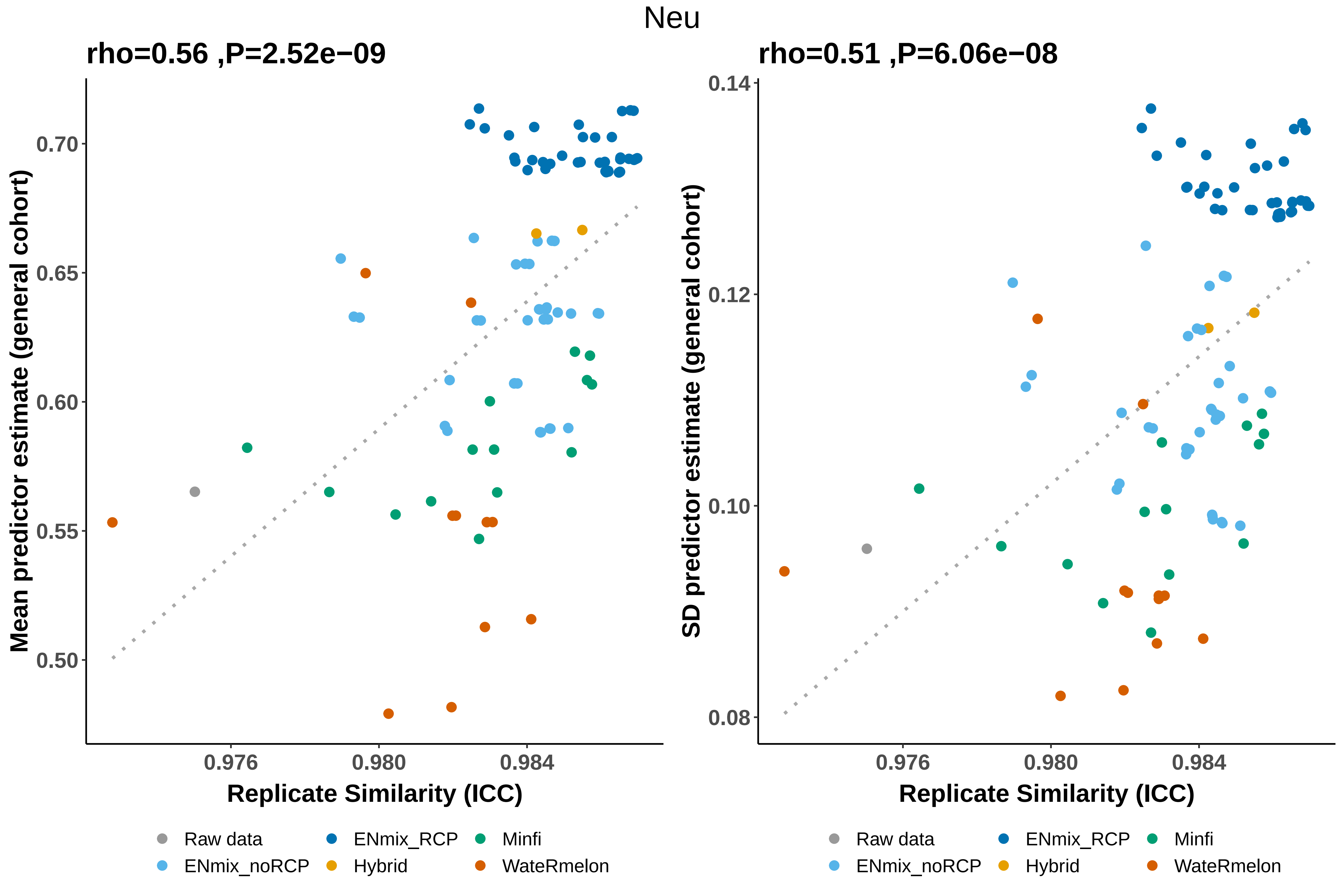
Neu
rho=0.56 ,P=2.52e−09
Mean predictor estimate (general cohort)
0.70
0.65
0.60
0.55
0.50
0.976
0.980
0.984
Replicate Similarity (ICC)
rho=0.51 ,P=6.06e−08
SD predictor estimate (general cohort)
0.14
0.12
0.10
0.08
0.976
0.980
0.984
Replicate Similarity (ICC)
Raw data
ENmix_noRCP
ENmix_RCP
Hybrid
Minfi
WateRmelon

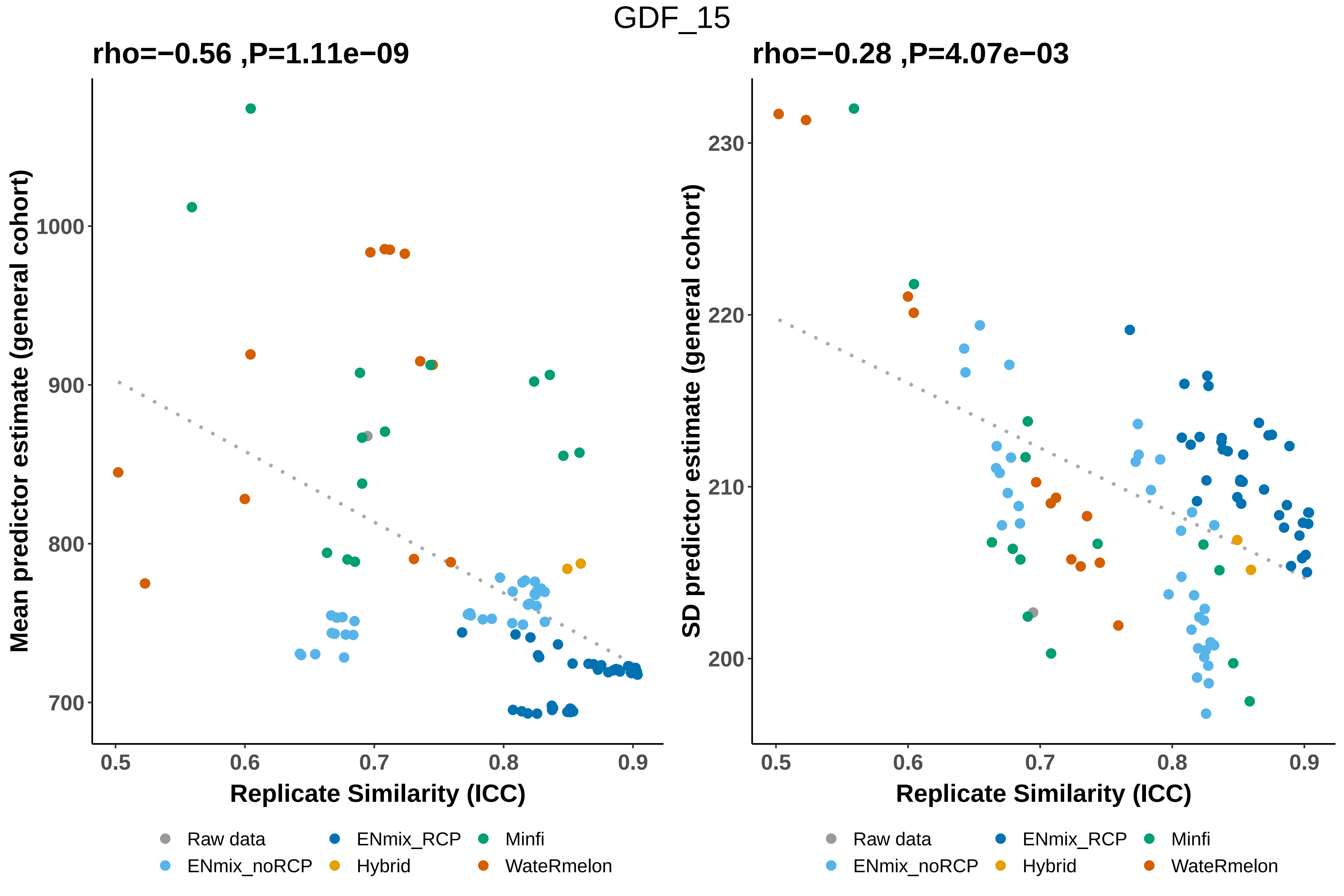
GDF_15
rho=−0.56 ,P=1.11e−09
rho=−0.28 ,P=4.07e−03
Mean predictor estimate (general cohort)
SD predictor estimate (general cohort)
1000
900
800
700
230
220
210
200
0.5
0.6
0.7
0.8
0.9
Replicate Similarity (ICC)
Raw data
ENmix_noRCP
ENmix_RCP
Hybrid
Minfi
WateRmelon

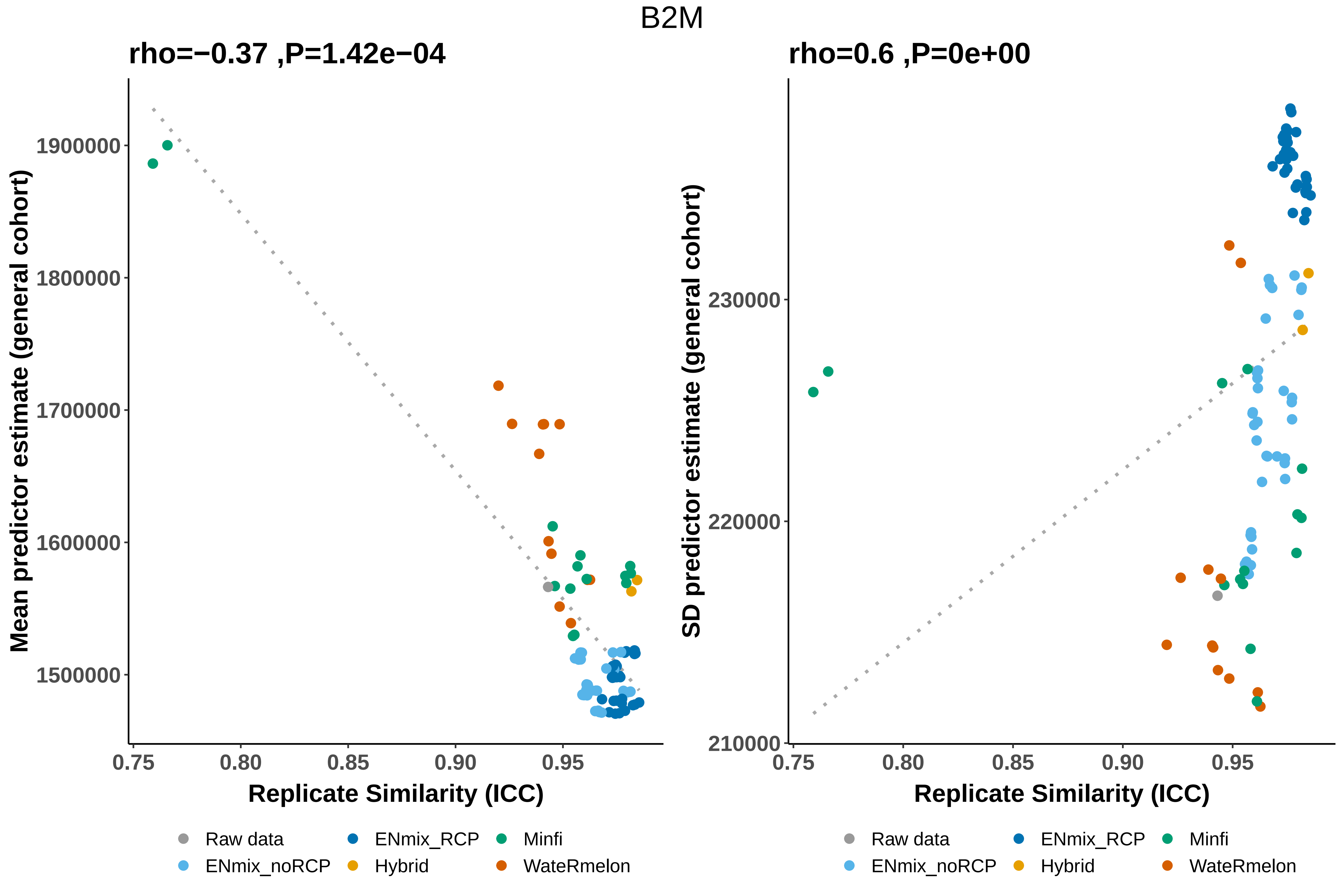
B2M
rho=−0.37 ,P=1.42e−04
Mean predictor estimate (general cohort)
1900000
1800000
1700000
1600000
1500000
0.75
0.80
0.85
0.90
0.95
Replicate Similarity (ICC)
rho=0.6 ,P=0e+00
SD predictor estimate (general cohort)
230000
220000
210000
Raw data
ENmix_RCP
Minfi
ENmix_noRCP
Hybrid
WateRmelon

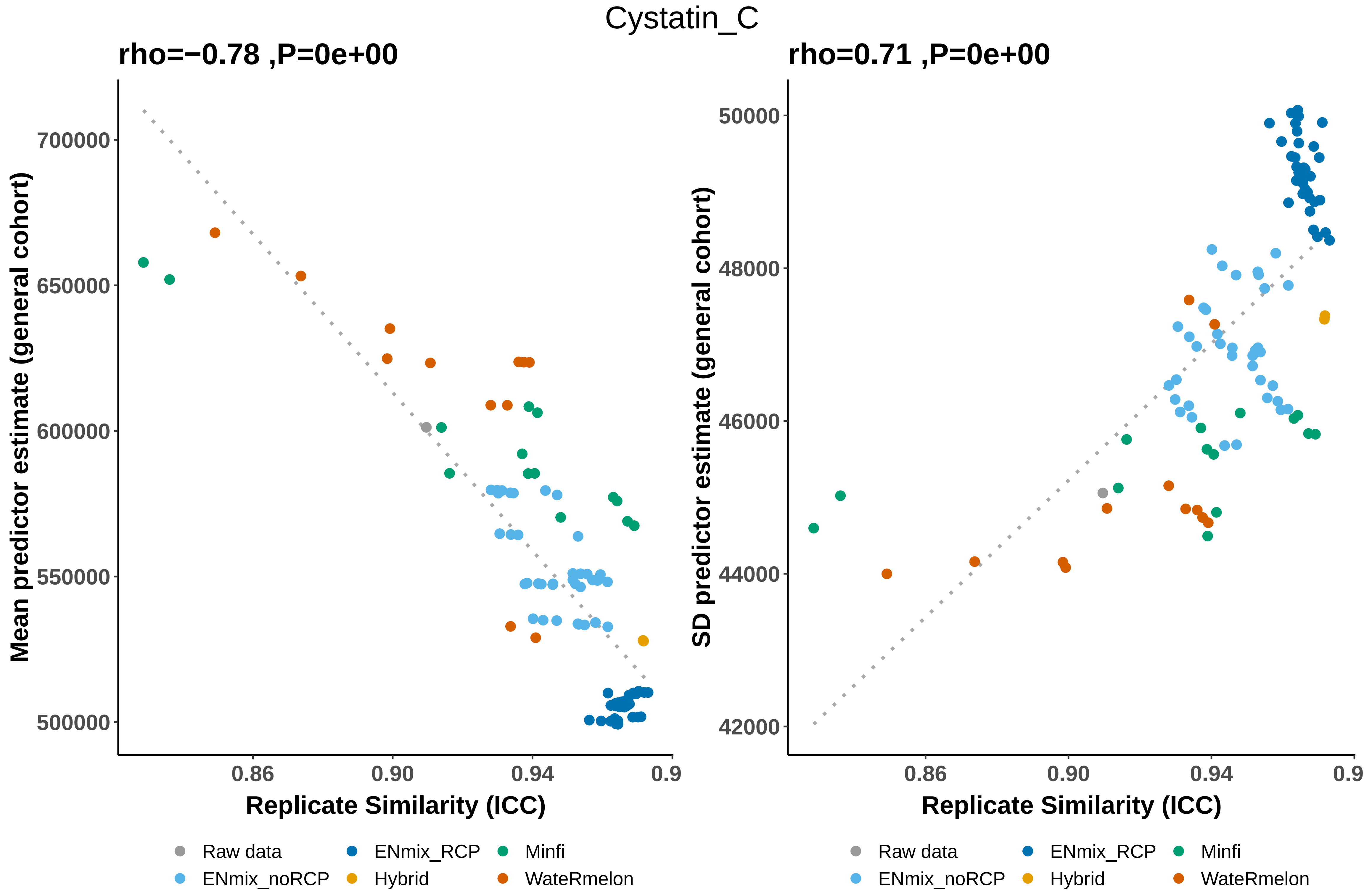

Cystatin_C
rho=−0.78 ,P=0e+00
rho=0.71 ,P=0e+00
Mean predictor estimate (general cohort)
SD predictor estimate (general cohort)
700000
650000
600000
550000
500000
50000
48000
46000
44000
42000
0.86
0.90
0.94
0.9
Replicate Similarity (ICC)
Raw data
ENmix_RCP
Minfi
ENmix_noRCP
Hybrid
WateRmelon

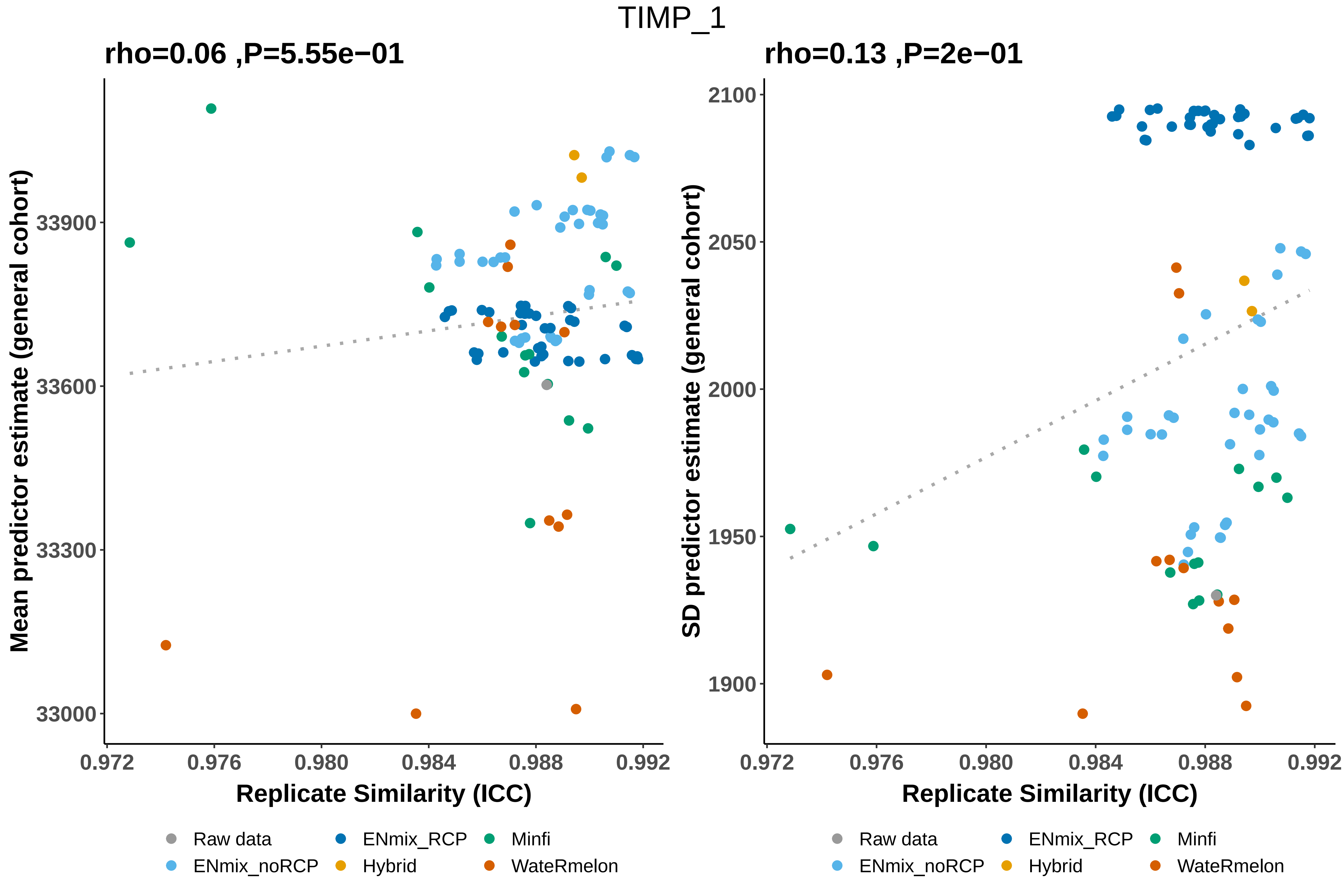

TIMP_1
rho=0.06 ,P=5.55e−01
rho=0.13 ,P=2e−01
Mean predictor estimate (general cohort)
SD predictor estimate (general cohort)
33000
33300
33600
33900
1900
1950
2000
2050
2100
0.972
0.976
0.980
0.984
0.988
0.992
Replicate Similarity (ICC)
Raw data
ENmix_RCP
Minfi
ENmix_noRCP
Hybrid
WateRmelon

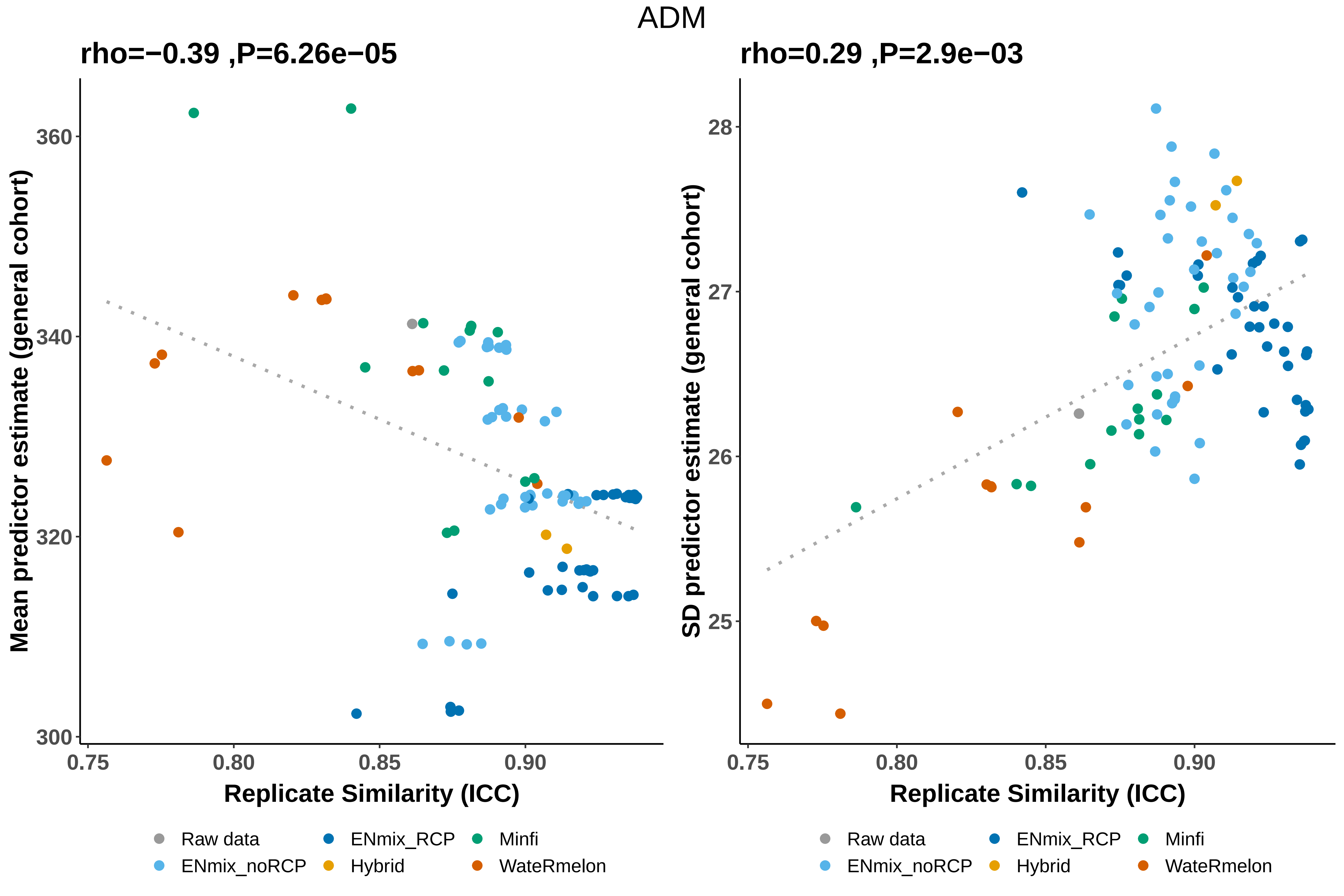
ADM
rho=−0.39 ,P=6.26e−05
Mean predictor estimate (general cohort)
360
340
320
300
0.75
0.80
0.85
0.90
Replicate Similarity (ICC)
rho=0.29 ,P=2.9e−03
SD predictor estimate (general cohort)
28
27
26
25
0.75
0.80
0.85
0.90
Replicate Similarity (ICC)
Raw data
ENmix_noRCP
ENmix_RCP
Hybrid
Minfi
WateRmelon

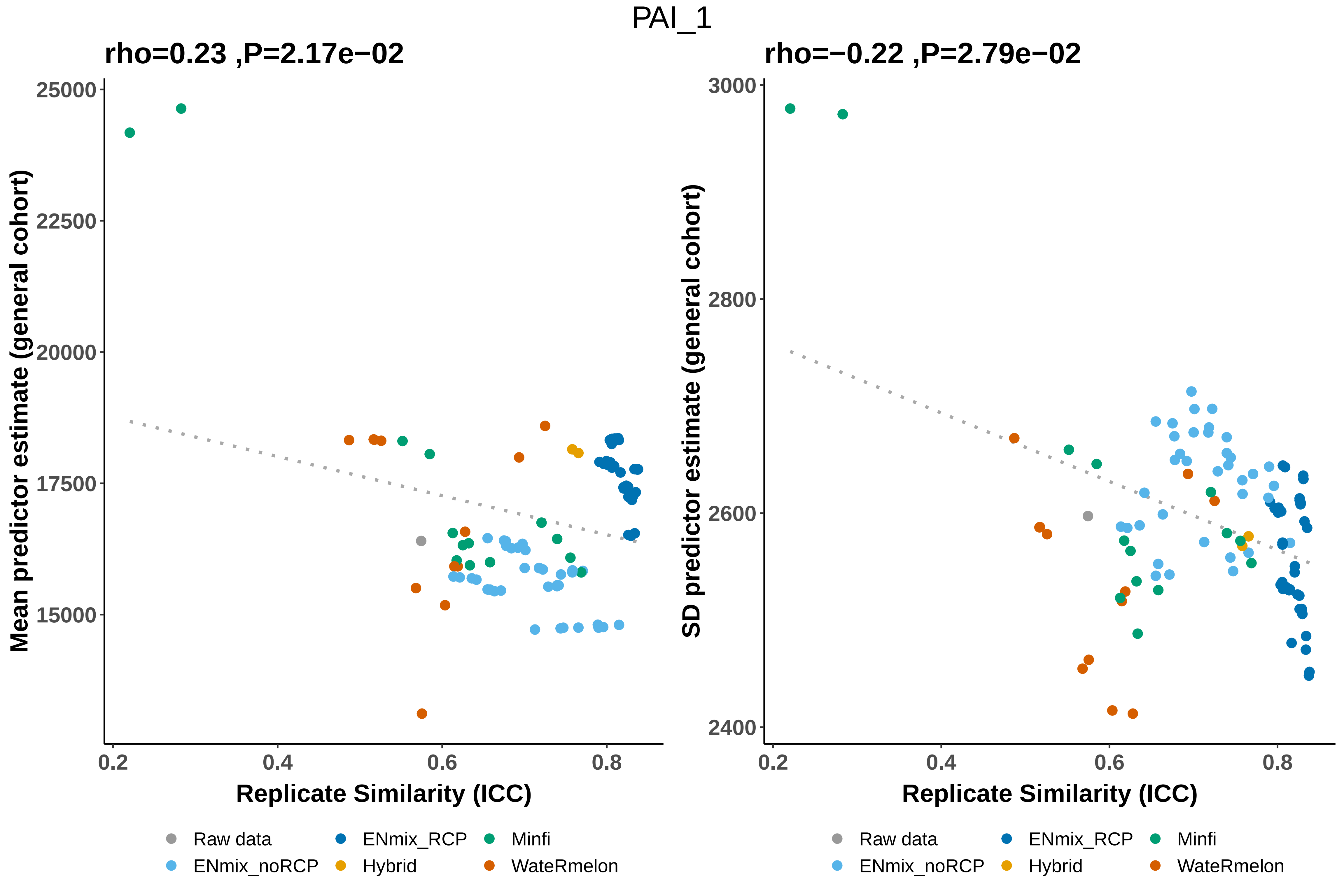
PAI_1
rho=0.23 ,P=2.17e−02
Mean predictor estimate (general cohort)
25000
22500
20000
17500
15000
0.2
0.4
0.6
0.8
Replicate Similarity (ICC)
rho=−0.22 ,P=2.79e−02
SD predictor estimate (general cohort)
3000
2800
2600
2400
0.2
0.4
0.6
0.8
Replicate Similarity (ICC)
Raw data
ENmix_RCP
Minfi
ENmix_noRCP
Hybrid
WateRmelon

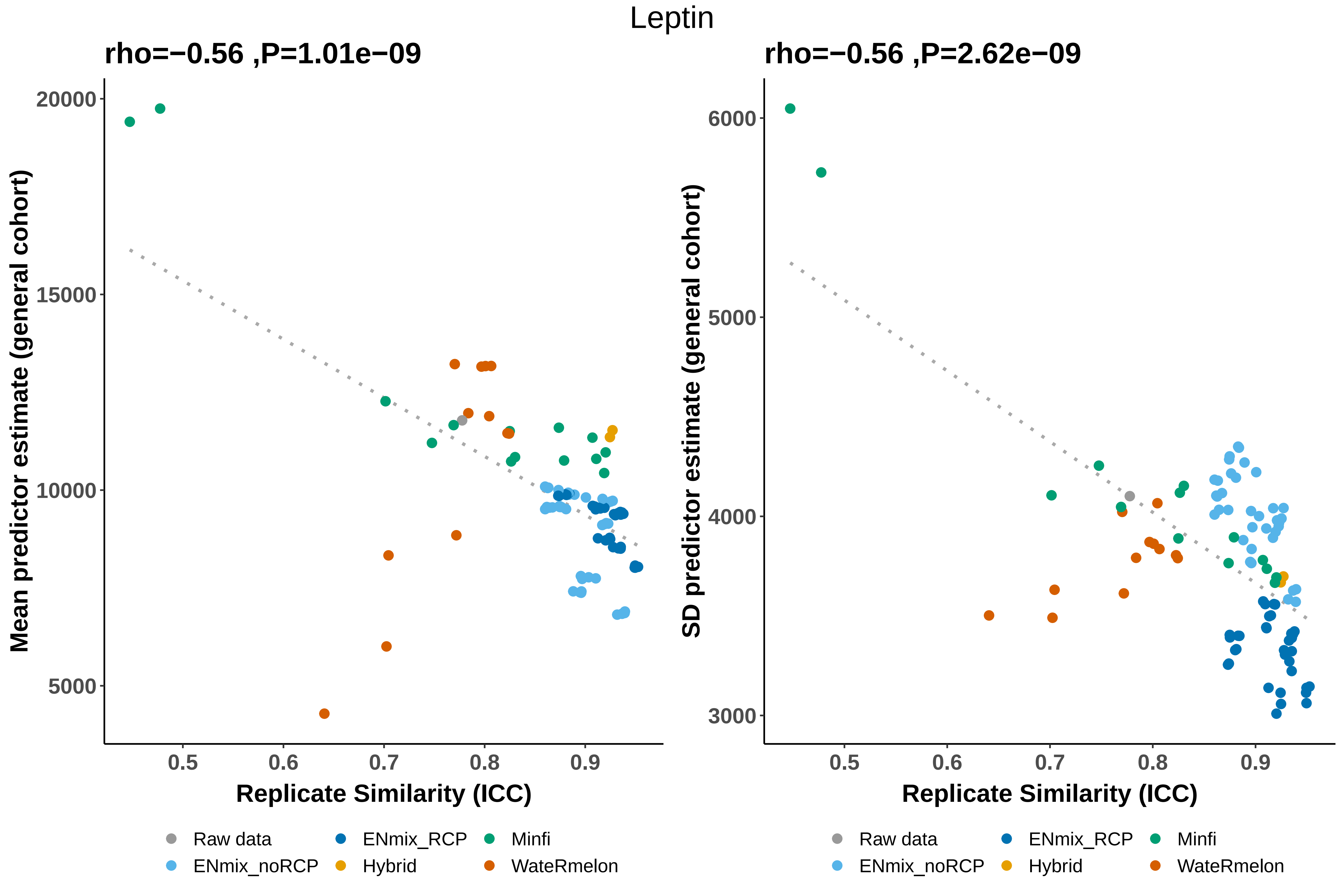
Leptin
rho=−0.56 ,P=1.01e−09
rho=−0.56 ,P=2.62e−09
Mean predictor estimate (general cohort)
SD predictor estimate (general cohort)
20000
15000
10000
5000
6000
5000
4000
3000
0.5
0.6
0.7
0.8
0.9
Replicate Similarity (ICC)
Replicate Similarity (ICC)
Raw data
ENmix_RCP
Minfi
ENmix_noRCP
Hybrid
WateRmelon

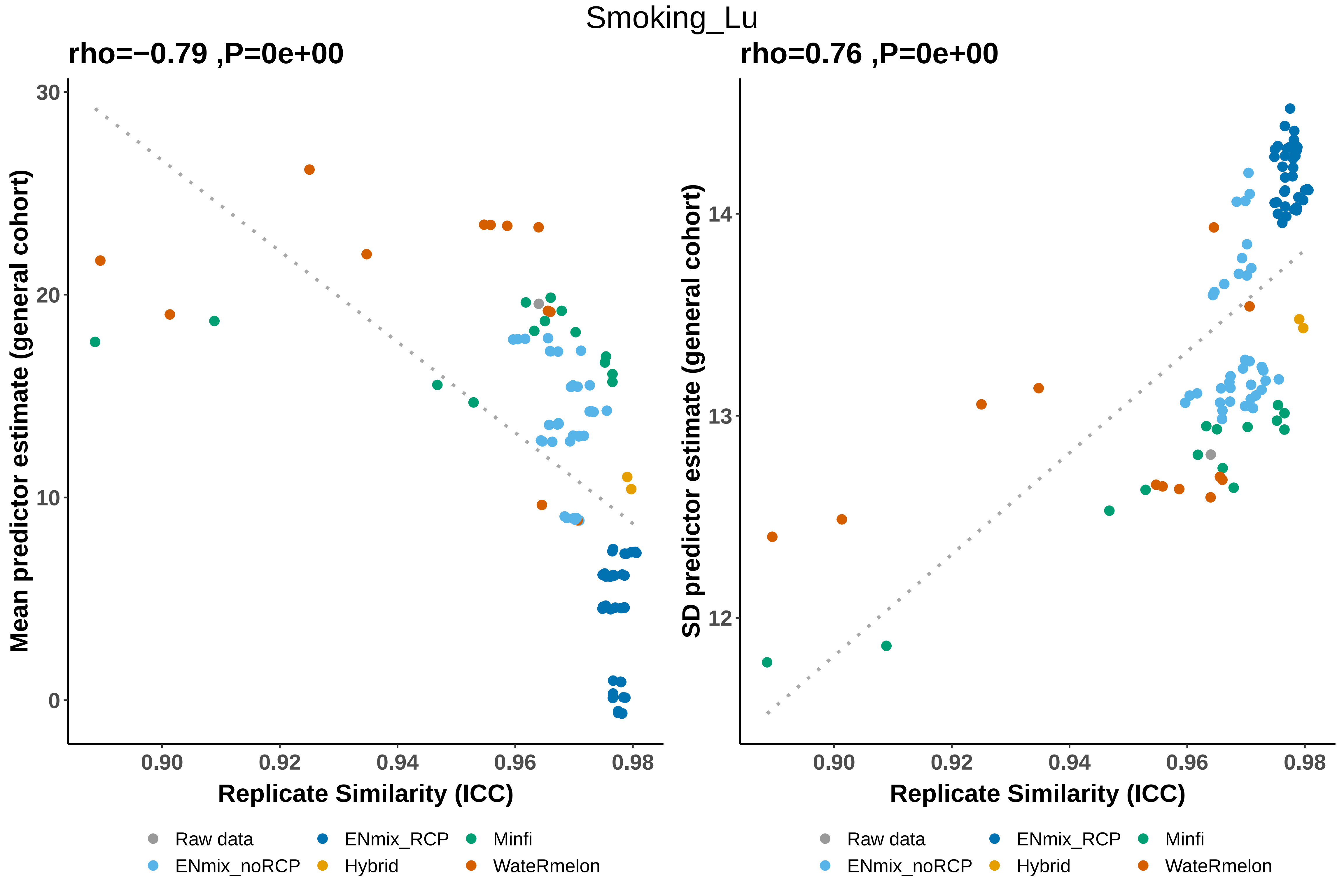
Smoking_Lu
rho=−0.79 ,P=0e+00
rho=0.76 ,P=0e+00
Mean predictor estimate (general cohort)
SD predictor estimate (general cohort)
Replicate Similarity (ICC)
Replicate Similarity (ICC)
0.90
0.92
0.94
0.96
0.98
0
10
20
30
12
13
14
Raw data
ENmix_noRCP
ENmix_RCP
Hybrid
Minfi
WateRmelon

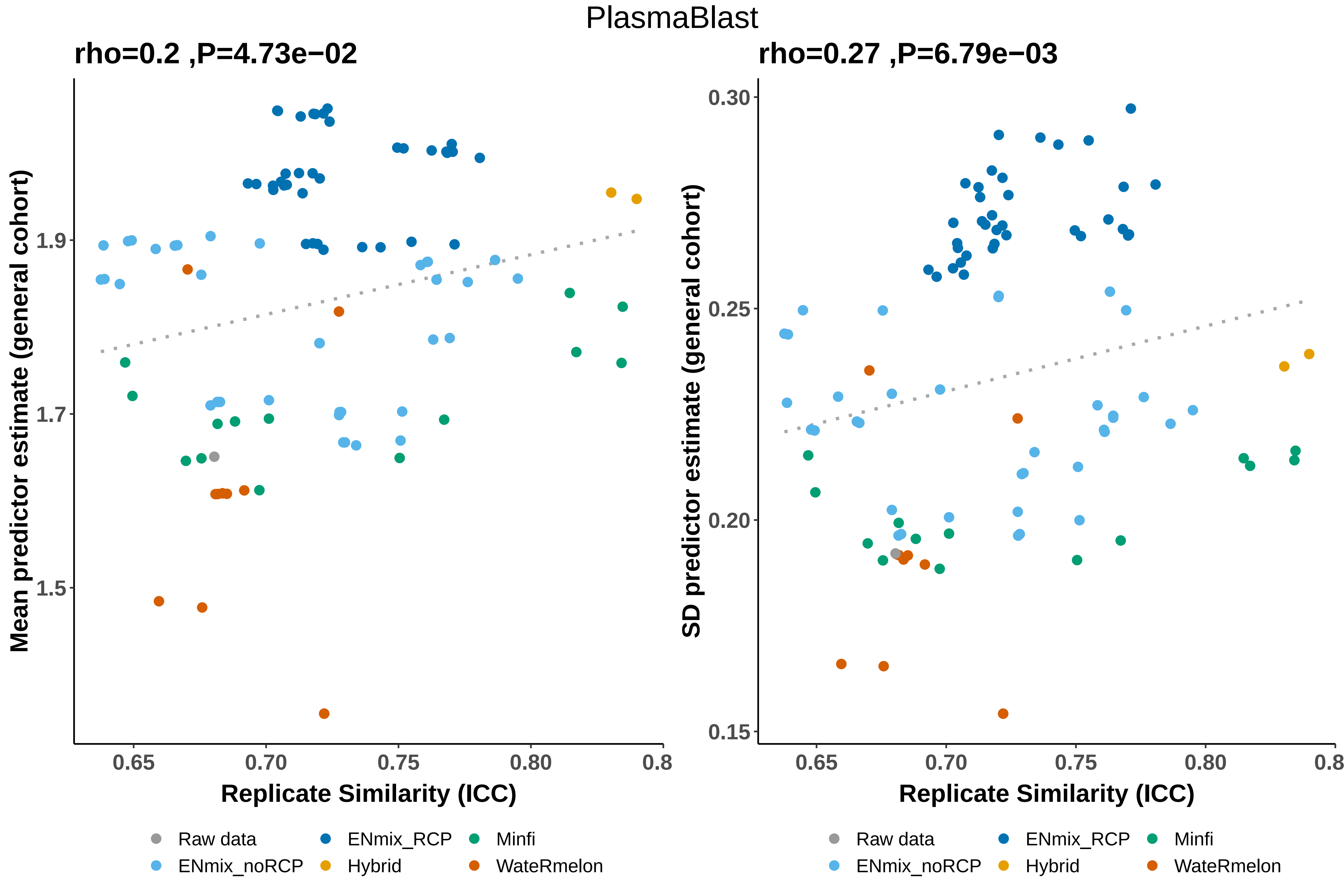
PlasmaBlast
rho=0.2 ,P=4.73e−02
Mean predictor estimate (general cohort)
1.9
1.7
1.5
0.65
0.70
0.75
0.80
0.8
Replicate Similarity (ICC)
rho=0.27 ,P=6.79e−03
SD predictor estimate (general cohort)
0.30
0.25
0.20
0.15
0.65
0.70
0.75
0.80
0.8
Replicate Similarity (ICC)
Raw data
ENmix_RCP
Minfi
ENmix_noRCP
Hybrid
WateRmelon

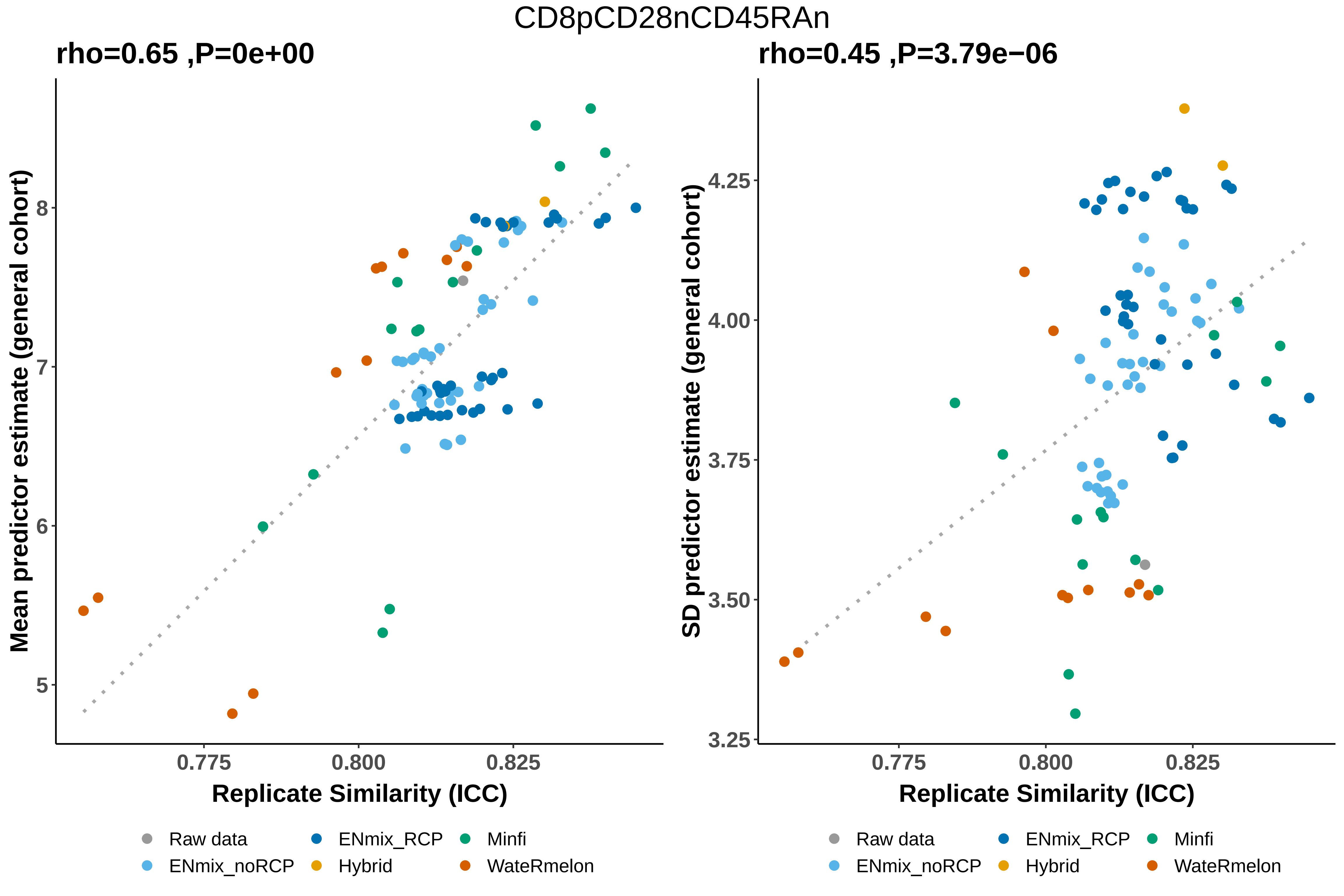

CD8pCD28nCD45RAn
rho=0.65 ,P=0e+00
rho=0.45 ,P=3.79e−06
Mean predictor estimate (general cohort)
SD predictor estimate (general cohort)
Replicate Similarity (ICC)
Replicate Similarity (ICC)
5
6
7
8
3.25
3.50
3.75
4.00
4.25
0.775
0.800
0.825
0.775
0.800
0.825
Raw data
ENmix_noRCP
ENmix_RCP
Hybrid
Minfi
WateRmelon
Raw data
ENmix_noRCP
ENmix_RCP
Hybrid
Minfi
WateRmelon

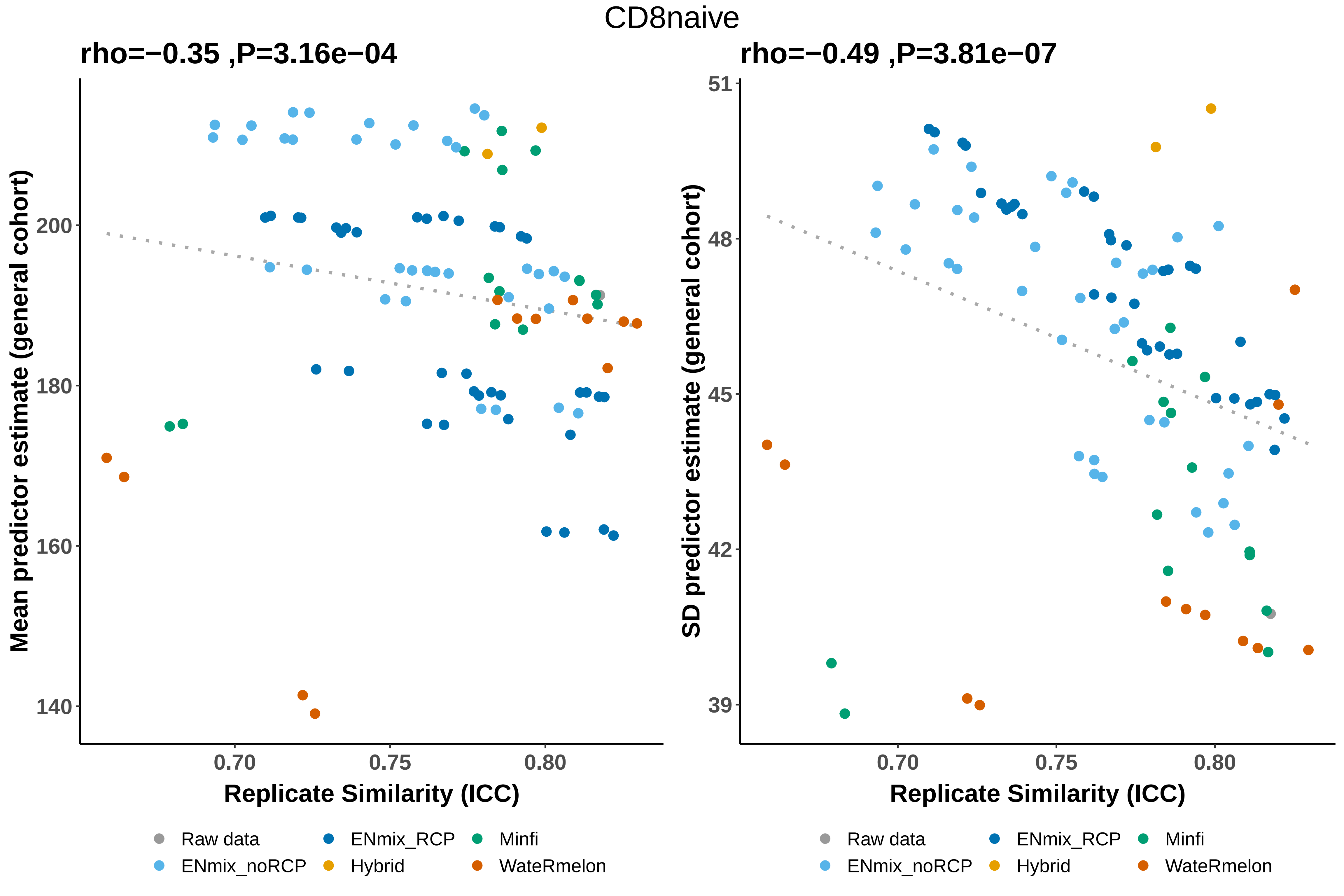
CD8naive
rho=−0.35 ,P=3.16e−04
rho=−0.49 ,P=3.81e−07
Mean predictor estimate (general cohort)
SD predictor estimate (general cohort)
Replicate Similarity (ICC)
Replicate Similarity (ICC)
140
160
180
200
39
42
45
48
51
0.70
0.75
0.80
0.70
0.75
0.80
Raw data
ENmix_noRCP
ENmix_RCP
Hybrid
Minfi
WateRmelon
Raw data
ENmix_noRCP
ENmix_RCP
Hybrid
Minfi
WateRmelon

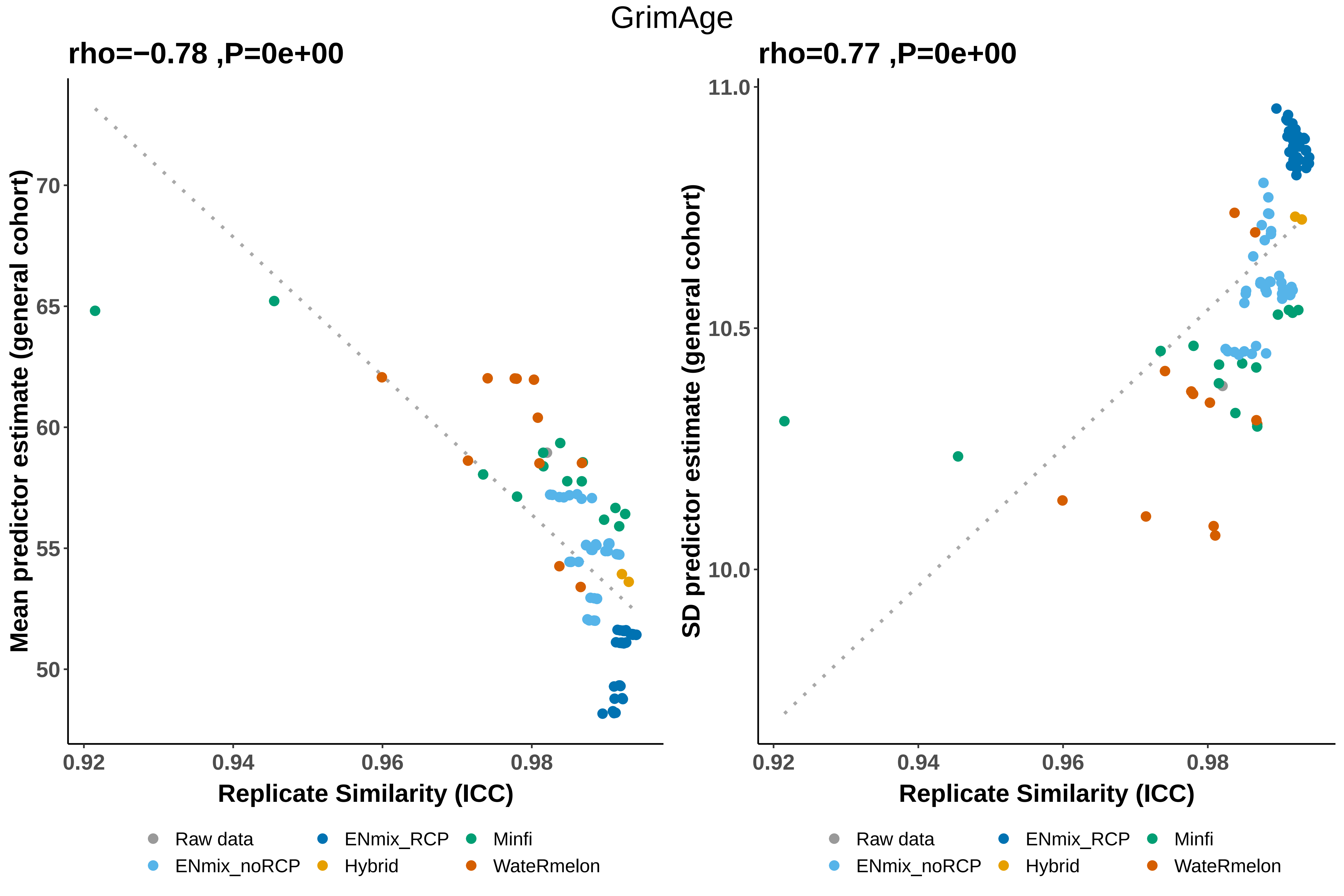
GrimAge
rho=−0.78 ,P=0e+00
Mean predictor estimate (general cohort)
70
65
60
55
50
0.92
0.94
0.96
0.98
Replicate Similarity (ICC)
rho=0.77 ,P=0e+00
SD predictor estimate (general cohort)
11.0
10.5
10.0
0.92
0.94
0.96
0.98
Replicate Similarity (ICC)
Raw data
ENmix_RCP
Minfi
ENmix_noRCP
Hybrid
WateRmelon

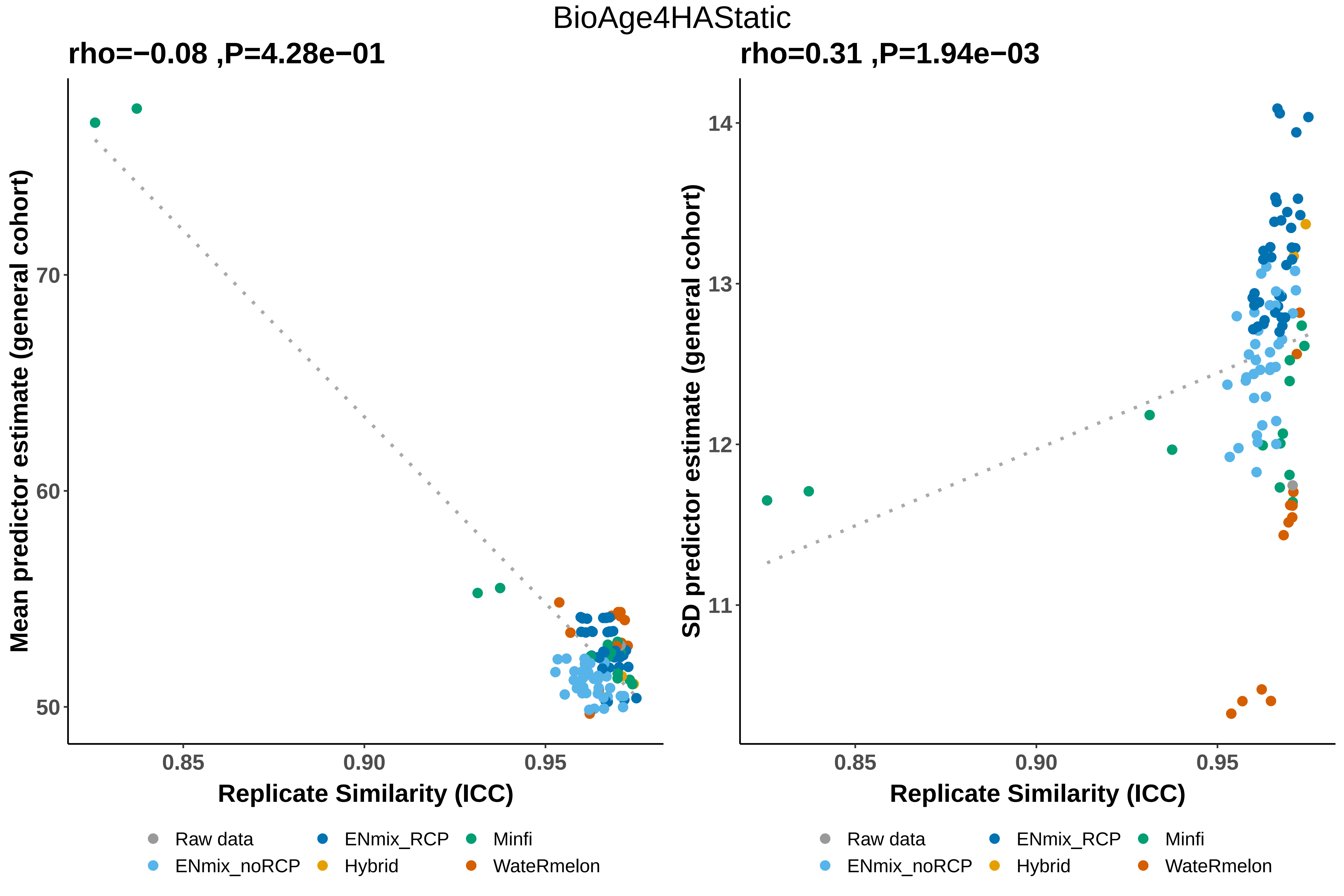

BioAge4HAStatic
rho=−0.08 ,P=4.28e−01
rho=0.31 ,P=1.94e−03
Mean predictor estimate (general cohort)
SD predictor estimate (general cohort)
Replicate Similarity (ICC)
70
60
50
14
13
12
11
0.85
0.90
0.95
Raw data
ENmix_noRCP
ENmix_RCP
Hybrid
Minfi
WateRmelon
