## Supplemental Note 2 for "A systematic evaluation of 41 DNA methylation predictors across 101 data preprocessing and normalization strategies highlights considerable variation in algorithm performance"

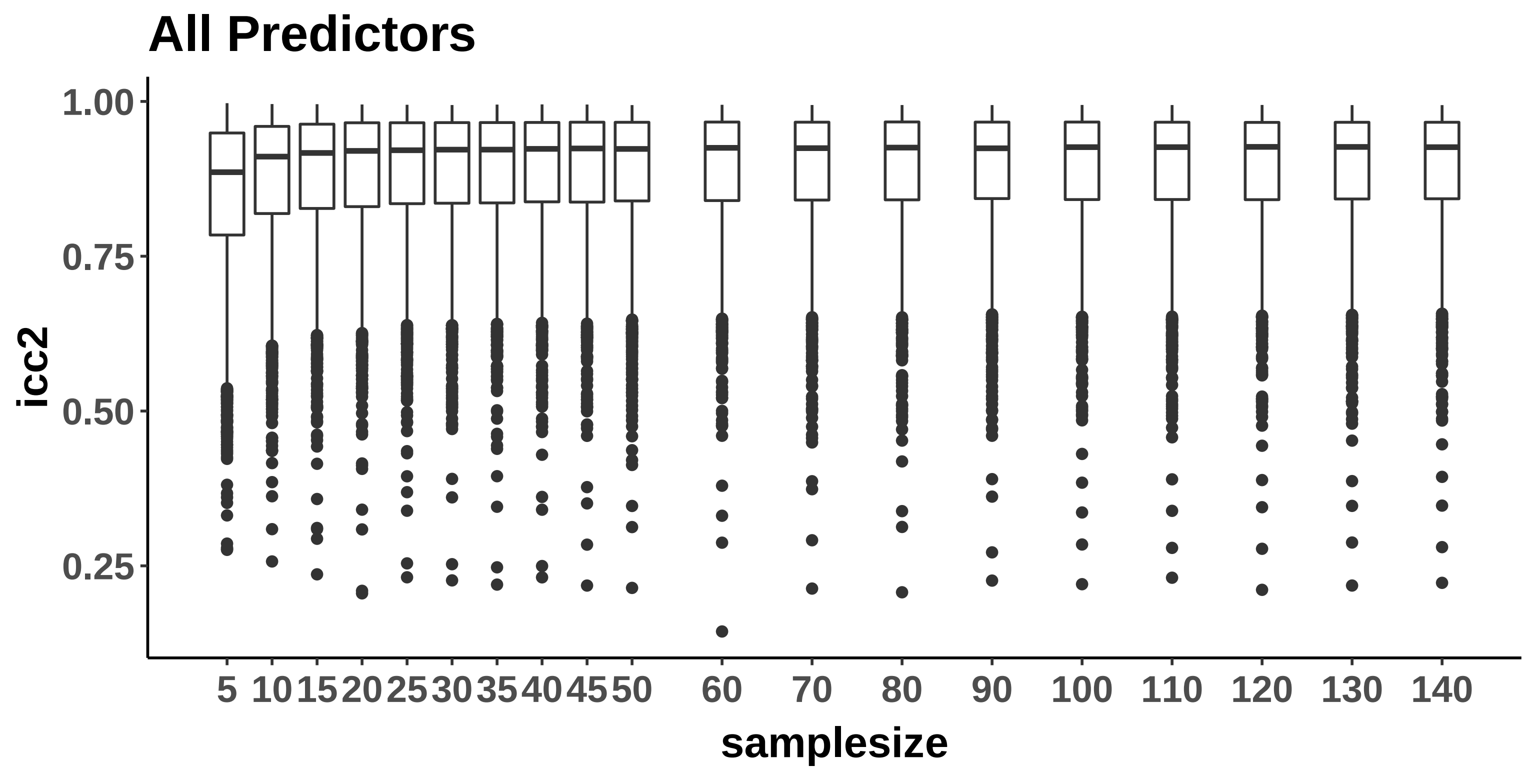
All Predictors
1.00
0.75
0.50
0.25
icc2
5
10
15
20
25
30
35
40
45
50
60
70
80
90
100
110
120
130
140
samplesize

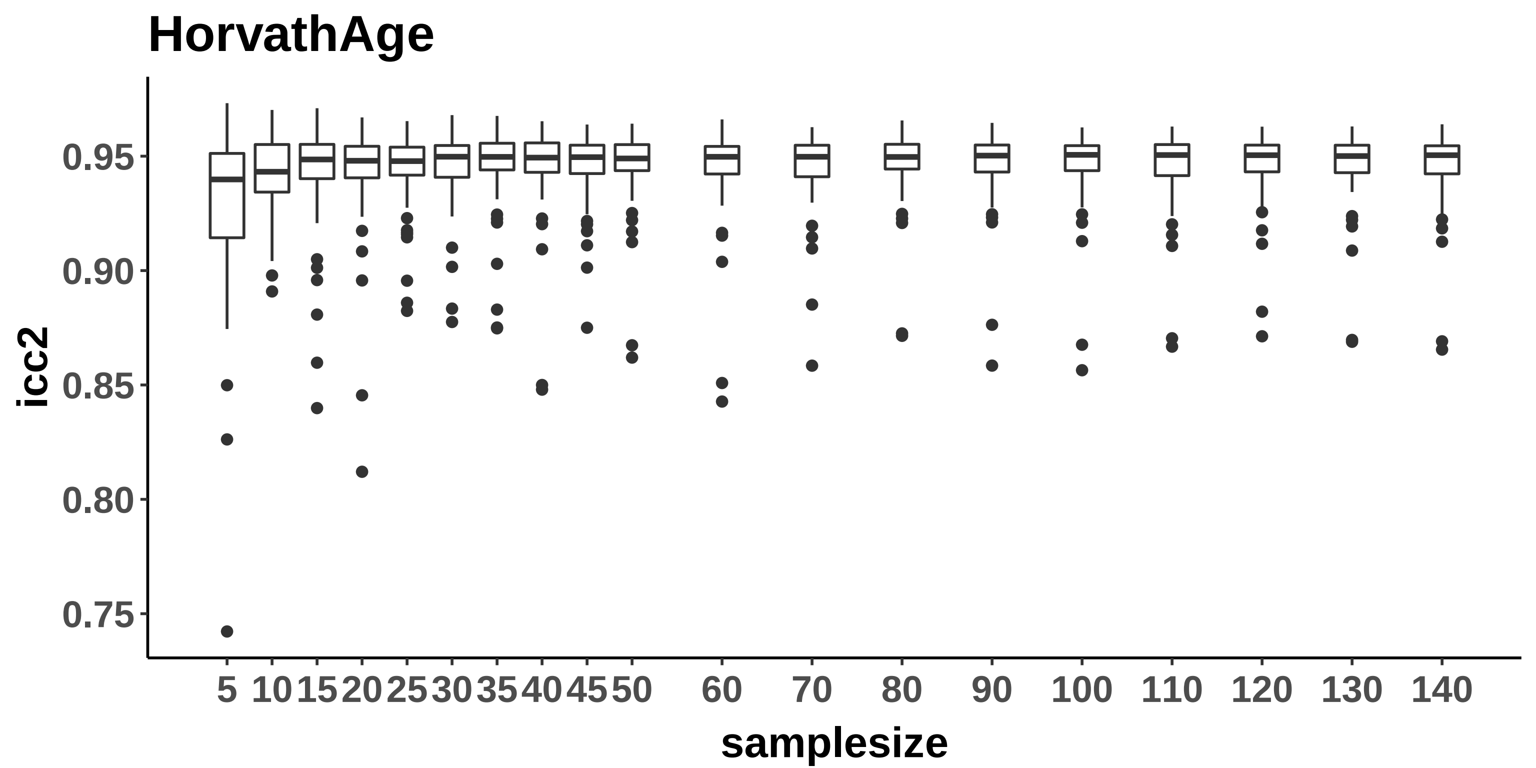
HorvathAge
icc2
0.95
0.90
0.85
0.80
0.75
5 10 15 20 25 30 35 40 45 50 60 70 80 90 100 110 120 130 140
samplesize

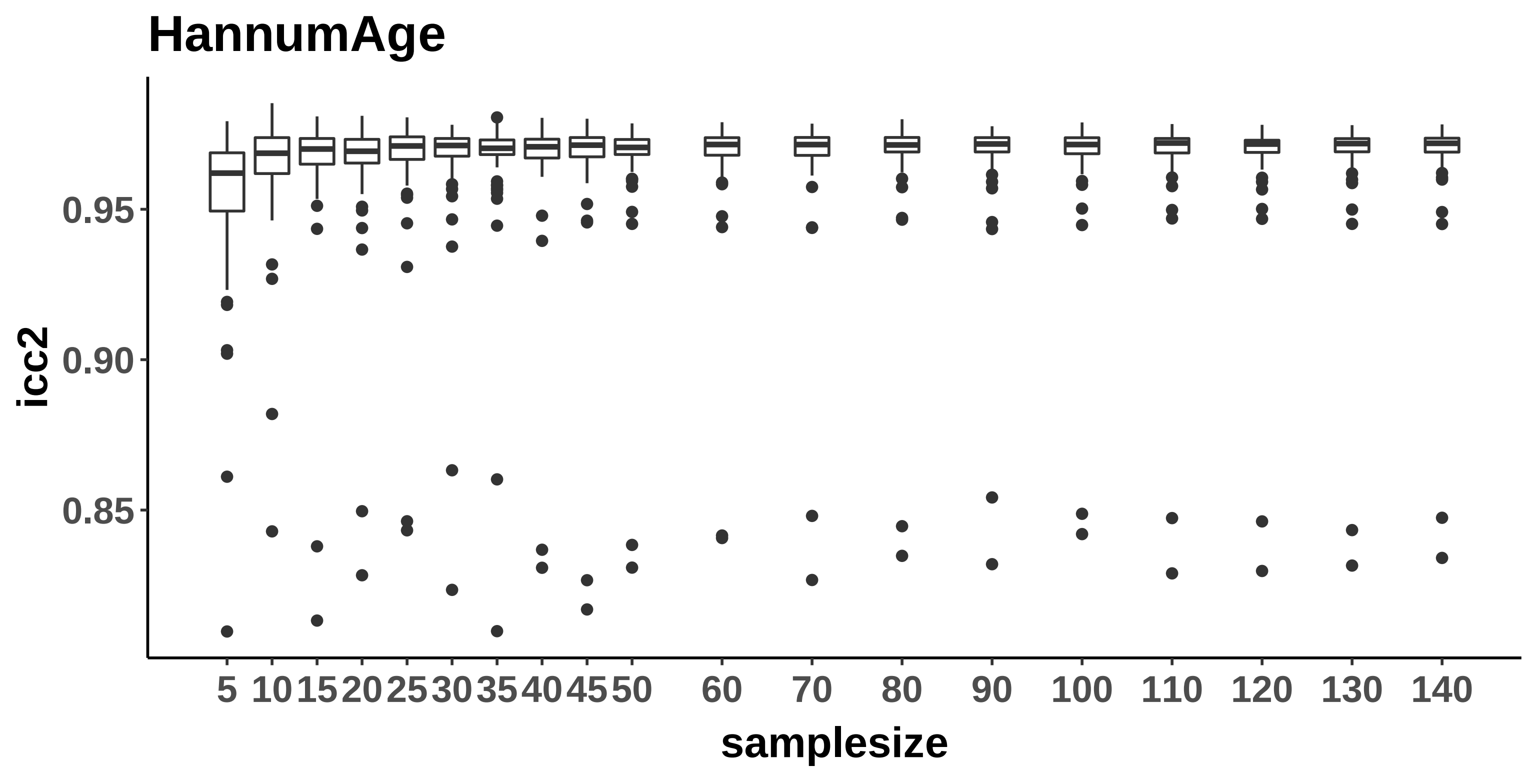

HannumAge
icc2
0.95
0.90
0.85
5
10
15
20
25
30
35
40
45
50
60
70
80
90
100
110
120
130
140
samplesize

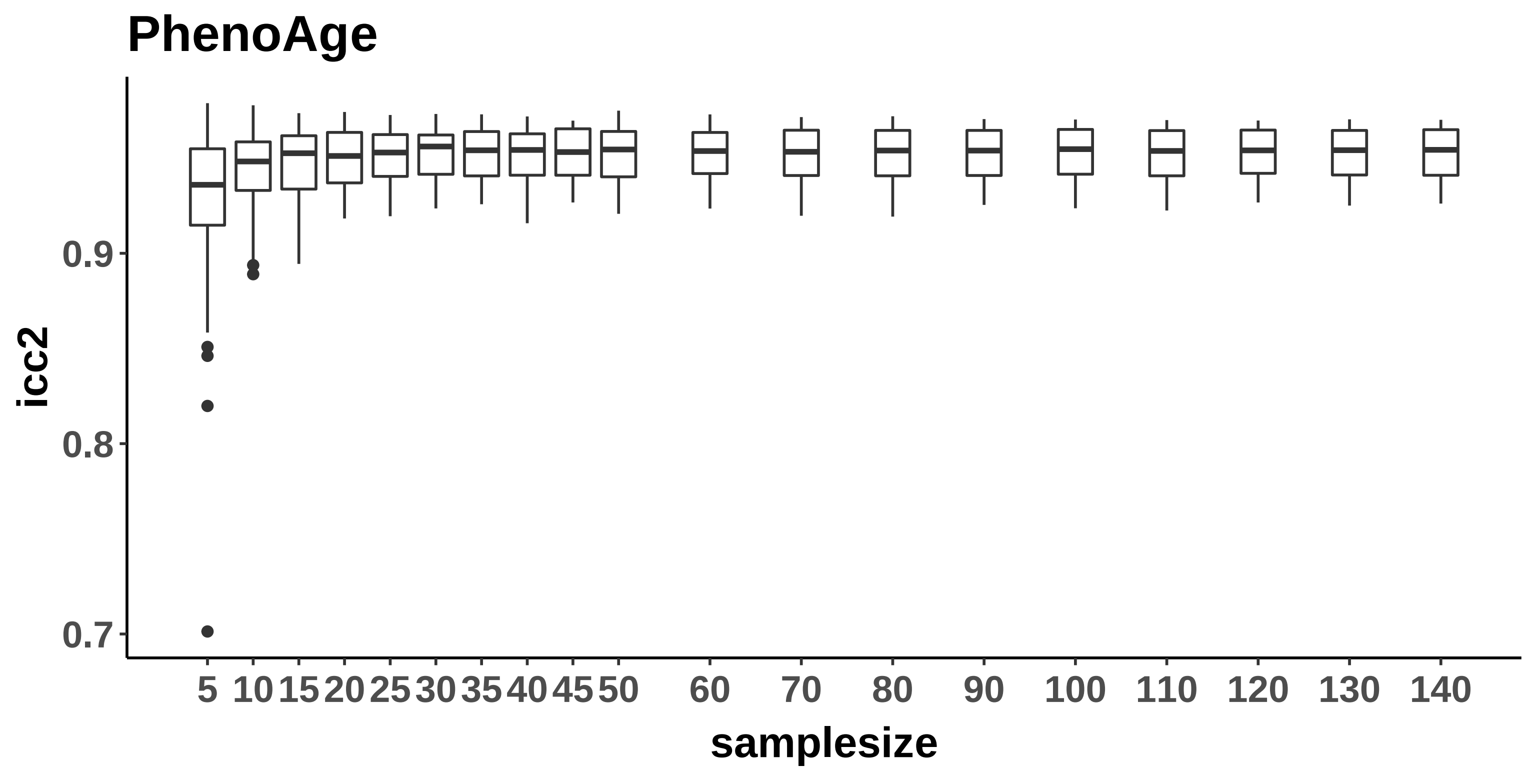

PhenoAge
icc2
0.9
0.8
0.7
5
10
15
20
25
30
35
40
45
50
60
70
80
90
100
110
120
130
140
samplesize

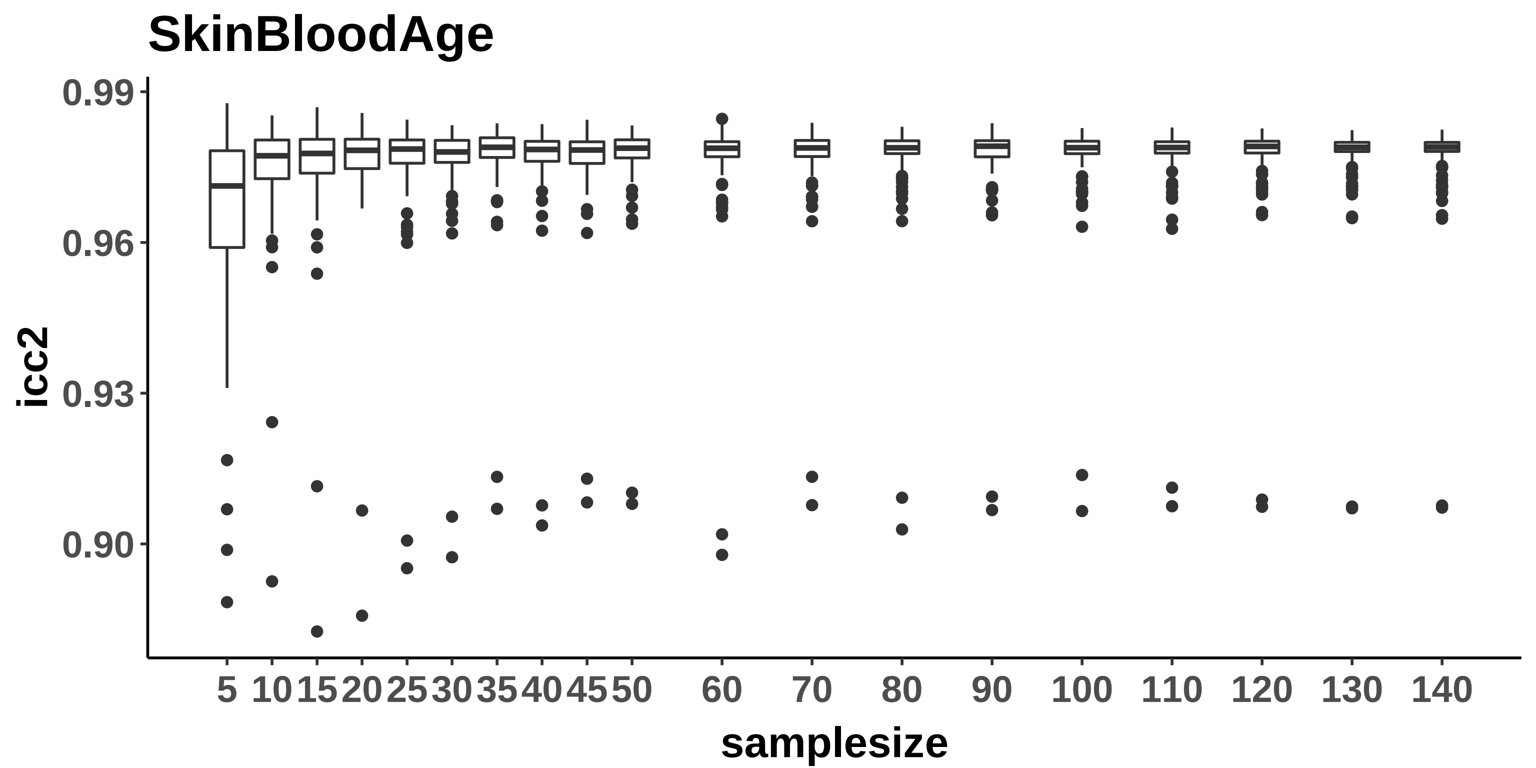
SkinBloodAge
0.99
0.96
0.93
0.90
icc2
5
10
15
20
25
30
35
40
45
50
60
70
80
90
100
110
120
130
140
samplesize

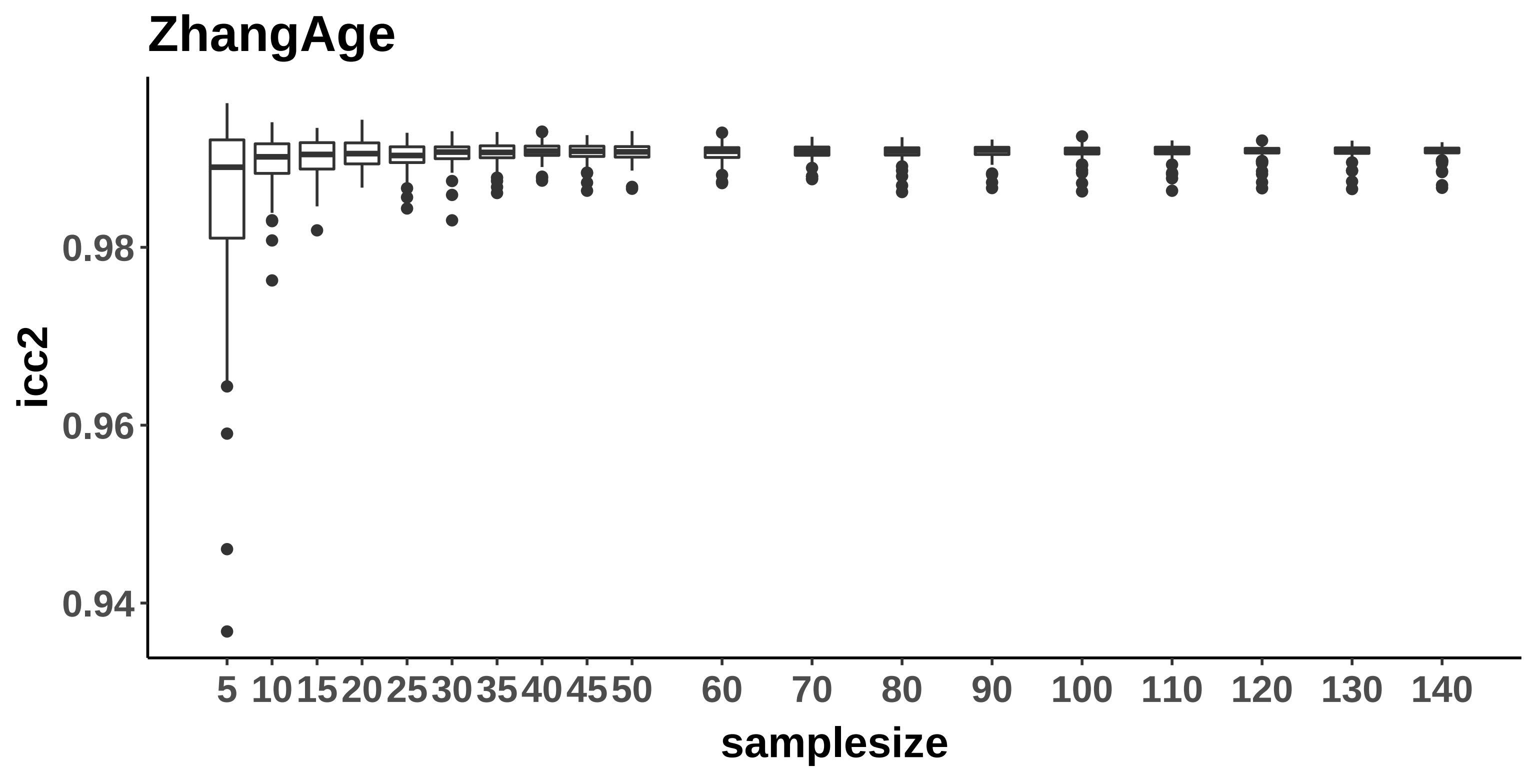
ZhangAge
icc2
0.98
0.96
0.94
samplesize
5
10
15
20
25
30
35
40
45
50
60
70
80
90
100
110
120
130
140

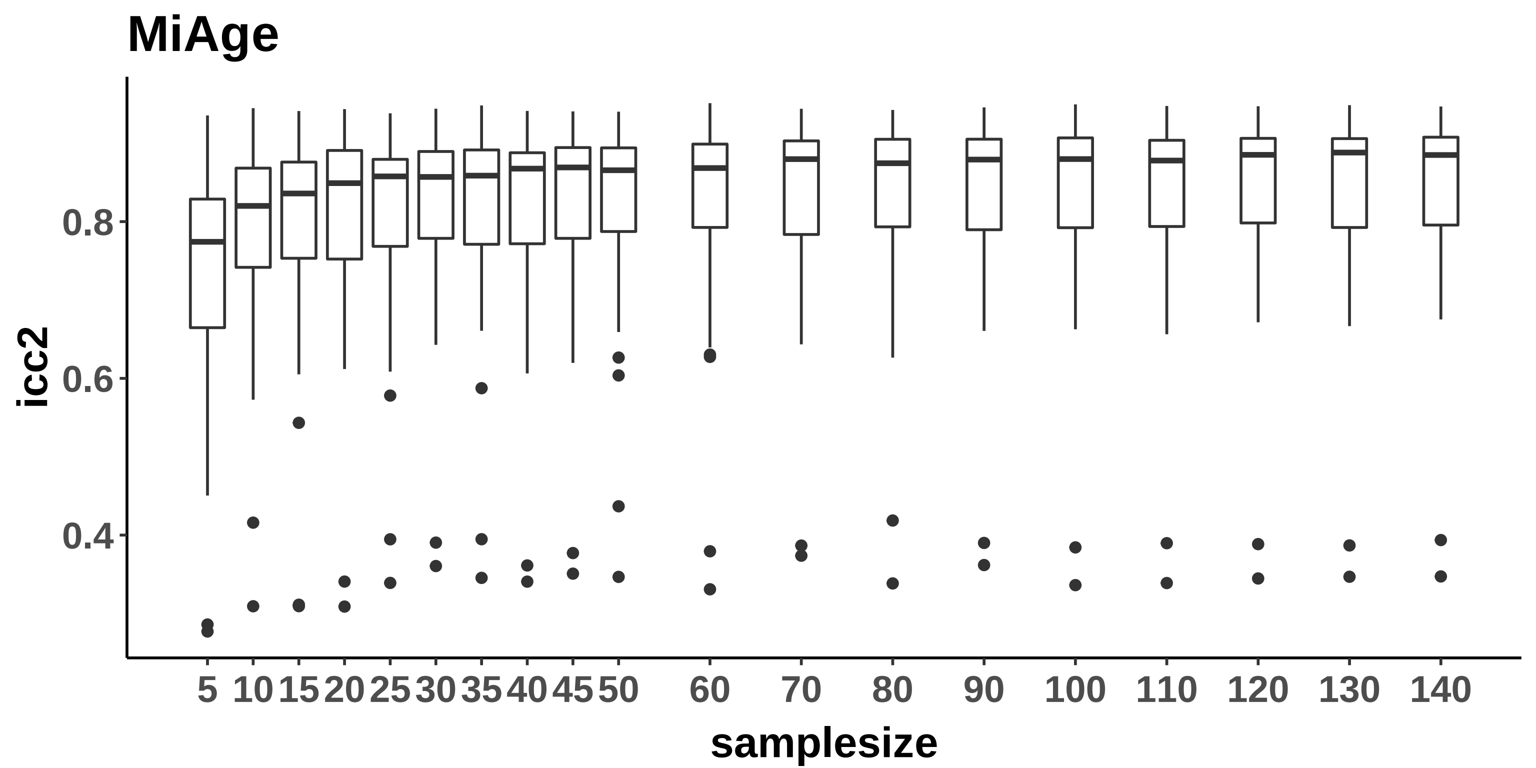
MiAge
icc2
0.8
0.6
0.4
samplesize
5
10
15
20
25
30
35
40
45
50
60
70
80
90
100
110
120
130
140

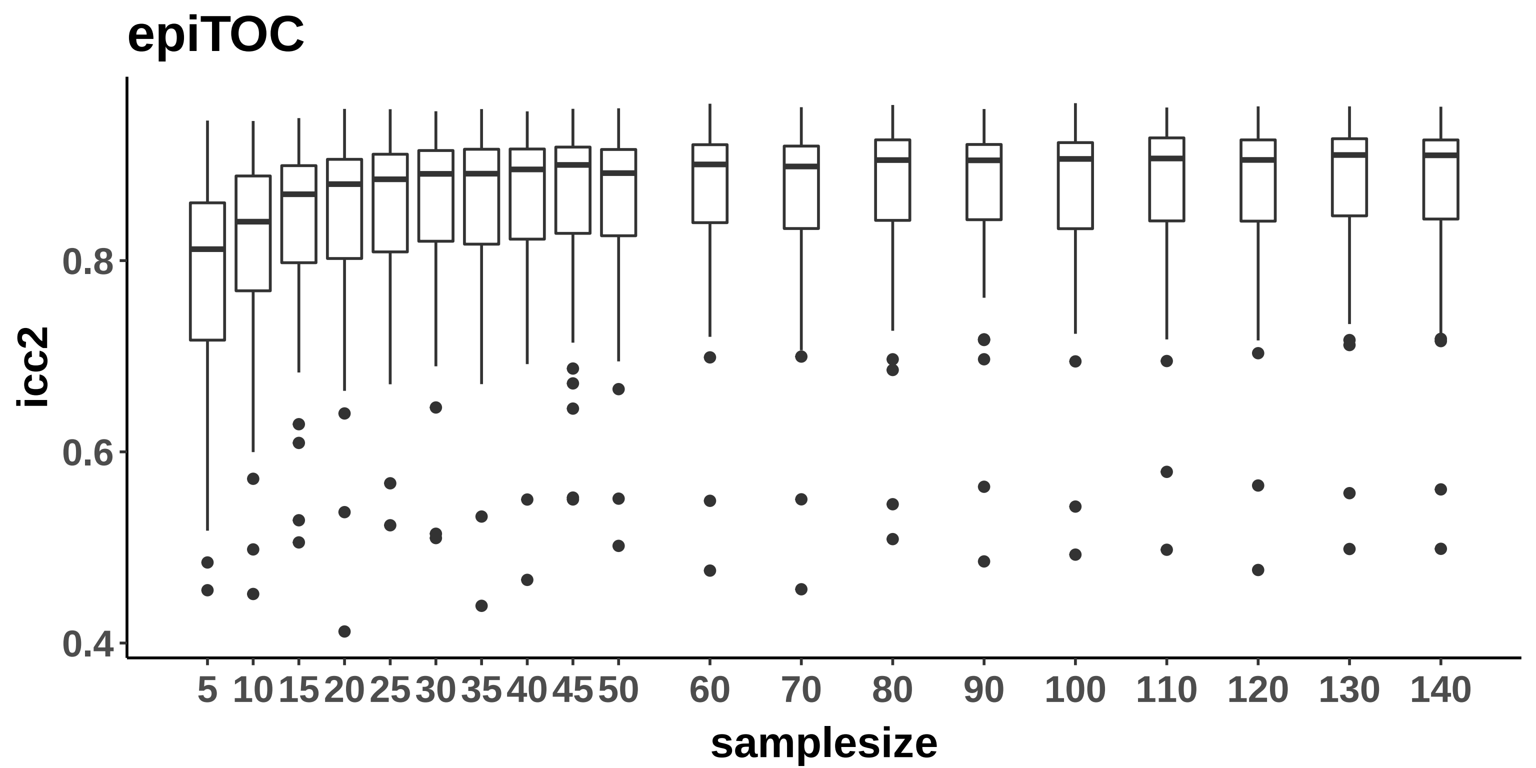

epiTOC
icc2
0.8
0.6
0.4
5
10
15
20
25
30
35
40
45
50
60
70
80
90
100
110
120
130
140
samplesize

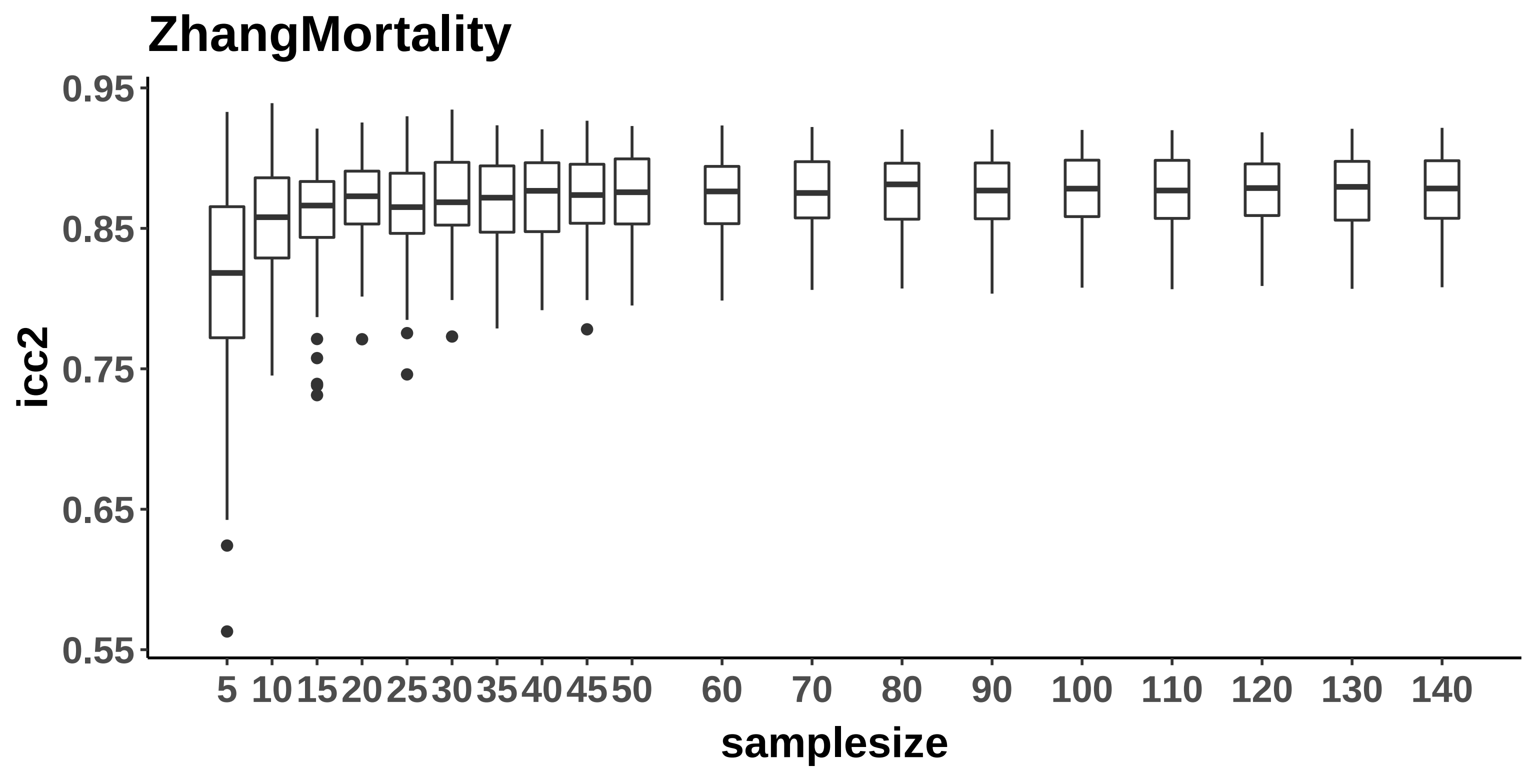
ZhangMortality
0.95
0.85
0.75
0.65
0.55
icc2
5
10
15
20
25
30
35
40
45
50
60
70
80
90
100
110
120
130
140
samplesize

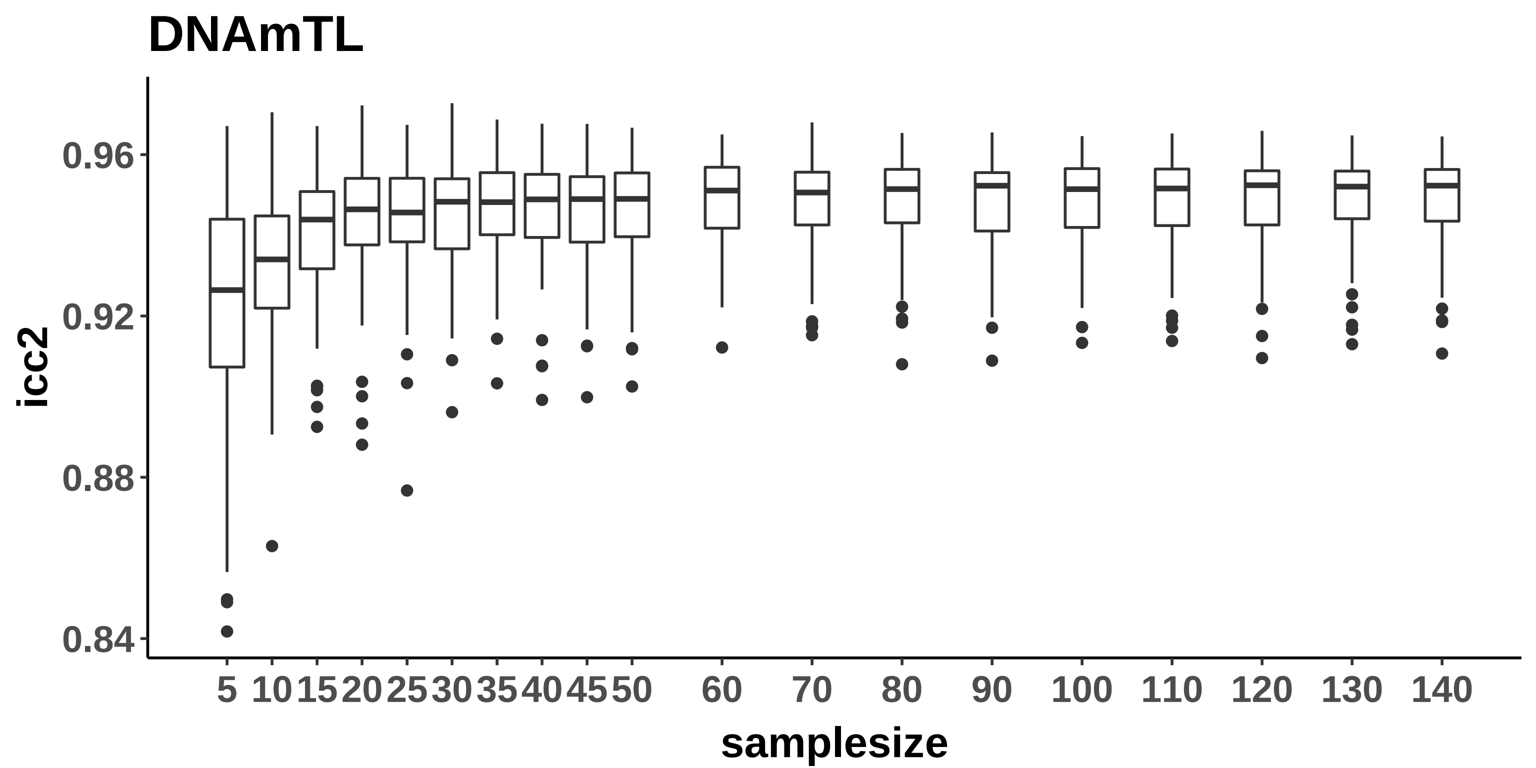
DNAmTL
icc2
0.96
0.92
0.88
0.84
5
10
15
20
25
30
35
40
45
50
60
70
80
90
100
110
120
130
140
samplesize

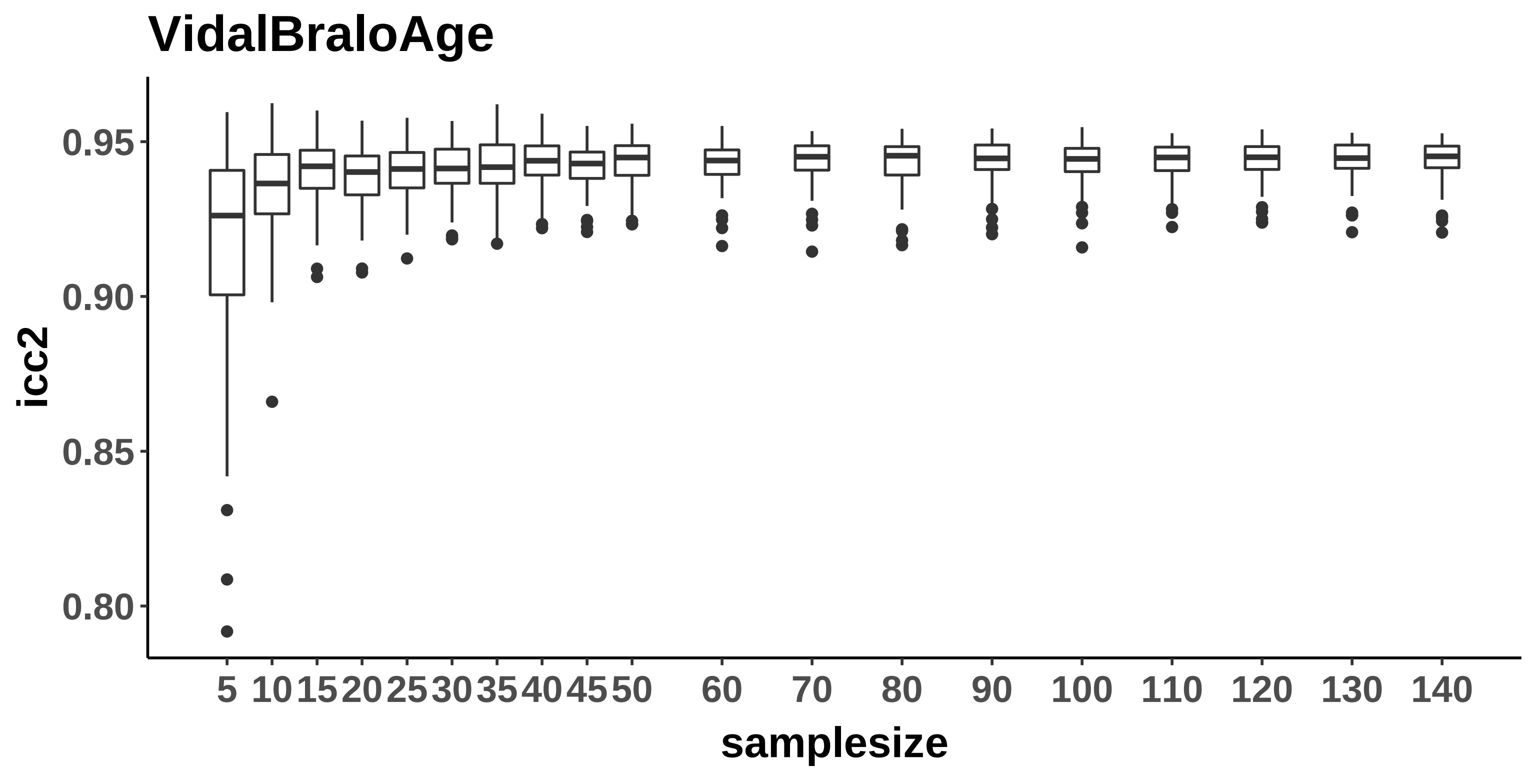

VidalBraloAge
icc2
0.95
0.90
0.85
0.80
5
10
15
20
25
30
35
40
45
50
60
70
80
90
100
110
120
130
140
samplesize

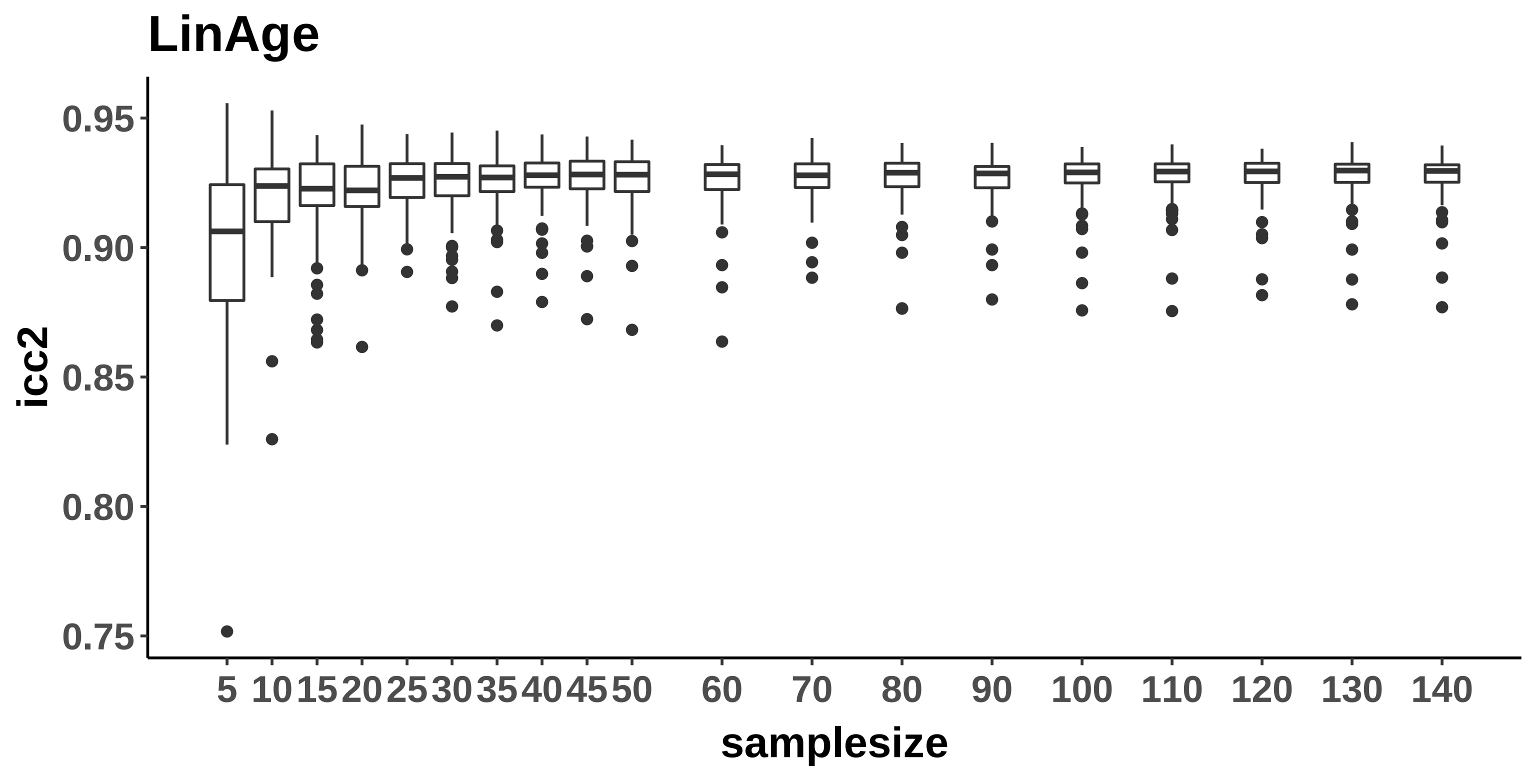
LinAge
icc2
0.95
0.90
0.85
0.80
0.75
5
10
15
20
25
30
35
40
45
50
60
70
80
90
100
110
120
130
140
samplesize

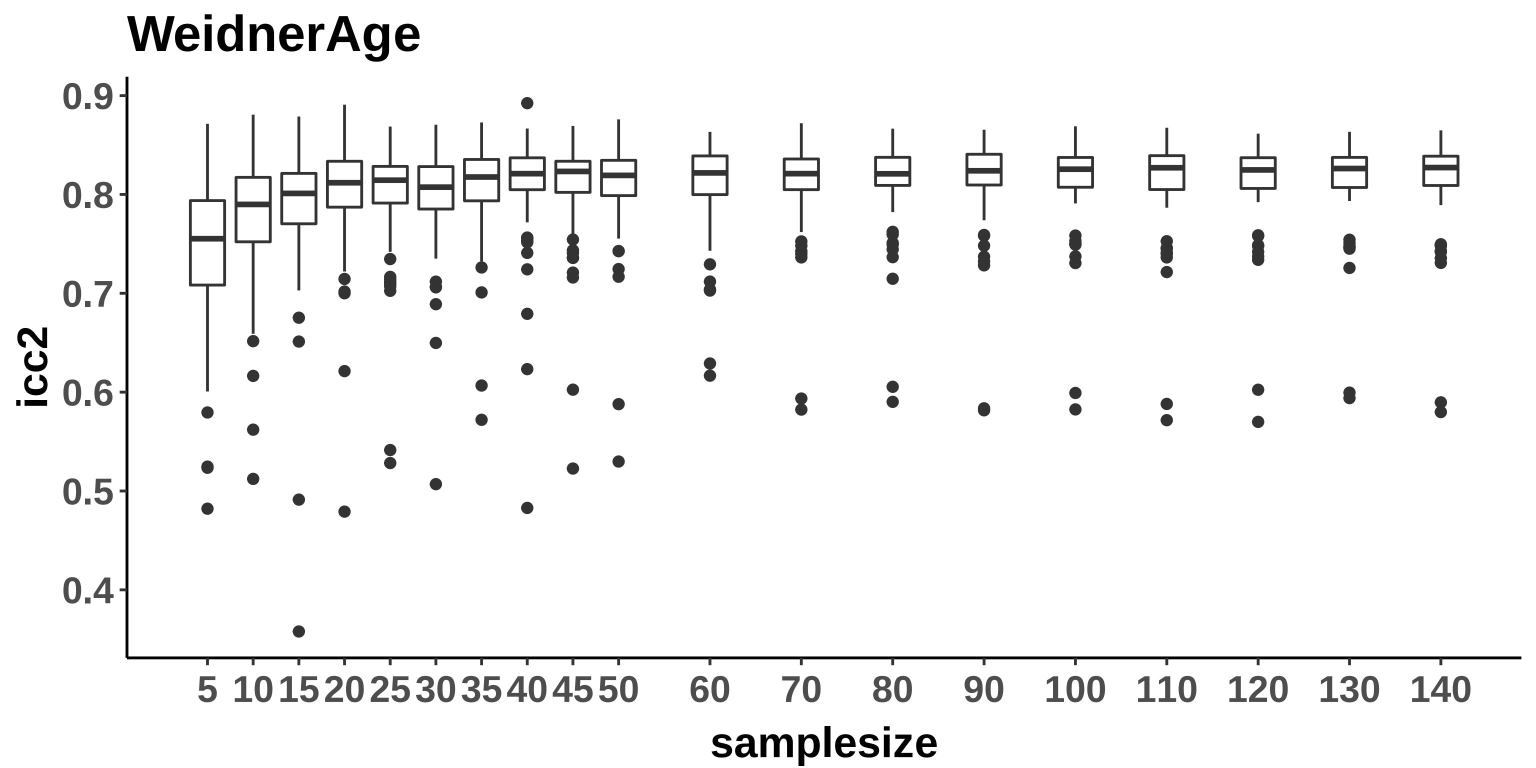
WeidnerAge
icc2
0.9
0.8
0.7
0.6
0.5
0.4
5
10
15
20
25
30
35
40
45
50
60
70
80
90
100
110
120
130
140
samplesize

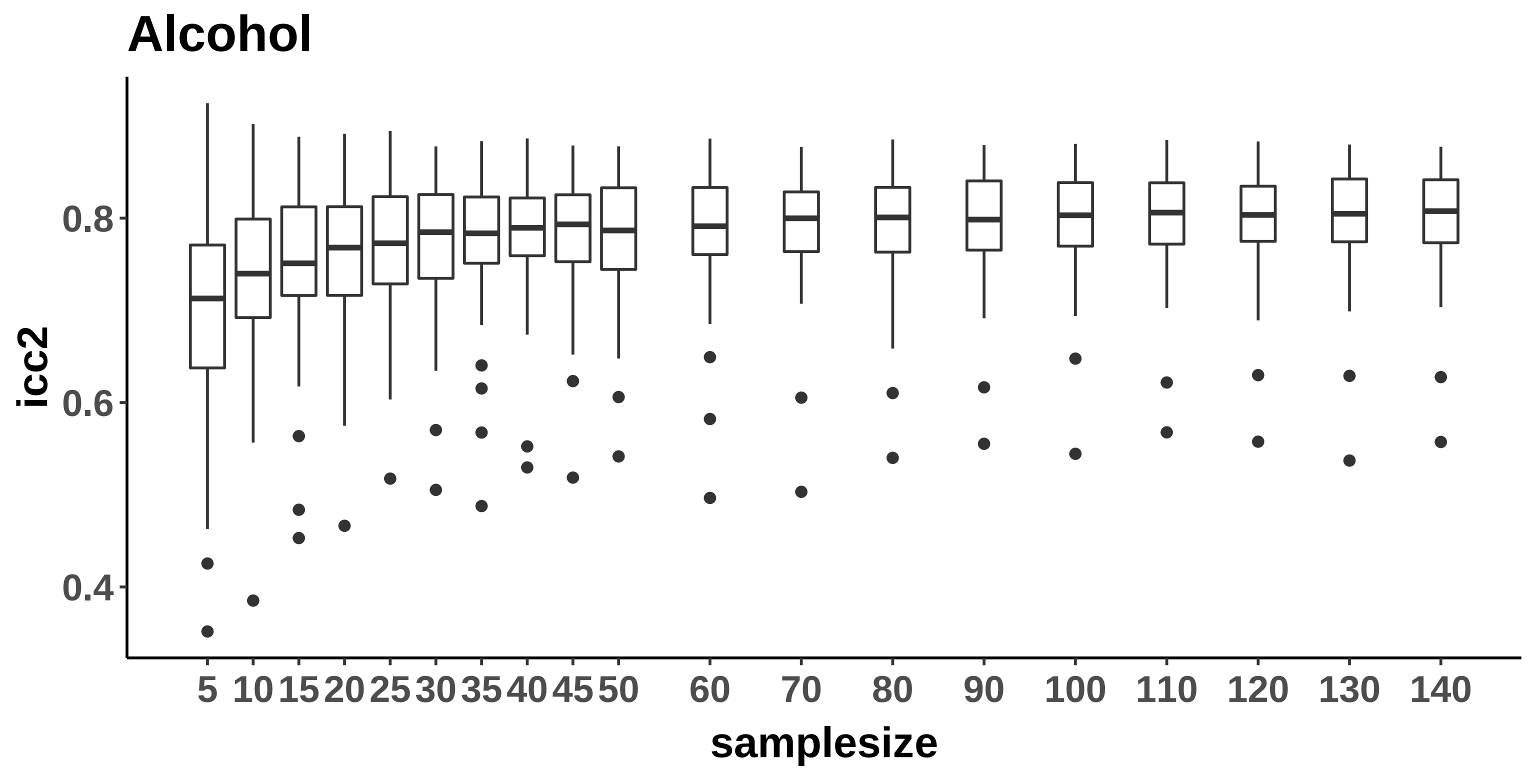

Alcohol
icc2
0.8
0.6
0.4
5
10
15
20
25
30
35
40
45
50
60
70
80
90
100
110
120
130
140
samplesize

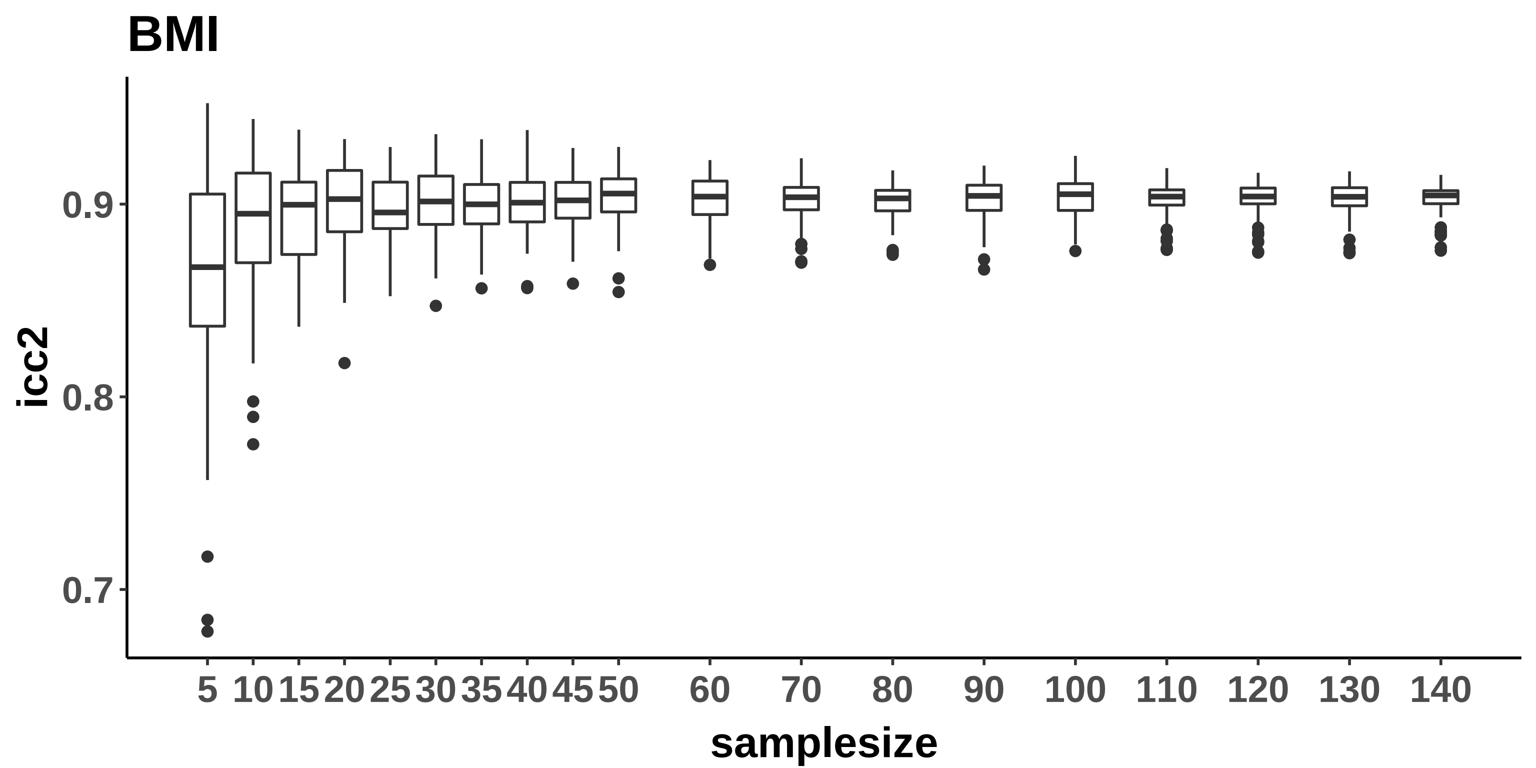
BMI
icc2
0.9
0.8
0.7
5
10
15
20
25
30
35
40
45
50
60
70
80
90
100
110
120
130
140
samplesize

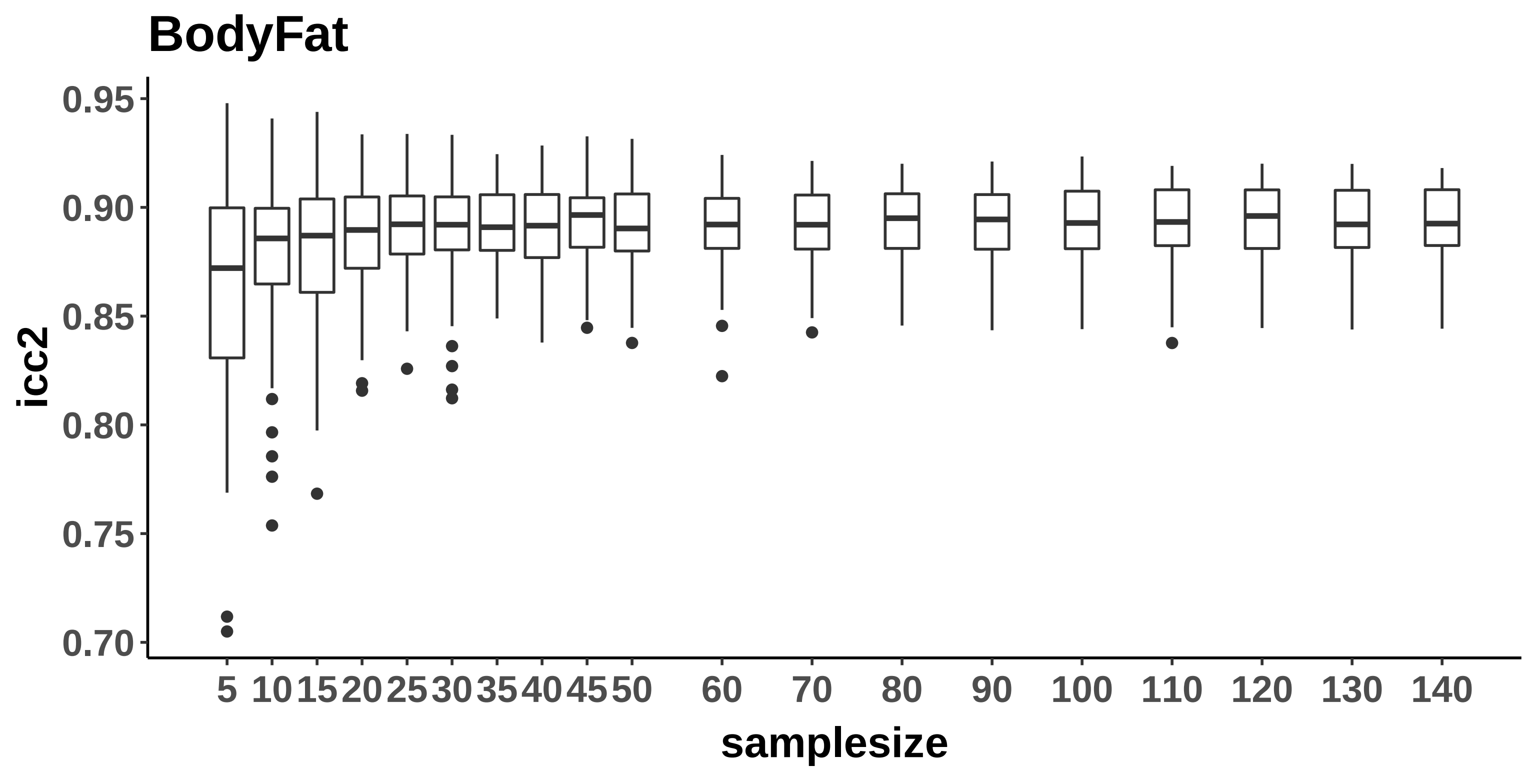
BodyFat
icc2
0.95
0.90
0.85
0.80
0.75
0.70
5
10
15
20
25
30
35
40
45
50
60
70
80
90
100
110
120
130
140
samplesize

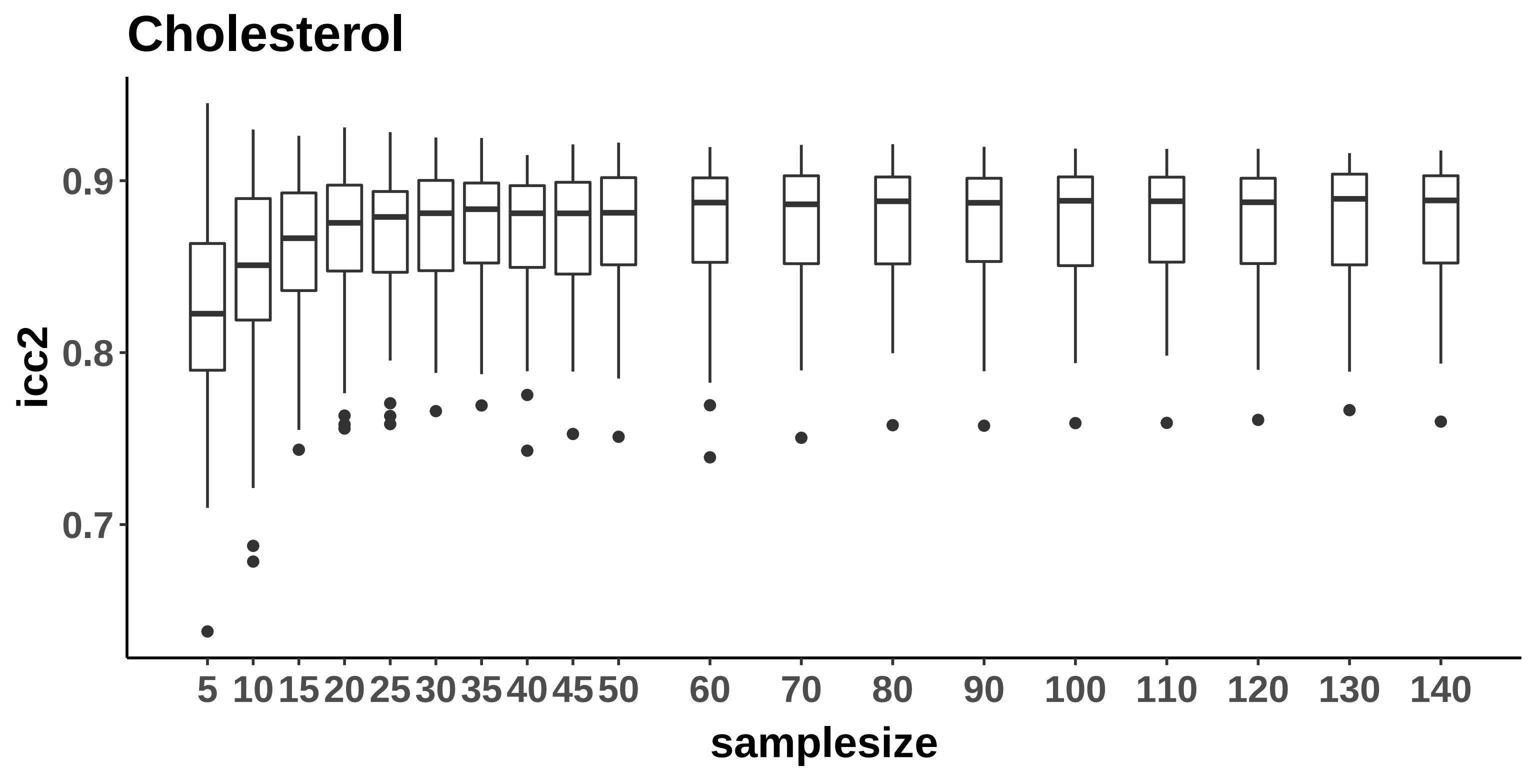
Cholesterol
icc2
0.9
0.8
0.7
samplesize
5 10 15 20 25 30 35 40 45 50 60 70 80 90 100 110 120 130 140

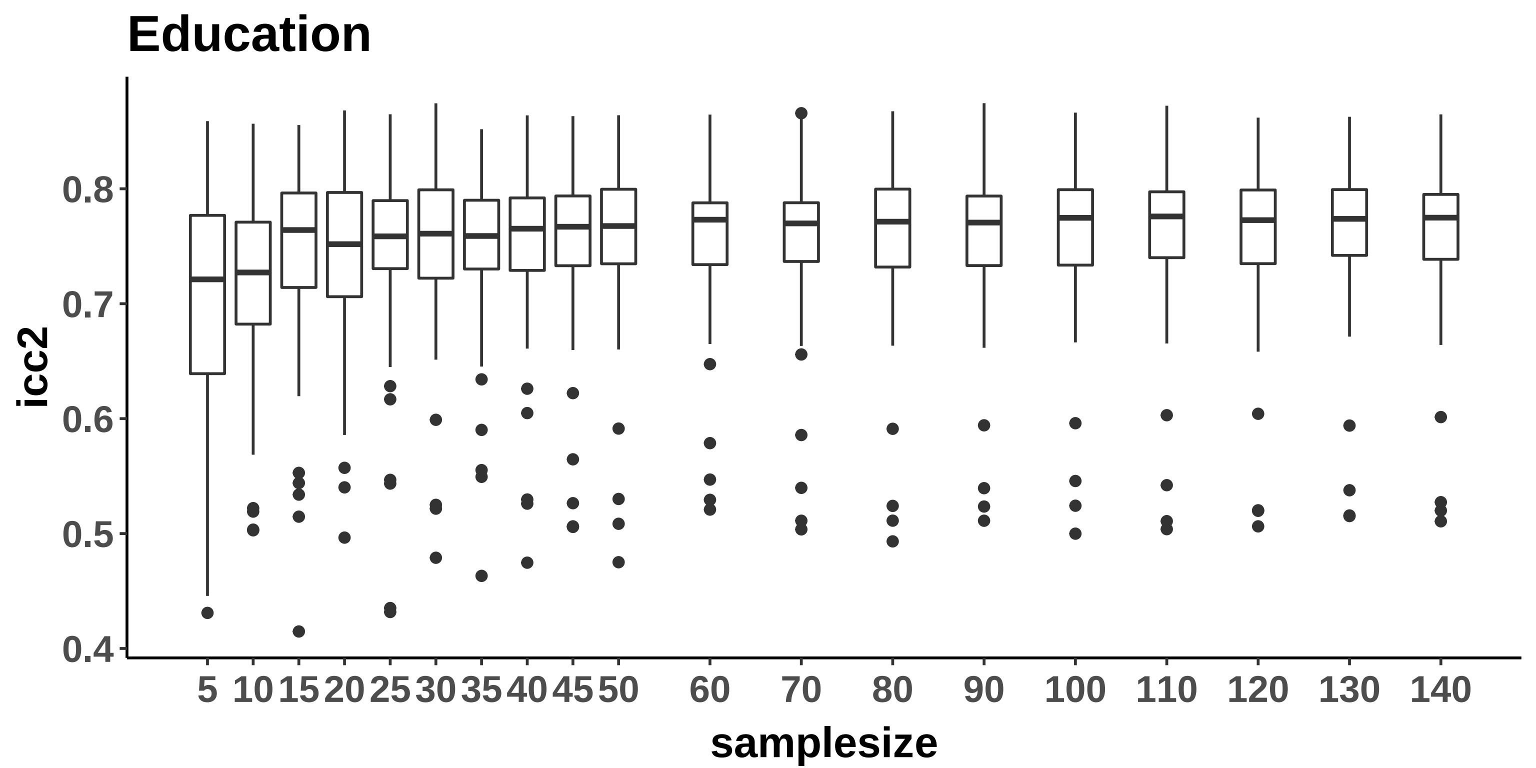
Education
icc2
0.8
0.7
0.6
0.5
0.4
5 10 15 20 25 30 35 40 45 50 60 70 80 90 100 110 120 130 140
samplesize

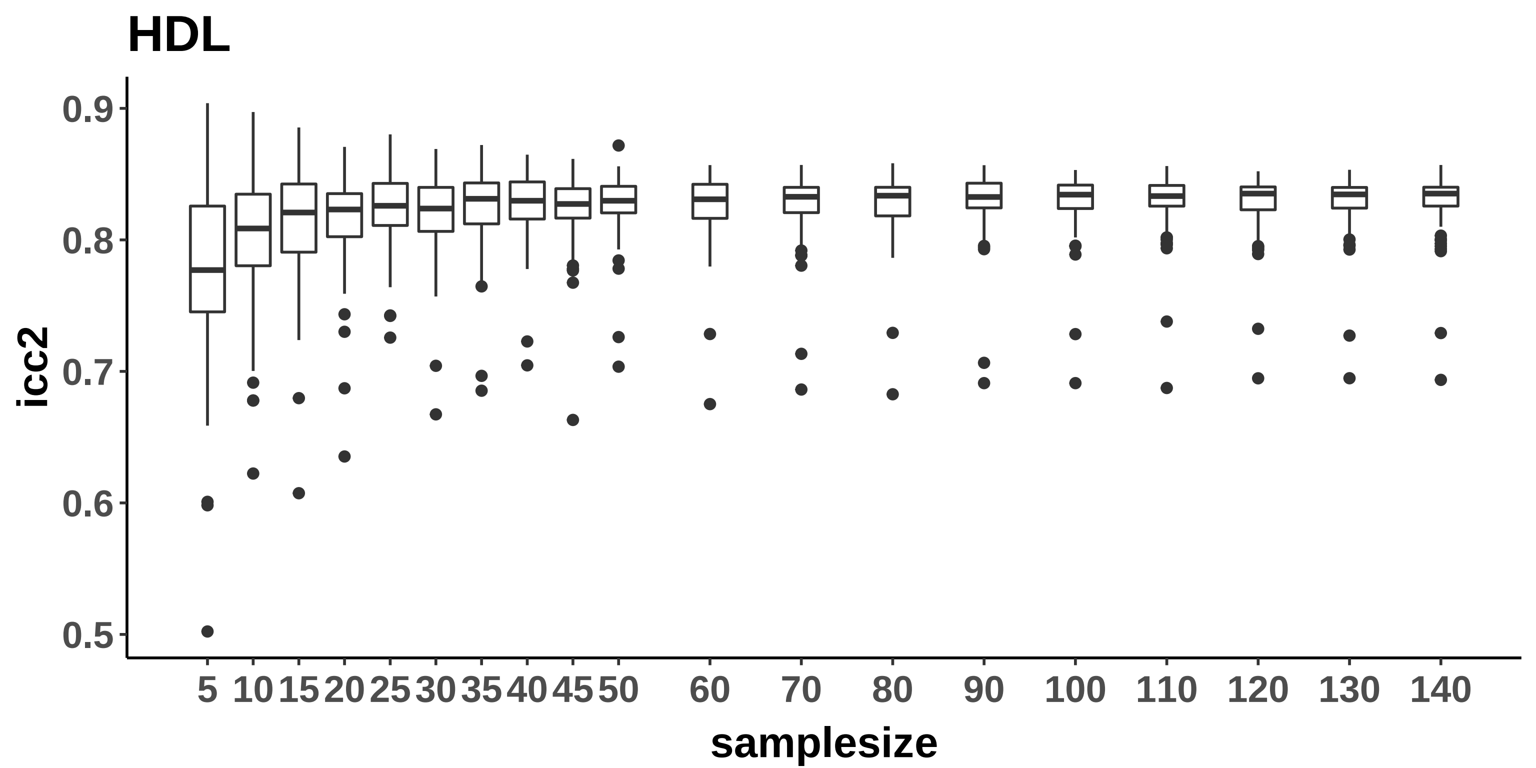
HDL
icc2
0.9
0.8
0.7
0.6
0.5
samplesize
5
10
15
20
25
30
35
40
45
50
60
70
80
90
100
110
120
130
140

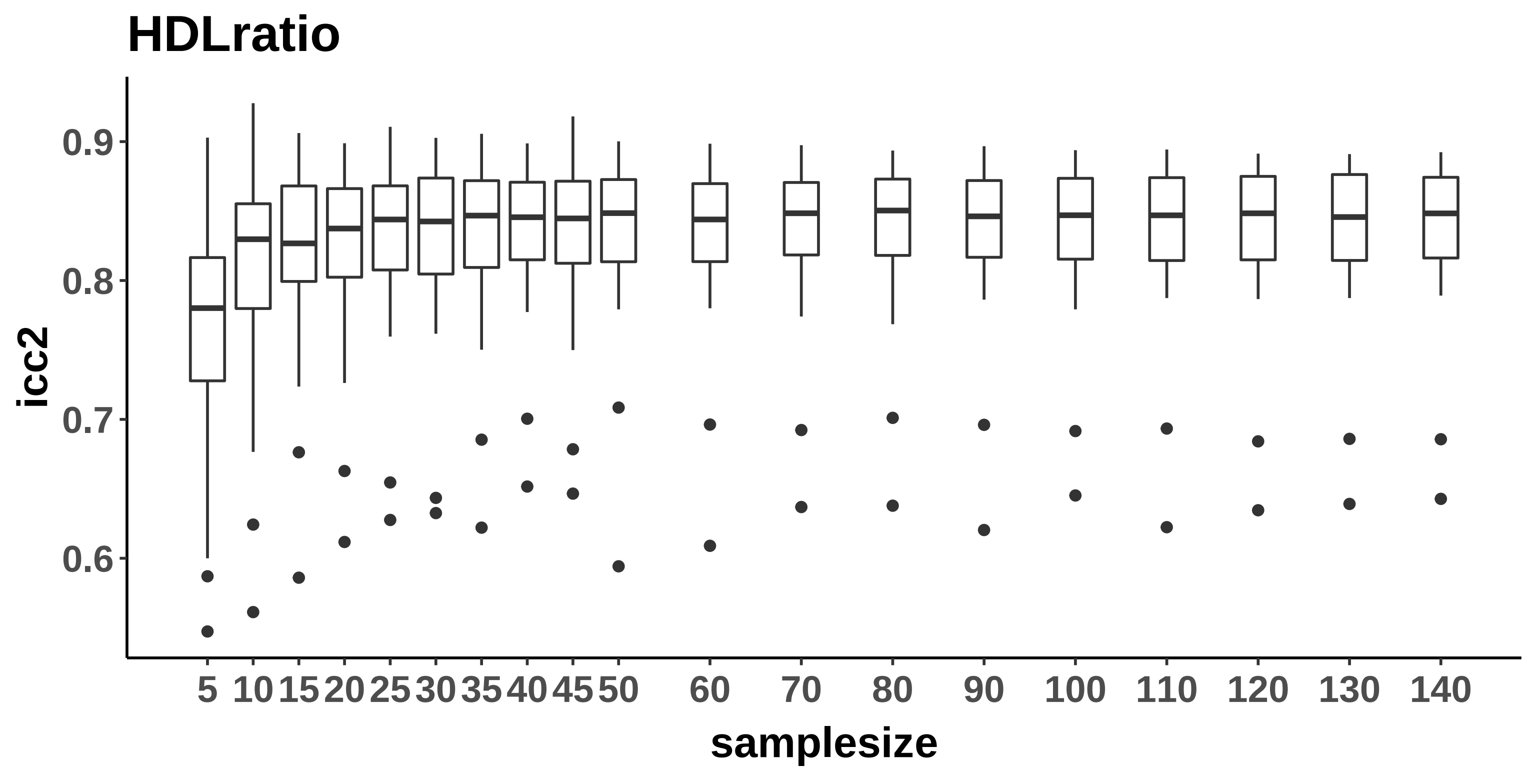
HDLratio
icc2
0.9
0.8
0.7
0.6
5
10
15
20
25
30
35
40
45
50
60
70
80
90
100
110
120
130
140
samplesize

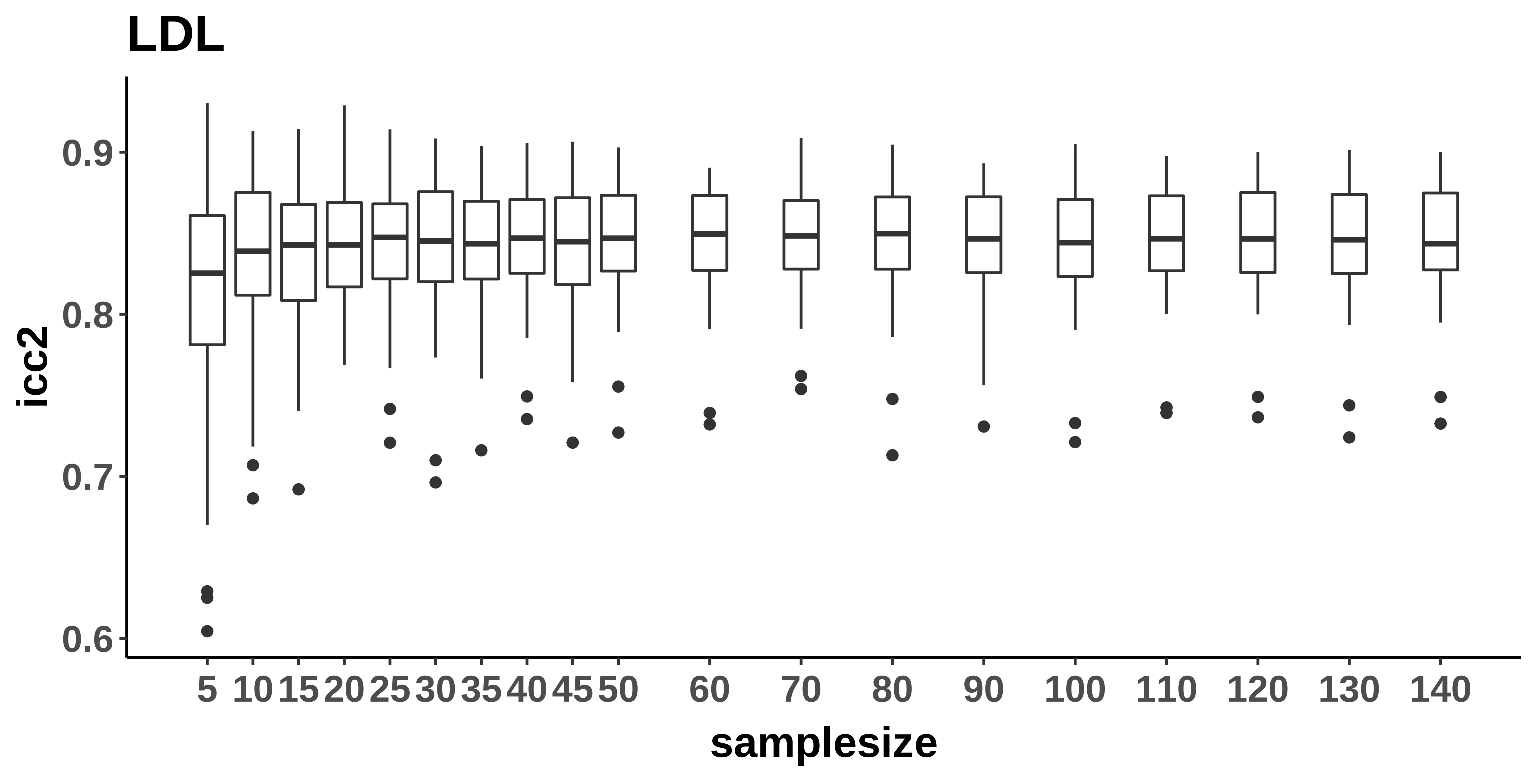
LDL
icc2
0.9
0.8
0.7
0.6
5
10
15
20
25
30
35
40
45
50
60
70
80
90
100
110
120
130
140
samplesize

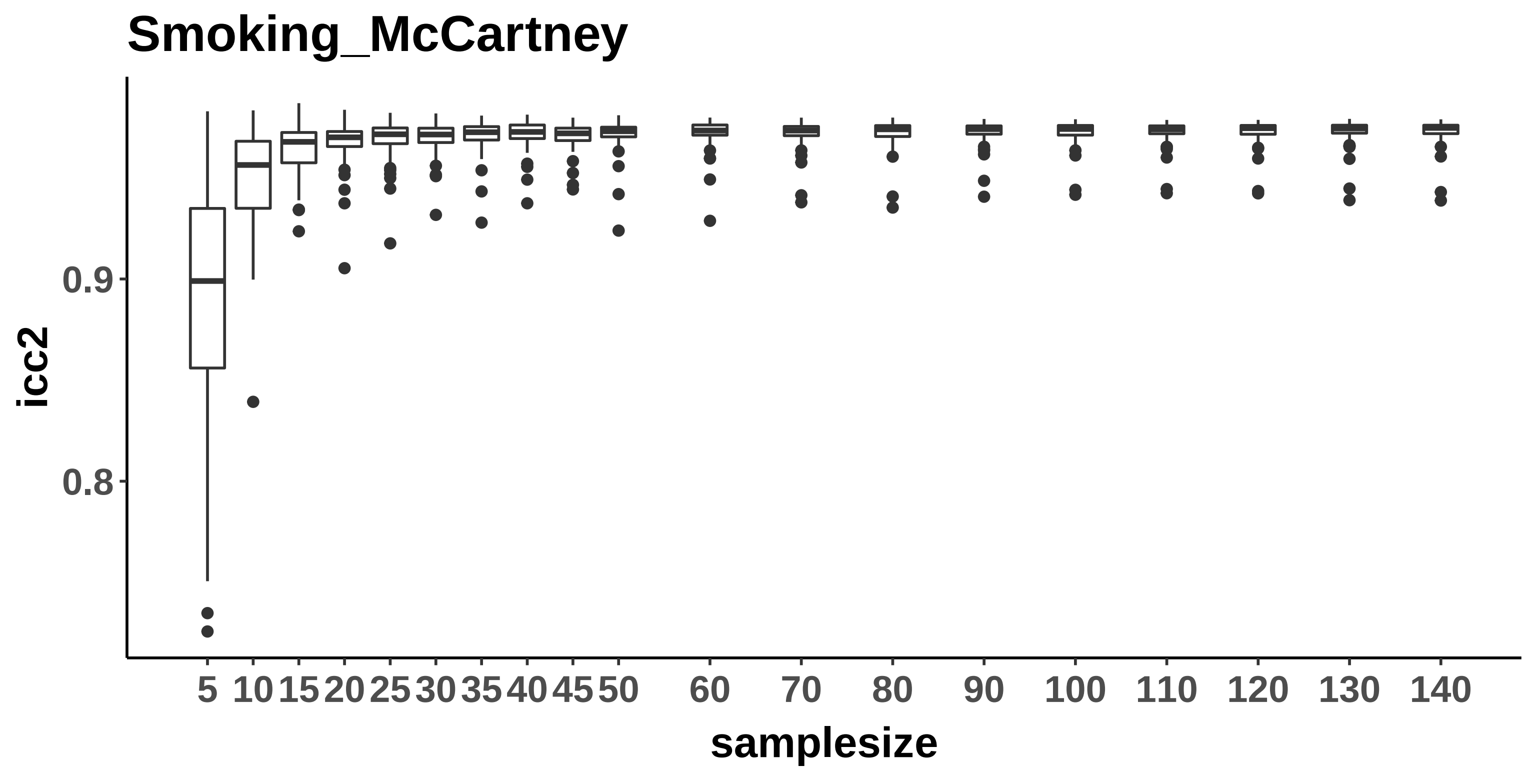

Smoking_McCartney
icc2
0.9
0.8
5
10
15
20
25
30
35
40
45
50
60
70
80
90
100
110
120
130
140
samplesize

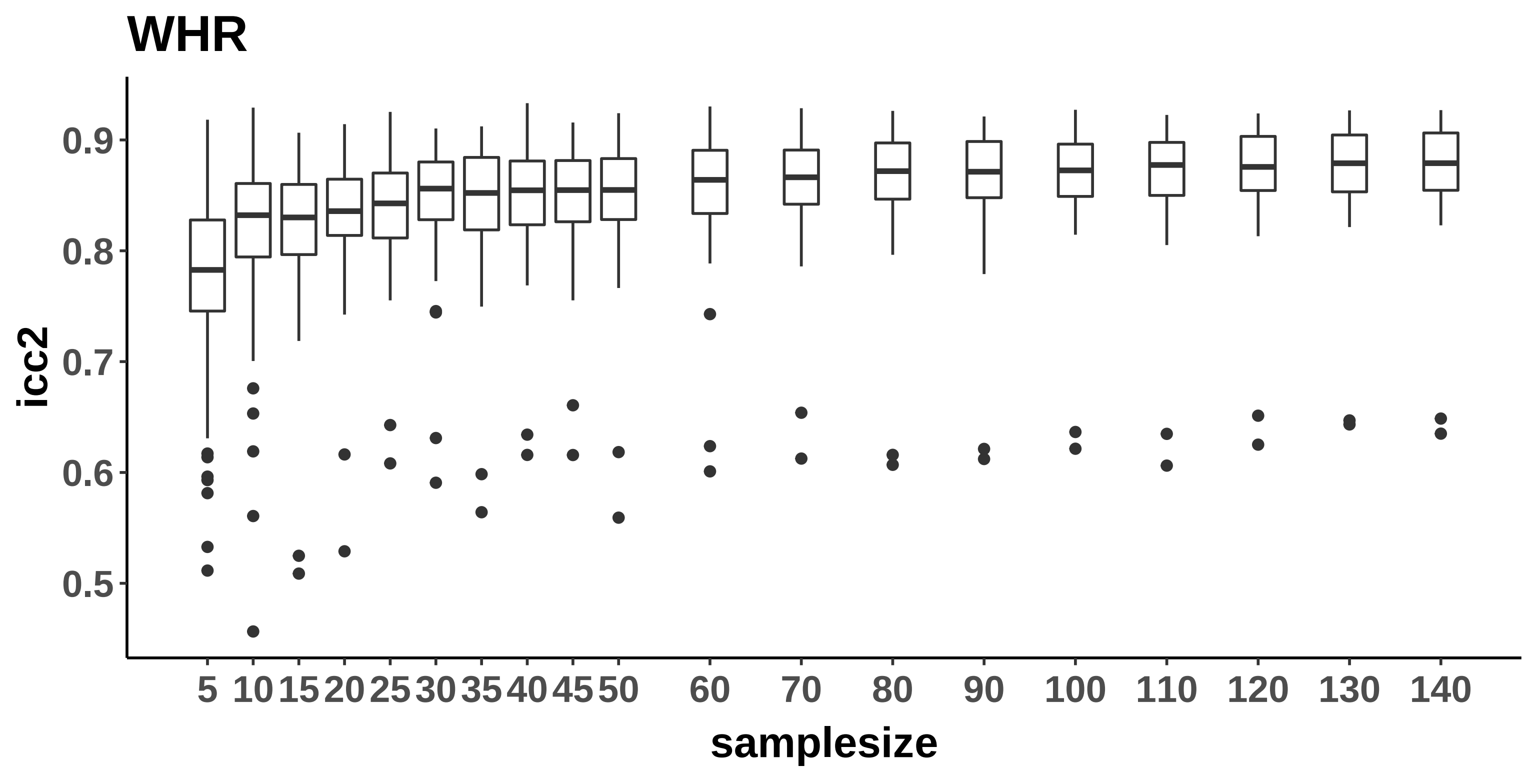

WHR
icc2
0.9
0.8
0.7
0.6
0.5
5
10
15
20
25
30
35
40
45
50
60
70
80
90
100
110
120
130
140
samplesize

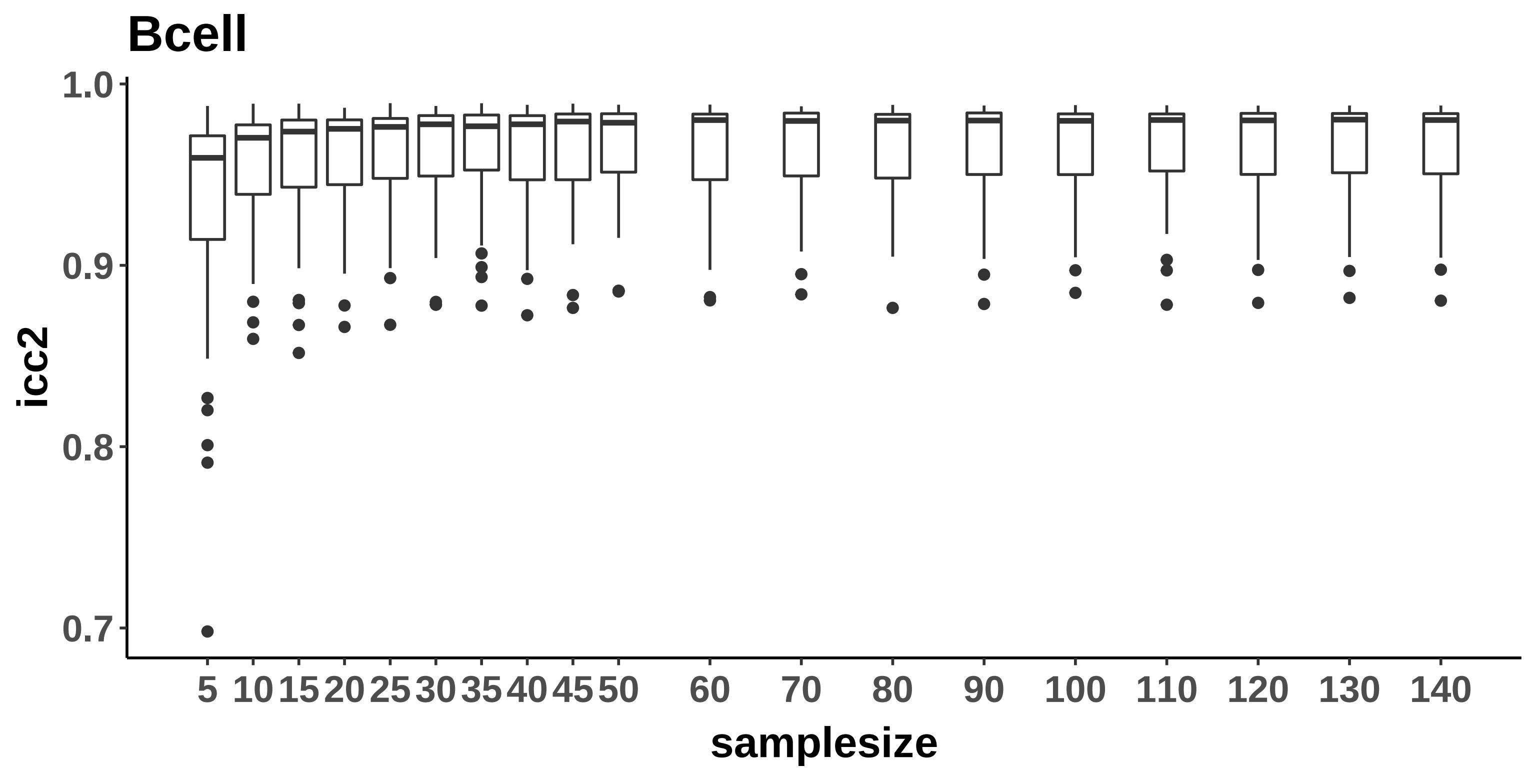

Bcell
1.0
0.9
0.8
0.7
icc2
5
10
15
20
25
30
35
40
45
50
60
70
80
90
100
110
120
130
140
samplesize

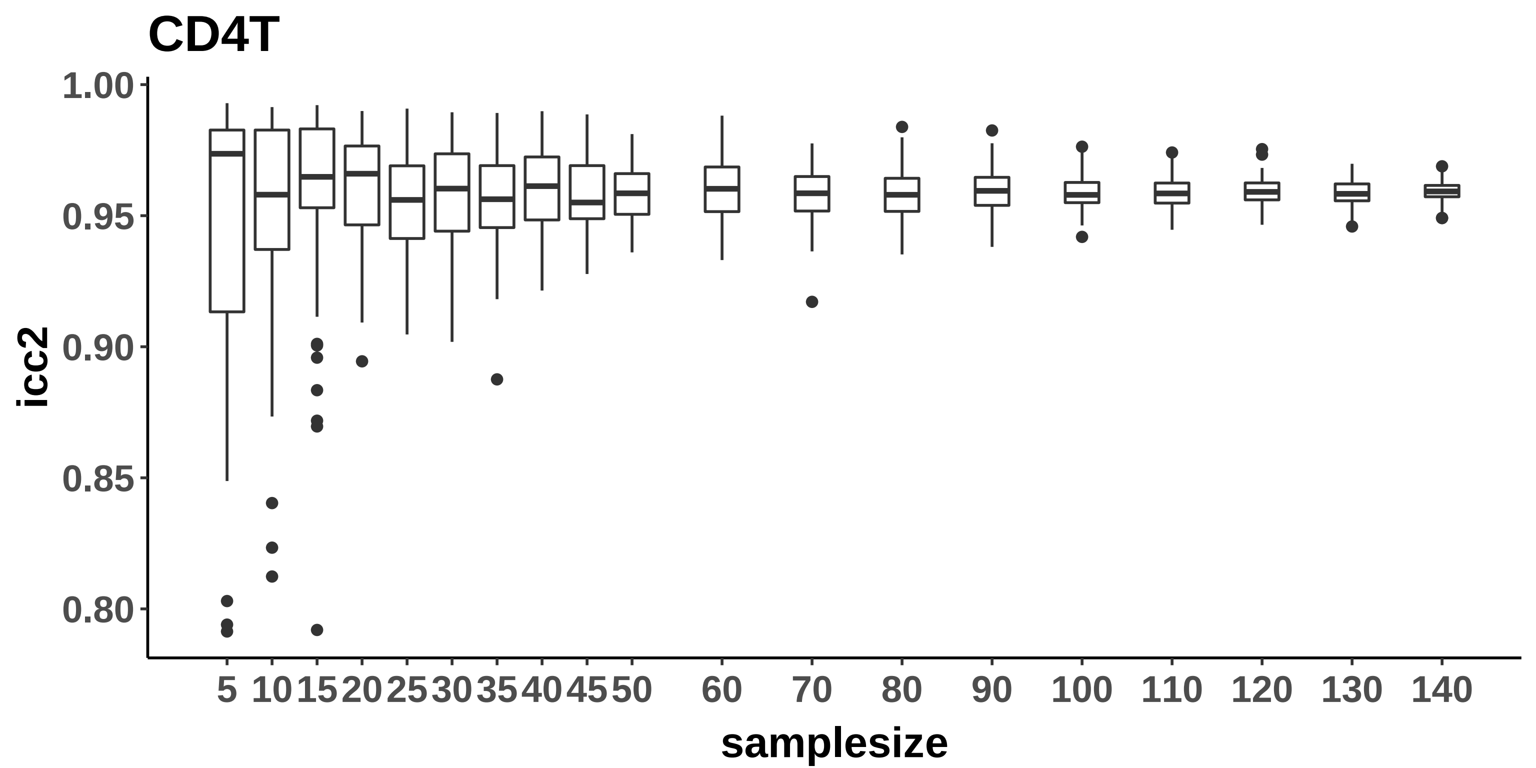

CD4T
1.00
0.95
0.90
0.85
0.80
icc2
5 10 15 20 25 30 35 40 45 50 60 70 80 90 100 110 120 130 140
samplesize

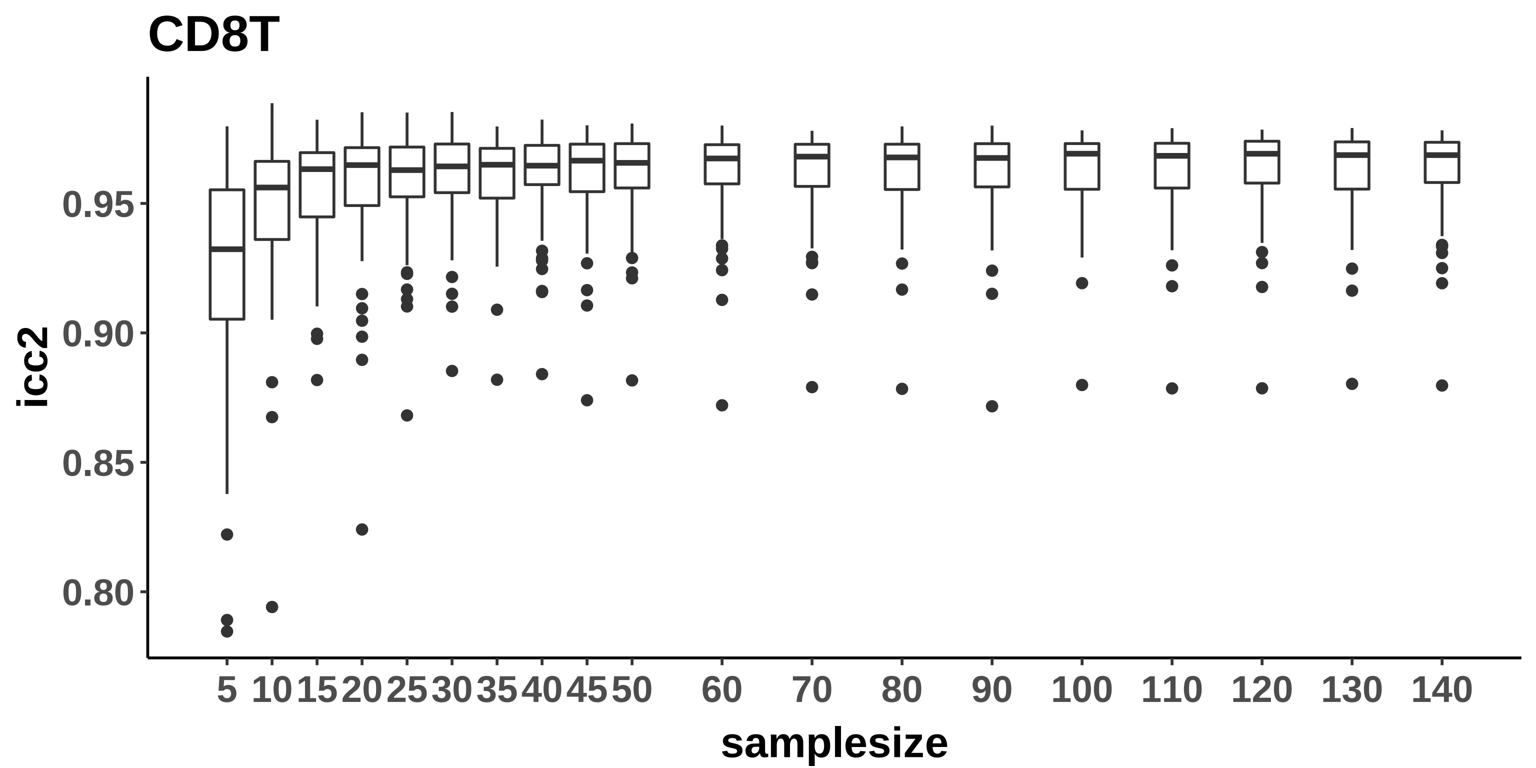
CD8T
icc2
0.95
0.90
0.85
0.80
5
10
15
20
25
30
35
40
45
50
60
70
80
90
100
110
120
130
140
samplesize

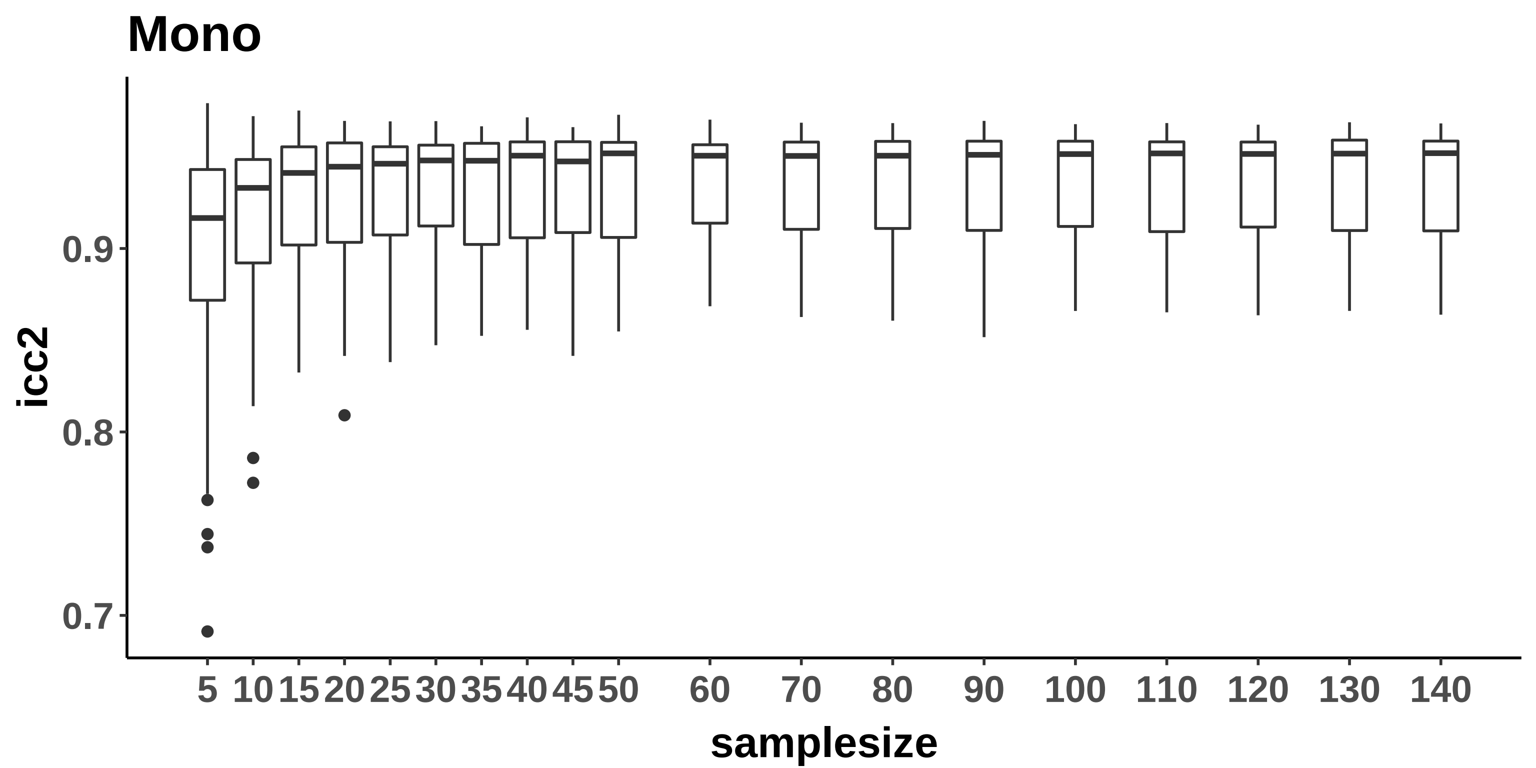

Mono
icc2
0.9
0.8
0.7
5
10
15
20
25
30
35
40
45
50
60
70
80
90
100
110
120
130
140
samplesize

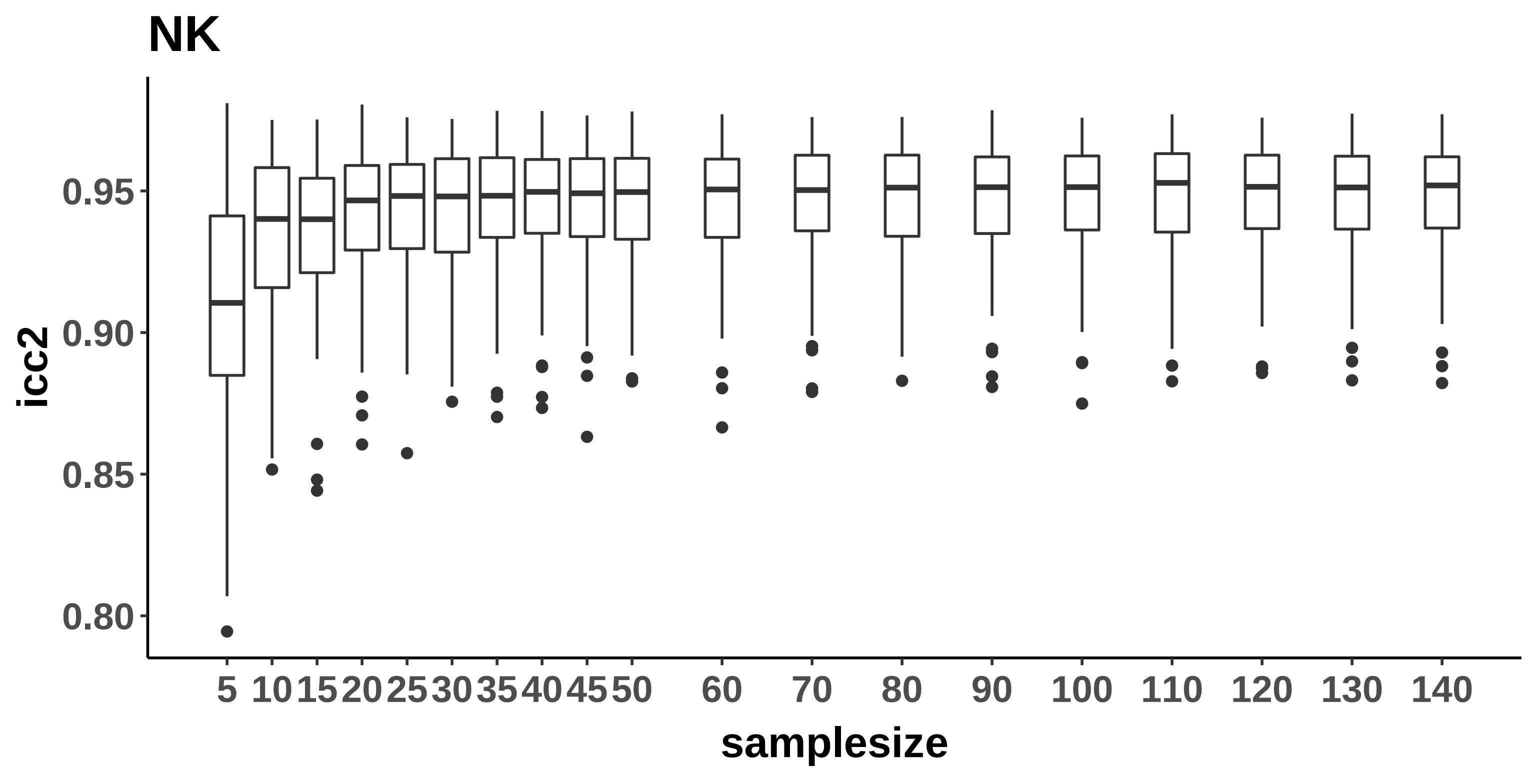

NK
icc2
0.95
0.90
0.85
0.80
5
10
15
20
25
30
35
40
45
50
60
70
80
90
100
110
120
130
140
samplesize

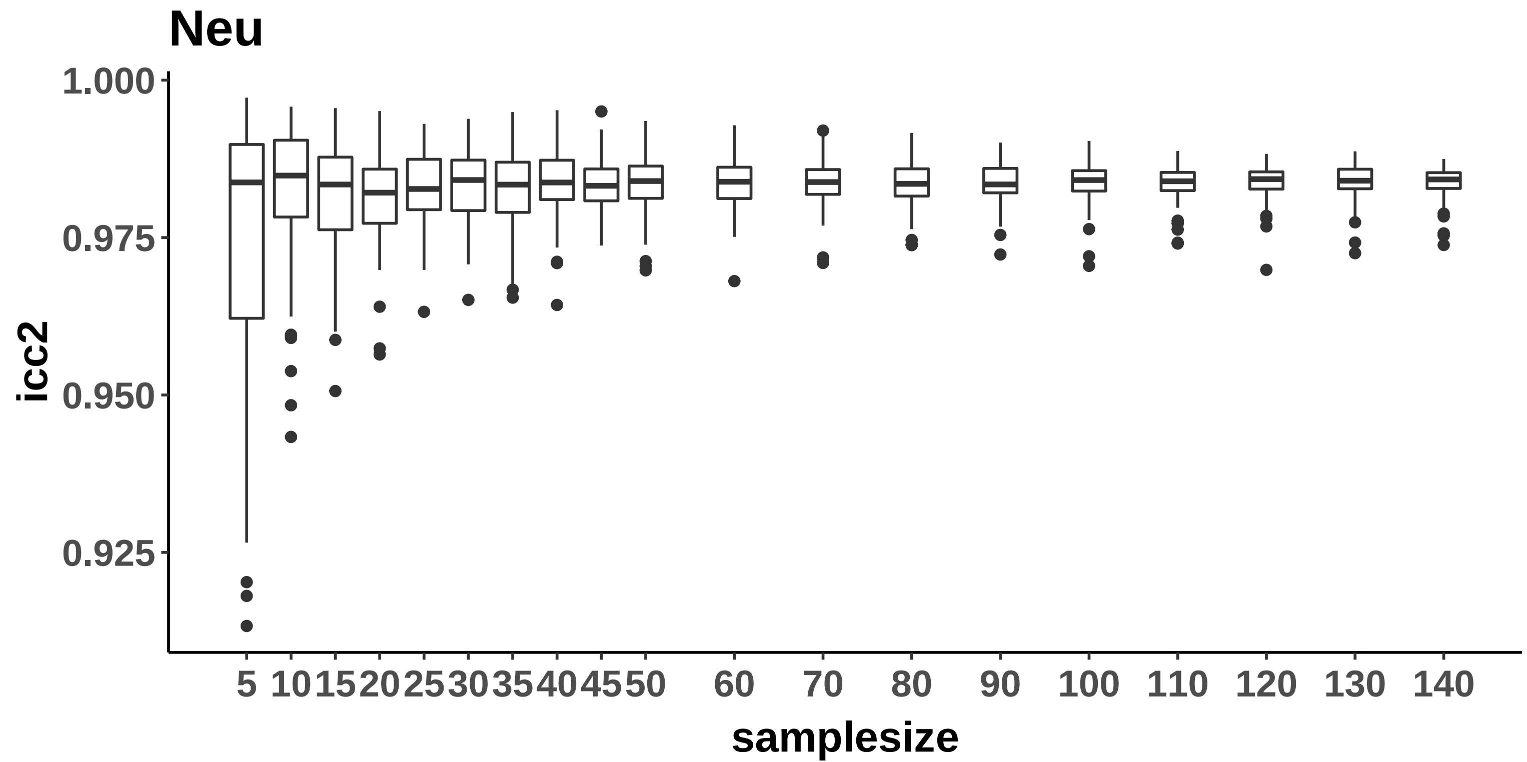

Neu
1.000
0.975
0.950
0.925
icc2
5
10
15
20
25
30
35
40
45
50
60
70
80
90
100
110
120
130
140
samplesize

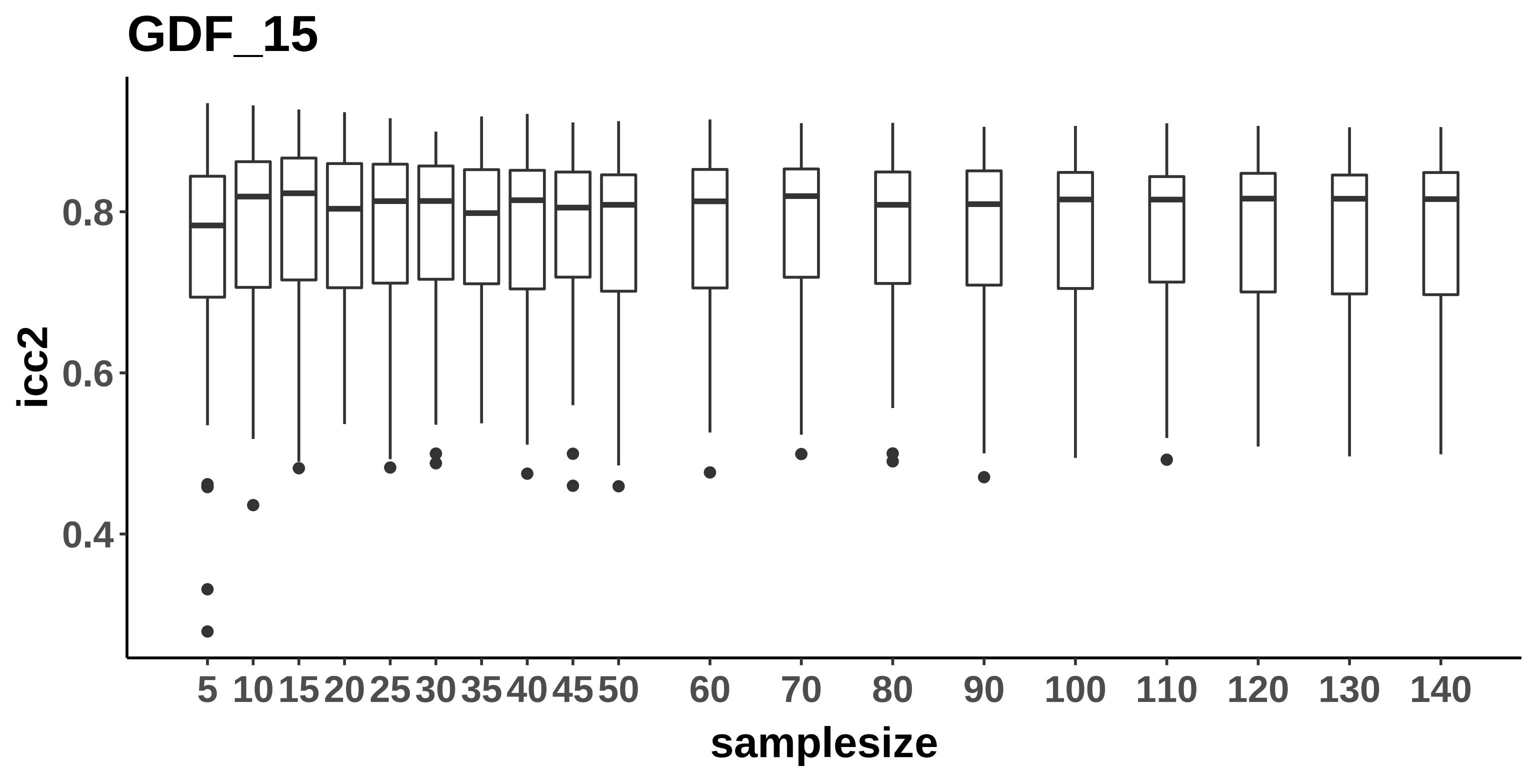
GDF_15
icc2
0.4
0.6
0.8
5
10
15
20
25
30
35
40
45
50
60
70
80
90
100
110
120
130
140
samplesize

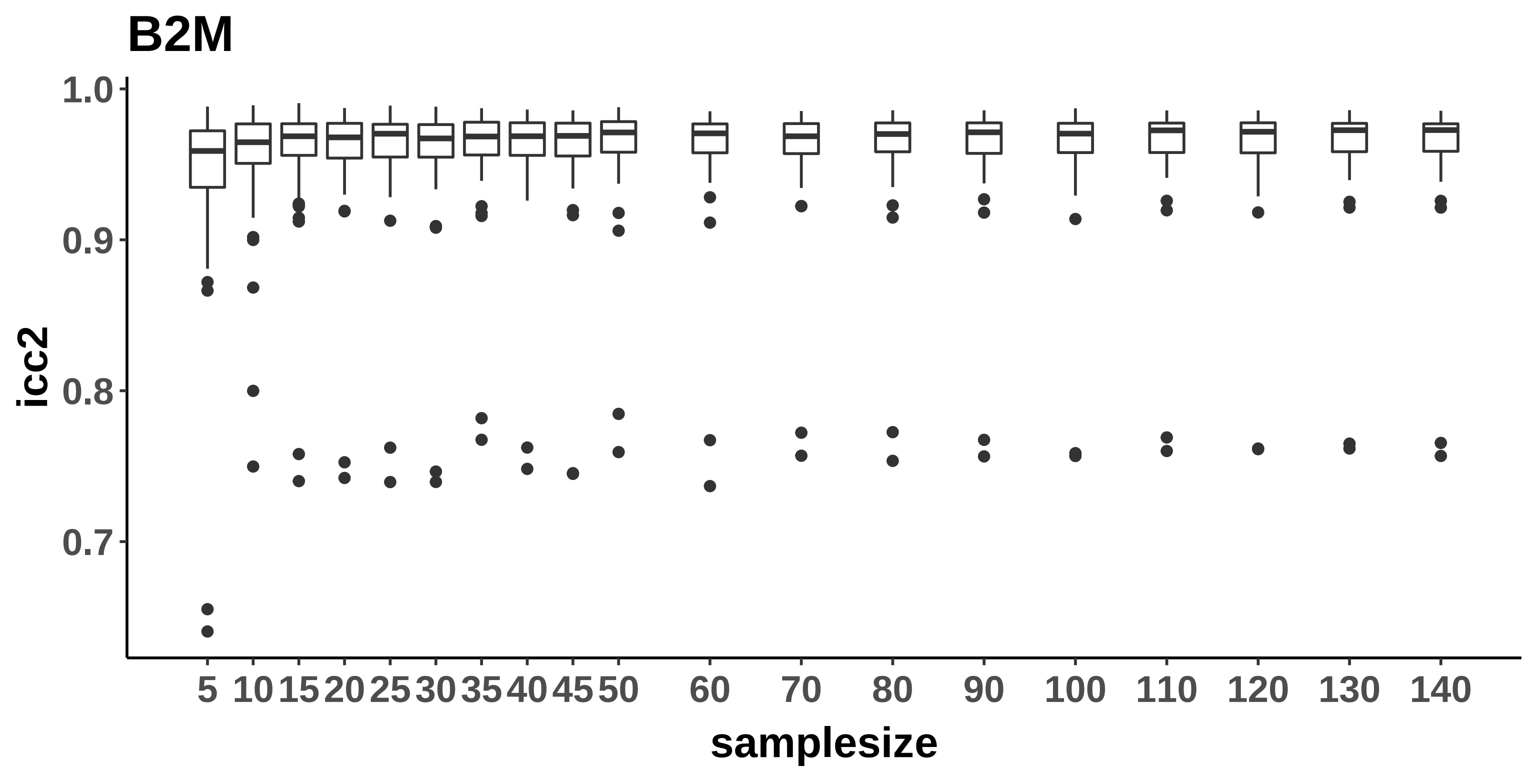
B2M
icc2
1.0
0.9
0.8
0.7
5
10
15
20
25
30
35
40
45
50
60
70
80
90
100
110
120
130
140
samplesize

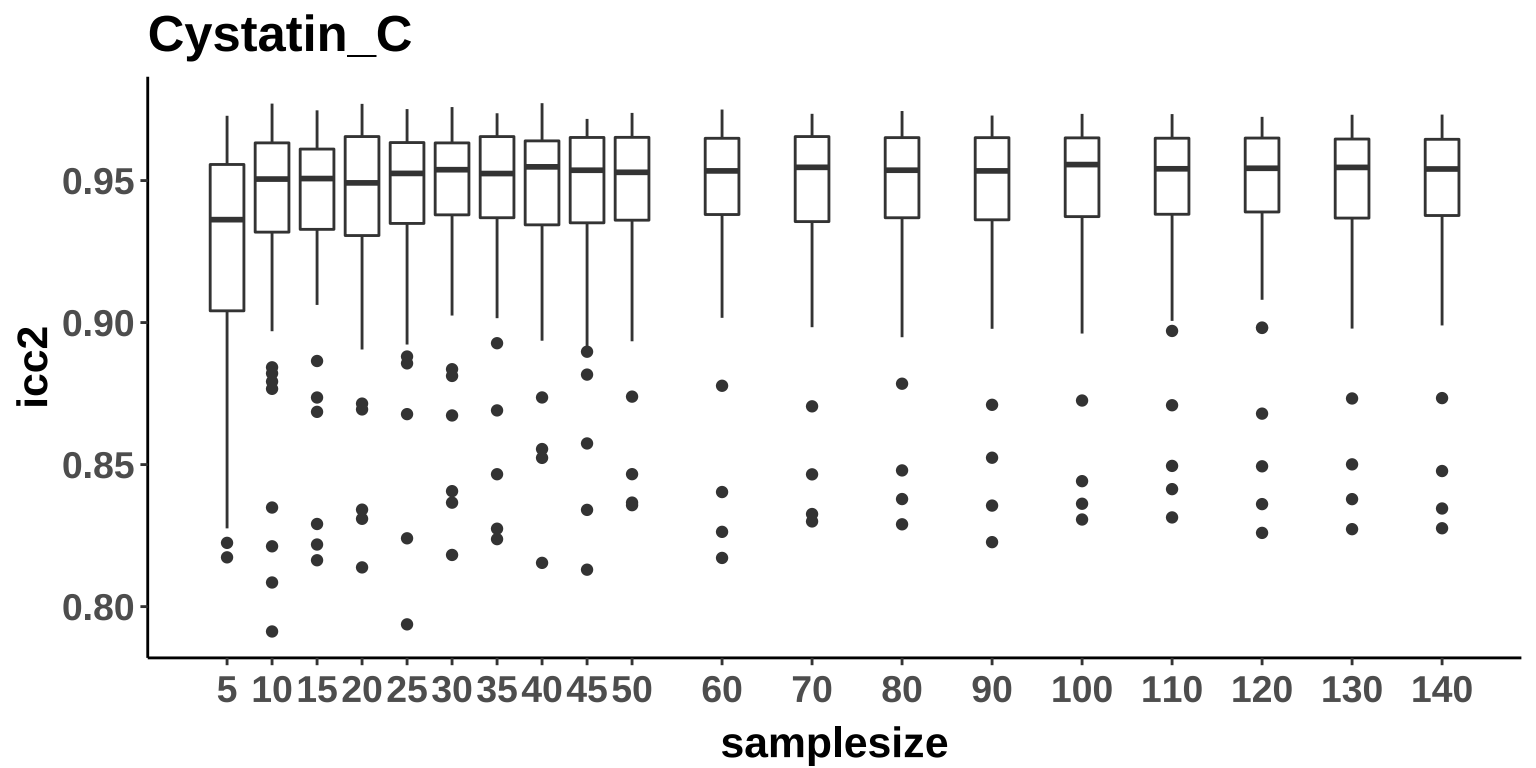
Cystatin_C
icc2
0.95
0.90
0.85
0.80
5 10 15 20 25 30 35 40 45 50 60 70 80 90 100 110 120 130 140
samplesize

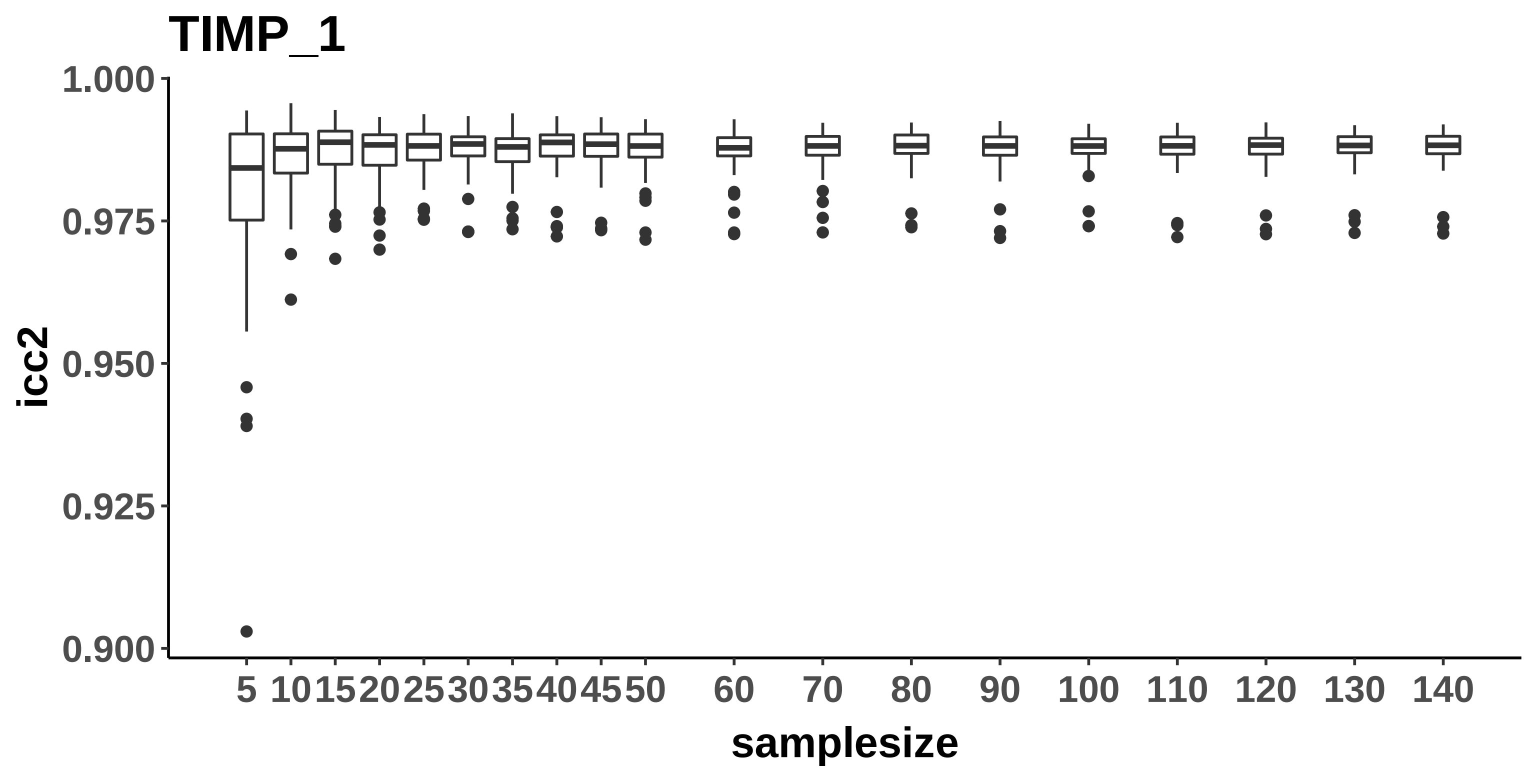
TIMP_1
icc2
1.000
0.975
0.950
0.925
0.900
5
10
15
20
25
30
35
40
45
50
60
70
80
90
100
110
120
130
140
samplesize

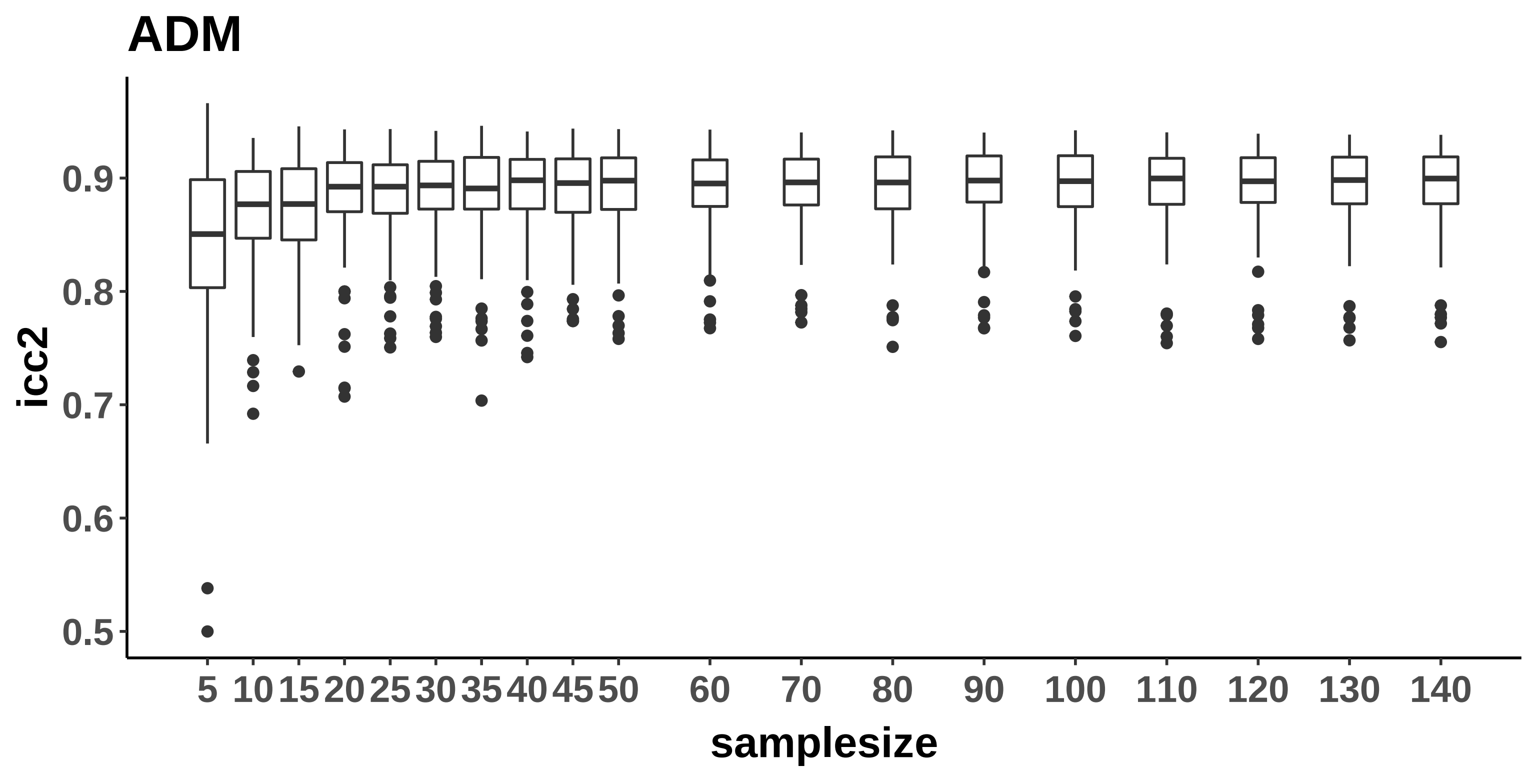
ADM
icc2
0.9
0.8
0.7
0.6
0.5
5
10
15
20
25
30
35
40
45
50
60
70
80
90
100
110
120
130
140
samplesize

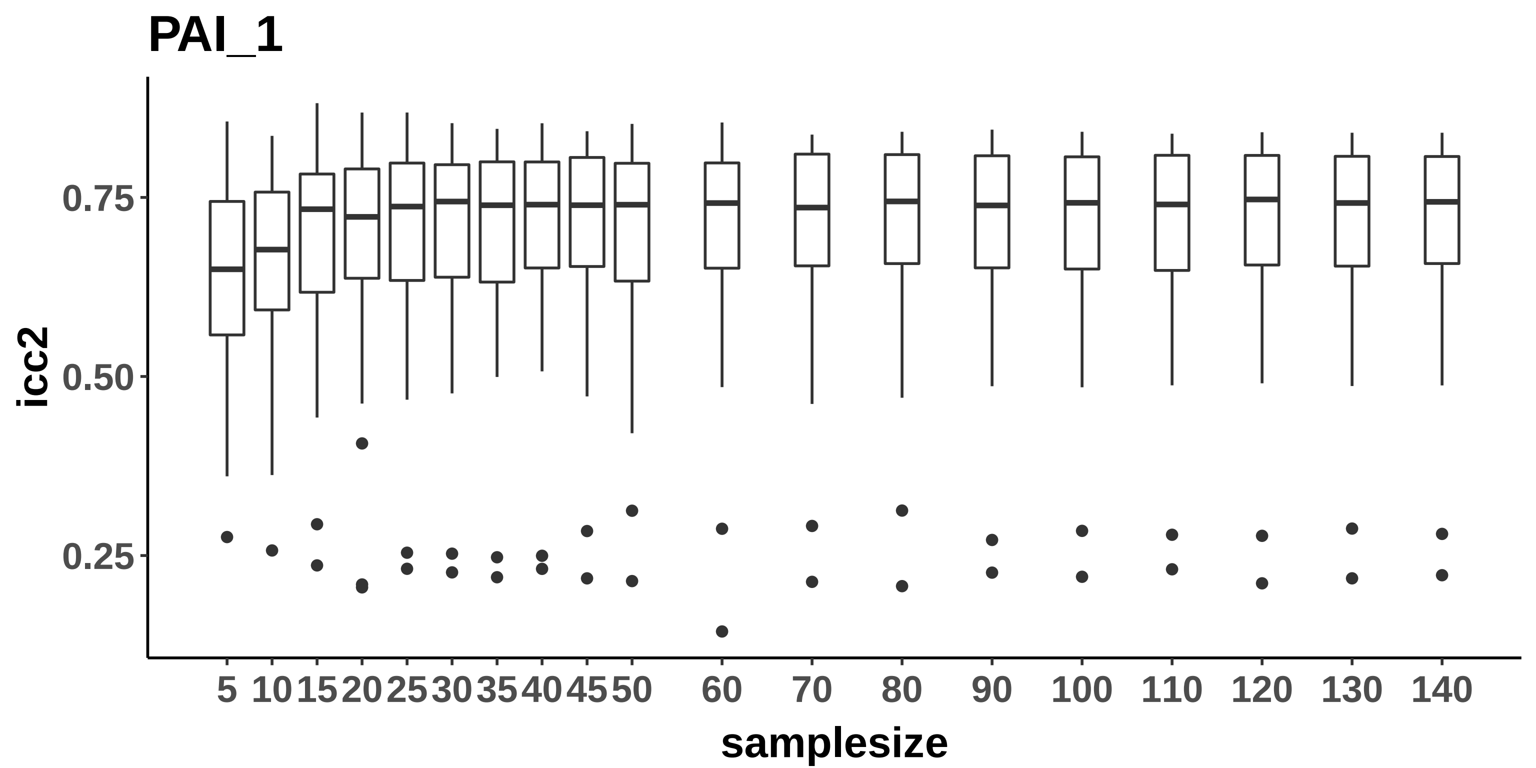
PAI_1
icc2
0.75
0.50
0.25
5
10
15
20
25
30
35
40
45
50
60
70
80
90
100
110
120
130
140
samplesize

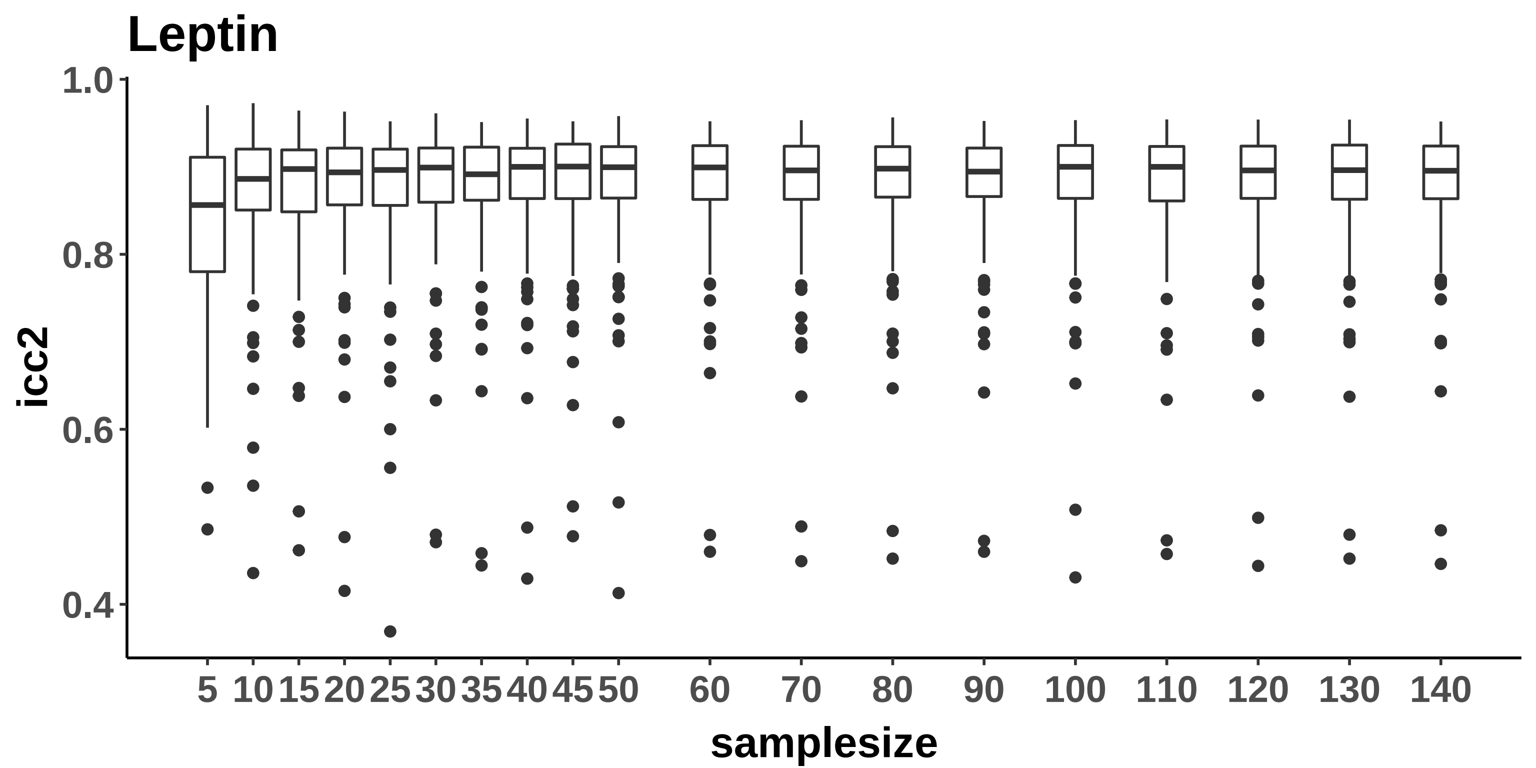

Leptin
1.0
0.8
0.6
0.4
icc2
5
10
15
20
25
30
35
40
45
50
60
70
80
90
100
110
120
130
140
samplesize

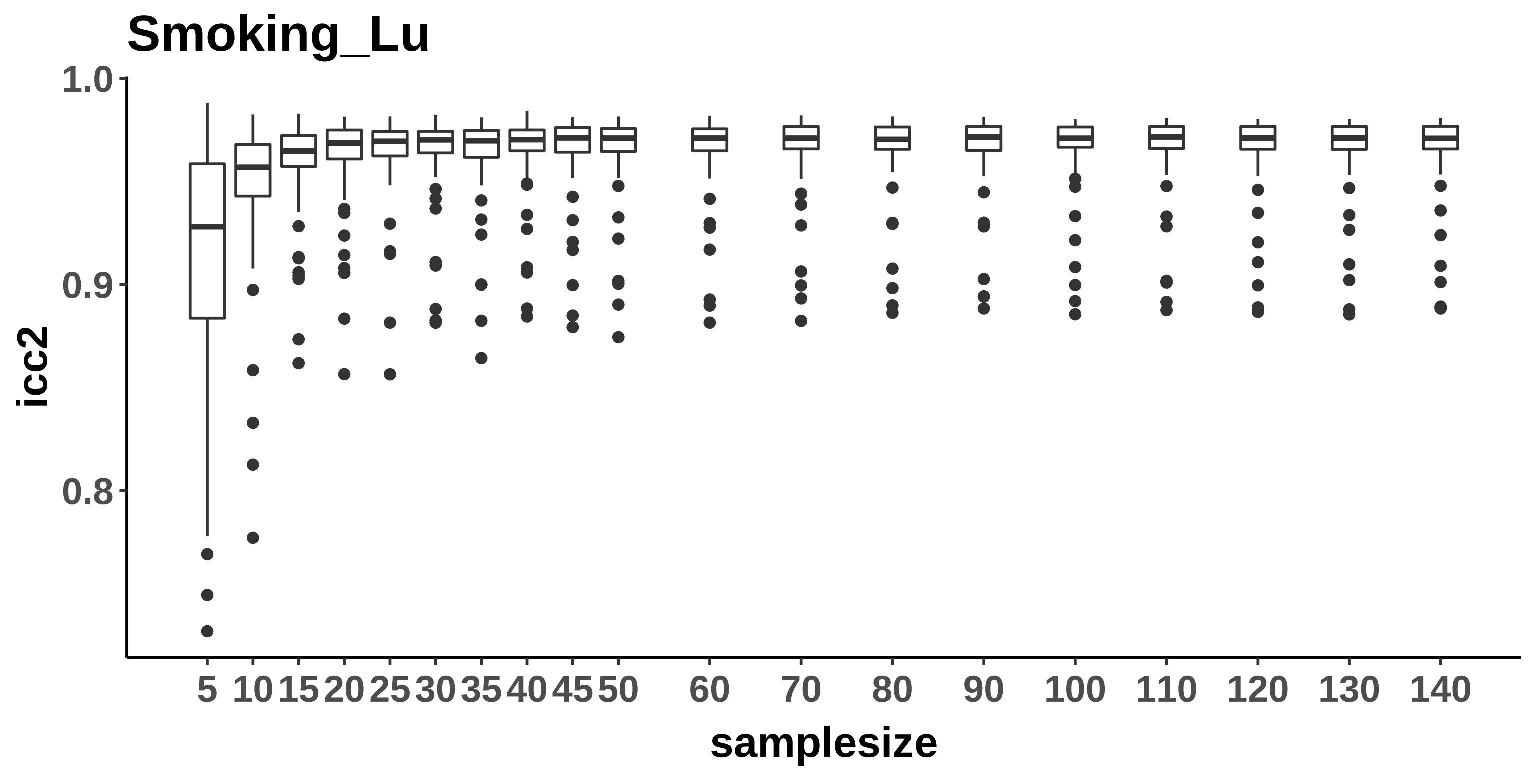
Smoking_Lu
1.0
0.9
0.8
icc2
5
10
15
20
25
30
35
40
45
50
60
70
80
90
100
110
120
130
140
samplesize

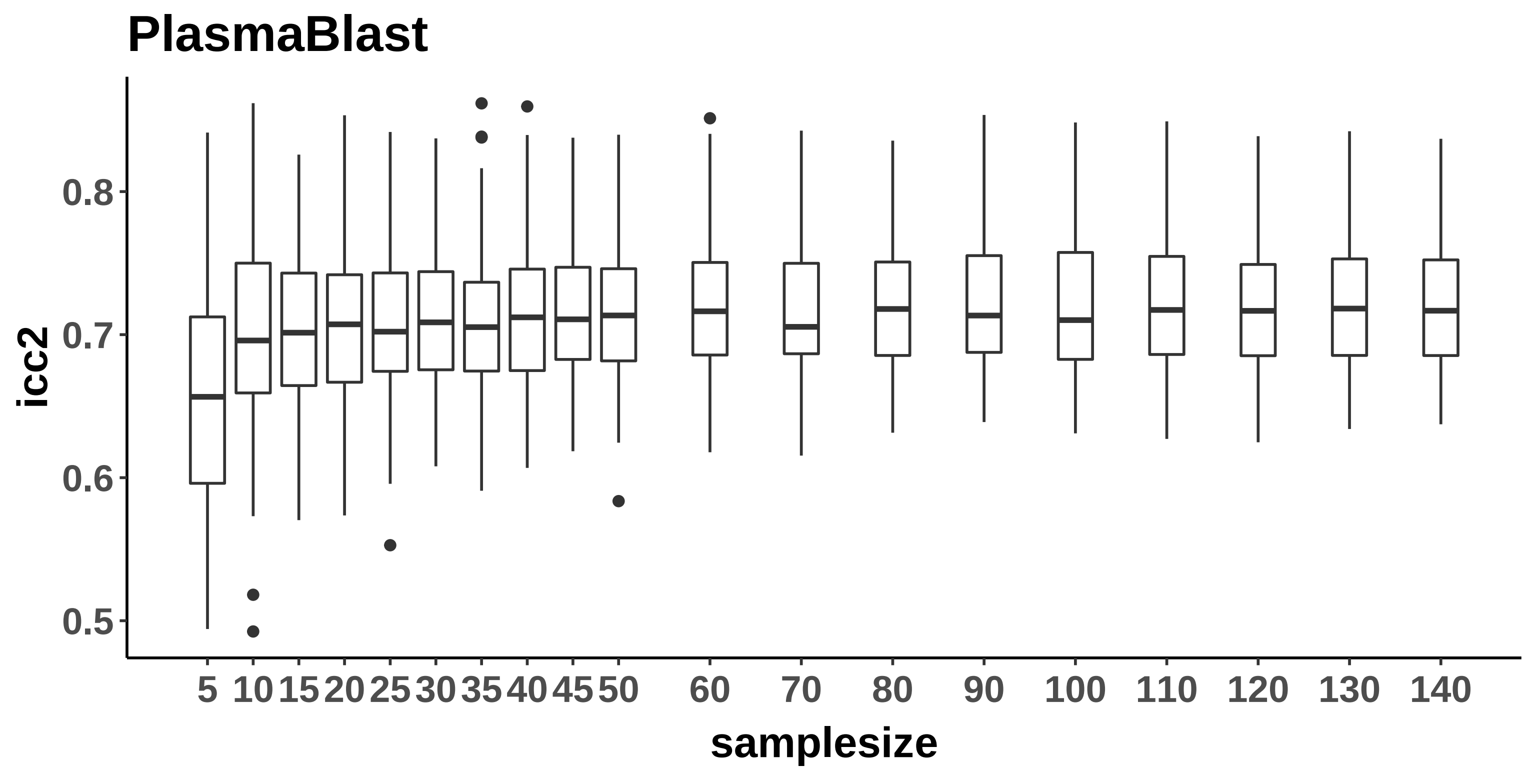
PlasmaBlast
icc2
0.8
0.7
0.6
0.5
5
10
15
20
25
30
35
40
45
50
60
70
80
90
100
110
120
130
140
samplesize

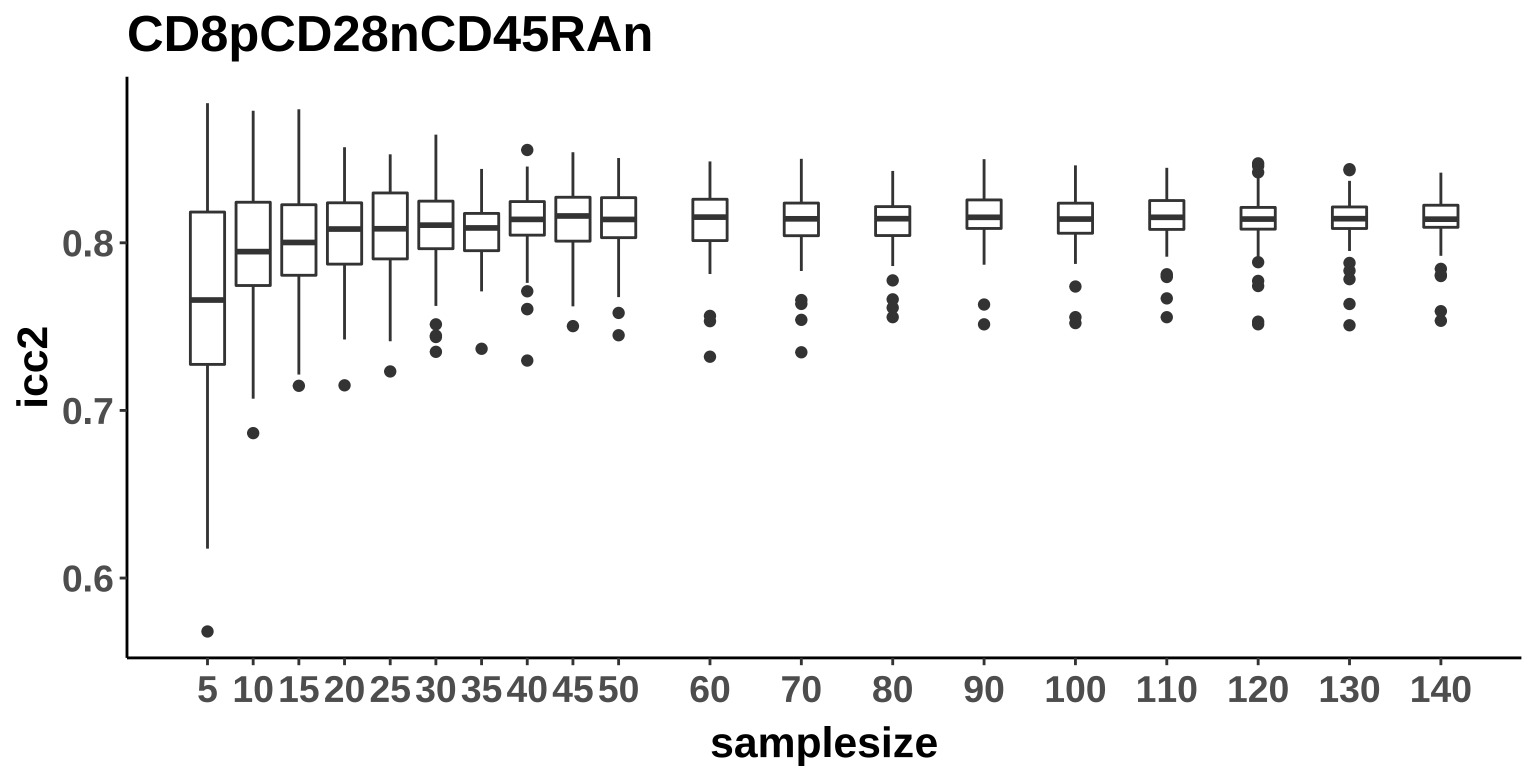
CD8pCD28nCD45RAn
icc2
0.8
0.7
0.6
5
10
15
20
25
30
35
40
45
50
60
70
80
90
100
110
120
130
140
samplesize

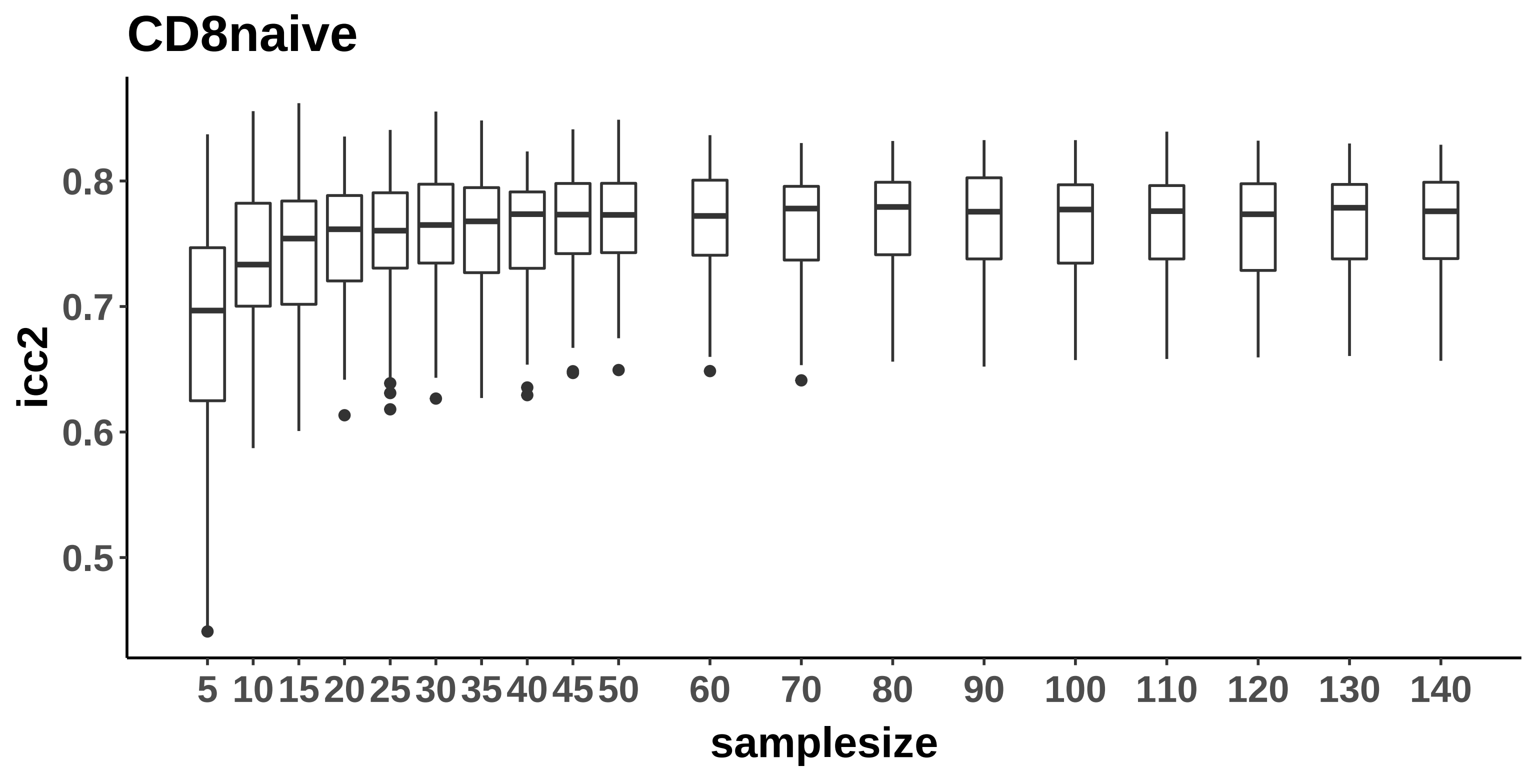
CD8naive
icc2
0.8
0.7
0.6
0.5
samplesize
5
10
15
20
25
30
35
40
45
50
60
70
80
90
100
110
120
130
140

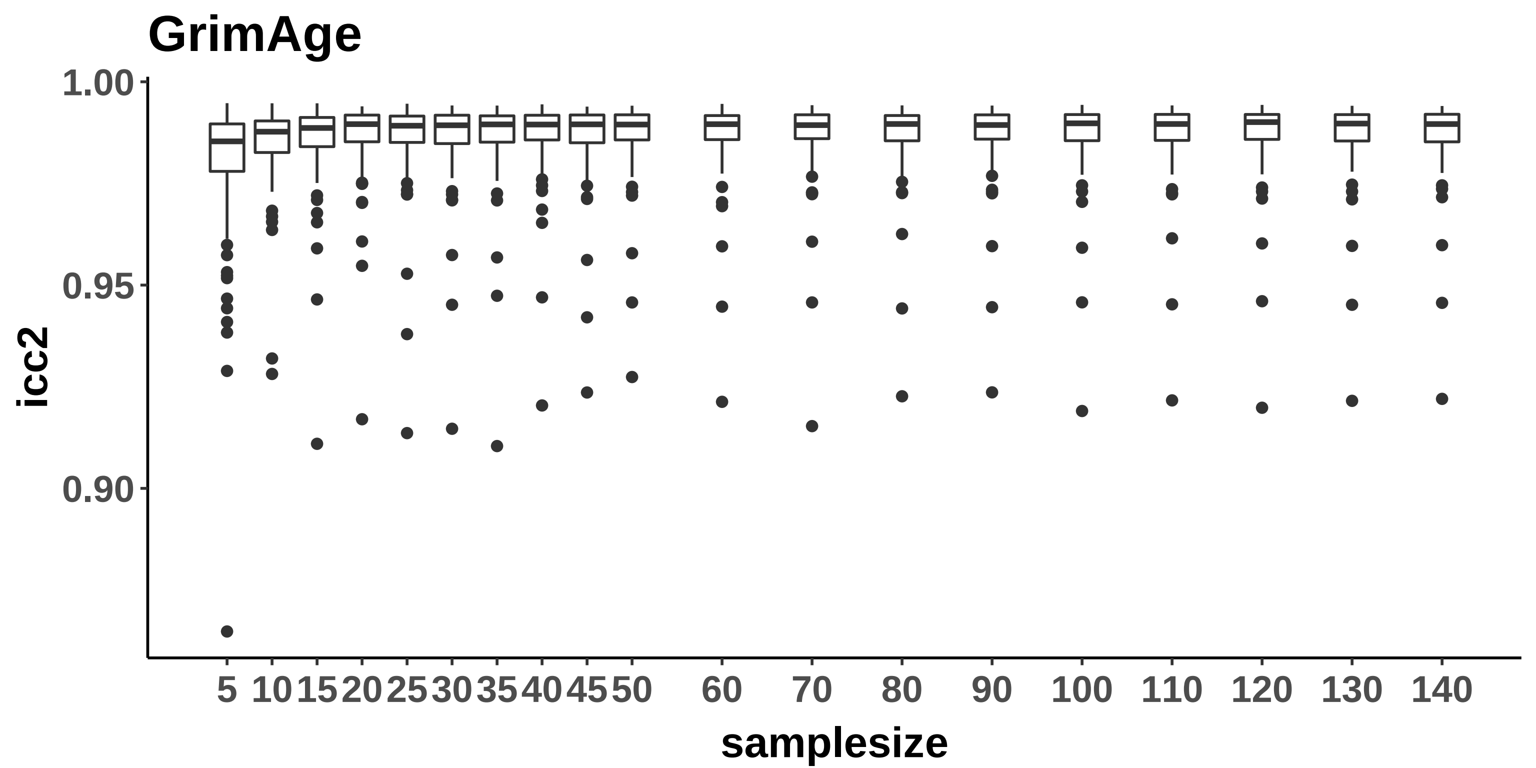
GrimAge
1.00
0.95
0.90
icc2
5
10
15
20
25
30
35
40
45
50
60
70
80
90
100
110
120
130
140
samplesize

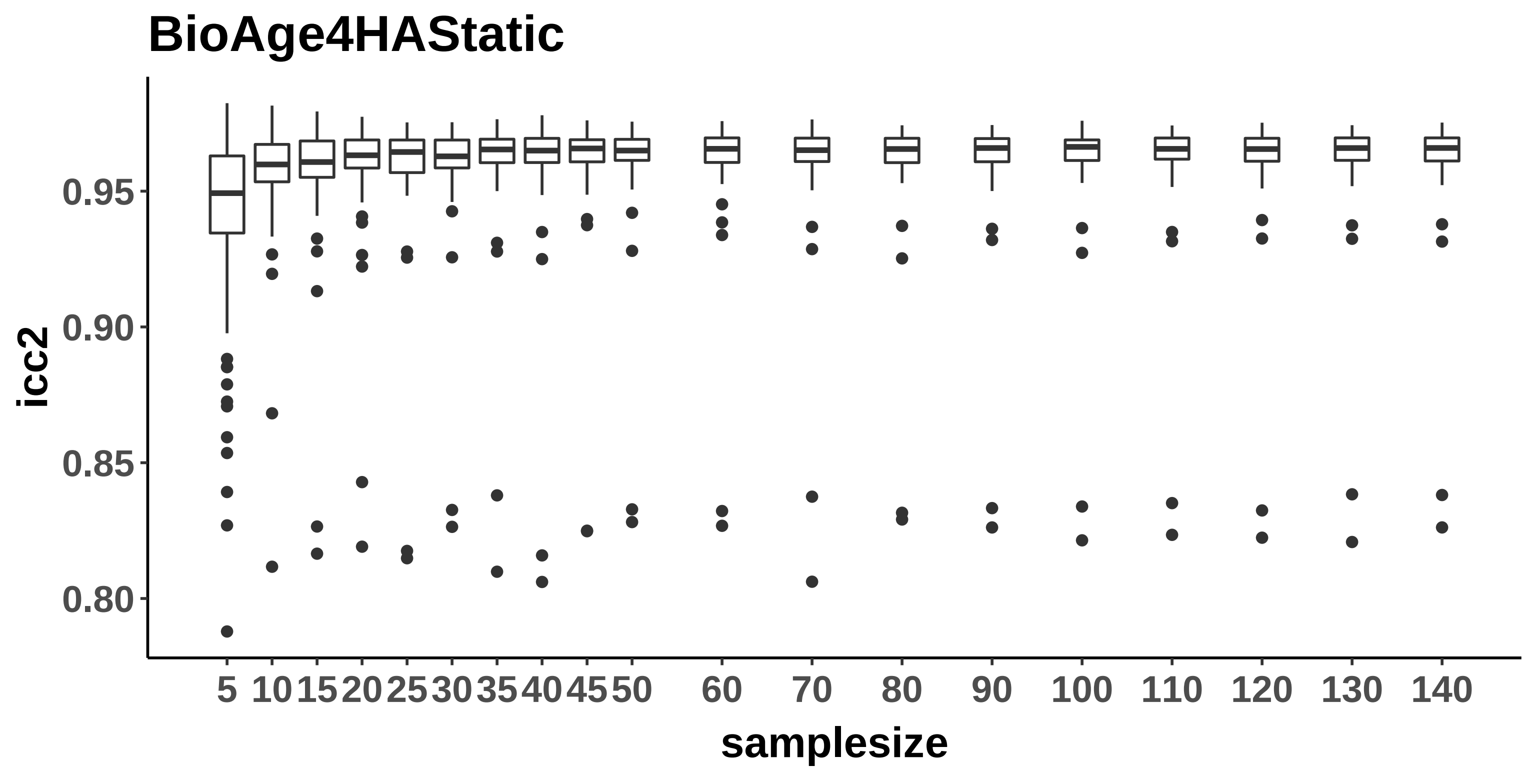
BioAge4HAStatic
icc2
0.95
0.90
0.85
0.80
samplesize
5 10 15 20 25 30 35 40 45 50 60 70 80 90 100 110 120 130 140
