## Supplemental Table 1 for "A systematic evaluation of 41 DNA methylation predictors across 101 data preprocessing and normalization strategies highlights considerable variation in algorithm performance"

|  | Replicate Sample | General Sample |
| --- | --- | --- |
| Individuals (N) | 146 | 1,761 |
| Age (years) | 57.4 (SD = 12.3) | 56.1 |
| Female (%) | 62.6 % | 62.2 % |
| Death (%) | - | 15.1% |
| EPIC arrays (N) | 292 | 1,761 |

**Table S1. Cohort characteristic of JHS.** In total, 1,907 individual and 2,053 850 EPIC array samples were included in our analysis. Our analyses were conducted in two subsets of this cohort, a subset that consist of only the technical replicate samples (n=146 individuals) and a subset that contains the remainder of the cohort. Shown above are standard cohort characteristics, including the percentage of individuals that died after follow-up for the general sample.
