## Supplemental Table 2 for "A systematic evaluation of 41 DNA methylation predictors across 101 data preprocessing and normalization strategies highlights considerable variation in algorithm performance"

**Table S2. Overview of data processing and normalization pipelines implemented.**

| **Pipeline number** | **Pipeline name** | **R package** | **Description** |
| --- | --- | --- | --- |
| 1 | enmix_oob_mean_nonorm_norcp | ENmix | Uses out-of-band Infinium I intensities to estimate normal distribution parameters to model background noise (bgParaEst="oob"). Dye bias correction applied based on averaged red/green ratio (dyeCorr="mean"). No normalization applied. No probe type bias correction (RCP) applied. |
| 2 | enmix_est_mean_nonorm_norcp | ENmix | Uses combined methylated and unmethylated intensities to estimate background distribution parameters separately for each color channel and each probe type (bgParaEst="est"). Dye bias correction applied based on averaged red/green ratio (dyeCorr="mean"). No normalization applied. No probe type bias correction (RCP) applied. |
| 3 | enmix_neg_mean_nonorm_norcp | ENmix | Uses 600 chip internal controls probes to estimate background distribution parameters (bgParaEst="neg"). Dye bias correction applied based on averaged red/green ratio (dyeCorr="mean"). No normalization applied. No probe type bias correction (RCP) applied. |
| 4 | enmix_oob_relic_nonorm_norcp | ENmix | Uses out-of-band Infinium I intensities to estimate normal distribution parameters to model background noise (bgParaEst="oob"). Dye bias correction applied based on REgression on Logarithm of Internal Control probes (dyeCorr="RELIC"). No normalization applied. No probe type bias correction (RCP) applied. |
| 5 | enmix_est_relic_nonorm_norcp | ENmix | Uses combined methylated and unmethylated intensities to estimate background distribution parameters separately for each color channel and each probe type (bgParaEst="est"). Dye bias correction applied based on REgression on Logarithm of Internal Control probes (dyeCorr="RELIC"). No normalization applied. No probe type bias correction (RCP) applied. |
| 6 | enmix_neg_relic_nonorm_norcp | ENmix | Uses 600 chip internal controls probes to estimate background distribution parameters (bgParaEst="neg"). Dye bias correction applied based on REgression on Logarithm of Internal Control probes (dyeCorr="RELIC"). No normalization applied. No probe type bias correction (RCP) applied. |
| 7 | enmix_oob_nodye_nonorm_norcp | ENmix | Uses out-of-band Infinium I intensities to estimate normal distribution parameters to model background noise (bgParaEst="oob"). No dye bias correction applied (dyeCorr="none"). No normalization applied. No probe type bias correction (RCP) applied. |
| 8 | enmix_est_nodye_nonorm_norcp | ENmix | Uses combined methylated and unmethylated intensities to estimate background distribution parameters separately for each color channel and each probe type (bgParaEst="est"). No dye bias correction applied (dyeCorr="none"). No normalization applied. No probe type bias correction (RCP) applied. |
| 9 | enmix_neg_nodye_nonorm_norcp | ENmix | Uses 600 chip internal controls probes to estimate background distribution parameters (bgParaEst="neg"). No dye bias correction applied (dyeCorr="none"). No normalization applied. No probe type bias correction (RCP) applied. |
| 10 | enmix_oob_mean_q1_norcp | ENmix | Uses out-of-band Infinium I intensities to estimate normal distribution parameters to model background noise (bgParaEst="oob"). Dye bias correction applied based on averaged red/green ratio (dyeCorr="mean"). Quantile normalization applied separately for methylated and unmethylated intensities of Infinium I and II probes (method="quantile1"). No probe type bias correction (RCP) applied. |
| 11 | enmix_est_mean_q1_norcp | ENmix | Uses combined methylated and unmethylated intensities to estimate background distribution parameters separately for each color channel and each probe type (bgParaEst="est"). Dye bias correction applied based on averaged red/green ratio (dyeCorr="mean"). Quantile normalization applied separately for methylated and unmethylated intensities of Infinium I and II probes (method="quantile1"). No probe type bias correction (RCP) applied. |
| 12 | enmix_neg_mean_q1_norcp | ENmix | Uses 600 chip internal controls probes to estimate background distribution parameters (bgParaEst="neg"). Dye bias correction applied based on averaged red/green ratio (dyeCorr="mean"). Quantile normalization applied separately for methylated and unmethylated intensities of Infinium I and II probes (method="quantile1"). No probe type bias correction (RCP) applied. |
| 13 | enmix_oob_relic_q1_norcp | ENmix | Uses out-of-band Infinium I intensities to estimate normal distribution parameters to model background noise (bgParaEst="oob"). Dye bias correction applied based on REgression on Logarithm of Internal Control probes (dyeCorr="RELIC"). Quantile normalization applied separately for methylated and unmethylated intensities of Infinium I and II probes (method="quantile1"). No probe type bias correction (RCP) applied. |
| 14 | enmix_est_relic_q1_norcp | ENmix | Uses combined methylated and unmethylated intensities to estimate background distribution parameters separately for each color channel and each probe type (bgParaEst="est"). Dye bias correction applied based on REgression on Logarithm of Internal Control probes (dyeCorr="RELIC"). Quantile normalization applied separately for methylated and unmethylated intensities of Infinium I and II probes (method="quantile1"). No probe type bias correction (RCP) applied. |
| 15 | enmix_neg_relic_q1_norcp | ENmix | Uses 600 chip internal controls probes to estimate background distribution parameters (bgParaEst="neg"). Dye bias correction applied based on REgression on Logarithm of Internal Control probes (dyeCorr="RELIC"). Quantile normalization applied separately for methylated and unmethylated intensities of Infinium I and II probes (method="quantile1"). No probe type bias correction (RCP) applied. |
| 16 | enmix_oob_nodye_q1_norcp | ENmix | Uses out-of-band Infinium I intensities to estimate normal distribution parameters to model background noise (bgParaEst="oob"). No dye bias correction applied (dyeCorr="none"). Quantile normalization applied separately for methylated and unmethylated intensities of Infinium I and II probes (method="quantile1"). No probe type bias correction (RCP) applied. |
| 17 | enmix_est_nodye_q1_norcp | ENmix | Uses combined methylated and unmethylated intensities to estimate background distribution parameters separately for each color channel and each probe type (bgParaEst="est"). No dye bias correction applied (dyeCorr="none"). Quantile normalization applied separately for methylated and unmethylated intensities of Infinium I and II probes (method="quantile1"). No probe type bias correction (RCP) applied. |
| 18 | enmix_neg_nodye_q1_norcp | ENmix | Uses 600 chip internal controls probes to estimate background distribution parameters (bgParaEst="neg"). No dye bias correction applied (dyeCorr="none"). Quantile normalization applied separately for methylated and unmethylated intensities of Infinium I and II probes (method="quantile1"). No probe type bias correction (RCP) applied. |
| 19 | enmix_oob_mean_q2_norcp | ENmix | Uses out-of-band Infinium I intensities to estimate normal distribution parameters to model background noise (bgParaEst="oob"). Dye bias correction applied based on averaged red/green ratio (dyeCorr="mean"). Quantile normalization applied on combined methylated or unmethylated intensities for Infinium I or II probes (method="quantile2"). No probe type bias correction (RCP) applied. |
| 20 | enmix_est_mean_q2_norcp | ENmix | Uses combined methylated and unmethylated intensities to estimate background distribution parameters separately for each color channel and each probe type (bgParaEst="est"). Dye bias correction applied based on averaged red/green ratio (dyeCorr="mean"). Quantile normalization applied on combined methylated or unmethylated intensities for Infinium I or II probes (method="quantile2"). No probe type bias correction (RCP) applied. |
| 21 | enmix_neg_mean_q2_norcp | ENmix | Uses 600 chip internal controls probes to estimate background distribution parameters (bgParaEst="neg"). Dye bias correction applied based on averaged red/green ratio (dyeCorr="mean"). Quantile normalization applied on combined methylated or unmethylated intensities for Infinium I or II probes (method="quantile2"). No probe type bias correction (RCP) applied. |
| 22 | enmix_oob_relic_q2_norcp | ENmix | Uses out-of-band Infinium I intensities to estimate normal distribution parameters to model background noise (bgParaEst="oob"). Dye bias correction applied based on REgression on Logarithm of Internal Control probes (dyeCorr="RELIC"). Quantile normalization applied on combined methylated or unmethylated intensities for Infinium I or II probes (method="quantile2"). No probe type bias correction (RCP) applied. |
| 23 | enmix_est_relic_q2_norcp | ENmix | Uses combined methylated and unmethylated intensities to estimate background distribution parameters separately for each color channel and each probe type (bgParaEst="est"). Dye bias correction applied based on REgression on Logarithm of Internal Control probes (dyeCorr="RELIC"). Quantile normalization applied on combined methylated or unmethylated intensities for Infinium I or II probes (method="quantile2"). No probe type bias correction (RCP) applied. |
| 24 | enmix_neg_relic_q2_norcp | ENmix | Uses 600 chip internal controls probes to estimate background distribution parameters (bgParaEst="neg"). Dye bias correction applied based on REgression on Logarithm of Internal Control probes (dyeCorr="RELIC"). Quantile normalization applied on combined methylated or unmethylated intensities for Infinium I or II probes (method="quantile2"). No probe type bias correction (RCP) applied. |
| 25 | enmix_oob_nodye_q2_norcp | ENmix | Uses out-of-band Infinium I intensities to estimate normal distribution parameters to model background noise (bgParaEst="oob"). No dye bias correction applied (dyeCorr="none"). Quantile normalization applied on combined methylated or unmethylated intensities for Infinium I or II probes (method="quantile2"). No probe type bias correction (RCP) applied. |
| 26 | enmix_est_nodye_q2_norcp | ENmix | Uses combined methylated and unmethylated intensities to estimate background distribution parameters separately for each color channel and each probe type (bgParaEst="est"). No dye bias correction applied (dyeCorr="none"). Quantile normalization applied on combined methylated or unmethylated intensities for Infinium I or II probes (method="quantile2"). No probe type bias correction (RCP) applied. |
| 27 | enmix_neg_nodye_q2_norcp | ENmix | Uses 600 chip internal controls probes to estimate background distribution parameters (bgParaEst="neg"). No dye bias correction applied (dyeCorr="none"). Quantile normalization applied on combined methylated or unmethylated intensities for Infinium I or II probes (method="quantile2"). No probe type bias correction (RCP) applied. |
| 28 | enmix_oob_mean_q3_norcp | ENmix | Uses out-of-band Infinium I intensities to estimate normal distribution parameters to model background noise (bgParaEst="oob"). Dye bias correction applied based on averaged red/green ratio (dyeCorr="mean"). Quantile normalization applied on combined methylated or unmethylated intensities for Infinium I and II probes together (method="quantile3"). No probe type bias correction (RCP) applied. |
| 29 | enmix_est_mean_q3_norcp | ENmix | Uses combined methylated and unmethylated intensities to estimate background distribution parameters separately for each color channel and each probe type (bgParaEst="est"). Dye bias correction applied based on averaged red/green ratio (dyeCorr="mean"). Quantile normalization applied on combined methylated or unmethylated intensities for Infinium I and II probes together (method="quantile3"). No probe type bias correction (RCP) applied. |
| 30 | enmix_neg_mean_q3_norcp | ENmix | Uses 600 chip internal controls probes to estimate background distribution parameters (bgParaEst="neg"). Dye bias correction applied based on averaged red/green ratio (dyeCorr="mean"). Quantile normalization applied on combined methylated or unmethylated intensities for Infinium I and II probes together (method="quantile3"). No probe type bias correction (RCP) applied. |
| 31 | enmix_oob_relic_q3_norcp | ENmix | Uses out-of-band Infinium I intensities to estimate normal distribution parameters to model background noise (bgParaEst="oob"). Dye bias correction applied based on REgression on Logarithm of Internal Control probes (dyeCorr="RELIC"). Quantile normalization applied on combined methylated or unmethylated intensities for Infinium I and II probes together (method="quantile3"). No probe type bias correction (RCP) applied. |
| 32 | enmix_est_relic_q3_norcp | ENmix | Uses combined methylated and unmethylated intensities to estimate background distribution parameters separately for each color channel and each probe type (bgParaEst="est"). Dye bias correction applied based on REgression on Logarithm of Internal Control probes (dyeCorr="RELIC"). Quantile normalization applied on combined methylated or unmethylated intensities for Infinium I and II probes together (method="quantile3"). No probe type bias correction (RCP) applied. |
| 33 | enmix_neg_relic_q3_norcp | ENmix | Uses 600 chip internal controls probes to estimate background distribution parameters (bgParaEst="neg"). Dye bias correction applied based on REgression on Logarithm of Internal Control probes (dyeCorr="RELIC"). Quantile normalization applied on combined methylated or unmethylated intensities for Infinium I and II probes together (method="quantile3"). No probe type bias correction (RCP) applied. |
| 34 | enmix_oob_nodye_q3_norcp | ENmix | Uses out-of-band Infinium I intensities to estimate normal distribution parameters to model background noise (bgParaEst="oob"). No dye bias correction applied (dyeCorr="none"). Quantile normalization applied on combined methylated or unmethylated intensities for Infinium I and II probes together (method="quantile3"). No probe type bias correction (RCP) applied. |
| 35 | enmix_est_nodye_q3_norcp | ENmix | Uses combined methylated and unmethylated intensities to estimate background distribution parameters separately for each color channel and each probe type (bgParaEst="est"). No dye bias correction applied (dyeCorr="none"). Quantile normalization applied on combined methylated or unmethylated intensities for Infinium I and II probes together (method="quantile3"). No probe type bias correction (RCP) applied. |
| 36 | enmix_neg_nodye_q3_norcp | ENmix | Uses 600 chip internal controls probes to estimate background distribution parameters (bgParaEst="neg"). No dye bias correction applied (dyeCorr="none"). Quantile normalization applied on combined methylated or unmethylated intensities for Infinium I and II probes together (method="quantile3"). No probe type bias correction (RCP) applied. |
| 37 | enmix_oob_mean_nonorm_rcp | ENmix | Uses out-of-band Infinium I intensities to estimate normal distribution parameters to model background noise (bgParaEst="oob"). Dye bias correction applied based on averaged red/green ratio (dyeCorr="mean"). No normalization applied. Probe design type bias correction applied using Regression on Correlated Probes (RCP) method. |
| 38 | enmix_est_mean_nonorm_rcp | ENmix | Uses combined methylated and unmethylated intensities to estimate background distribution parameters separately for each color channel and each probe type (bgParaEst="est"). Dye bias correction applied based on averaged red/green ratio (dyeCorr="mean"). No normalization applied. Probe design type bias correction applied using Regression on Correlated Probes (RCP) method. |
| 39 | enmix_neg_mean_nonorm_rcp | ENmix | Uses 600 chip internal controls probes to estimate background distribution parameters (bgParaEst="neg"). Dye bias correction applied based on averaged red/green ratio (dyeCorr="mean"). No normalization applied. Probe design type bias correction applied using Regression on Correlated Probes (RCP) method. |
| 40 | enmix_oob_relic_nonorm_rcp | ENmix | Uses out-of-band Infinium I intensities to estimate normal distribution parameters to model background noise (bgParaEst="oob"). Dye bias correction applied based on REgression on Logarithm of Internal Control probes (dyeCorr="RELIC"). No normalization applied. Probe design type bias correction applied using Regression on Correlated Probes (RCP) method. |
| 41 | enmix_est_relic_nonorm_rcp | ENmix | Uses combined methylated and unmethylated intensities to estimate background distribution parameters separately for each color channel and each probe type (bgParaEst="est"). Dye bias correction applied based on REgression on Logarithm of Internal Control probes (dyeCorr="RELIC"). No normalization applied. Probe design type bias correction applied using Regression on Correlated Probes (RCP) method. |
| 42 | enmix_neg_relic_nonorm_rcp | ENmix | Uses 600 chip internal controls probes to estimate background distribution parameters (bgParaEst="neg"). Dye bias correction applied based on REgression on Logarithm of Internal Control probes (dyeCorr="RELIC"). No normalization applied. Probe design type bias correction applied using Regression on Correlated Probes (RCP) method. |
| 43 | enmix_oob_nodye_nonorm_rcp | ENmix | Uses out-of-band Infinium I intensities to estimate normal distribution parameters to model background noise (bgParaEst="oob"). No dye bias correction applied (dyeCorr="none"). No normalization applied. Probe design type bias correction applied using Regression on Correlated Probes (RCP) method. |
| 44 | enmix_est_nodye_nonorm_rcp | ENmix | Uses combined methylated and unmethylated intensities to estimate background distribution parameters separately for each color channel and each probe type (bgParaEst="est"). No dye bias correction applied (dyeCorr="none"). No normalization applied. Probe design type bias correction applied using Regression on Correlated Probes (RCP) method. |
| 45 | enmix_neg_nodye_nonorm_rcp | ENmix | Uses 600 chip internal controls probes to estimate background distribution parameters (bgParaEst="neg"). No dye bias correction applied (dyeCorr="none"). No normalization applied. Probe design type bias correction applied using Regression on Correlated Probes (RCP) method. |
| 46 | enmix_oob_mean_q1_rcp | ENmix | Uses out-of-band Infinium I intensities to estimate normal distribution parameters to model background noise (bgParaEst="oob"). Dye bias correction applied based on averaged red/green ratio (dyeCorr="mean"). Quantile normalization applied separately for methylated and unmethylated intensities of Infinium I and II probes (method="quantile1"). Probe design type bias correction applied using Regression on Correlated Probes (RCP) method. |
| 47 | enmix_est_mean_q1_rcp | ENmix | Uses combined methylated and unmethylated intensities to estimate background distribution parameters separately for each color channel and each probe type (bgParaEst="est"). Dye bias correction applied based on averaged red/green ratio (dyeCorr="mean"). Quantile normalization applied separately for methylated and unmethylated intensities of Infinium I and II probes (method="quantile1"). Probe design type bias correction applied using Regression on Correlated Probes (RCP) method. |
| 48 | enmix_neg_mean_q1_rcp | ENmix | Uses 600 chip internal controls probes to estimate background distribution parameters (bgParaEst="neg"). Dye bias correction applied based on averaged red/green ratio (dyeCorr="mean"). Quantile normalization applied separately for methylated and unmethylated intensities of Infinium I and II probes (method="quantile1"). Probe design type bias correction applied using Regression on Correlated Probes (RCP) method. |
| 49 | enmix_oob_relic_q1_rcp | ENmix | Uses out-of-band Infinium I intensities to estimate normal distribution parameters to model background noise (bgParaEst="oob"). Dye bias correction applied based on REgression on Logarithm of Internal Control probes (dyeCorr="RELIC"). Quantile normalization applied separately for methylated and unmethylated intensities of Infinium I and II probes (method="quantile1"). Probe design type bias correction applied using Regression on Correlated Probes (RCP) method. |
| 50 | enmix_est_relic_q1_rcp | ENmix | Uses combined methylated and unmethylated intensities to estimate background distribution parameters separately for each color channel and each probe type (bgParaEst="est"). Dye bias correction applied based on REgression on Logarithm of Internal Control probes (dyeCorr="RELIC"). Quantile normalization applied separately for methylated and unmethylated intensities of Infinium I and II probes (method="quantile1"). Probe design type bias correction applied using Regression on Correlated Probes (RCP) method. |
| 51 | enmix_neg_relic_q1_rcp | ENmix | Uses 600 chip internal controls probes to estimate background distribution parameters (bgParaEst="neg"). Dye bias correction applied based on REgression on Logarithm of Internal Control probes (dyeCorr="RELIC"). Quantile normalization applied separately for methylated and unmethylated intensities of Infinium I and II probes (method="quantile1"). Probe design type bias correction applied using Regression on Correlated Probes (RCP) method. |
| 52 | enmix_oob_nodye_q1_rcp | ENmix | Uses out-of-band Infinium I intensities to estimate normal distribution parameters to model background noise (bgParaEst="oob"). No dye bias correction applied (dyeCorr="none"). Quantile normalization applied separately for methylated and unmethylated intensities of Infinium I and II probes (method="quantile1"). Probe design type bias correction applied using Regression on Correlated Probes (RCP) method. |
| 53 | enmix_est_nodye_q1_rcp | ENmix | Uses combined methylated and unmethylated intensities to estimate background distribution parameters separately for each color channel and each probe type (bgParaEst="est"). No dye bias correction applied (dyeCorr="none"). Quantile normalization applied separately for methylated and unmethylated intensities of Infinium I and II probes (method="quantile1"). Probe design type bias correction applied using Regression on Correlated Probes (RCP) method. |
| 54 | enmix_neg_nodye_q1_rcp | ENmix | Uses 600 chip internal controls probes to estimate background distribution parameters (bgParaEst="neg"). No dye bias correction applied (dyeCorr="none"). Quantile normalization applied separately for methylated and unmethylated intensities of Infinium I and II probes (method="quantile1"). Probe design type bias correction applied using Regression on Correlated Probes (RCP) method. |
| 55 | enmix_oob_mean_q2_rcp | ENmix | Uses out-of-band Infinium I intensities to estimate normal distribution parameters to model background noise (bgParaEst="oob"). Dye bias correction applied based on averaged red/green ratio (dyeCorr="mean"). Quantile normalization applied on combined methylated or unmethylated intensities for Infinium I or II probes (method="quantile2"). Probe design type bias correction applied using Regression on Correlated Probes (RCP) method. |
| 56 | enmix_est_mean_q2_rcp | ENmix | Uses combined methylated and unmethylated intensities to estimate background distribution parameters separately for each color channel and each probe type (bgParaEst="est"). Dye bias correction applied based on averaged red/green ratio (dyeCorr="mean"). Quantile normalization applied on combined methylated or unmethylated intensities for Infinium I or II probes (method="quantile2"). Probe design type bias correction applied using Regression on Correlated Probes (RCP) method. |
| 57 | enmix_neg_mean_q2_rcp | ENmix | Uses 600 chip internal controls probes to estimate background distribution parameters (bgParaEst="neg"). Dye bias correction applied based on averaged red/green ratio (dyeCorr="mean"). Quantile normalization applied on combined methylated or unmethylated intensities for Infinium I or II probes (method="quantile2"). Probe design type bias correction applied using Regression on Correlated Probes (RCP) method. |
| 58 | enmix_oob_relic_q2_rcp | ENmix | Uses out-of-band Infinium I intensities to estimate normal distribution parameters to model background noise (bgParaEst="oob"). Dye bias correction applied based on REgression on Logarithm of Internal Control probes (dyeCorr="RELIC"). Quantile normalization applied on combined methylated or unmethylated intensities for Infinium I or II probes (method="quantile2"). Probe design type bias correction applied using Regression on Correlated Probes (RCP) method. |
| 59 | enmix_est_relic_q2_rcp | ENmix | Uses combined methylated and unmethylated intensities to estimate background distribution parameters separately for each color channel and each probe type (bgParaEst="est"). Dye bias correction applied based on REgression on Logarithm of Internal Control probes (dyeCorr="RELIC"). Quantile normalization applied on combined methylated or unmethylated intensities for Infinium I or II probes (method="quantile2"). Probe design type bias correction applied using Regression on Correlated Probes (RCP) method. |
| 60 | enmix_neg_relic_q2_rcp | ENmix | Uses 600 chip internal controls probes to estimate background distribution parameters (bgParaEst="neg"). Dye bias correction applied based on REgression on Logarithm of Internal Control probes (dyeCorr="RELIC"). Quantile normalization applied on combined methylated or unmethylated intensities for Infinium I or II probes (method="quantile2"). Probe design type bias correction applied using Regression on Correlated Probes (RCP) method. |
| 61 | enmix_oob_nodye_q2_rcp | ENmix | Uses out-of-band Infinium I intensities to estimate normal distribution parameters to model background noise (bgParaEst="oob"). No dye bias correction applied (dyeCorr="none"). Quantile normalization applied on combined methylated or unmethylated intensities for Infinium I or II probes (method="quantile2"). Probe design type bias correction applied using Regression on Correlated Probes (RCP) method. |
| 62 | enmix_est_nodye_q2_rcp | ENmix | Uses combined methylated and unmethylated intensities to estimate background distribution parameters separately for each color channel and each probe type (bgParaEst="est"). No dye bias correction applied (dyeCorr="none"). Quantile normalization applied on combined methylated or unmethylated intensities for Infinium I or II probes (method="quantile2"). Probe design type bias correction applied using Regression on Correlated Probes (RCP) method. |
| 63 | enmix_neg_nodye_q2_rcp | ENmix | Uses 600 chip internal controls probes to estimate background distribution parameters (bgParaEst="neg"). No dye bias correction applied (dyeCorr="none"). Quantile normalization applied on combined methylated or unmethylated intensities for Infinium I or II probes (method="quantile2"). Probe design type bias correction applied using Regression on Correlated Probes (RCP) method. |
| 64 | enmix_oob_mean_q3_rcp | ENmix | Uses out-of-band Infinium I intensities to estimate normal distribution parameters to model background noise (bgParaEst="oob"). Dye bias correction applied based on averaged red/green ratio (dyeCorr="mean"). Quantile normalization applied on combined methylated or unmethylated intensities for Infinium I and II probes together (method="quantile3"). Probe design type bias correction applied using Regression on Correlated Probes (RCP) method. |
| 65 | enmix_est_mean_q3_rcp | ENmix | Uses combined methylated and unmethylated intensities to estimate background distribution parameters separately for each color channel and each probe type (bgParaEst="est"). Dye bias correction applied based on averaged red/green ratio (dyeCorr="mean"). Quantile normalization applied on combined methylated or unmethylated intensities for Infinium I and II probes together (method="quantile3"). Probe design type bias correction applied using Regression on Correlated Probes (RCP) method. |
| 66 | enmix_neg_mean_q3_rcp | ENmix | Uses 600 chip internal controls probes to estimate background distribution parameters (bgParaEst="neg"). Dye bias correction applied based on averaged red/green ratio (dyeCorr="mean"). Quantile normalization applied on combined methylated or unmethylated intensities for Infinium I and II probes together (method="quantile3"). Probe design type bias correction applied using Regression on Correlated Probes (RCP) method. |
| 67 | enmix_oob_relic_q3_rcp | ENmix | Uses out-of-band Infinium I intensities to estimate normal distribution parameters to model background noise (bgParaEst="oob"). Dye bias correction applied based on REgression on Logarithm of Internal Control probes (dyeCorr="RELIC"). Quantile normalization applied on combined methylated or unmethylated intensities for Infinium I and II probes together (method="quantile3"). Probe design type bias correction applied using Regression on Correlated Probes (RCP) method. |
| 68 | enmix_est_relic_q3_rcp | ENmix | Uses combined methylated and unmethylated intensities to estimate background distribution parameters separately for each color channel and each probe type (bgParaEst="est"). Dye bias correction applied based on REgression on Logarithm of Internal Control probes (dyeCorr="RELIC"). Quantile normalization applied on combined methylated or unmethylated intensities for Infinium I and II probes together (method="quantile3"). Probe design type bias correction applied using Regression on Correlated Probes (RCP) method. |
| 69 | enmix_neg_relic_q3_rcp | ENmix | Uses 600 chip internal controls probes to estimate background distribution parameters (bgParaEst="neg"). Dye bias correction applied based on REgression on Logarithm of Internal Control probes (dyeCorr="RELIC"). Quantile normalization applied on combined methylated or unmethylated intensities for Infinium I and II probes together (method="quantile3"). Probe design type bias correction applied using Regression on Correlated Probes (RCP) method. |
| 70 | enmix_oob_nodye_q3_rcp | ENmix | Uses out-of-band Infinium I intensities to estimate normal distribution parameters to model background noise (bgParaEst="oob"). No dye bias correction applied (dyeCorr="none"). Quantile normalization applied on combined methylated or unmethylated intensities for Infinium I and II probes together (method="quantile3"). Probe design type bias correction applied using Regression on Correlated Probes (RCP) method. |
| 71 | enmix_est_nodye_q3_rcp | ENmix | Uses combined methylated and unmethylated intensities to estimate background distribution parameters separately for each color channel and each probe type (bgParaEst="est"). No dye bias correction applied (dyeCorr="none"). Quantile normalization applied on combined methylated or unmethylated intensities for Infinium I and II probes together (method="quantile3"). Probe design type bias correction applied using Regression on Correlated Probes (RCP) method. |
| 72 | enmix_neg_nodye_q3_rcp | ENmix | Uses 600 chip internal controls probes to estimate background distribution parameters (bgParaEst="neg"). No dye bias correction applied (dyeCorr="none"). Quantile normalization applied on combined methylated or unmethylated intensities for Infinium I and II probes together (method="quantile3"). Probe design type bias correction applied using Regression on Correlated Probes (RCP) method. |
| 73 | watermelon_dasen | wateRmelon | Quantile normalizes methylated and unmethylated and type I and type II intensities separately. Same as the watermelon_nasen pipeline but type I and type II backgrounds are normalized first. |
| 74 | watermelon_naten | wateRmelon | Quantile normalizes methylated and unmethylated intensities separately. |
| 75 | watermelon_nanet | wateRmelon | Quantile normalizes methylated and unmethylated intensities together. |
| 76 | watermelon_nanes | wateRmelon | Quantile normalizes methylated and unmethylated intensities separately, except for type II probes where methylated and unmethylated are normalized together. |
| 77 | watermelon_danes | wateRmelon | Same as watermelon_nanes, except typeI and type II background are equalized first. |
| 78 | watermelon_danet | wateRmelon | Same as watermelon_nanet, except typeI and type II background are equalized first. |
| 79 | watermelon_danen | wateRmelon | Background equalization only, no normalization. |
| 80 | watermelon_daten1 | wateRmelon | Same as watermelon_naten, except typeI and type II background are equalized first (smoothed only for methylated intensities). |
| 81 | watermelon_daten2 | wateRmelon | Same as watermelon_naten, except typeI and type II background are equalized first (smoothed for both methylated and unmethylated intensities). |
| 82 | watermelon_nasen | wateRmelon | Same as watermelon_naten but typeI and typeII intensities quantile normalized separately. |
| 83 | watermelon_raw_BMIQ | wateRmelon | Using an intra-sample normalisation procedure, the bias of typeII intensity values is corrected. BMIQ uses a 3-step procedure: (i) fitting of a 3-state beta mixture model, (ii) transformation of state membership probabilities of typeII probes into quantiles of the typeI distribution, and (iii) a conformal transformation for the hemi-methylated probes. |
| 84 | watermelon_raw_PBC | wateRmelon | Peak correction only, no normalization. |
| 85 | minfi_raw | Minfi | Converts the Red/Green channel for an Illumina methylation array into methylation signal, without using any data processing or normalization. |
| 86 | minfi_illumina_bg_nonorm | Minfi | Implements Illumina GenomeStudio background correction. No control normalization applied. |
| 87 | minfi_illumina_bg_normcontrol | Minfi | Implements Illumina GenomeStudio background correction and control normalization. |
| 88 | minfi_illumina_nobg_normcontrol | Minfi | implements only Illumina GenomeStudio control normalization. No background correction applied. |
| 89 | minfi_noob_dyecorr | Minfi | Noob (normal-exponential out-of-band) background correction is applied with dye-bias normalization (dyeCorr=T,dyeMethod="single"). |
| 90 | minfi_noob_nodyecorr | Minfi | Noob (normal-exponential out-of-band) background correction is applied without dye-bias normalization (dyeCorr=F,dyeMethod="single"). |
| 91 | minfi_funnorm_nobg_nodyecorr | Minfi | Between array-normalization is applied that removes unwanted variation by regressing out variability explained by the control probes present on the array. No NOOB background correction and dye normalization applied before normalization (bgCorr=F,dyeCorr=F) |
| 92 | minfi_funnorm_bg_dyecorr | Minfi | Between array-normalization is applied that removes unwanted variation by regressing out variability explained by the control probes present on the array. NOOB background correction and dye normalization applied before normalization (bgCorr=T,dyeCorr=T) |
| 93 | minfi_funnorm_bg_nodyecorr | Minfi | Between array-normalization is applied that removes unwanted variation by regressing out variability explained by the control probes present on the array. Only NOOB background correction applied before normalization and no dye normalization (bgCorr=T,dyeCorr=F) |
| 94 | minfi_raw_quantile_strat | Minfi | Stratified quantile normalization was applied by region (CpG island, shore, etc.) |
| 95 | minfi_raw_quantile_nostrat | Minfi | Quantile normalization was applied with no stratification by region (CpG island, shore, etc.) |
| 96 | minfi_illumina_bg_quantile_strat | Minfi | Stratified quantile normalization was applied by region (CpG island, shore, etc.) after Illumina GenomeStudio background correction. |
| 97 | minfi_illumina_bg_quantile_nostrat | Minfi | Quantile normalization was applied with no stratification by region (CpG island, shore, etc.) after Illumina GenomeStudio background correction. |
| 98 | minfi_raw_SWAN | Minfi | Subset_quantile within array normalization (SWAN) was applied. It allows Infinium I and II type probes on a single array to be normalized together. |
| 99 | minfi_illumina_bg_SWAN | Minfi | Illumina GenomeStudio background correction was applied followed by subset quantile within array normalization (SWAN). |
| 100 | cross_noob_dyecorr_BMIQ | Minfi/  wateRmelon | Noob (normal-exponential out-of-band) background correction is applied with dye-bias normalization (dyeCorr=T,dyeMethod="single") followed by subset quantile within array normalization (SWAN). |
| 101 | cross_noob_nodyecorr_BMIQ | Minfi/  wateRmelon | Noob (normal-exponential out-of-band) background correction is applied without dye-bias normalization (dyeCorr=F,dyeMethod="single") followed by subset quantile within array normalization (SWAN). |
